# Chemical Synthesis of Oligosaccharides Derived from *Streptococcus pneumoniae* Serotype 35B and D to Investigate Binding of Complement and Serum Factors

**DOI:** 10.1101/2025.01.28.635287

**Authors:** Ivan A. Gagarinov, Lin Liu, Geert-Jan Boons

## Abstract

The wide use of *Streptococcus pneumoniae* capsular polysaccharide (CPS) conjugate vaccines is causing serotype replacement and the emergence of serotype 35B is concerning because of its multidrug resistance. CPS of 35B is composed of pentasaccharide repeating units that are linked through phosphodiester linkages. One of the galactofuranose residues of the pentasaccharide is acetylated, which distinguishes it from the invasive serotype 35D lacking the acetyl ester. Here, we describe a synthetic approach that can provide oligosaccharides derived of CPS 35B and 35D composed of up to 15 monosaccharides using a pentasaccharide building block equipped with four orthogonal protecting groups. The synthetic compounds were used to examine binding properties of ficolin-2, which is a protein that can activate the lectin pathway of the complement system. It was found that *O*-acetylation is essential for recognition by ficolin-2 requiring at least two repeating units. The data provides a rationale why 35D may escape immune detection and be more invasive. The synthetic oligosaccharides were also investigated for binding to pneumococcal serum factor 35a and 29b, which are employed for serotype identification. It indicates that immunization with 35B CPS will not provide protection against 35D and thus inclusion of 35B in vaccines may result in serotype replacement by 35D. On the other hand, antibodies that can bind 35D can recognize 35B and thus 35D CPS may provide cross protection. Our findings have direct implementation for the development of the next generation pneumococcal vaccine and provide an understanding of disease severity by the emerging serotype 35B and 35D.

## INTRODUCTION

*Streptococcus pneumoniae* are gram-positive bacteria that are a common cause of pneumonia and can cause other illnesses including severe ear infection, meningitis, and bacteraemia,^1^ which can result in long-lasting problems such as hearing loss, brain damage, and even death. The most effective way to combat pneumococcal disease is by preventative vaccination. More than 90 serologically distinct pneumococcal capsular polysaccharides (serotypes) have been identified. Antibodies against the capsule are protective and therefore polysaccharide conjugate vaccines have been developed against the most prevalent serotypes, which has resulted in a great reduction in pneumococcal disease.^2^ The most widely employed vaccine, which is known as PCV13, is composed of capsular polysaccharides (CPS) conjugated to carrier proteins and provide protection against 13 serotypes that commonly cause pneumococcal disease. The wide use of PCV13, and the earlier PCV7 vaccine, has caused other serotypes to become more prevalent.^3^ This so-called antigenic shift is favouring the spread of invasive nonvaccine serotypes such as 12F, 15B and 15C (15B/C), and 35B/D. Serotype 35B is of particular concern because it is associated with high rates of multidrug resistance.^4-6^ The emergence of new serotypes has led to the recent introduction of a twenty and twenty one valent vaccine in which the capsular polysaccharides are conjugated to the carrier protein CRM_197_.

CPS of 35B is a high molecular weight polymer composed of D-galactofuranose, D-glucose, *N*-acetyl-D-galactosamine, and ribitol (Figure 1).^7^ These monosaccharides are *trans*-linked through glycosidic linkages to give a pentasaccharide repeating unit that is further polymerized through phosphodiesters. One of the galactofuranose residues is modified at C-2 by an acetyl ester. This feature distinguishes 35B from closely related and invasive serotype 35D that lacks the acetyl ester.^8^

**Figure 1.**
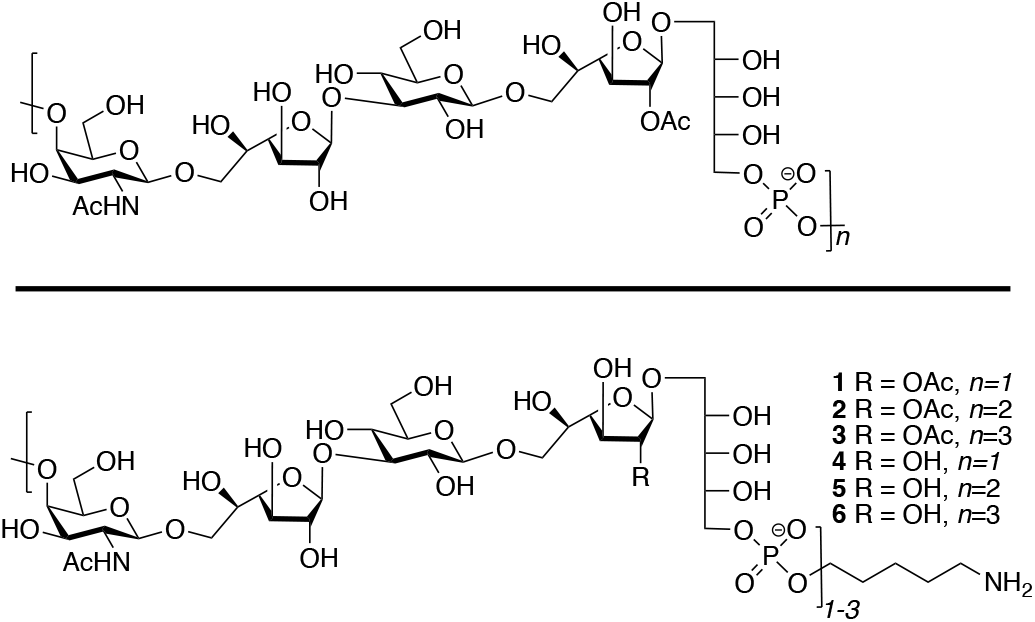
Structure of *S. pneumoniae* 35B CPS and well-defined linker-equipped target oligosaccharides.

*O*-acetylation is important for functional immunity for a number of polysaccharides such as meningococcal serogroup A^9^ and *Salmonella typhi* V.^10^ Furthermore, *O*-acetylation can be important for innate immune detection and for example *O*-acetylated capsular polysaccharides can be recognized by the serum protein ficolin-2, which is an initiator of the lectin complement pathway.^11^ Ficolin-2 mediated complement deposition has been observed for serotype 11A and 35B but not for *O*-acetyl transferase deficient derivatives of these serotypes. The ability of ficolin-2 to recognize serotype 11A and 35B capsular polysaccharides has been associated with low invasiveness in children.^12^ The elderly are, however, susceptible to invasive pneumococcal diseases by these serotypes and it has been proposed that this is due to reduced ficolin-2-mediated immunity. Thus, the elderly is an important focus group for vaccination by *S. pneumonia* serotype 35B. Furthermore, serotype 35D, which lacks the C-2 acetyl ester at the galactofuranoside, is more invasive^13, 14^ which may be due to a lack of recognition by ficolin-2.

Ficolin 1, 2, and 3, which are also known as ficolin-M, -L, and -H, are human serum-associated pattern-recognition receptors that structurally and functionally are similar to mannose-binding lectin (MBL).^11^ Ficolins can form complexes with MBL-associated serine proteases and, upon binding, initiate the lectin complement pathway resulting in opsonophagocytosis. Ficolins have a collagen-like domain that is important for oligomerization and fibrinogen-like domain that can recognize specific carbohydrate-motifs. The three human ficolins have different ligand requirements and can distinguish self-from non-self. Ficolin-2 can recognize a diverse range of carbohydrates, including acetylated *N*- and *O*-structures, as well neutral sugars polysaccharides. The ligand requirements of ficolins have mainly been studied by genetic approaches, and for example the importance of *O*-acetylation of CPS of *S. pneumonia* serotypes 11A and 35B has been determined by mutants lacking *O*-acetyl transferases.^12^ A series of biological relevant oligosaccharides is required to dissect how ficolin-2 selectively recognize microbial polysaccharide for complement mediated neutralization.

Here, we report the chemical synthesis of well-defined oligomers derived from CPS 35B. By saponification of the acetyl esters, compounds derived from serotype 35D were easily obtained. Binding studies by glycan microarray technology demonstrated that *O*-acetylation is essential for recognition by ficolin-2. At least two repeating units are required for binding and furthermore a trimer showed a much greater responsiveness. The data provides a rationale why the closely related serotype 35D might escape immune detection and be more invasive.^13^ The synthetic oligosaccharides were also investigated for binding to pneumococcal serum factor 35a and 29b, which are employed for serotype identification. Factor sera 35a is reported to only bind type serotype 35B and as expected it only recognized *O*-acetylated oligosaccharides in a length dependant manner. Factor sera 29b is reported to recognize both 35B and 35D and bind to all synthetic compounds. The findings have direct implementation for the development of the next generation pneumococcal vaccine and provide an understanding of disease severity by the emerging serotype 35B and 35D.

## RESULTS AND DISCUSSION

### Chemical Synthesis of *S. pneumoniae* 35B Oligosaccharides

The chemical synthesis of the pentasaccharide repeating unit of *S. pneumoniae* 35B and its oligomers has not been reported yet. We envisaged a scalable synthetic strategy that can provide oligosaccharides **1**–**3** (Figure 1) that are composed of one, two, and three repeating units of the CPS of *S. pneumoniae* 35B. The compounds are equipped with an artificial aminopentyl linker to facilitate bioconjugation and microarray printing. Our strategy employs glycosyl acceptor **7** and thioglycosyl donors **8**-**11** to assemble key oligosaccharide **12** (Figure 2). The latter compound is equipped with the four orthogonal protecting groups, *t-*butyldiphenylsilyl (TBDPS), fluorenylmethyl (Fmoc), levulinolyl (Lev), and 2,2,2-trichloroethylcarbonyl (NHTroc), which allow oligomer assembly and deprotection without affecting the critical acetyl esters. The TBDPS and Fmoc protecting group of **12** are at sites where a phosphodiester needs to be installed and can independently be cleaved by HF-pyridine and triethylamine, respectively.^15^ Removal of the TBDPS ether of **12** will give an alcohol that can be modified as an *H*-phosphonate to provide building block **13** that was linked to a properly protected aminopentyl spacer. Alternatively, the Fmoc protecting group of **12** can be removed by a hindered base to give an alcohol that can be coupled with *H*-phosphonate **13** to give, after oxidation with iodine, a phosphodiester. The process of Fmoc removal and coupling with **13** can be repeated to give larger oligomers.

**Figure 2.**
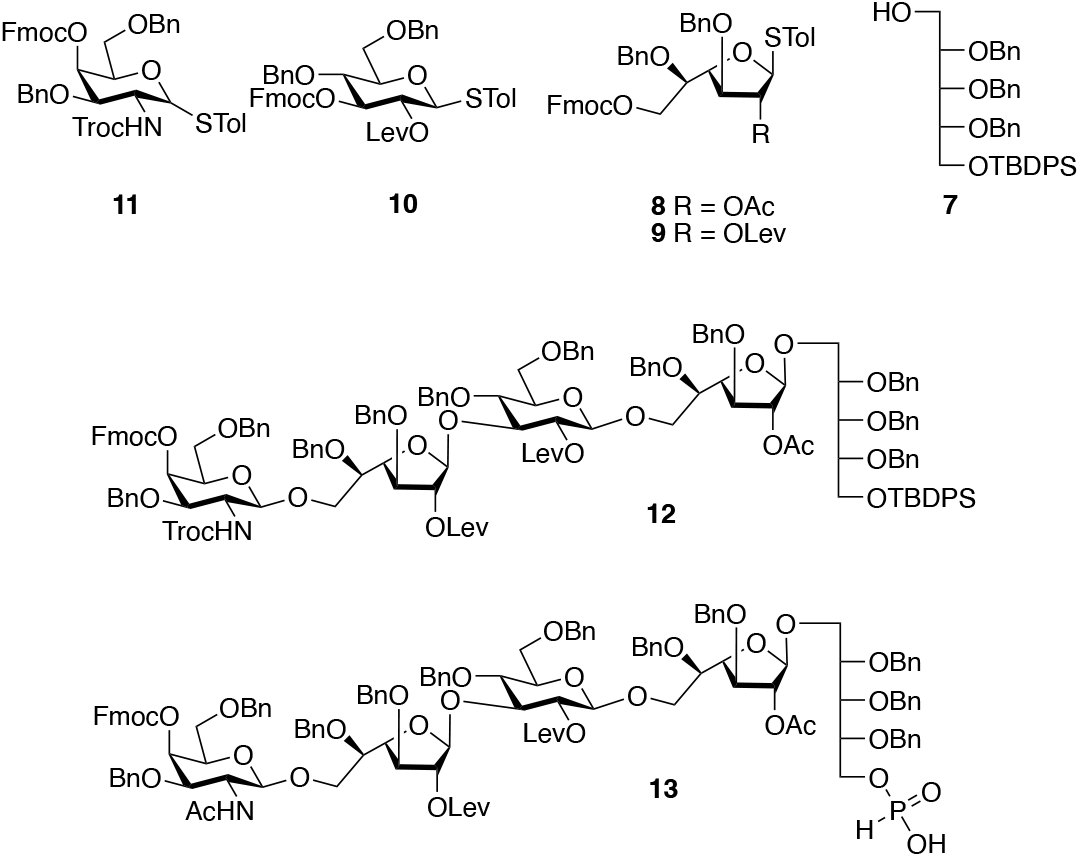
Key building blocks required for the assembly of *S. pneumoniae* 35B/D.

The building blocks **8** – **11** were strategically selected for efficient oligosaccharide assembly and final deprotection. The anomeric STol^16^ of donors **8** – **11** ensures shelf stability yet allow highly efficient glycosylations using NIS/TMSOTf as the promotor system.^17^ All donors are protected by Fmoc at sites where a subsequent glycosylation needs to be performed and thus only one set of reaction conditions is needed for glycosylation and acceptor generation opening possibilities for automated synthesis.^16^ The Lev esters of **9** and **10** ensure that the glucoside and non-acetylated Gal*f* moiety are selectively installed as *β*-anomers due to neighbouring group participation.^18^ Building block **8** is used to install the Gal*f* moiety carrying the critical acetyl ester. At the final deprotection stage, the Lev esters can be cleaved by hydrazine acetate and these conditions preserve the base sensitive acetyl esters. Finally, donor **11** is derivatized with a NHTroc at C2 to ensure selective *β*-glycoside formation and this group can be selectively converted into a native acetamido moiety by treatment with Zn powder without affecting other parts of the oligosaccharide.

Ribitol acceptor **7** was prepared in a large quantity (∼45.0 g) in four steps from crystalline ribose dithioacetal (Scheme S1).^19^ Previous syntheses of ribitol acceptors are lengthy^20^ and not scalable. Glycosyl donors **8**-**11** could also be prepared on large scales and details are provided in Schemes S2-S4. An NIS/TMSOTf-catalyzed^21^ glycosylation of **7** with **8** furnished disaccharide **14** in 95% yield and as expected only the β-anomer was formed due to neighbouring group participation of the acetyl ester (Scheme 1). Et_3_N-mediated cleavage of the Fmoc protecting group liberated a hydroxyl to give acceptor **15** that was coupled with the glucosyl donor **10** to provide trisaccharide **16** in a yield of 72%. Next, the Fmoc protecting group of **16** was removed using standard conditions to afford acceptor **17** that was coupled with the Gal*f* donor **9** using NIS/TMSOTf as the promotor to provide tetrasaccharide **18** in 86% yield. The Fmoc protecting group of **18** was cleaved to give acceptor **19**, which was further coupled with glycosyl donor **11** to provide **12** in a yield of 91%. Interestingly, **12** was obtained in a much lower yield when a trichloroacetimidate^22^ donor was used. The synthetic route made it easy to obtain pentasaccharide **12** in a large quantity (15.7 g). The results demonstrate that Fmoc is an attractive temporary protecting group that can repeatedly be cleaved without affecting the other functionalities.

**Scheme 1.**
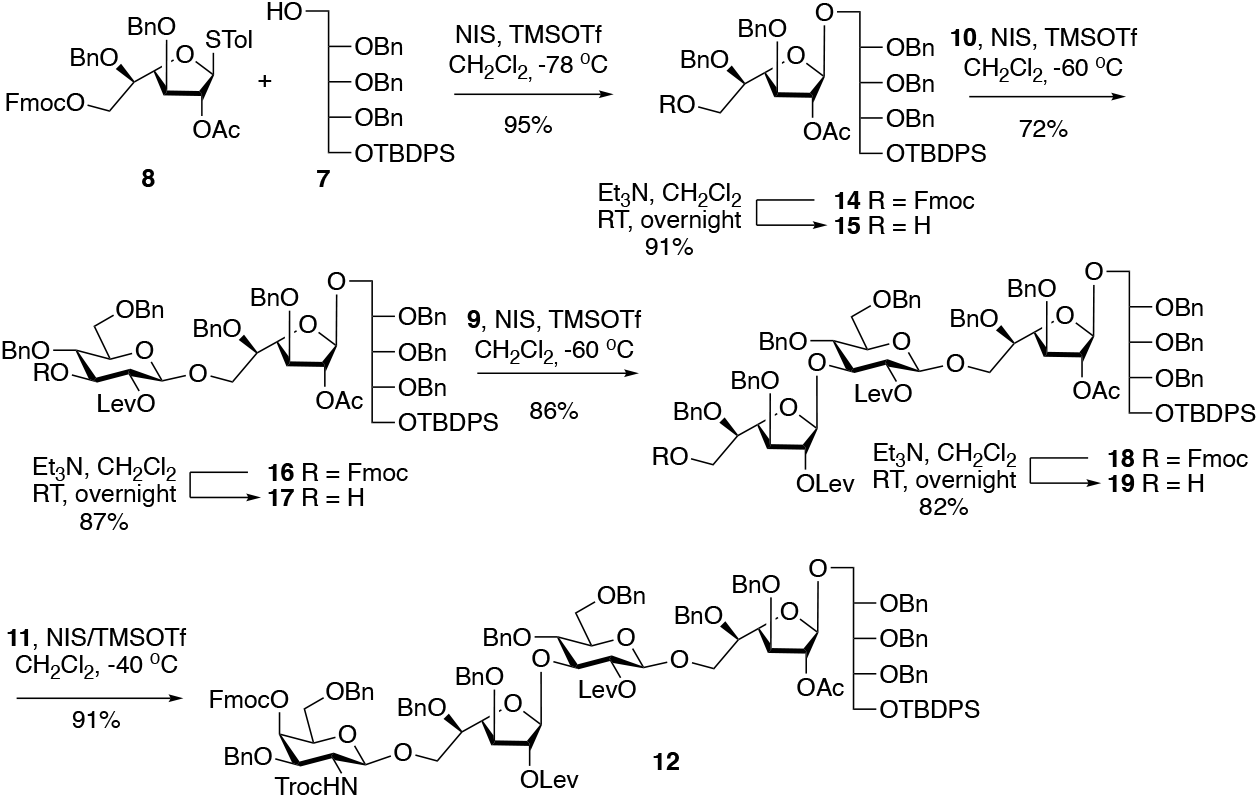
Synthesis of the core pentasaccharide **12**.

The flexibility of building block **12** was demonstrated by performing a phosphitylation followed by linker attachment and deprotection to give compound **1** (Scheme 2). Thus, the Troc carbamate of **12** was converted into an acetamido moiety by treatment with Zn dust in a solution of THF/AcOH/Ac_2_O^17^ to afford **20** in high yield. Next, the TBDPS ether of **20** was cleaved using a hydrogen fluoride–pyridine complex to provide alcohol **21** in a quantitative yield. Subsequent installation of the crucial *H*-phosphonate group was accomplished by using salicyl chlorophosphite^23^ to give **13** in a yield of 89%. Coupling *N*-(benzyl)benzyloxycarbonyl-protected aminopentanol^24^ and **13** using pivaloyl chloride (PivCl) as activator, followed by *in situ* oxidation with iodine in pyridine/water and subsequent removal of the Fmoc group generated fragment **22** in 88% yield over two steps.^20^ Finally, the Lev esters were selectively cleaved using hydrazine acetate in a mixture of CH_2_Cl_2_ and CH_3_OH,^18^ and the remaining benzyl ethers were by hydrogenation over palladium hydroxide (Degussa type) ^18^ at ambient pressure to furnish the first desired target compound **1**.

**Scheme 2.**
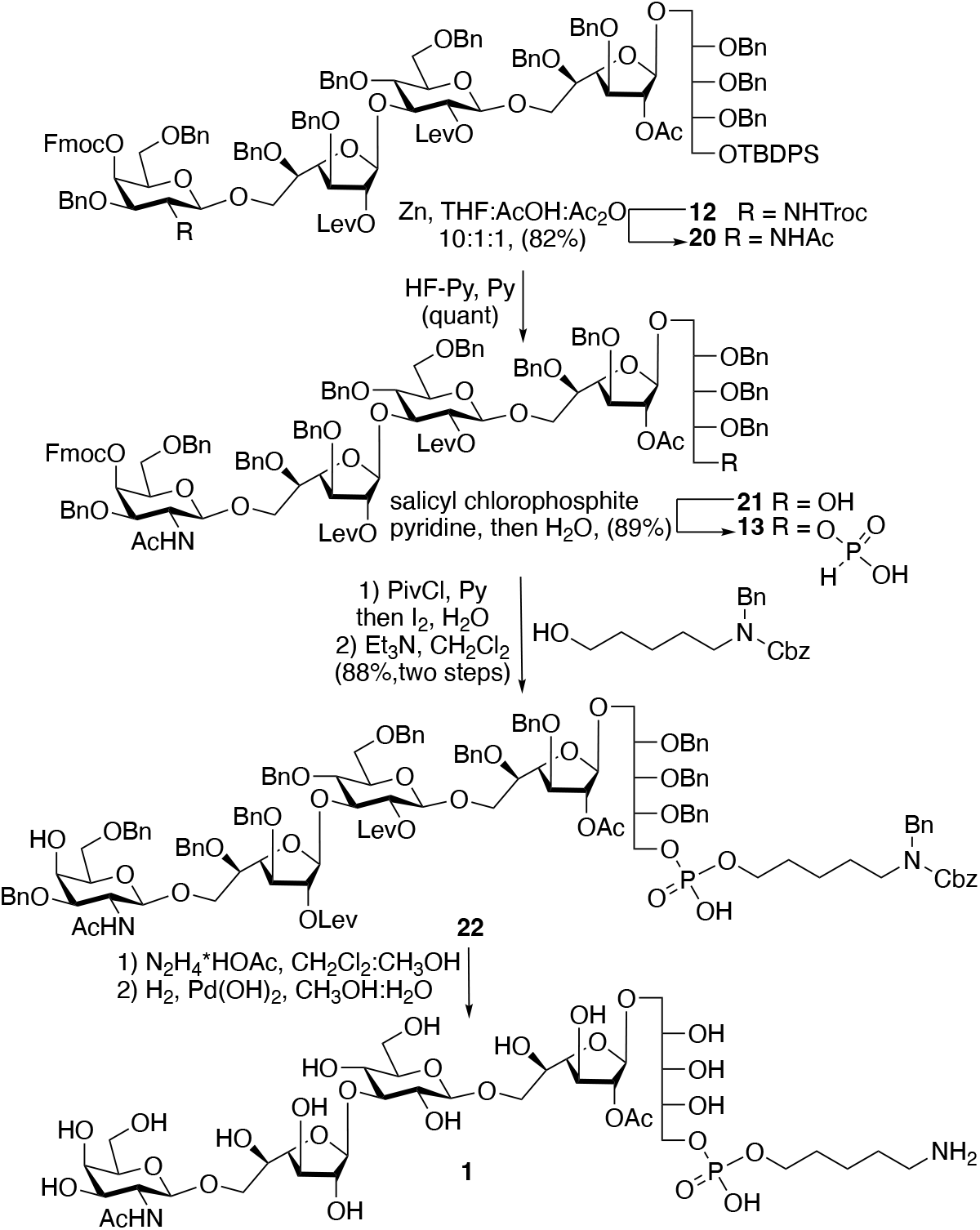
Synthesis of key phosphonate **13** and its further functionalization with a C5 linker to give **1**.

The versatility of phosphonate **13** was showcased by synthesizing deca- and pentadecasaccharides **2** and **3**, respectively (Scheme 3). A 5 + 5 block coupling of **22** and **13** using PivCl as activator afforded **23** in 46% yield after purification by silica gel chromatography. Remaining starting material **22** could only be removed from the desired **23** by incorporation of acetonitrile into eluent system, which is unusual for standard silica gel columns (see SI). Subsequent removal of the Fmoc group gave alcohol **24** which was further treated with hydrazine acetate to remove the Lev esters, and then subjected to catalytic hydrogenation over Pd(OH)_2_ to remove all 24 benzyl ethers to provide decasaccharide **2** in an overall yield of 88%. Pentadecasaccharide **3** was synthesized according to the above methodology, albeit the coupling yield between **24** and **13** was lower (41%), which was attributed to low nucleophilicity of C-4 hydroxyl of GalNAc (Scheme 3).

**Scheme 3.**
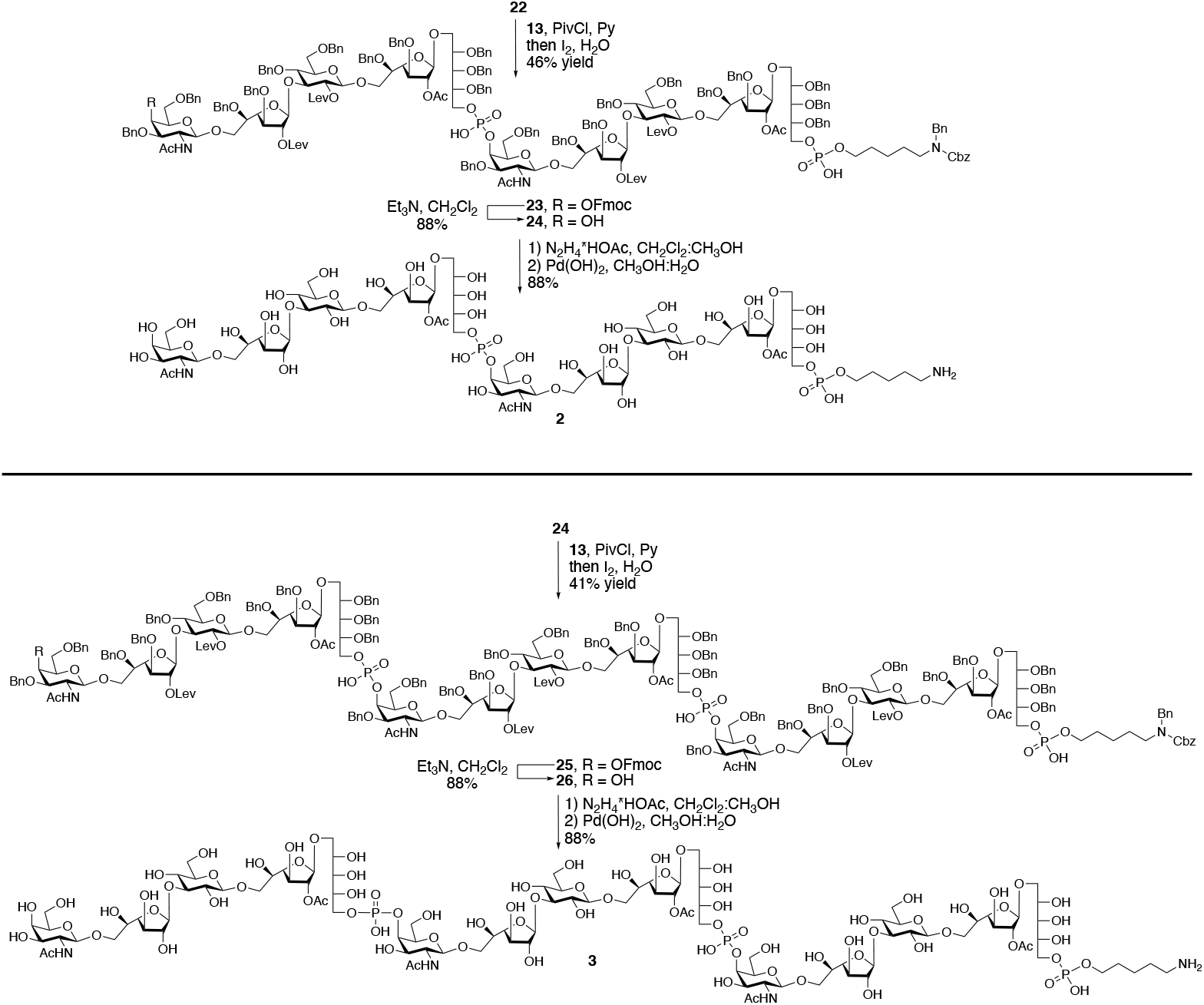
Synthesis of decasaccharide **2** and pentadecasaccharide **3**.

### Microarray Development and Binding Studies with Ficolins and Factor Antisera

Next, attention was focused on developing a microarray to investigate binding with ficolins and antisera. First, samples of compounds **1 – 3** were incubated with 100 mM NaOH (pH 11.0) at 40 ^°^C for 2 h to effect de-*O*-acetylation to give compounds **4 – 6**. Next, the oligosaccharides were dissolved in a printing buffer (pH 8.5) and exposed to *N*-hydroxysuccinimide (NHS)-activated glass slides at a concentration of 100 *μ*M in replicates of 10. The acetyl esters were stable under these conditions.

The slides were incubated with different concentrations of recombinant ficolin-2 having a C-terminal His-10 tag (2, 20, and 100 *μ*g/mL) for 1 h, followed by washing and re-incubation with a AlexaFluor®647 conjugated anti-His antibody to detect binding. The results uncovered that the length of the CPS fragments together with *O*-acetylation is important for binding to ficolin-2 (Figure 3, left). Strong binding of ficolin-2 was observed for *O*-acetylated trimer **3** whereas no binding was detected for *O*-acetylated monomer **1** and a weak response for *O*-acetylated dimer **2**. Removal of the acetyl esters abolished binding. Furthermore, compounds **1–6** exhibited no binding to ficolin-1 (ficolin-M). The data indicate that ficolin-2 has complex binding requirements, which are not only dependent on *O*-acetylation but also the size of the ligand. A loss of *O*-acetylation of CPS is probably a mechanism of immune evasion, which has recently been reflected as a microevolution of serotype 35B into a genetically similar serotype 35D.^13^ The serotype 35D capsule is identical to that of serotype 35B except for the absence of acetyl esters at one of the galactofuranose residues. *S. pneumoniae* has a remarkable ability to adapt and for example there is an increase in reported global occurrence of 35D infections, in young children.^14^ Our data underline the invasive potential conferred by the loss of *O*-acetylation of the pneumococcal capsule by loss of detection by ficolin-2.^25^

**Figure 3.**
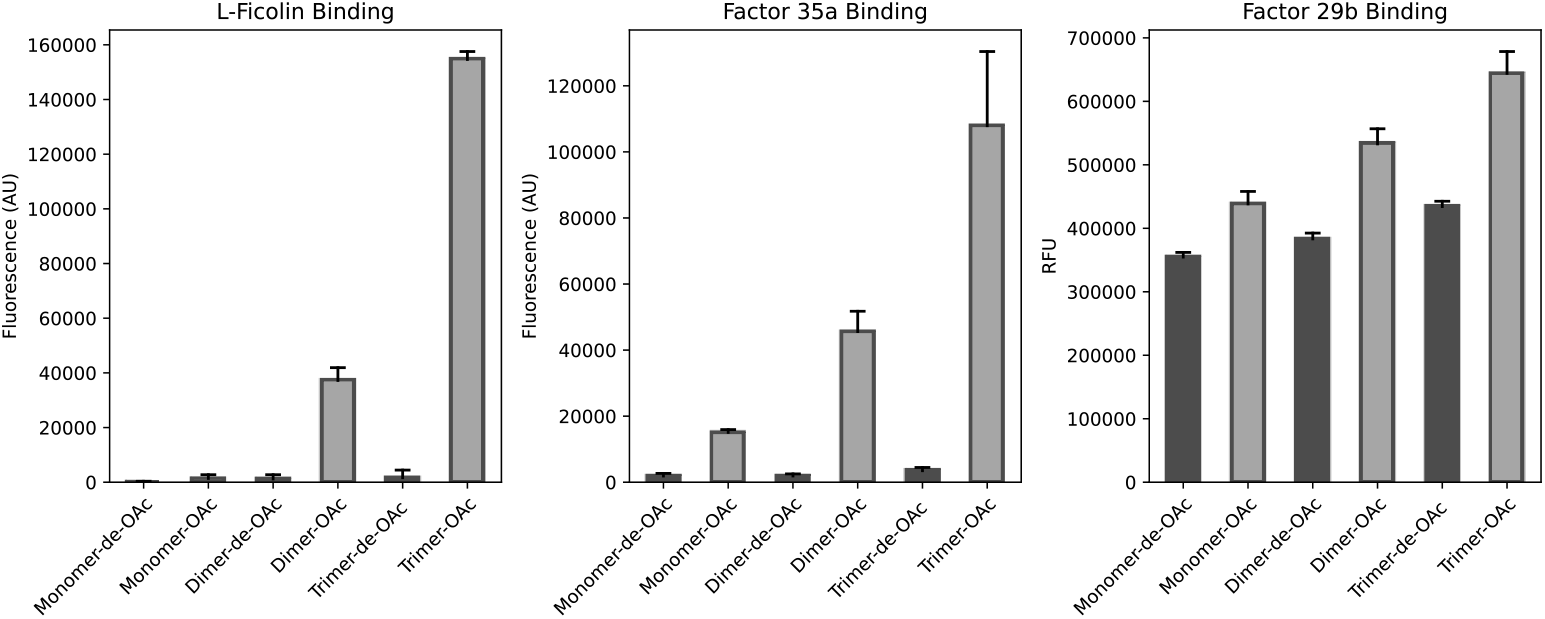
Results of microarray binding studies of synthetic oligosaccharides **1 – 6** with L-ficolin (left), factor 35a (middle), and factor 29b (right).

To further demonstrate the importance of previously neglected *S. pneumoniae* CPS *O*-acetylation, another set of microarray binding experiments was performed with serum factor 35a and 29b.^26^ Factor sera bind to specific serotypes and is used to perform detailed serotype identification. Factor sera 29b is reported to recognize both 35B and 35D, while factor sera 35a only recognize type 35B.^13^ Thus, the slides were incubated with the factor-sera 29b and 35a at different dilutions, followed by washing and re-incubation with a premixed solution of biotinylated Goat-anti-Rabbit antibody and Streptavidin-AlexaFluor®647 conjugate for detection. The data showed that factor 35a only recognizes the *O*-acetylated oligosaccharides derived from CPS of *S. pneumoniae* 35B and thus the acetyl esters are a critical part of the recognition epitope (Figure 3, middle). Length was also an important for binding and an increase in oligosaccharide length resulted in better responsiveness. Factor 29b bound to both acetylated 35B and the non-acetylated oligosaccharides (Figure 3, right), and in this case the length of the oligosaccharides did not impact binding.

## CONCLUSIONS

A scalable synthetic route for oligosaccharides composed of multiple repeating units derived from the CPS of *S. pneumoniae* 35B is described which is an emerging serotype and its immuno. Our modular synthetic approach provided compounds composed of up to 15 monosaccharides in length using a key pentasaccharide building block equipped with four orthogonal protecting groups. Careful selection of the protecting groups was important to preserve the biologically important *O*-acetyl esters and phosphodiester linkages. The synthetic compounds made it possible to examine binding properties of ficolin-2, which is one of the few proteins that can activate the lectin pathway of the complement system.^27^ There are indications that ficolin-2 can recognize a diverse range of microbial *O*- and *N*-acetylated carbohydrate structures as well polysaccharides such as 1,3-β-glucan.^11^ Ficolin-2 mediated complement deposition has been observed for serotype 11A and 35B but not variants deficient in the *O*-acetyl transferase.^12^ Furthermore, it has been shown that ficolin-A-deficient knockout mice exhibited reduced survival rates following infection with *S. pneumonia* compared to the wild-type mice.^28^ Direct binding of ficolin-2 to CPS of serotype 35B has not been demonstrated and furthermore molecular mechanisms by which ficolin-2 distinguishes self from non-self are not well understood. Multiple binding sites have been identified within ficolin-2, and the S3 site, which includes Arg132, Asp133, Thr136, and Lys221, is the putative binding pocket for acetylated saccharides.^29^ Ficolins occur as oligomers due to a collagen-like region, and thus it is likely that these immune receptors bind to saccharides that present multiple binding partners.^30^ Our data indicates that a single repeating unit of 35B CPS, even when presented on a surface, is not sufficient for ficolin-2 binding. On the other hand, it was found that a compound composed of three repeating units (**3**) bound avidly to ficolin-2 at a concentration that mimics its present in serum. Furthermore, *O*-acetylation was critical for binding and oligosaccharides derived for CPS of serotype 35D that is devoid of acetylation was not recognized. Despite the latter compounds contain GlcNAc moieties, which have been identified as ligand for ficolin-2, they did not exhibit binding. Thus, it a multivalent display of *O*-acetylated epitopes as part of a polysaccharides may be important for detection by ficolin-2. Despite immune detection by ficolin-2, bacteria may have an advantage by *O*-acetylation of their capsular polysaccharides and for example loss of WciG-mediated *O*-acetylation of serotypes 33A and 33F resulted in a less stable capsule with an increase in cell wall accessibility, increased nonspecific opsonophagocytic killing, enhanced biofilm formation, and increased adhesion to nasopharyngeal cells.^31^

Binding studies with antisera indicated that the acetyl ester of the D-galactofuranose moiety of 35B CPS is critical for binding and appears to be an immune-dominant feature. The results indicate that immunization with 35B CPS will not provide protection against 35D and thus inclusion of 35B CPS serotype in a vaccine may result serotype replacement by 35D. Interestingly, antibodies that can bind 35D can also recognize 35B and thus 35D CPS may provide cross protection, but this needs further investigation.

It is a challenge to preserve acetyl esters during industrial scale production of conjugate vaccines. The synthetic antigens described here are modified by an aminopentyl linker that will facilitate controlled coupling to carrier proteins such as CRM_197_ and will facilitate conjugate vaccine development. The synthetic strategy described here is also relevant to the preparation of other emerging serotypes of *S. pneumoniae* that share structural features such as repeating units linked by phosphodiester bonds and acetyl esters.

## ASSOCIATED CONTENT

### Supporting Information

The Supporting Information is available free of charge at https://pubs.cs.org/doi/ Methods, analytical data, and additional figures (PDF).

## AUTHOR INFORMATION

### Authors

**Ivan A. Gagarinov -** Chemical Biology and Drug Discovery, Utrecht Institute for Pharmaceutical Sciences, Utrecht University, Utrecht 3584 CG, The Netherlands. Present address: Institute for Biomedicine and Glycomics, Griffith University, Gold Coast, QLD 4222, Australia; orcid.org/0000-0003-1742-3870.

**Lin Liu** - Complex Carbohydrate Research Center, University of Georgia, Athens, Georgia 30602, United States.

## ACKNOWLEDGMENTS

This research was supported by NIH (Grants P41GM103390 and HLBI R01HL151617 to G.J.B.).

## References

(1) O’Brien, K. L.; Wolfson, L. J.; Watt, J. P.; Henkle, E.; Deloria-Knoll, M.; McCall, N.; Lee, E.; Mulholland, K.; Levine, O. S.; Cherian, T. Burden of Disease Caused by Streptococcus pneumoniae in Children Younger than 5 years: Global Estimates. Lancet 2009, 374 (9693), 893–902.

(2) Johnson, H. L.; Deloria-Knoll, M.; Levine, O. S.; Stoszek, S. K.; Freimanis Hance, L.; Reithinger, R.; Muenz, L. R.; O’Brien, K. L. Systematic Evaluation of Serotypes Causing Invasive Pneumococcal Disease among Children Under Five: The Pneumococcal Global Serotype Project. PLoS Med. 2010, 7 (10), e1000348.

(3) Domenech, M.; Damián, D.; Ardanuy, C.; Liñares, J.; Fenoll, A.; García, E. Emerging, Non-PCV13 Serotypes 11A and 35B of Streptococcus pneumoniae Show High Potential for Biofilm Formation In Vitro. PLoS One 2015, 10 (4), e0125636.

(4) Liset, O.; Sheldon, L. K.; William, J. B.; José, R. R.; Philana Ling, L.; Tina, Q. T.; Jill, A. H.; John, S. B.; Laurence, B. G.; Edward, O. M.; et al. Invasive Serotype 35B Pneumococci Including an Expanding Serotype Switch Lineage. Emerg. Infect. Dis. 2018, 24 (2), 405.

(5) Olarte, L.; Kaplan, S. L.; Barson, W. J.; Romero, J. R.; Lin, P. L.; Tan, T. Q.; Hoffman, J. A.; Bradley, J. S.; Givner, L. B.; Mason, E. O.; et al. Emergence of Multidrug-Resistant Pneumococcal Serotype 35B among Children in the United States. J. Clin. Microbiol. 2017, 55 (3), 724–734.

(6) Beall, B.; McEllistrem, M. C.; Gertz, R. E., Jr.; Boxrud, D. J.; Besser, J. M.; Harrison, L. H.; Jorgensen, J. H.; Whitney, C. G. Emergence of a Novel Penicillin-Nonsusceptible, Invasive Serotype 35B Clone of Streptococcus pneumoniae within the United States. J. Infect. Dis. 2002, 186 (1), 118–122.

(7) Beynon, L. M.; Richards, J. C.; Perry, M. B.; Kniskern, P. J. Characterization of the Capsular Antigen of Streptococcus pneumoniae Serotype 35B. Can. J. Chem. 1995, 73 (1), 41–48.

(8) Staples, M.; Graham, R. M. A.; Hicks, V.; Strachan, J.; da Silva, A. G.; Peverall, J.; Wicks, V.; Jennison, A. V. Discovery of Serogroup 35 Variants in Australian Patients. Clin. Microbiol. Infec. 2017, 23 (7), 476–479.

(9) Fiebig, T.; Freiberger, F.; Pinto, V.; Romano, M. R.; Black, A.; Litschko, C.; Bethe, A.; Yashunsky, D.; Adamo, R.; Nikolaev, A.; et al. Molecular Cloning and Functional Characterization of Components of the Capsule Biosynthesis Complex of Neisseria meningitidis Serogroup A: Toward In Vitro Vaccine Production. J. Biol. Chem. 2014, 289 (28), 19395–19407.

(10) Jarvis, F. G.; Mesenko, M. T.; Martin, D. G.; Perrine, T. D. Physicochemical Properties of the Vi Antigen Before and After Mild Alkaline Hydrolysis. J. Bacteriol. 1967, 94 (5), 1406.

(11) Ma, Y. J.; Lee, B. L.; Garred, P. An Overview of the Synergy and Crosstalk between Pentraxins and Collectins/Ficolins: Their Functional Relevance in Complement Activation. Exp. Mol. Med. 2017, 49.

(12) Geno, K. A.; Spencer, B. L.; Bae, S.; Nahm, M. H. Ficolin-2 Binds to Serotype 35B Pneumococcus as it does to Serotypes 11A and 31, and these Serotypes Cause More Infections in Older Adults than in Children. PLoS One 2018, 13 (12).

(13) Geno, K. A.; Saad, J. S.; Nahm, M. H. Discovery of Novel Pneumococcal Serotype 35D, a Natural WciG-Deficient Variant of Serotype 35B. J. Clin. Microbiol. 2017, 55 (5), 1416.

(14) Lo, S. W.; Gladstone, R. A.; van Tonder, A. J.; Hawkins, P. A.; Kwambana-Adams, B.; Cornic, J. E.; Madhi, S. A.; Nzenze, S. A.; du Plessis, M.; Kandasamy, R.; et al. Global Distribution of Invasive Serotype 35D Isolates following Introduction of 13-Valent Pneumococcal Conjugate Vaccine. J. Clin. Microbiol. 2018, 56 (7).

(15) Arungundram, S.; Al-Mafraji, K.; Asong, J.; Leach, F. E.; Amster, I. J.; Venot, A.; Turnbull, J. E.; Boons, G.-J. Modular Synthesis of Heparan Sulfate Oligosaccharides for Structure−Activity Relationship Studies. J. Am. Chem. Soc. 2009, 131 (47), 17394–17405.

(16) Hsu, C.-H.; Hung, S.-C.; Wu, C.-Y.; Wong, C.-H. Toward Automated Oligosaccharide Synthesis. Angew. Chem. Int. Ed. 2011, 50 (50), 11872–11923.

(17) Gagarinov, I. A.; Fang, T.; Liu, L.; Srivastava, A. D.; Boons, G.-J. Synthesis of Staphylococcus aureus Type 5 Trisaccharide Repeating Unit: Solving the Problem of Lactamization. Org. Lett. 2015, 17 (4), 928–931.

(18) Gagarinov, I. A.; Li, T.; Toraño, J. S.; Caval, T.; Srivastava, A. D.; Kruijtzer, J. A. W.; Heck, A. J. R.; Boons, G.-J. Chemoenzymatic Approach for the Preparation of Asymmetric Bi-, Tri-, and Tetra-Antennary N-Glycans from a Common Precursor. J. Am. Chem. Soc. 2017, 139 (2), 1011–1018.

(19) Roberts, J. C.; Nagasawa, H. T.; Zera, R. T.; Fricke, R. F.; Goon, D. J. W. Prodrugs of L-Cysteine as Protective Agents against Acetaminophen-induced Hepatotoxicity. 2-(Polyhydroxyalkyl)- and 2-(polyacetoxyalkyl)thiazolidine-4(R)-carboxylic acids. J. Med. Chem. 1987, 30 (10), 1891–1896.

(20) Baek, J. Y.; Geissner, A.; Rathwell, D. C. K.; Meierhofer, D.; Pereira, C. L.; Seeberger, P. H. A Modular Synthetic Route to Size-defined Immunogenic Haemophilus influenzae b Antigens is Key to the Identification of an Octasaccharide Lead Vaccine Candidate. Chem. Sci. 2018, 9 (5), 1279–1288.

(21) Veeneman, G. H.; van Leeuwen, S. H.; van Boom, J. H. Iodonium Ion Promoted Reactions at the Anomeric Centre. II An Efficient Thioglycoside Mediated Approach Toward the Formation of 1,2-trans Linked Glycosides and Glycosidic Esters. Tetrahedron Lett. 1990, 31 (9), 1331–1334.

(22) Wegmann, B.; Schmidt, R. R. The Application of the Trichloroacetimidate Method to the Synthesis of α-D-Gluco- and α-D-Galactopyranosides. J. Carbohydr. Chem. 1987, 6 (3), 357–375.

(23) Marugg, J. E.; Tromp, M.; Kuyl-Yeheskiely, E.; van der Marel, G. A.; van Boom, J. H. A Convenient and General Approach to the Synthesis of Properly Protected D-Nucleoside-3′-Hydrogenphosphonates via Phosphite intermediates. Tetrahedron Lett. 1986, 27 (23), 2661–2664.

(24) Castelli, R.; Overkleeft, H. S.; van der Marel, G. A.; Codée, J. D. C. 2,2-Dimethyl-4-(4-methoxy-phenoxy) butanoate and 2,2-Dimethyl-4-azido Butanoate: Two New Pivaloate-ester-like Protecting Groups. Org. Lett. 2013, 15 (9), 2270–2273.

(25) Brady, A. M.; Calix, J. J.; Yu, J.; Geno, K. A.; Cutter, G. R.; Nahm, M. H. Low Invasiveness of Pneumococcal Serotype 11A Is Linked to Ficolin-2 Recognition of O-acetylated Capsule Epitopes and Lectin Complement Pathway Activation. J. Infect. Dis. 2014, 210 (7), 1155–1165.

(26) Henrichsen, J. Six Newly Recognized Types of Streptococcus pneumoniae. J. Clin. Microbiol. 1995, 33 (10), 2759–2762.

(27) Bidula, S.; Sexton, D. W.; Schelenz, S. Ficolins and the Recognition of Pathogenic Microorganisms: An Overview of the Innate Immune Response and Contribution of Single Nucleotide Polymorphisms. J. Immunol. Res. 2019, 2019.

(28) Endo, Y.; Takahashi, M.; Iwaki, D.; Ishida, Y.; Nakazawa, N.; Kodama, T.; Matsuzaka, T.; Kanno, K.; Liu, Y.; Tsuchiya, K.; et al. Mice Deficient in Ficolin, a Lectin Complement Pathway Recognition Molecule, Are Susceptible to Streptococcus pneumoniae Infection. J. Immunol. 2012, 189 (12), 5860.

(29) Garlatti, V.; Belloy, N.; Martin, L.; Lacroix, M.; Matsushita, M.; Endo, Y.; Fujita, T.; Fontecilla-Camps, J. C.; Arlaud, G. J.; Thielens, N. M.; et al. Structural Insights into the Innate Immune Recognition Specificities of L- and H-Ficolins. EMBO J. 2007, 26 (2), 623–633.

(30) Howard, M.; Farrar, C. A.; Sacks, S. H. Structural and Functional Diversity of Collectins and Ficolins and Their Relationship to Disease. Semin. Immunopathol. 2018, 40 (1), 75–85.

(31) Geno, K. A.; Bush, C. A.; Wang, M. N.; Jin, C.; Nahm, M. H.; Yang, J. H. WciG -Acetyltransferase Functionality Differentiates Pneumococcal Serotypes 35C and 42. J. Clin. Microbiol. 2017, 55 (9), 2775–2784.

