## Supplementary material for "Chemical Synthesis of Oligosaccharides Derived from *Streptococcus pneumoniae* Serotype 35B and D to Investigate Binding of Complement and Serum Factors": SI

### Table of Contents

**Experimental**……………………………………………………………………………S1

**General Methods**…………………………………………………………………………..S1

**Microarray Printing Procedure**…………………………………………………………...S24

**Microarray Binding Studies with Factor Sera 29b and 35a**…………………………...S24

**Microarray Binding Studies with Ficolin-2**……………………………………………..S24

**References**…………………………………………………………………………………S25

**NMR Spectra**...……………………………………………………………………………S26

### Experimental

**General Methods**. All reagents, unless otherwise stated, were purchased from Sigma-Aldrich. $^{1}$H and $^{13}$C NMR spectra were recorded either on Varian Mercury 400 MHz, Bruker 600 or 750 MHz instrument. Chemical shifts are reported in parts per million (ppm) relative to $D_2O$ or $CDCl_3$ as the internal standards. NMR data are presented as follows: Chemical shift, multiplicity (s = singlet, d = doublet, t = triplet, dd = doublet of doublets, m = multiplet and/or multiple resonances, app = apparent); coupling constants are reported in Hertz (Hz). All NMR signals were assigned on the basis of $^{1}$H NMR, COSY and HSQC experiments. For oligosaccharide assignments the following abbreviations were used: $\boldsymbol{\delta}^{\mathbf{Rib}}$ – D-ribose; $\boldsymbol{\delta}^{\mathbf{Gal1}}$ – D-galactofuranose (acetylated); $\boldsymbol{\delta}^{\mathbf{Glc}}$ – D-glucose; $\boldsymbol{\delta}^{\mathbf{Gal2}}$ – galactofuranose (non-acetylated); $\boldsymbol{\delta}^{\mathbf{GalN}}$ –*N*-acetyl-D-galactosamine. Mass spectra were recorded on either on an Applied Biosystems SCIEX MALDI TOF/TOF 5800 mass spectrometer, a Shimadzu Biotech Axima-CFR MALDI-TOF, or a high resolution Shimadzu LCMS-IT-TOF mass spectrometer. Column chromatography was performed on silica gel G60 (Silicycle, 60-200 µm, 60 Å). TLC analysis was conducted on Silicagel 60 F254 (EMD Chemicals Inc.) with detection by UV light (254 nm) where applicable, and by charring with 10% sulfuric acid in ethanol or a solution of $(NH_4)_6Mo_7O_2*4H_2O$ (25 g/L) in 10% sulfuric acid in ethanol. All reactions were carried out under argon atmosphere unless specified otherwise. Unless otherwise stated, all reactions were carried out at room temperature (RT) in glassware with magnetic stirring. Solutions in organic solvents were dried with $Na_2SO_4$ and concentrated at 40 °C/2 kPa. Molecular sieves were flame-dried under vacuum immediately prior to use.

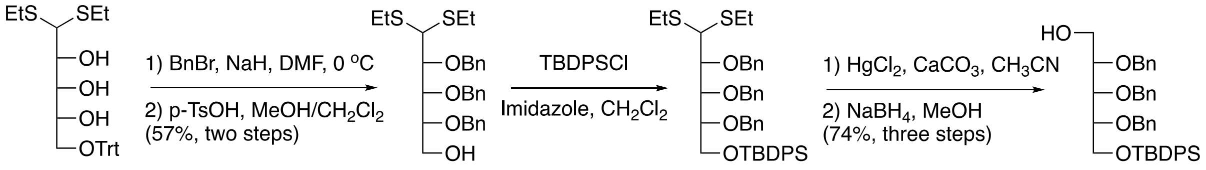


**S1** **S2** **S3** **7**

**Scheme S1.** Synthesis of acceptor **7**

#### 2,3,4-Tri-*O*-benzyl-D-ribose diethyl dithioacetal (S2)

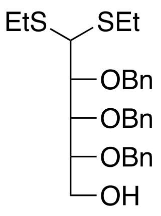


Compound **S1**[1] (4.3 g, 8.58 mmol) was dissolved in anhydrous DMF (20 mL), after which NaH (1.7 g, 42.9 mmol) and BnBr (4.6 mL, 38.61 mmol) were added at 0 °C, and the reaction mixture was allowed to stir at that temperature for 2 h. The mixture was then quenched with MeOH/AcOH (10 mL, 1:1) and concentrated to dryness. The obtained syrup was dissolved in $CH_2Cl_2$ (100 mL), and the solution was washed with water, and the organic phase was dried and concentrated. The resulting syrup was dissolved in methanol, after which a catalytic amount of *p*-TsOH was added, and the reaction mixture was stirred at rt for 1 h. The mixture was concentrated, absorbed on silica gel, and purified using EtOAc:Hexane (5:95 to 1:9) to afford the title product as a clear syrup (2.7 g, 57% over 2 steps). $^{1}$H NMR (400 MHz, $CDCl_3$): δ 1.19 (6H, m, $SCH_2\underline{CH_3}$ x 2), 2.22 (1H, dd, *J* = 7.1, 5.9 Hz, 6-OH), 2.52 – 2.68 (4H, m, $S\underline{CH_2}CH_3$ x 2), 3.68 – 3.79 (2H, m, H-6 x 2), 3.84 – 3.91 (2H, m, H-4, H-5), 4.19 (2H, m, H-2, H-1), 4.55 – 4.73 (4H, m, $OC\underline{H_2}Ph$ x 2), 4.83 (1H, d, *J* = 11.0 Hz, $OCH\underline{H}Ph$), 4.93 (1H, d, *J* = 11.0 Hz, $OC\underline{H}HPh$), 7.24 – 7.36 (15H, m, Ar-H). $^{13}$C NMR (101 MHz, $CDCl_3$) δ 138.2, 138.1, 137.9, 128.5, 128.4, 128.3, 128.2, 128.1, 128.0, 127.9 x 2, 127.8, 127.8 x 3, 127.7 x 2, 127.6, 126.9, 82.3, 79.7, 79.6, 74.9, 73.8, 72.0, 61.7, 53.7, 26.0, 25.2, 14.4 x 2. MALDI-MS: $[M + Na]^+$ $C_{30}H_{38}NaO_4S_2$ calcd. 549.2109, found 549.2123.

**$t$-Butyldiphenylsilyl 2,3,4-tri-$O$-benzyl-D-ribose (7)**

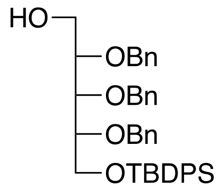


Compound **S2** (40.4 g, 76.69 mmol) was dissolved in $CH_2Cl_2$ (100 mL) followed by adding imidazole (16.0 g, 230.07 mmol) and TBDPSCl (26 mL, 92.03 mmol), after which the reaction mixture was left stirring at rt for 1 h. The mixture was washed with 1M HCl (100 mL), and the organic phase was dried and concentrated to give crude **S3** as a syrup, which was directly dissolved in $CH_3CN$ (100 mL) followed by adding $CaCO_3$ (15.5 g, 153.38 mmol) and $HgCl_2$ (42.0 g, 153.38 mmol) at 0 °C. The reaction mixture was stirred at that temperature for 1 h, and then at rt for 1 h. The mixture was then filtered and concentrated *in vacuo* to dryness. This material was dissolved in $CH_3OH$ (200 mL), after which $NaBH_4$ (10.0 g) was added at 0 °C, and stirring was continued at that temperature for 30 min. The mixture was neutralized with AcOH, and concentrated *in vacuo* to dryness. The residue was loaded on silica gel and purified using EtOAc:Hexane (5:95 to 1:9) as a mobile phase to give the target product as a clear oil (42.0 g, 82% over three steps). $^1$H NMR (400 MHz, $CDCl_3$): δ 1.09 (9H, s, $(CH_3)_3$ of TBDPS), 2.34 (1H, t, $J$ = 5.98 Hz, 1-OH), 3.77 (4H, m, incl. H-1, H-4), 3.92 (2H, d, $J$ = 4.5 Hz, H-5), 3.99 (1H, t, $J$ = 4.8 Hz), 4.54 – 4.74 (6H, m, <u>CH2</u>Ph), 7.21 – 7.71 (25H, m, Ar-H). $^{13}$C NMR (101 MHz, $CDCl_3$) δ 138.3, 138.2, 138.1, 135.7, 135.6, 133.4, 133.3, 129.7, 129.7, 128.4, 128.4, 128.3, 128.0, 127.8, 127.7, 127.7, 127.7, 127.7, 127.5, 79.7, 79.0, 79.0, 73.9, 72.6, 71.8, 63.5, 61.5, 26.9, 19.2. MALDI-MS: $[M + Na]^+$ $C_{42}H_{48}NaO_5Si$ calcd. 683.3169, found 683.3154.

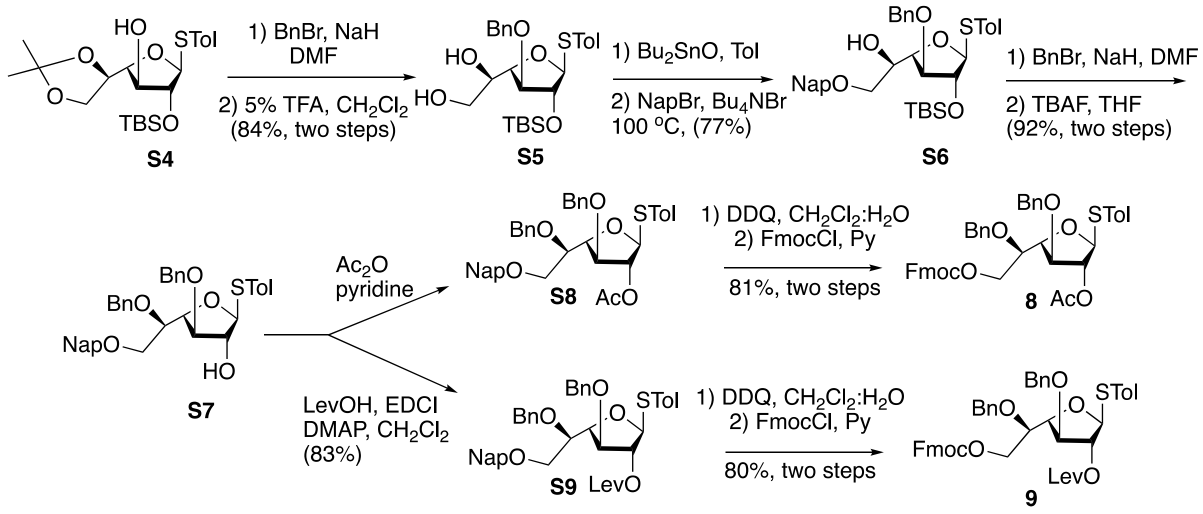


**Scheme S2.** Synthesis of **8** and **9**

**4-Methyphenyl 2-$O$-$t$-butyldimethylsilyl-3-$O$-benzyl-1-thio-$\beta$-D-galactofuranoside (S5)**

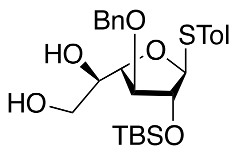


Compound **S4**[2] (26.0 g, 59.0 mmol) was dissolved in DMF (100 mL), and BnBr (10.5 mL, 88.5 mmol) and NaH (3.6 g, 88.5 mmol) were added at 0 °C in succession. The reaction mixture was then stirred at 0 °C for 15 min, and then at rt for 1 h. The reaction mixture was quenched by the addition of MeOH:AcOH (1:1), concentrated under reduced pressure to give the product as an oil. This material was dissolved in $CH_2Cl_2$ (100 mL), after which TFA (10 mL) and water (2 mL) were added, and the reaction mixture was left stirring at rt for 3 h, after which it was diluted with $CH_2Cl_2$ (100 mL), washed with water and sat. $NaHCO_3$ in succession. The organic phase was concentrated under reduced pressure, and the product was absorbed on silica gel and purified using EtOAc:Hexane (3:7) as a phase to give the target product as a clear oil (24.5 g, 84% over two steps). $^1$H NMR (400 MHz, $CDCl_3$): δ 0.12 (6H, s, $CH_3$ x 2 of TBS), 0.88 (9H, s, $t$Bu of TBS), 2.17 (1H, dd, $J$ = 8.7,

4.6 Hz, 6-OH), 2.32 (3H, s, $CH_3$ of STol), 2.62 (1H, d, $J$ = 6.8 Hz, 5-OH), 3.61 – 3.72 (2H, m, H-6), 3.77 (1H, m, H-5), 3.99 (1H, dd, $J$ = 6.0, 2.7 Hz, H-3), 4.30 (1H, dd, $J$ = 3.7, 6.0 Hz, H-4), 4.34 (1H, t, $J$ = 2.5 Hz, H-2), 4.57 (1H, d, $J$ = 11.8 Hz, CHHPh), 4.70 (1H, d, $J$ = 11.8 Hz, CHHPh), 5.24 (1H, d, $J$ = 2.17 Hz, H-1), 7.09 – 7.39 (9H, m, Ar-H). $^{13}$C NMR (101 MHz, $CDCl_3$) δ 137.8, 137.4, 132.5, 130.8, 129.8, 128.5, 128.0, 127.7, 94.4, 85.7, 85.7, 82.9, 81.6, 72.5, 70.8, 64.6, 25.7, 25.7, 25.6, 21.1, 17.9, -4.4, -5.0. MALDI-MS: $[M + Na]^+$ $C_{26}H_{38}NaO_5SSi$ calcd. 513.2107, found 513.2112.

**4-Methyphenyl 2-*O*-*t*-butyldimethylsilyl-3-*O*-benzyl-6-*O*-(2-naphthyl)-1-thio-*β*-D-galactofuranoside (S6)**

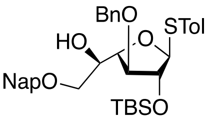


Compound **S5** (3.51 g, 7.15 mmol) was dissolved in toluene (20 mL), followed by adding $Bu_2SnO$ (2.1 g, 8.58 mmol), and the reaction mixture was stirred at 100 °C for 2 h. After complete dissolution of $Bu_2SnO$, NapBr (2.3 g, 10.72 mmol) and $Bu_4NBr$ (2.3 g, 7.15 mmol) were added, and stirring was further continued for 6 h. The reaction mixture was then concentrated, diluted with $CH_2Cl_2$, and washed with water and sat. $NaHCO_3$ in succession. The organic phase was then filtered, dried and concentrated. The crude product was purified on silica gel using EtOAc:Hexane (5:95 to 1:9) as a mobile phase, giving the target product as clear oil (3.53 g, 77%). $^1$H NMR (400 MHz, $CDCl_3$): δ 0.12 (6H, s, $CH_3$ x 2 of TBS), 0.88 (9H, s, *t*Bu of TBS), 2.32 (3H, s, $CH_3$ of STol), 2.57 (1H, d, $J$ = 6.8 Hz, 5-OH), 3.60 (2H, m, H-6), 3.95 – 4.02 (2H, m, H-3, H-5), 4.36 (1H, t, $J$ = 2.39 Hz, H-2), 4.40 (1H, dd, $J$ = 5.40, 3.04 Hz, H-4), 5.27 (1H, d, $J$ = 2.1 Hz, H-1), 7.02 – 7.84 (16H, m, Ar-H). $^{13}$C NMR (101 MHz, $CDCl_3$) δ 137.6, 137.4, 135.6, 133.2, 133.0, 132.3, 131.2, 129.7, 128.5, 128.2, 127.9, 127.9, 127.7, 127.7, 126.5, 126.1, 125.8, 125.7, 94.3, 85.6, 82.2, 81.6, 73.5, 72.4, 71.5, 69.9, 25.7, 21.1, 17.9, -4.4, -5.0. MALDI-MS: $[M + Na]^+$ $C_{37}H_{46}NaO_5SSi$ calcd. 653.2733, found 653.2737.

**4-Methyphenyl 3,5-di-*O*-benzyl-6-*O*-(2-naphthyl)-1-thio-*β*-D-galactofuranoside (S7)**

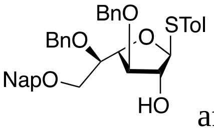


Compound **S6** (3.39 g, 5.37 mmol) was dissolved in DMF (20 mL), after which BnBr (963 µL, 8.05 mmol) and NaH (322 mg, 8.05 mmol) were added at 0 °C, and the reaction mixture was left stirring at that temperature for 1 h, after which it was quenched with $CH_3OH$, followed by adding AcOH. The reaction mixture was then concentrated, dissolved in THF (20 mL) followed by adding TBAF (3.4 g, 10.74 mmol, hydrate form), and stirring was continued at rt for 3 h. The solution was then concentrated, diluted with $CH_2Cl_2$ and washed with water. The organic phase was then dried and concentrated to give a residue, which was purified on silica gel using EtOAc:Hexane (3:7) as a mobile phase to give the product as a clear oil (3.03 g, 92%, over two steps). $^1$H NMR (400 MHz, $CDCl_3$): δ 2.29 (3H, s, $CH_3$ of STol), 3.66 (1H, d, $J$ = 10.7 Hz, 2-OH), 3.72 – 3.79 (2H, m, H-5, H-6a), 3.79 – 3.85 (2H, m, H-6b, H-3), 4.27 (1H, d, $J$ = 10.8 Hz, H-2), 4.44 (1H, d, $J$ = 11.8 Hz, CHHAr), 4.47 – 4.50 (2H, m, H-4, CHHAr), 4.64 (1H, d, $J$ = 11.9 Hz, CHHAr), 4.70 (2H, s, $CH_2$Ar), 4.76 (1H, d, $J$ = 11.2 Hz, CHHAr), 5.40 (1H, s, H-1), 7.02 – 7.85 (21H, m, Ar-H). $^{13}$C NMR (101 MHz, $CDCl_3$) δ 137.5, 136.9, 136.9, 135.4, 133.2, 133.0, 131.9, 131.6, 129.6, 128.6, 128.5, 128.4, 128.3, 128.2, 127.9, 127.8, 127.8, 127.7, 126.5, 126.1, 125.9, 125.6, 95.1, 85.3, 83.8, 79.0, 76.9, 73.7, 73.4, 72.0, 70.4, 21.0. MALDI-MS: $[M + Na]^+$ $C_{38}H_{38}NaO_5S$ calcd. 629.2338, found 629.2345.

**4-Methyphenyl 2-*O*-acetyl-3,5-di-*O*-benzyl-6-*O*-(2-naphthyl)-1-thio-*β*-D-galactofuranoside (S8)**

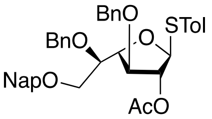

Compound **S7** (3.03 g, 4.99 mmol) was dissolved in $CH_2Cl_2$ (20 mL), followed by adding pyridine (1.6 mL, 19.96 mmol) and $Ac_2O$ (943 µL, 9.98 mmol). The reaction mixture was left stirring at rt for 2 h, after which it was diluted with $CH_2Cl_2$ (50 mL), washed with 1 M HCl, concentrated, and purified on silica gel using EtOAc:Hexane (1:4) as a mobile phase, to give the product as a clear oil (3.18 g, quant.). $^1$H NMR (400 MHz, $CDCl_3$): δ 1.90 (3H, s, OAc), 2.29 (3H, s, $CH_3$ of STol), 3.67 (1H, dd, *J* = 9.9, 5.1 Hz, H-6a), 3.71 – 3.81 (2H, m, H-6b, H-5), 3.97 (1H, d, *J* = 6.0 Hz, H-3), 4.34 (1H, d, *J* = 11.5 Hz, CH*H*Ar), 4.41 (1H, d, *J* = 11.5 Hz, CH*H*Ar), 4.49 (1H, dd, *J* = 6.2, 2.9 Hz, H-4), 4.61 – 4.71 (4H, m, $CH_2$Ar), 5.19 (1H, t, *J* = 1.5 Hz, H-2), 5.49 (1H, s, H-1), 6.97 – 7.86 (21H, m, Ar-H). $^{13}$C NMR (101 MHz, $CDCl_3$) δ 169.9, 138.2, 137.6, 137.4, 135.6, 133.2, 133.0, 132.2, 130.3, 129.6, 128.4, 128.2, 128.2, 128.2, 128.1, 127.9, 127.8, 127.7, 127.7, 126.3, 126.1, 125.8, 125.6, 91.0, 82.6, 82.3, 82.0, 76.4, 73.6, 73.5, 72.1, 70.6, 21.1, 20.8. MALDI-MS: $[M + Na]^+$ $C_{40}H_{40}NaO_6S$ calcd. 671.2443, found 671.2450.

**4-Methyphenyl 2-*O*-acetyl-3,5-di-*O*-benzyl-6-*O*-fluorenylmethoxycarbonyl-1-thio-*β*-D-galactofuranoside (8)**

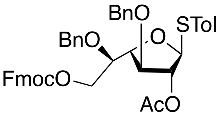

To a solution of compound **S8** (13.8 g, 21.2 mmol) in $CH_2Cl_2$ (100 mL) were added $H_2O$ (10 mL) and DDQ (7.1 g, 31.9 mmol), and the reaction mixture was left stirring at rt for 1 h. The reaction mixture was quenched with sat. $NaHCO_3$, filtered, and washed with sat. $NaHCO_3$. The organic phase was dried, concentrated, and the product purified on silica gel to give the alcohol as a clear oil. This material was then dissolved in pyridine (50 mL), after which Fmoc-Cl (7.6 g, 29.3 mmol) was added at 0 °C and the reaction mixture was left stirring at rt for 1 h. After that time pyridine was removed *in vacuo*, the crude mixture was dissolved in $CH_2Cl_2$ (100 mL), washed with 1 M HCl, the organic phase was then dried and concentrated to give the product, which was purified on silica gel using EtOAc:Hexane (1:4) as a mobile phase, providing the target donor as a white foam (12.6 g, 81%, over two steps). $^1$H NMR (400 MHz, $CDCl_3$): δ 1.96 (3H, s, OAc), 2.29 (3H, s, STol), 3.85 (1H, m, H-5), 3.97 (1H, d, *J* = 5.7 Hz, H-3), 4.23 (1H, t, *J* = 7.4 Hz, CH of Fmoc), 4.32 – 4.40 (5H, m, H-6, CHHPh, $CH_2$ of Fmoc), 4.45 (1H, d, *J* = 12.4 Hz, CHHPh), 4.50 (1H, dd, *J* = 5.7, 3.4 Hz, H-4), 4.64 – 4.70 (2H, m, CHHPh x 2), 5.22 (1H, t, *J* = 1.4 Hz, H-2), 5.50 (1H, s, H-1), 7.05 – 7.77 (22H, m, Ar-H). $^{13}$C NMR (101 MHz, $CDCl_3$) δ 169.8, 155.0, 143.3, 141.3, 137.7, 137.4, 132.4, 130.1, 129.7, 128.4, 128.3, 128.2, 128.1, 128.0, 127.9, 127.9, 127.9, 127.2, 125.1, 125.1, 120.0, 91.3, 82.6, 81.8, 81.7, 75.0, 73.5, 72.2, 70.0, 67.4, 46.7, 21.1, 21.0, 20.8. MALDI-MS: $[M + Na]^+$ $C_{44}H_{42}NaO_8S$ calcd. 753.2498, found 753.2494.

**4-Methyphenyl 2-*O*-levulinolyl-3,5-di-*O*-benzyl-6-*O*-(2-naphthyl)-1-thio-*β*-D-galactofuranoside (S9)**

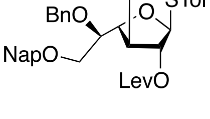

Compound **S7** (12.54 g, 20.6 mmol) was dissolved in $CH_2Cl_2$ (100 mL), followed by adding LevOH (3.6 mL, 31 mmol), EDCI (6.0 g, 31 mmol) and DMAP (catalytic amount). The reaction mixture was left stirring at rt for 2 h., after which it was diluted with $CH_2Cl_2$ (100 mL), washed with sat. $NaHCO_3$, the organic phase was dried ($Na_2SO_4$), concentrated and purified on silica gel using EtOAc:Hexane (3:7 to 2:3) as a mobile system to give the target product as a clear oil (12.0 g, 83%). $^1$H NMR (400 MHz, $CDCl_3$): δ 2.12 (3H, s, $CH_3$ of Lev), 2.27 (3H, s, $CH_3$ of STol), 2.41 (2H, m, $CH_2$ of Lev), 2.61 (2H, m, $CH_2$ of Lev), 3.66 (1H, m, H-6a), 3.73 (1H, m, H-6b), 3.79 (1H, m, H-5), 4.00 (1H, d, *J* = 6.1 Hz, H-3), 4.38 (2H, m, $CH_2$Ar), 4.49 (1H, dd, *J* = 5.7, 3.3 Hz, H-4), 4.66 (4H, m, $CH_2$Ar x 2), 5.22 (1H, t, *J* = 1.6 Hz, H-2), 5.48 (1H, s, H-1), 6.99 –

7.83 (21H, m, Ar-H), $^{13}$C NMR (101 MHz, $CDCl_3$) δ 206.1, 171.7, 138.2, 137.6, 137.4, 135.6, 133.2, 132.9, 132.3, 130.3, 129.6, 128.4, 128.4, 128.2, 128.2, 128.2, 127.9, 127.9, 127.7, 126.3, 126.1, 125.8, 125.6, 90.9, 82.4, 82.1, 82.1, 76.4, 73.6, 73.5, 72.1, 70.5, 37.7, 29.7, 27.8, 21.1. MALDI-MS: $[M + Na]^+$ $C_{43}H_{44}NaO_7S$ calcd. 727.2705, found 727.2712.

**4-Methyphenyl 2-*O*-levulinoyl-3,5-di-*O*-benzyl-6-*O*-fluorenylmethoxycarbonyl-1-thio-*β*-D-galactofuranoside (9)**

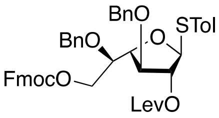


Compound **S9** (15.0 g, 21.3 mmol) was dissolved in $CH_2Cl_2$ (100 mL), $H_2O$ (10 mL) and DDQ (6.3 g, 31.9 mmol) were then added, and the reaction mixture was left stirring at rt for 1 h. It was quenched with sat. $NaHCO_3$, filtered, and washed with sat. $NaHCO_3$. The organic phase was dried, concentrated, and the product purified on silica gel to give the alcohol as a clear oil. This material was then dissolved in pyridine (50 mL), after which Fmoc-Cl (8.3 g, 31.9 mmol) was added at 0 $^oC$ and the reaction mixture was left stirring at rt for 1 h. After that time pyridine was removed *in vacuo*, the crude mixture was dissolved in $CH_2Cl_2$ (100 mL), washed with 1 M HCl, the organic phase was then dried and concentrated to give the product, which was purified on silica gel using EtOAc:Hexane (3:7) system as a mobile phase, providing the target donor as a white foam (13.3 g, 80%, over two steps). $^1$H NMR (400 MHz, $CDCl_3$): δ 2.14 (3H, s, $CH_3$ of Lev), 2.28 (3H, s, $CH_3$ of STol), 2.48 (2H, m, $CH_2$ of Lev), 2.68 (2H, m, $CH_2$ of Lev), 3.84 (1H, m, H-5), 4.22 (1H, t, $J$ = 7.4 Hz, CH of Fmoc), 4.29 – 4.45 (6H, m, H-6, $CH_2$ of Fmoc, C<u>H</u>HAr x 2), 4.49 (1H, dd, $J$ = 3.5, 5.8 Hz, H-4), 4.63 (1H, d, $J$ = 11.3 Hz, C<u>H</u>HAr), 4.68 (1H, d, $J$ = 11.8 Hz, C<u>H</u>HAr), 5.24 (1H, t, $J$ = 1.68 Hz, H-2), 5.47 (1H, s, H-1), 7.04 – 7.78 (22H, m, Ar-H). $^{13}$C NMR (101 MHz, $CDCl_3$) δ 206.1, 171.7, 154.9, 143.3, 143.3, 141.3, 141.2, 137.7, 137.7, 137.4, 132.5, 130.0, 129.6, 128.4, 128.3, 128.2, 128.2, 127.9, 127.9, 127.9, 127.9, 127.2, 125.1, 125.1, 120.1, 120.0, 91.1, 82.5, 81.9, 81.6, 75.0, 73.5, 72.2, 69.9, 69.9, 67.4, 46.8, 46.7, 37.7, 29.7, 27.8, 21.1. MALDI-MS: $[M + Na]^+$ $C_{47}H_{46}NaO_9S$ calcd. 809.2760, found 809.2778.

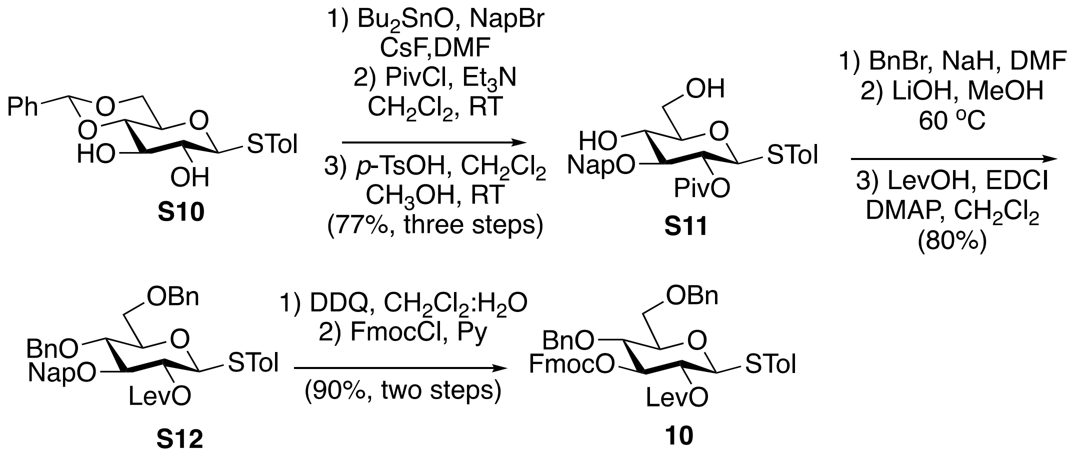


**Scheme S3.** Synthesis of **10**

**4-Methyphenyl 2-*O*-pivaloyl-3-*O*-(2-naphthyl)-1-thio-*β*-D-glucopyranoside (S11)**

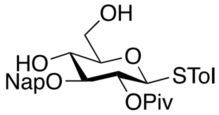

Diol **S10**[3] (33.75 g, 90.13 mmol) was suspended in toluene (200 mL), followed by adding $Bu_2SnO$ (27.0 g, 108.15 mmol), and the mixture was heated at 110 °C until clear solution was obtained. The reaction mixture was concentrated, dissolved in DMF (100 mL) after which CsF (21.0 g, 135.19 mmol) and NapBr (26.0 g, 117.169 mmol) were added, and the reaction mixture was left stirring at rt until the starting material was fully consumed. The reaction mixture was concentrated, the crude intermediate was dissolved in $CH_2Cl_2$ and washed with water and sat. $NaHCO_3$. The organic phase was then filtered, dried and concentrated to give the intermediate as a white solid. To a solution of this material in $CH_2Cl_2$ (200 mL) was added $Et_3N$ (38 mL, 270.4 mmol) and DMAP (11.0 g, 90.13 mmol), after which the solution was cooled to 0 °C, and PivCl (17 mL, 135.2 mmol) was added dropwise, and the reaction mixture was left stirring at rt for 1 h, after which it was diluted with $CH_2Cl_2$, washed with 1 M HCl, and the organic phase was dried and concentrated to provide a crude intermediate as a syrup, which was dissolved in $CH_2Cl_2$:$CH_3OH$ (1:1, 200 mL), and CSA (5.0 g) was then added. The reaction mixture was left stirring at rt overnight, after which it was concentrated, diluted with $CH_2Cl_2$, washed with sat. $NaHCO_3$, the organic phase was dried ($Na_2SO_4$) and concentrated to give a crude product, which was crystallized from ethanol to provide a pure title compound as an off-yellow powder (35.0 g, 77% over three steps). $^1$H NMR (400 MHz, $CDCl_3$): δ 1.25 (9H, s, $CH_3$ x 3 Piv), 2.32 (3H, s, $CH_3$ of STol), 2.69 (1H, br m, 4-OH), 3.37 (1H, m, H-5), 3.61 (1H, t, $J$ = 8.6 Hz, H-3), 3.68 (1H, t, $J$ = 9.0 Hz, H-4), 3.76 (1H, dd, $J$ = 11.7, 4.59 Hz, H-6a), 3.87 (1H, dd, $J$ = 12.5, 3.4 Hz, H-6b), 4.63 (1H, d, $J$ = 9.7 Hz, H-1), 4.78 (1H, d, $J$ = 11.5 Hz, C*H*HAr), 4.88 (1H, d, $J$ = 11.5 Hz, CH*H*Ar), 5.05 (1H, dd, $J$ = 9.7, 9.0 Hz, H-2), 7.08 – 7.8 (11H, m, Ar-H). $^{13}$C NMR (101 MHz, $CDCl_3$) δ 176.8, 138.3, 138.2, 135.4, 135.4, 133.2, 133.0, 132.7, 132.7, 129.7, 129.2, 129.1, 128.4, 128.4, 127.9, 127.7, 126.3, 126.3, 126.2, 126.0, 126.0, 125.8, 125.3, 87.0, 84.3, 79.3, 79.3, 74.7, 71.4, 70.2, 62.5, 27.2, 21.1. MALDI-MS: $[M + Na]^+$ $C_{29}H_{34}NaO_6S$ calcd. 533.1974, found 533.1983.

**4-Methyphenyl 4,6-di-*O*-benzyl-2-*O*-levulinoyl-3-*O*-(2-naphthyl)-1-thio-*β*-D-glucopyranoside (S12)**

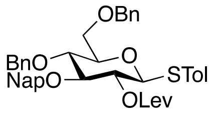

Diol **S11** (34.0 g, 66.5 mmol) was dissolved in anhydrous DMF (200 mL) followed by adding NaH (7.8 g, 199.5 mmol) and BnBr (24 mL, 199.5 mmol) at 0 °C and the reaction mixture was stirred at that temperature for 30 min. It was then neutralized with $CH_3OH$, concentrated, and the resulting intermediate was dissolved in $CH_3OH$/$CH_2Cl_2$ (200 mL, 3:1), LiOH (5.0 g, 199.5 mmol) was then added and the reaction was heated at 60 °C for 12 h. The mixture was then concentrated, dissolved in $CH_2Cl_2$ (300 mL), and washed with $H_2O$ (200 mL), the organic phase was dried and concentrated to afford an intermediate which was directly dissolved in $CH_2Cl_2$ (200 mL), followed by adding LevOH (12 mL, 99.75 mmol), EDCI (19.2 g, 99.75) and DMAP (1.6 g, 13.3 mmol). The reaction mixture was stirred at RT for 2 h, after which it was washed with sat. $NaHCO_3$ (200 mL), dried and concentrated to give the crude product as an oil, which was chromatographed using EtOAc:Hexane (3:7), (37.0 g, 80% over three steps). $R_f$ = 0.4 (EtOAc:Hexane, 2:3). $^1$H NMR (400 MHz, $CDCl_3$): δ 2.08 (3H, s, $CH_3$ of Lev), 2.28 (3H, s, $CH_3$ of STol), 2.37 – 2.62 (4H, m, $CH_2$ x 2 of Lev), 3.52 (1H, m, H-5), 3.67 (1H, dd, *J* = 8.9 Hz, H-4), 3.72 (1H, m, H-3), 3.71 – 3.80 (3H, m, H-6 x 2, H-3), 4.50 – 4.60 (3H, C*H*HAr x 2, H-1), 4.79 (1H, d, *J* = 10.9 Hz, C*H*HAr), 4.85 (1H, d, *J* = 11.7 Hz, C*H*HAr), 4.93 (1H, d, *J* = 11.7 Hz, C*H*HAr), 5.00 (1H, dd, *J* = 9.0, 9.9 Hz, H-2), 7.00 – 7.84 (21H, m, Ar-H). $^{13}$C NMR (101 MHz, $CDCl_3$) δ 206.1, 171.4, 138.2, 138.1, 137.9, 135.7, 133.2, 133.2, 132.9, 129.6, 128.7, 128.4, 128.3, 128.1, 127.9, 127.8, 127.7, 127.6, 127.5, 126.6, 126.0, 126.0, 125.9, 86.2, 84.4, 79.4, 77.8, 75.3, 75.1, 73.5, 72.2, 68.9, 37.7, 29.7, 28.1, 21.1. MALDI-MS: $[M + Na]^+$ $C_{43}H_{44}NaO_7S$ calcd. 727.2705, found 727.2717.

**4-Methyphenyl 4,6-di-*O*-benzyl-2-*O*-levulinoyl-3-*O*-fluorenylmethyl-1-thio-*β*-D-glucopyranoside (10)**

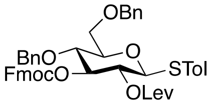


Compound **S12** (5.0 g, 7.09 mmol) was dissolved in $CH_2Cl_2$ (20 mL), after which $H_2O$ (2 mL) was added followed by DDQ (2.4 g, 10.64 mmol). The reaction mixture was left stirring at RT for 1 h, after which it was neutralized with sat. $NaHCO_3$, diluted with $CH_2Cl_2$ (50 mL), and the organic layer was then dried and concentrated to give an oil, which was purified on silica gel using EtOAc:Hexane (3:7) as a mobile phase. This intermediate was then dissolved in pyridine (20 mL), after which Fmoc-Cl (2.7 g, 10.6 mmol) was added and the mixture was left stirring at RT for 1 h, after which the reaction mixture was concentrated, re-dissolved in $CH_2Cl_2$ (50 mL) and washed with 1M HCl. The organic layer was then dried and concentrated to give the product which was chromatographed using EtOAc:Hexane (1:10 to 3:7), (6.74 g, 90% over two steps). $R_f$ = 0.6 (EtOAc:Hexane, 3:7). $^1$H NMR (400 MHz, $CDCl_3$): δ 2.04 (3H, s, $CH_3$ of Lev), 2.29 (3H, s, $CH_3$ of STol), 2.54 – 2.75 (4H, m, $CH_2$ x 2 of Lev), 3.54 (1H, m, H-5), 3.70 – 3.79 (3H, m, H-4, H-6 x 2), 4.22 (2H, m, CH of Fmoc, CH*H*Fmoc), 4.48 (1H, m, C*H*HFmoc), 4.50 – 4.59 (4H, m, $CH_2$Ar), 4.61 (1H, d, *J* = 9.7 Hz, H-1), 7.01 – 7.73 (18H, m, Ar-H). $^{13}$C NMR (101 MHz, $CDCl_3$) δ 205.8, 171.3, 154.6, 143.6, 143.2, 141.2, 141.1, 138.4, 138.1, 137.5, 133.7, 129.6, 128.3, 128.3, 127.9, 127.8, 127.8, 127.8, 127.6, 127.6, 127.1, 125.3, 125.2, 119.9, 85.6, 80.7, 79.1, 75.6, 74.9, 73.5, 70.4, 70.4, 68.6, 46.6, 37.7, 29.6, 28.0, 21.1. MALDI-MS: $[M + Na]^+$ $C_{47}H_{46}NaO_9S$ calcd. 809.2760, found 809.2748.

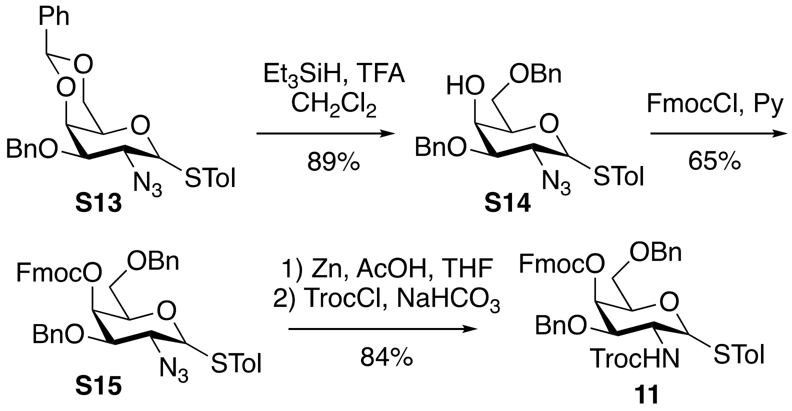

**Scheme S4.** Synthesis of donor **11**

**4-Methyphenyl 2-azido-2-deoxy-3,6-di-*O*-benzyl-1-thio-*α,β*-D-galactopyranoside (S14)**

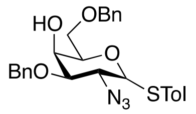

To a solution of **S13**[4] (21.67 g, 44.26 mmol) in $CH_2Cl_2$ (160 mL) containing $Et_3SiH$ (36 mL, 221.3 mmol) was added TFA (17 mL, 221.3 mmol) at 0 °C. The reaction mixture was stirred at that temperature for 30 min, and further at RT for 2 h. The reaction mixture was diluted with $CH_2Cl_2$ (50 mL) and washed with sat. $NaHCO_3$ (100 mL). The organic phase was then dried and concentrated to give the crude product as an oil, which was chromatographed using EtOAc:Hexane (1:4 to 3:7) to give the pure product as a clear oil, (21.0 g, 89%). $^1$H NMR (400 MHz, $CDCl_3$): δ 2.29 (3H, s, $CH_3$ of STol), 2.30 (3H, s, $CH_3$ of STol), 2.42 (1H, m, 4-OH), 2.62 (1H, m, 4-OH), 3.37 (1H, dd, *J* = 9.0, 3.7 Hz, H-3), 3.53 (1H, m, *J* = 5.61 Hz, H-5), 3.58 (1H, dd, *J* = 9.76 Hz, H-2), 3.67 – 3.81 (3H, m, H-6 x 2, H-3), 4.04 (1H, m, H-4), 4.14 (1H, m, H-4), 4.27 (1H, dd, *J* = 10.1, 5.2 Hz), 4.30 (1H, d, *J* = 10.1 Hz, H-1), 4.51 (1H, m, H-5), 4.52 – 4.76 (4H, m, $CH_2$Ar), 5.52 (1H, d, *J* = 5.5 Hz, H-1), 6.99 – 7.50 (14H, m, Ar-H). $^{13}$C NMR (101 MHz, $CDCl_3$) δ 138.5, 138.0, 137.9, 137.8, 137.0, 137.0, 133.8, 133.0, 129.8, 129.7, 129.2, 128.7, 128.6, 128.6, 128.4, 128.4, 128.3, 128.2, 128.0, 128.0, 128.0, 127.8, 127.8, 127.8, 127.7, 127.6, 127.4, 87.7, 86.3, 81.1, 77.8, 77.0, 73.7, 73.6, 72.0, 71.9, 69.7, 69.5, 69.3, 66.7, 65.6, 60.8, 59.7, 21.1, 21.1. MALDI-MS: $[M + Na]^+$ $C_{27}H_{29}N_3NaO_4S$ calcd. 514.1776, found 514.1790.

**4-Methyphenyl 2-azido-2-deoxy-3,6-di-*O*-benzyl-4-*O*-fluorenylmethyl-1-thio-,*β*-D-galactopyranoside (S15)**

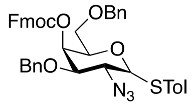

To a solution of compound **S14** (21.0 g, 42.71mmol) in pyridine (100 mL) was added DMAP (cat.) and the solution was cooled to 0 °C, followed by adding Fmoc-Cl (16.7 g, 64.06 mmol). The reaction mixture was left stirring at that temperature for 3 h. It was concentrated, dissolved in $CH_2Cl_2$ (150 mL) and washed with 1M HCl (100 mL). The organic phase was then dried and concentrated to give the crude product as an oil, which was chromatographed using EtOAc:Toluene (0:1 to 5:95) as a solvent system, (19.5 g, 65%). $^1$H NMR (400 MHz, $CDCl_3$): δ 2.27 (3H, s, $CH_3$ of STol), 2.30 (3H, s, $CH_3$ of STol), 3.48 (1H, dd, *J* = 9.7, 3.1 Hz, H-3), 3.55 – 3.74 (4H, H-6 x 2, H-5, H-2), 3.84 (1H, dd, *J* = 10.5, 3.18 Hz, H-3), 4.14 (1H, t, *J* = 7.0 Hz, CH of Fmoc), 4.22 – 4.30 (3H, m, H-2, C*H*HAr x 2), 4.34 – 4.54 (4H, m, H-1, C*H*HAr, $CH_2$ of Fmoc), 4.58 (1H, d, *J* = 10.8 Hz, C*H*HAr), 4.71 (1H, t, *J* = 6.7 Hz, H-5), 4.77 (1H, d, *J* = 11.0 Hz, C*H*HAr), 4.81 (1H, d, *J* = 10.8 Hz, C*H*HAr), 5.41 (1H, d, *J* = 3.15 Hz, H-4), 5.51 (1H, d, *J* = 2.0 Hz, H-4), 5.55 (1H, d, *J* = 5.5 Hz, H-1), 6.99 – 7.80 (22H, m, Ar-H). $^{13}$C NMR (101 MHz, $CDCl_3$) δ 154.8, 154.8, 143.5, 143.5, 143.1, 143.0, 141.3, 141.2, 141.2, 138.5, 138.1, 137.6, 137.5, 136.8, 133.6, 133.1, 129.8, 129.7, 128.9, 128.4, 128.4, 128.3, 128.3, 128.2, 127.9, 127.9, 127.9, 127.9, 127.9, 127.8, 127.8, 127.7, 127.6, 127.2, 127.1, 127.1, 125.3, 125.0, 125.0, 120.0, 120.0, 119.9,

87.6, 86.5, 79.4, 77.2, 76.2, 75.9, 73.7, 73.6, 71.9, 71.8, 70.9, 70.1, 70.1, 69.6, 68.7, 68.2, 67.9, 61.0, 59.9, 46.5, 46.5, 21.2, 21.1. MALDI-MS: $[M + Na]^+$ $C_{42}H_{39}N_3NaO_6S$ calcd. 736.2457, found 736.2478.

**4-Methyphenyl 2-(2,2,2-trichloroethylcarbonylamino)-2-deoxy-3,6-di-*O*-benzyl-4-*O*-fluorenylmethyl-1-thio-*α,β*-D-galactopyranoside (11)**

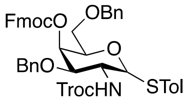


Azide **S15** (19.2 g, 26.89 mmol) was dissolved in THF (50 mL), followed by adding AcOH (5 mL). Zn dust (20.0 g) was then added, and the reaction mixture was stirred at RT for 1 h, after which it was filtered and concentrated to dryness. The resulting crude amine was dissolved in THF (20 mL), after which solid $NaHCO_3$ (10.0 g) and Troc-Cl (5.5 mL, 40.34 mmol) were added, and the reaction mixture was stirred at RT for further 10 mins. The reaction mixture was diluted with $CH_2Cl_2$ (100 mL) and washed with sat. $NaHCO_3$. The organic phase was dried and concentrated affording a solid, which was crystallized from EtOAc to give product as a white powder, (19.5 g, 84%). $^1$H NMR (400 MHz, $CDCl_3$): δ 2.27 (3H, s, $CH_3$ of STol), 2.30 (3H, s, $CH_3$ of STol), 3.56 – 3.72 (4H, m, H-3, H-2, H-6 x 2), 3.81 (1H, br t, H-5), 3.98 (1H, br d, H-3), 4.20 (1H, m, H of Fmoc), 4.29 (2H, m, CHHAr), 4.38 – 4.53 (6H, m, $CH_2$ of Fmoc, CHHAr, $CH_2$ of Troc), 4.56 (1H, m, H-2), 4.64 – 4.87 (4H, m, H-5, CHHAr x 3), 5.06 (1H, br d, H-1), 5.49 (1H, d, $J$ = 2.8 Hz, H-4), 5.54 (1H, br d, H-4), 5.69 (1H, d, $J$ = 5.1 Hz, H-1), 7.02 – 7.78 (22H, m, Ar-H). $^{13}$C NMR (101 MHz, $CDCl_3$) δ 154.9, 154.9, 154.0, 153.7, 143.6, 143.5, 143.1, 143.1, 141.3, 141.3, 141.2, 141.2, 138.1, 138.1, 137.7, 137.6, 137.2, 137.2, 132.8, 132.7, 129.9, 129.9, 129.7, 129.3, 128.9, 128.5, 128.4, 128.4, 128.4, 128.3, 128.1, 128.0, 127.9, 127.9, 127.9, 127.8, 127.8, 127.8, 127.8, 127.2, 127.2, 125.5, 125.4, 125.2, 125.1, 120.0, 120.0, 119.9, 105.0, 95.4, 89.5, 86.3, 78.5, 77.2, 75.9, 75.7, 75.6, 74.7, 74.4, 74.0, 73.7, 73.6, 71.5, 71.1, 70.4, 70.2, 70.2, 70.1, 69.1, 68.2, 68.1, 52.8, 51.3, 46.6, 46.5, 21.1, 21.1. MALDI-MS: $[M + Na]^+$ $C_{45}H_{42}Cl_3NNaO_8S$ calcd. 884.1594, found 884.1612.

**2-*O*-Acetyl-3,5-di-*O*-benzyl-6-*O*-fluorenylmethoxycarbonyl-*β*-D-galactofuranosyl-(1→1)-*t*-butyldiphenylsilyl 2,3,4-tri-*O*-benzyl-D-ribose (14)**

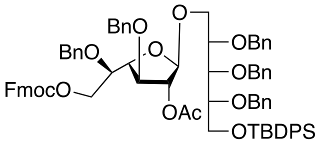


To a mixture of the donor **8** (9.0 g, 12.3 mmol), acceptor **7** (9.7 g, 14.7 mmol), NIS (4.1 g, 18.5 mmol), and activated molecular sieves in dry $CH_2Cl_2$ (50 mL) was added TMSOTf (222 µL, 1.23 mmol) at -78 °C. The reaction mixture was stirred at that temperature for 10 min, and then gradually warmed to -40 °C at which point it was quenched with pyridine. The reaction mixture was diluted with $CH_2Cl_2$ (50 mL) and washed with 10% $Na_2S_2O_3$. The organic layer was dried and concentrated to give a syrup which was chromatographed using EtOAc:Hexane (1:4) to give the pure product as a clear foam, (15.8 g, 95%). $^1$H NMR (400 MHz, $CDCl_3$): δ 1.03 (9H, s, $CH_3$ x 3 of TBDPS), 1.93 (3H, s, OAc), 3.65 (1H, dd, *J* = 9.9, 2.0 Hz, H-$5_a^{Rib}$), 3.71 – 3.81 (2H, m, H-$5^{Gal1}$,$H^{Rib}$), 3.84 (1H, d, *J* = 5.3 Hz, H-3), 3.87 – 3.94 (5H, m, H-$5_b^{Rib}$, H-$1^{Rib}$ x 2, $H^{Rib}$ x 2), 4.16 (1H, dd, *J* = 5.5, 3.6 Hz, H-$4^{Gal1}$), 4.19 – 4.24 (2H, m, H-$6_a^{Gal}$, CH of Fmoc), 4.27 – 4.40 (5H, m, H-$6_b^{Gal1}$, $CH_2$ of Fmoc, CHHAr x 2), 4.51 – 4.72 (8H, m, CHHAr), 4.97 (1H, s, H-$1^{Gal1}$), 5.07 (1H, s, H-$2^{Gal1}$), 7.1 – 7.77 (43H, m, ArH). $^{13}$C NMR (101 MHz, $CDCl_3$) δ 169.8, 154.9, 143.3, 141.2, 138.7, 138.7, 138.5, 137.8, 137.7, 135.7, 135.7, 135.6, 135.6, 133.6, 133.4, 129.5, 129.5, 128.4, 128.3, 128.3, 128.3, 128.2, 128.2, 128.2, 128.2, 128.0, 128.0, 127.9, 127.8, 127.8, 127.8, 127.7, 127.7, 127.6, 127.6, 127.6, 127.6, 127.3, 127.2, 127.2, 127.1, 127.1, 125.1, 120.0, 105.7, 83.0, 82.2, 81.0, 79.9, 78.6, 77.9, 77.2, 75.3, 73.6, 73.5, 72.5, 72.4, 72.1, 69.9, 68.1, 67.2, 63.7, 46.7, 26.9, 20.8, 19.2, 0.0. MALDI-MS: $[M + Na]^+$ $C_{79}H_{82}NaO_{13}Si$ calcd. 1289.5422, found 1289.5435.

**2-*O*-Acetyl-3,5-di-*O*-benzyl-*β*-D-galactofuranosyl-(1→1)-*t*-butyldiphenylsilyl 2,3,4-tri-*O*-benzyl-D-ribose (15)**

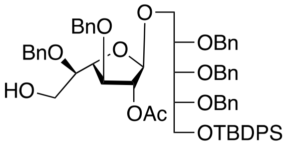


Disaccharide **14** (15.0 g, 11.8 mmol) was dissolved in $CH_2Cl_2$ (50 mL), after which $Et_3N$ (16 mL, 118.3 mmol) was added and the reaction mixture was stirred at RT overnight. It was then concentrated, loaded on silica gel and chromatographed using EtOAc:Hexane (3:7), (11.3 g, 91%). $^1$H NMR (400 MHz, $CDCl_3$): δ 1.03 (9H, s, $CH_3$ x 3 of TBDPS), 1.98 (3H, s, OAc), 3.52 (1H, m, H-$5^{Gal1}$), 3.60 (2H, m, H-$6^{Gal1}$), 3.64 (1H, m, H-$5_a^{Rib}$), 3.76 (1H, m, $H^{Rib}$), 3.86 (1H, d, *J* = 5.6 Hz, H-$3^{Gal1}$), 3.87 – 3.96 (5H, m, H-$1^{Rib}$, H-$5_b^{Rib}$, $H^{Rib}$ x 2), 4.20 (1H, dd, *J* = 4.1, 5.6 Hz, H-$4^{Gal}$), 4.39 (1H, d, *J* = 6.2 Hz, C*H*HAr), 4.42 (1H, d, *J* = 6.4 Hz, C*H*HAr), 4.50 – 4.60 (5H, m, C*H*HAr), 4.64 – 4.70 (4H, m, C*H*HAr), 4.94 (1H, s, H-$1^{Gal1}$), 5.08 (1H, d, *J* = 1.0 Hz, H-$2^{Gal1}$), 7.11 – 7.43 (30H, m, Ar-H), 7.63 (5H, m, Ar-H). $^{13}$C NMR (101 MHz, $CDCl_3$) δ 169.8, 138.7, 138.6, 138.5, 138.1, 137.6, 135.7, 135.6, 133.5, 133.3, 129.5, 129.5, 128.3, 128.3, 128.2, 128.2, 128.1, 128.0, 127.8, 127.8, 127.8, 127.7, 127.7, 127.6, 127.6, 127.6, 127.4, 127.3, 127.2, 105.6, 83.2, 82.9, 81.0, 79.8, 78.6, 78.0, 77.6, 77.2, 73.6, 72.8, 72.5, 72.4, 72.2, 67.2, 63.6, 62.1, 26.9, 20.8, 19.2, 0.0. MALDI-MS: $[M + Na]^+$ $C_{79}H_{82}NaO_{13}Si$ calcd. 1067.4742 found 1067.4756.

**4,6-Di-*O*-benzyl-2-*O*-levulinoyl-3-*O*-fluorenylmethyl-*β*-D-glucopyranosyl-(1→6)-2-*O*-acetyl-3,5-di-*O*-benzyl-*β*-D-galactofuranosyl-(1→1)-*t*-butyldiphenylsilyl 2,3,4-tri-*O*-benzyl-D-ribose (16)**

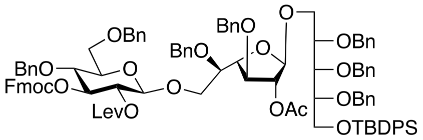


To a mixture of the donor **10** (2.5 g, 3.19 mmol), acceptor **15** (2.57 g, 2.45 mmol), NIS (715 mg, 3.19 mmol), and activated molecular sieves in dry $CH_2Cl_2$ (20 mL) was added TMSOTf (88 μL, 0.49 mmol) at -78 °C. The reaction mixture was stirred at that temperature for 10 min, and then gradually warmed to -40 °C at which point it was quenched with pyridine. The reaction mixture was diluted with $CH_2Cl_2$ (50 mL) and washed with 10% $Na_2S_2O_3$. The organic layer was dried and concentrated to give a syrup which was chromatographed using EtOAc:Hexane (1:4) to give the pure product as a clear foam, (3.0 g, 72%). $^1$H NMR (400 MHz, $CDCl_3$): δ 1.04 (9H, s, $CH_3$ x 3 of TBDPS), 1.87 (3H, s, OAc), 1.88 (3H, s, OLev), 2.23 – 2.54 (4H, m, $CH_2$ of Lev), 3.43 (1H, m, H-5$^{Glc}$), 3.63 (1H, m, H-5$_a^{Rib}$), 3.69 (2H, m, H-6$^{Gal1}$), 3.74 (2H, m, H-5$^{Gal1}$, H-6$_a^{Glc}$), 3.76 (1H, m, H-3$^{Gal1}$), 3.78 (1H, m, H$^{Rib}$), 3.86 (1H, m, H-4$^{Glc}$), 3.87 (1H, m, H-5$_b^{Rib}$), 3.89 (4H, m, H-1$^{Rib}$, H$^{Rib}$ x 2), 3.99 (1H, m, H-6$_b^{Glc}$), 4.00 (1H, m, H-4$^{Gal1}$), 4.19 (1H, d, *J* = 11.4 Hz, C*H*HAr), 4.23 – 4.25 (2H, m, CH of Fmoc, C*H*HAr), 4.30 (1H, d, *J* = 11.3 Hz, C*H*HAr), 4.45 (1H, d, H-1$^{Glc}$), 4.42 – 4.69 (14H, m, $CH_2$Ar, $CH_2$ of Fmoc, H-1$^{Glc}$), 4.95 (1H, s, H-1$^{Gal1}$), 5.02 (1H, s, H-2$^{Gal1}$), 5.06 (1H, m, H-3$^{Glc}$), 5.08 (1H, m, H-2$^{Glc}$), 7.00 – 7.76 (53H, m, Ar-H). $^{13}$C NMR (101 MHz, $CDCl_3$) δ 205.6, 171.4, 171.1, 169.8, 154.6, 143.6, 143.3, 141.2, 141.1, 138.7, 138.7, 138.5, 138.4, 137.8, 137.7, 137.5, 135.7, 135.6, 133.5, 133.4, 129.5, 129.5, 128.4, 128.3, 128.3, 128.3, 128.2, 128.2, 128.2, 128.2, 128.1, 128.0, 127.8, 127.8, 127.8, 127.8, 127.8, 127.8, 127.7, 127.7, 127.7, 127.6, 127.6, 127.6, 127.6, 127.4, 127.3, 127.2, 127.1, 125.3, 125.2, 119.9, 105.8, 100.8, 83.1, 83.0, 81.1, 79.9, 79.5, 78.6, 78.0, 77.2, 76.4, 75.7, 74.9, 74.6, 74.0, 73.6, 73.5, 72.5, 72.4, 72.1, 72.0, 71.8, 70.4, 68.0, 67.3, 63.7, 46.6, 37.6, 29.4, 27.8, 26.9, 21.0, 20.7, 19.2, 14.2, 0.0. MALDI-MS: [M + Na]$^+$ $C_{104}H_{110}NaO_{20}Si$ calcd. 1729.7257 found 1729.7269.

**4,6-Di-*O*-benzyl-2-*O*-levulinoyl-3-*β*-D-glucopyranosyl-(1→6)-2-*O*-acetyl-3,5-di-*O*-benzyl-*β*-D-galactofuranosyl-(1→1)-*t*-butyldiphenylsilyl 2,3,4-tri-*O*-benzyl-D-ribose (17)**

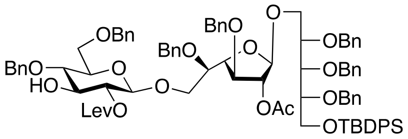


Trisaccharide **16** (2.8 g, 1.64 mmol) was dissolved in $CH_2Cl_2$ (20 mL), after which $Et_3N$ (1.14 mL, 8.19 mmol) was added and the reaction mixture was stirred at RT overnight. It was then concentrated, loaded on silica gel and chromatographed using EtOAc:Toluene (1:4), (2.1 g, 87%). $^1$H NMR (400 MHz, $CDCl_3$): δ 1.04 (9H, s, $CH_3$ x 3 of TBDPS), 1.85 (3H, s, OAc), 2.07 (3H, s, OLev) 2.20 (1H, m, C*H*H of OLev), 2.44 (1H, m, C*H*H of OLev), 2.52 (1H, m, C*H*H of OLev), 2.70 (1H, m, C*H*H of OLev), 3.07 (1H, d, *J* = 3.17 Hz, 3-OH$^{Glc}$), 3.39 (1H, m, H-5$^{Glc}$), 3.62 (1H, m, H-5$_a^{Rib}$), 3.63 (1H, m, H-4$^{Glc}$), 3.67 (2H, m, H-6$^{Gal1}$), 3.71 (2H, m, H-5$^{Gal1}$, H-6$_a^{Glc}$), 3.75 (1H, m, H-3$^{Gal1}$), 3.79 (1H, m, H-3$^{Glc}$), 3.84 (1H, m, H-5$_b^{Rib}$), 3.88 (4H, m, H-1$^{Rib}$, H$^{Rib}$ x 2), 3.99 (1H, m, H-6$_b^{Glc}$), 3.99 (1H, m, H-4$^{Gal1}$), 4.17 (1H, d, *J* = 11.6 Hz, C*H*HAr), 4.30 (1H, d, *J* = 11.7 Hz, C*H*HAr), 4.39 (1H, d, *J* = 7.8 Hz, H-1$^{Glc}$), 4.42 – 4.69 (11H, m, $CH_2$Ar), 4.87 (1H, d, *J* = 11.0 Hz), 4.91 (1H, m, H-2$^{Glc}$), 4.95 (1H, s, H-1$^{Gal1}$), 5.00 (1H, s, H-2$^{Gal1}$), 7.09 – 7.69 (45H, m, Ar-H). $^{13}$C NMR (101 MHz, $CDCl_3$) δ 207.8, 172.2, 169.8, 138.7, 138.6, 138.5, 138.5, 138.2, 137.9, 137.7, 135.7, 135.6, 133.5, 133.4, 129.5, 129.5, 129.0, 128.4, 128.4, 128.3, 128.3, 128.2, 128.2, 128.2, 128.1, 128.0, 127.8, 127.8, 127.8, 127.7, 127.7, 127.6, 127.6, 127.6, 127.6, 127.5, 127.5, 127.4, 127.3, 127.2, 127.2, 125.3, 105.8, 100.6, 83.0, 82.9, 81.1, 79.9, 78.6, 77.9, 77.2, 76.4, 76.1, 74.8, 74.8, 74.5, 74.0,

73.6, 73.5, 72.5, 72.4, 72.0, 68.5, 67.2, 63.7, 38.4, 29.6, 28.1, 26.9, 21.4, 20.7, 19.2, 0.0. MALDI-MS: [M + Na]+ $C_{89}H_{100}NaO_{18}Si$ calcd. 1507.6577 found 1507.6598.

**2-*O*-Levulinoyl-3,5-di-*O*-benzyl-6-*O*-fluorenylmethyl-*β*-D-galactofuranosyl-(1→3)-4,6-di-*O*-benzyl-2-*O*-levulinoyl-*β*-D-glucopyranosyl-(1→6)-2-*O*-acetyl-3,5-di-*O*-benzyl-*β*-D-galactofuranosyl-(1→1)-*t*-butyldiphenylsilyl 2,3,4-tri-*O*-benzyl-D-ribose (18)**

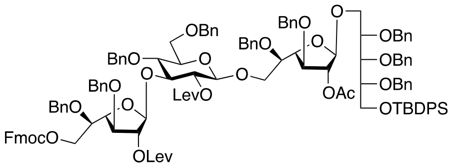


To a mixture of the donor **9** (1.6 g, 2.12 mmol), acceptor **17** (2.1 g, 1.41 mmol), NIS (480 mg, 2.12 mmol), and activated molecular sieves in dry $CH_2Cl_2$ (20 mL) was added TMSOTf (51 μL, 0.282 mmol) at -78 °C. The reaction mixture was stirred at that temperature for 10 min, and then gradually warmed to -40 °C at which point it was quenched with pyridine. The reaction mixture was diluted with $CH_2Cl_2$ (50 mL) and washed with 10% $Na_2S_2O_3$. The organic layer was dried and concentrated to give a syrup which was chromatographed using EtOAc:Toluene (5:95 to 1:9) to give the pure product as a clear foam, (2.6 g, 86%). $^1$H NMR (400 MHz, $CDCl_3$): δ 1.04 (9H, s, $CH_3$ x 3 of TBDPS), 1.86 (3H, s, OAc), 1.94 (3H, s, OLev), 2.09 (3H, s, OLev), 2.44 (4H, m, C*H*H of OLev), 2.62 (4H, m, C*H*H of OLev), 3.40 (1H, m, H-5$^{Glc}$), 3.59 (2H, m, incl. H-5$^{Gal2}$), 3.60 – 3.68 (m, H-5$_a^{Rib}$, H-6$^{Gal1}$), 3.70 (2H, m, H-6$_a^{Glc}$), 3.72 (1H, m, H-5$^{Gal1}$), 3.75 (1H, m, H-3$^{Gal1}$), 3.78 (m, H$^{Rib}$), 3.86 (1H, m, H-5$_b^{Rib}$), 3.88 (m, H-1$^{Rib}$, H$^{Rib}$, H-3$^{Gal2}$), 3.93 – 3.94 (m, H-3$^{Glc}$, H-4$^{Glc}$), 3.96 (m, H-6$_b^{Glc}$), 3.98 (1H, m, H-6$_a^{Gal2}$), 4.00 (1H, m, H-4$^{Gal1}$), 4.13 – 4.20 (m, H-6$_b^{Gal2}$, C*H*HAr), 4.20 (1H, m, CH of Fmoc), 4.22 (1H, m, H-4$^{Gal2}$), 4.29 (1H, d, C*H*HAr), 4.31 (1H, m, H-1$^{Glc}$), 4.32 – 4.60 (m, CH*H*Ar), 4.62 – 4.70 (4H, m, C*H*HAr), 4.88 (1H, d, *J* = 11.4 Hz, C*H*HAr), 4.95 (1H, s, H-1$^{Gal1}$), 5.0 (1H, s, H-2$^{Gal1}$), 5.08 (2H, m, H-2$^{Gal2}$, H-2$^{Glc}$), 5.31 (1H, s, H-1$^{Gal2}$), 7.04 – 7.80 (63H, Ar-H). $^{13}$C NMR (101 MHz, $CDCl_3$) δ 206.2, 205.9, 171.6, 171.2, 169.8, 154.9, 143.3, 141.3, 138.7, 138.6, 138.6, 138.5, 138.4, 138.0, 137.8, 137.7, 137.7, 135.7, 135.6, 133.6, 133.4, 129.5, 129.5, 128.3, 128.3, 128.3, 128.3, 128.2, 128.2, 128.2, 128.1, 128.0, 128.0, 128.0, 127.9, 127.8, 127.8, 127.8, 127.7, 127.7, 127.7, 127.7, 127.7, 127.6, 127.6, 127.6, 127.6, 127.6, 127.6, 127.5, 127.4, 127.3, 127.3, 127.2, 127.2, 127.1, 127.1, 125.1, 120.0, 105.8, 105.7, 100.9, 83.1, 83.1, 82.7, 82.6, 81.1, 80.7, 79.9, 78.6, 77.9, 77.2, 76.1, 75.4, 75.2, 74.4, 74.0, 73.8, 73.6, 73.4, 73.2, 72.5, 72.4, 72.1, 72.0, 71.8, 69.8, 68.6, 68.4, 67.2, 63.7, 46.7, 37.7, 37.5, 29.7, 29.5, 27.9, 27.8, 26.9, 20.7, 19.2, 0.0. MALDI-MS: [M + Na]+ $C_{129}H_{138}NaO_{27}Si$ calcd. 2169.9092 found 2169.9105

**2-*O*-Levulinoyl-3,5-di-*O*-benzyl-*β*-D-galactofuranosyl-(1→3)-4,6-di-*O*-benzyl-2-*O*-levulinoyl-*β*-D-glucopyranosyl-(1→6)-2-*O*-acetyl-3,5-di-*O*-benzyl-*β*-D-galactofuranosyl-(1→1)-*t*-butyldiphenylsilyl 2,3,4-tri-*O*-benzyl-D-ribose (19)**

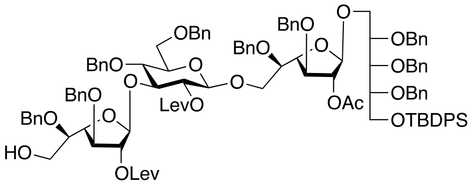


Tetrasaccharide **18** (2.6 g, 1.21 mmol) was dissolved in $CH_2Cl_2$ (20 mL), after which $Et_3N$ (1.7 mL, 12.1 mmol) was added and the reaction mixture was stirred at RT overnight. It was then concentrated, loaded on silica gel and chromatographed using EtOAc:Toluene (1:4), (1.9 g, 82%). $^1$H NMR (400 MHz, $CDCl_3$): δ 1.03 (9H, s, $CH_3$ x 3 of TBDPS), 1.86 (3H, s, OAc), 1.94 (3H, s, OLev), 2.09 (3H, s, OLev), 2.44 (4H, m, C*H*H of OLev), 2.64 (4H, m, C*H*H of OLev), 3.37 (1H, m, H-5$^{Gal2}$), 3.38 (1H, m, H-6$_a^{Gal2}$), 3.40 (1H, m, H-5$^{Glc}$), 3.47 (1H, m, H-6$_b^{Gal2}$), 3.58 – 3.67 (4H, m, H-6$^{Gal1}$, H-5$_a^{Rib}$), 3.70 (1H, m, H-6$^{Glc}$), 3.71 (1H, m, H-5$^{Gal1}$), 3.74 (1H, m, H-3$^{Gal1}$), 3.77 (1H, H$^{Rib}$), 3.86 (1H, m, H-5$_b^{Rib}$), 3.88 (m, H-1$^{Rib}$, H-4$^{Glc}$), 3.90 (1H, m, H-3$^{Gal2}$), 3.94 (1H, m, H-3$^{Glc}$), 3.96 (1H, m, H-6$_b^{Glc}$), 3.99 (1H, dd, *J* = 5.8, 2.2 Hz, H-4$^{Gal1}$), 4.16 – 4.23 (2H, m, C*H*HAr x 2), 4.26 (1H, dd, *J* = 3.9, 4.9 Hz, H-4$^{Gal2}$), 4.27 – 4.59 (11H, m, C*H*HAr), 4.31 (1H, m, H-1$^{Glc}$), 4.62 – 4.72 (4H, m,

C*H*HAr), 4.89 (1H, d, *J* = 11.4 Hz, C*H*HAr), 4.95 (1H, s, H-$1^{Gal1}$), 5.01 (1H, s, H-$2^{Gal1}$), 5.08 (1H, dd, *J* = 9.3, 8.2 Hz, H-$2^{Glc}$), 5.09 (1H, s, H-$2^{Gal2}$), 5.31 (1H, s, H-$1^{Gal2}$). $^{13}$C NMR (101 MHz, $CDCl_3$) δ 206.3, 206.0, 171.6, 171.2, 169.8, 138.7, 138.6, 138.6, 138.5, 138.3, 138.1, 137.8, 137.7, 137.6, 135.7, 135.6, 133.6, 133.4, 129.5, 129.5, 128.3, 128.3, 128.3, 128.3, 128.2, 128.2, 128.2, 128.2, 128.2, 128.2, 128.1, 128.1, 128.0, 128.0, 127.9, 127.9, 127.9, 127.8, 127.8, 127.8, 127.7, 127.7, 127.7, 127.7, 127.6, 127.6, 127.6, 127.6, 127.5, 127.4, 127.3, 127.3, 127.2, 105.8, 105.6, 100.9, 83.2, 83.1, 83.0, 82.8, 81.1, 80.8, 79.9, 78.6, 78.0, 77.2, 76.5, 76.1, 75.2, 74.5, 74.0, 73.8, 73.6, 73.4, 72.5, 72.4, 72.2, 72.0, 71.8, 68.4, 67.2, 63.7, 61.8, 37.8, 37.5, 29.7, 29.5, 27.8, 26.9, 20.7, 19.2, 0.0. MALDI-MS: [M + Na]$^+$ $C_{114}H_{128}NaO_{25}Si$ calcd. 1947.8412 found 1947.8427.

**2-(2,2,2-Trichloroethylcarbonylamino)-2-deoxy-3,6-di-*O*-benzyl-4-*O*-fluorenylmethyl-*β*-D-galactopyranosyl-(1→6)-2-*O*-levulinoyl-3,5-di-*O*-benzyl-*β*-D-galactofuranosyl-(1→3)-4,6-di-*O*-benzyl-2-*O*-levulinoyl-*β*-D-glucopyranosyl-(1→6)-2-*O*-acetyl-3,5-di-*O*-benzyl-*β*-D-galactofuranosyl-(1→1)-*t*-butyldiphenylsilyl 2,3,4-tri-*O*-benzyl-D-ribose (12)**

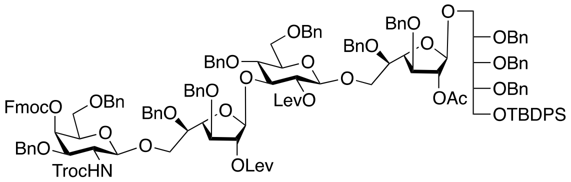


To a mixture of the donor **11** (8.4 g, 9.7 mmol), acceptor **19** (12.5 g, 6.49 mmol), NIS (2.1 g, 9.7 mmol), and activated molecular sieves in dry $CH_2Cl_2$ (100 mL) was added TMSOTf (235 μL, 1.298 mmol) at -50 °C. The reaction mixture was stirred at that temperature for 10 min, and then gradually warmed to -40 °C at which point it was quenched with pyridine. The reaction mixture was diluted with $CH_2Cl_2$ (50 mL) and washed with 10% $Na_2S_2O_3$. The organic layer was dried and concentrated to a syrup which was chromatographed using EtOAc:Toluene (1:4 to 3:7) to give the pure product as a white solid, (15.7 g, 91%). $^1$H NMR (400 MHz, $CDCl_3$): δ 1.03 (9H, s, $CH_3$ x 3 of TBDPS), 1.86 (3H, s, OAc), 1.94 (3H, s, OLev), 2.09 (3H, s, OLev), 2.44 (4H, m, C*H*H of OLev), 2.64 (4H, m, C*H*H of OLev), 3.37 (1H, m, H-$5^{Glc}$), 3.56 – 3.65 (8H, m, H-$2^{GalN}$, H-$6^{Gal1}$, H-$6^{Gal2}$, H-$6_a^{GalN}$), 3.70 (2H, m, H-$6_b^{GalN}$, H-$5^{Gal1}$), 3.73 – 3.81 (5H, m, H-$3^{GalN}$, H-$6_a^{Glc}$, H-$3^{Gal1}$, H-$3^{Gal2}$, $H^{Rib}$), 3.84 (1H, m, H-$5^{Rib}$), 3.88 (m, inc. H-$1^{Rib}$, H-$3^{Glc}$, H-$4^{Glc}$, $H^{Rib}$), 3.95 (1H, m, H-$6_b^{Glc}$), 3.99 (1H, dd, *J* = 5.4, 1.9 Hz, H-$4^{Gal1}$), 4.10 (1H, at, H-$4^{Gal2}$), 4.12 – 4.22 (3H, m, C*H*HAr, CH of Fmoc), 4.23 – 4.35 (m, C*H*HAr), 4.30 (1H, H-$1^{Glc}$), 4.37 – 4.60 (m, CHHAr), 4.49 (1H, H-$1^{GalN}$), 4.61 – 4.69 (3H, m, CHHAr), 4.74 (1H, d, *J* = 10.8 Hz, CHHAr), 4.88 (1H, d, *J* = 11.0 Hz, CHHAr), 4.95 (1H, s, H-$2^{Gal1}$), 5.01 (1H, s, H-$2^{Gal2}$), 5.06 (1H, dd, *J* = 8.2, 9.5 Hz, H-$2^{Glc}$), 5.30 (1H, s, H-$1^{Gal2}$), 5.46 (1H, d, *J* = 2.7 Hz, H-$4^{GalN}$), 7.04 – 7.79 (73H, m, Ar-H). $^{13}$C NMR (101 MHz, $CDCl_3$) δ 206.4, 206.0, 171.7, 171.2, 169.8, 155.0, 153.9, 143.6, 143.1, 141.3, 141.2, 138.7, 138.6, 138.6, 138.6, 138.5, 138.4, 137.9, 137.7, 137.7, 137.5, 137.4, 135.7, 135.6, 133.5, 133.4, 129.5, 129.5, 128.5, 128.3, 128.3, 128.3, 128.3, 128.3, 128.2, 128.2, 128.2, 128.2, 128.1, 128.1, 128.0, 128.0, 127.9, 127.9, 127.9, 127.8, 127.8, 127.7, 127.7, 127.7, 127.6, 127.6, 127.6, 127.5, 127.4, 127.4, 127.3, 127.2, 127.2, 127.2, 125.5, 125.2, 119.9, 105.9, 100.8, 83.4, 83.1, 82.9, 81.1, 80.9, 79.9, 78.6, 77.9, 77.2, 77.1, 76.6, 76.2, 75.1, 74.5, 74.0, 73.9, 73.7, 73.6, 73.4, 72.5, 72.4, 72.0, 71.8, 71.7, 71.4, 70.1, 69.7, 67.6, 67.2, 63.7, 54.4, 46.6, 37.7, 37.5, 29.7, 29.6, 27.9, 27.7, 26.9, 20.7, 19.2, 0.0. MALDI-MS: [M + Na]$^+$ $C_{152}H_{162}Cl_3NNaO_{33}Si$ calcd. 2684.9762 found 2684.9778.

**2-Acetamido-2-deoxy-3,6-di-*O*-benzyl-4-*O*-fluorenylmethyl-*β*-D-galactopyranosyl-(1→6)-2-*O*-levulinoyl-3,5-di-*O*-benzyl-*β*-D-galactofuranosyl-(1→3)-4,6-di-*O*-benzyl-2-*O*-levulinoyl-*β*-D-glucopyranosyl-(1→6)-2-*O*-acetyl-3,5-di-*O*-benzyl-*β*-D-galactofuranosyl-(1→1)-*t*-butyldiphenylsilyl 2,3,4-tri-*O*-benzyl-D-ribose (20)**

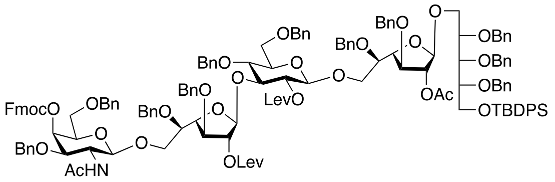


To a solution of pentasaccharide **12** (1.0 g, 0.37 mmol) in THF (5 mL) was added Zn dust (1.0 g), $Ac_2O$ (0.5 mL) and AcOH (0.5 mL), and the reaction mixture was left stirring at RT for 1 h, after which it was filtered, concentrated, absorbed on silica gel and purified using acetone:toluene (1:9 to 1:4) to give the product as a white solid (787 mg, 82%).$^1$H NMR (400 MHz, $CDCl_3$): δ 1.04 (9H, s, $CH_3$ x 3 of TBDPS), 1.63 (3H, s, NHAc), 1.86 (3H, s, OAc), 1.94 (3H, s, OLev), 2.09 (3H, s, OLev), 2.44 (4H, m, C*H*H of OLev), 2.64 (4H, m, C*H*H of OLev), 3.36 – 3.44 (2H, m, H-$5^{Glc}$, H-$2^{GalN}$), 3.53 – 3.67 (7H, m, H-$6^{Gal1}$, H-$6^{Gal2}$, H-$5^{Gal2}$, H-$5_a^{Rib}$, H-$6_a^{GalN}$), 3.70 (1H, m, H-$6_a^{Glc}$), 3.71 (1H, m, H-$5^{Gal1}$), 3.74 (1H, m, H-$3^{Gal1}$), 3.77 (3H, m, H-$5^{GalN}$, H-$3^{Gal2}$, $H^{Rib}$), 3.81 (1H, m, H-$6_b^{GalN}$), 3.84 (1H, m, H-$5_b^{Rib}$), 3.87 – 3.93 (4H, m, H-$3^{Glc}$, H-$4^{Glc}$, H-$1^{Rib}$), 3.96 (1H, m, H-$6_b^{Glc}$), 3.99 (1H, dd, $J$ = 2.3, 5.6 Hz, H-$4^{Gal1}$), 4.11 (1H, m, H-$4^{Gal2}$), 4.15 – 4.22 (3H, m, H-$3^{GalN}$, CH of Fmoc, C*H*HAr), 4.27 – 4.33 (3H, m, H-$1^{Glc}$, CH*H*Ar x 2), 4.33 – 4.70 (m, C*H*HAr), 4.74 (1H, d, $J$ = 11.6 Hz, C*H*HAr), 4.81 (1H, d, $J$ = 8.3 Hz, H-$1^{GalN}$), 4.92 (1H, d, $J$ = 11.6 Hz, C*H*HAr), 4.96 (1H, s, H-$1^{Gal1}$), 5.01 (1H, s, H-$2^{Gal1}$), 5.04 (1H, s, H-$2^{Gal2}$), 5.06 (1H, dd, $J$ = 8.4, 9.4 Hz, H-$2^{Glc}$), 5.19 (1H, d, $J$ = 7.8 Hz, NH), 5.31 (1H, s, H-$1^{Gal2}$), 5.46 (1H, d, $J$ = 2.5 Hz, H-$4^{GalN}$), 7.06 – 7.77 (73H, m, Ar-H). $^{13}$C NMR (101 MHz, $CDCl_3$) δ 206.4, 206.0, 171.7, 171.3, 170.4, 169.8, 155.1, 143.6, 143.2, 141.3, 141.2, 138.7, 138.6, 138.6, 138.6, 138.5, 138.5, 137.9, 137.8, 137.7, 137.7, 137.6, 135.7, 135.6, 135.6, 133.6, 133.4, 129.5, 129.5, 128.4, 128.4, 128.3, 128.3, 128.2, 128.2, 128.2, 128.2, 128.2, 128.1, 128.0, 128.0, 128.0, 127.9, 127.9, 127.8, 127.8, 127.8, 127.8, 127.7, 127.7, 127.7, 127.7, 127.6, 127.6, 127.6, 127.6, 127.5, 127.4, 127.3, 127.3, 127.2, 127.2, 127.1, 125.4, 125.2, 120.0, 119.9, 105.9, 105.8, 105.0, 100.8, 100.3, 83.3, 83.1, 83.0, 81.1, 79.9, 78.6, 77.9, 77.2, 76.6, 76.4, 75.1, 74.7, 74.5, 74.0, 73.9, 73.7, 73.6, 73.4, 73.1, 72.5, 72.4, 72.0, 71.7, 71.6, 70.8, 70.2, 70.1, 68.4, 67.7, 67.2, 63.7, 54.6, 46.6, 37.7, 37.5, 29.7, 29.6, 27.9, 27.7, 26.9, 23.4, 20.7, 19.2, 0.0. MALDI-MS: $[M + Na]^+$ $C_{151}H_{163}NNaO_{32}Si$ calcd. 2553.0825 found 2553.0843.

**2-Acetamido-2-deoxy-3,6-di-*O*-benzyl-4-*O*-fluorenylmethyl-*β*-D-galactopyranosyl-(1→6)-2-*O*-levulinoyl-3,5-di-*O*-benzyl-*β*-D-galactofuranosyl-(1→3)-4,6-di-*O*-benzyl-2-*O*-levulinoyl-*β*-D-glucopyranosyl-(1→6)-2-*O*-acetyl-3,5-di-*O*-benzyl-*β*-D-galactofuranosyl-(1→1)-2,3,4-tri-*O*-benzyl-D-ribose (21)**

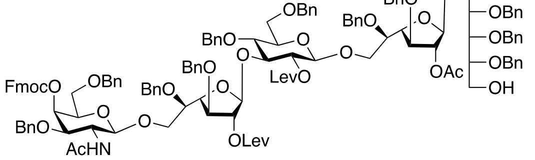


Pentasaccharide **20** (787 mg, 0.31 mmol) was dissolved in pyridine (6 mL) and cooled to 0 $^o$C, followed by adding HF-Py (4 mL). The reaction mixture was left stirring at RT for 30 min, after which it was diluted with $CH_2Cl_2$ (20 mL), washed with water (20 mL), and then with sat. $NaHCO_3$ (20 mL). The organic phase was dried and concentrated *in vacuo* to give a crude residue, which was chromatographed using acetone:toluene (1:4) to give the desired product as a white solid (680 mg, 86%).$^1$H NMR (400 MHz, $CDCl_3$): δ 1.63 (3H, s, NHAc), 1.86 (3H, s, OAc), 1.94 (3H, s, OLev), 2.09 (3H, s, OLev), 2.24 (1H, m, 1-$OH^{Rib}$), 2.44 (4H, m, C*H*H of OLev), 2.64 (4H, m, C*H*H of OLev), 3.36 – 3.44 (2H, m, H-$5^{Glc}$, H-$2^{GalN}$), 3.53 – 3.67 (7H, m, H-$6^{Gal1}$, H-$6^{Gal2}$, H-$5^{Gal2}$, H-$5_a^{Rib}$, H-$6_a^{GalN}$), 3.70 (1H, m, H-$6_a^{Glc}$), 3.71 (1H, m, H-$5^{Gal1}$), 3.74 (1H, m, H-

$3^{Gal1}$), 3.77 (3H, m, H-$5^{GalN}$, H-$3^{Gal2}$, $H^{Rib}$), 3.81 (1H, m, H-$6_b^{GalN}$), 3.84 (1H, m, H-$5_b^{Rib}$), 3.87 – 3.93 (4H, m, H-$3^{Glc}$, H-$4^{Glc}$, H-$1^{Rib}$), 3.96 (1H, m, H-$6_b^{Glc}$), 3.99 (1H, m, H-$4^{Gal1}$), 4.11 (1H, m, H-$4^{Gal2}$), 4.15 – 4.22 (3H, m, H-$3^{GalN}$, CH of Fmoc, C*H*HAr), 4.27 – 4.33 (3H, m, H-$1^{Glc}$, CH*H*Ar x 2), 4.33 – 4.70 (m, C*H*HAr), 4.74 (1H, d, *J* = 11.6 Hz, C*H*HAr), 4.81 (1H, d, *J* = 8.3 Hz, H-$1^{GalN}$), 4.92 (1H, d, *J* = 11.6 Hz, C*H*HAr), 4.96 (1H, s, H-$1^{Gal1}$), 5.01 (1H, s, H-$2^{Gal1}$), 5.04 (1H, s, H-$2^{Gal2}$), 5.06 (1H, dd, *J* = 8.4, 9.4 Hz, H-$2^{Glc}$), 5.19 (1H, d, *J* = 7.8 Hz, NH), 5.31 (1H, s, H-$1^{Gal2}$), 5.46 (1H, s, H-$4^{GalN}$), 7.03 – 7.79 (63H, m, Ar-H). $^{13}$C NMR (101 MHz, $CDCl_3$) δ 120.0, 125.5, 125.2, 128.0, 127.0, 70.3, 105.7, 81.2, 105.9, 74.6, 100.3, 71.8, 72.5, 73.9, 72.2, 72.0, 72.3, 72.2, 73.6, 74.5, 70.1, 73.7, 71.9, 100.9, 72.2, 70.1, 70.1, 72.2, 73.9, 73.4, 46.7, 74.9, 73.2, 83.4, 83.2, 71.6, 78.9, 66.7, 77.7, 70.6, 83.0, 71.7, 76.5, 78.9, 61.5, 71.6, 66.7, 68.4, 68.1, 70.9, 76.7, 75.3, 54.7, 37.8, 29.9, 29.8, 20.9, 23.6. MALDI-MS: $[M + Na]^+$ $C_{135}H_{145}NNaO_{32}$ calcd. 2314.9647 found 2316.4943.

**2-Acetamido-2-deoxy-3,6-di-*O*-benzyl-4-*O*-fluorenylmethyl-*β*-D-galactopyranosyl-(1→6)-2-*O*-levulinoyl-3,5-di-*O*-benzyl-*β*-D-galactofuranosyl-(1→3)-4,6-di-*O*-benzyl-2-*O*-levulinoyl-*β*-D-glucopyranosyl-(1→6)-2-*O*-acetyl-3,5-di-*O*-benzyl-*β*-D-galactofuranosyl-(1→1)-5-[5-benzyl(benzyloxycarbonyl)aminopentyl]-phosphoryl-2,3,4-tri-*O*-benzyl-D-ribose (13)**

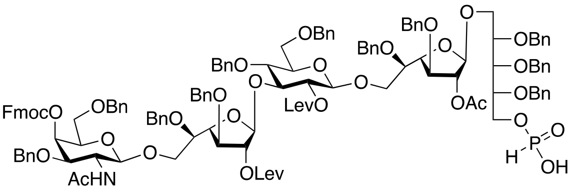


To a solution of pentasaccharide **21** (2.0 g, 0.87 mmol) in pyridine (10 mL), was added dioxane (5 mL), and after that he reaction was cooled to 0 °C, followed by adding salicyl chlorophosphite (883 mg, 4.35 mmol). Stirring was continued at RT for 2 h. A second portion of salicyl chlorophospite (883 mg, 4.35 mmol) was added to bring the reaction to completion. Water was then added, and the solution was then concentrated *in vacuo*, loaded on silica gel and chromatographed using $CH_3CN$, then $CH_3CN:CH_2Cl_2$ (1:1), and $CH_3OH:CH_2Cl_2:CH_3CN$ (1:4:5) to give the desired phosphonate as a white solid. (2.04 g, 89%). $^1$H NMR (400 MHz, $CDCl_3$): δ 1.65 (3H, s, NHAc), 1.84 (3H, s, OAc), 1.93 (3H, s, OLev), 2.09 (3H, s, OLev), 2.29 – 2.60 (8H, m, C*H*H of OLev), 3.36 – 3.44 (2H, m, H-$5^{Glc}$, H-$2^{GalN}$), 3.53 – 3.67 (7H, m, H-$6^{Gal1}$, H-$6^{Gal2}$, H-$5^{Gal2}$, H-$5_a^{Rib}$, H-$6_a^{GalN}$), 3.70 (1H, m, H-$6_a^{Glc}$), 3.71 (1H, m, H-$5^{Gal1}$), 3.74 (1H, m, H-$3^{Gal1}$), 3.77 (3H, m, H-$5^{GalN}$, H-$3^{Gal2}$, $H^{Rib}$), 3.81 (1H, m, H-$6_b^{GalN}$), 3.84 (1H, m, H-$5_b^{Rib}$), 3.87 – 3.93 (4H, m, H-$3^{Glc}$, H-$4^{Glc}$, H-$1^{Rib}$), 3.96 (1H, m, H-$6_b^{Glc}$), 3.99 (1H, m, H-$4^{Gal1}$), 4.11 (1H, m, H-$4^{Gal2}$), 4.15 – 4.22 (3H, m, H-$3^{GalN}$, CH of Fmoc, C*H*HAr), 4.27 – 4.33 (3H, m, H-$1^{Glc}$, CH*H*Ar x 2), 4.33 – 4.70 (m, C*H*HAr), 4.74 (1H, d, *J* = 11.6 Hz, C*H*HAr), 4.81 (1H, d, *J* = 8.3 Hz, H-$1^{GalN}$), 4.92 (1H, d, *J* = 11.6 Hz, C*H*HAr), 4.96 (1H, s, H-$1^{Gal1}$), 5.01 (1H, s, H-$2^{Gal1}$), 5.04 (1H, s, H-$2^{Gal2}$), 5.06 (1H, dd, *J* = 8.4, 9.4 Hz, H-$2^{Glc}$), 5.19 (1H, d, *J* = 7.8 Hz, NH), 5.31 (1H, s, H-$1^{Gal2}$), 5.46 (1H, s, H-$4^{GalN}$), 5.90 (1H, br s, H-phosphonate), 7.03 – 7.79 (63H, m, Ar-H). $^{13}$C NMR (101 MHz, $CDCl_3$) δ 206.4, 205.9, 183.0, 171.7, 171.2, 170.4, 169.9, 155.1, 143.6, 143.2, 141.3, 141.2, 138.5, 138.5, 138.2, 138.1, 138.1, 137.9, 137.8, 137.7, 137.6, 128.4, 128.4, 128.4, 128.3, 128.3, 128.3, 128.2, 128.2, 128.2, 128.2, 128.1, 128.0, 128.0, 128.0, 127.9, 127.9, 127.8, 127.7, 127.7, 127.7, 127.6, 127.6, 127.5, 127.5, 127.3, 127.2, 127.1, 125.4, 125.2, 120.0, 106.0, 105.8, 100.8, 100.3, 83.1, 83.0, 81.2, 78.9, 78.8, 77.6, 77.2, 75.1, 74.7, 74.5, 74.0, 73.9, 73.8, 73.7, 73.4, 73.1, 72.5, 72.1, 72.0, 71.9, 71.7, 71.6, 70.2, 70.0, 68.5, 67.7, 66.6, 61.4, 54.6, 46.6, 37.7, 37.5, 29.7, 29.6, 27.9, 27.7, 23.4, 20.7. MALDI-MS: $[M + Na]^+$ $C_{135}H_{146}NNaO_{34}P$ calcd. 2378.9362 found 2378.9464.

**2-Acetamido-2-deoxy-3,6-di-*O*-benzyl-4-*O*-fluorenylmethyl-*β*-D-galactopyranosyl-(1→6)-2-*O*-levulinoyl-3,5-di-*O*-benzyl-*β*-D-galactofuranosyl-(1→3)-4,6-di-*O*-benzyl-2-*O*-levulinoyl-3-*O*-*β*-D-glucopyranosyl-(1→6)-2-*O*-acetyl-3,5-di-*O*-benzyl-*β*-D-galactofuranosyl-(1→1)-5-[5-benzyl(benzyloxycarbonyl)aminopentyl]-phosphoryl-2,3,4-tri-*O*-benzyl-D-ribose (22a)**

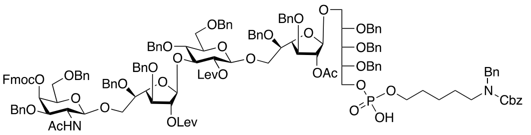


To a solution of the phosphonate **13** (500 mg, 0.212 mmol) and linker (347 mg, 1.06 mmol) in pyridine (10 mL ) was added PivCl (104 μL, 0.848 mmol), and the reaction mixture was left stirring at that temperature for 2 h. $I_2$ (538 mg, 2.1 mmol) and $H_2O$ (0.5 mL) were added, and the reaction mixture was additionally stirred at RT for 2 h. The reaction mixture was neutralized with sat. $Na_2S_2O_3$, concentrated *in vacuo*, re-dissolved in $CH_2Cl_2$, and washed with water. The organic phase was dried and concentrated *in vacuo* to give the residue which was further chromatographed to give the desired product as a white solid (500 mg, 88%). $^1$H NMR (750 MHz, $CDCl_3$): δ 1.15 – 1.39 (4H, br m, $CH_2$ x 2 linker), 1.65 (3H, s, NHAc), 1.84 (3H, s, OAc), 1.93 (3H, s, OLev), 2.09 (3H, s, OLev), 2.29 – 2.60 (8H, m, C*H*H of OLev), 2.87 – 3.04 (4H, m, $CH_2$ x 2 linker), 3.36 – 3.44 (2H, m, H-5$^{Glc}$, H-2$^{GalN}$), 3.53 – 3.67 (7H, m, H-6$^{Gal1}$, H-6$^{Gal2}$, H-5$^{Gal2}$, H-5$_a^{Rib}$, H-6$_a^{GalN}$), 3.70 (1H, m, H-6$_a^{Glc}$), 3.71 (1H, m, H-5$^{Gal1}$), 3.74 (1H, m, H-3$^{Gal1}$), 3.77 (3H, m, H-5$^{GalN}$, H-3$^{Gal2}$, H$^{Rib}$), 3.81 (1H, m, H-6$_b^{GalN}$), 3.84 (1H, m, H-5$_b^{Rib}$), 3.87 – 3.93 (4H, m, H-3$^{Glc}$, H-4$^{Glc}$, H-1$^{Rib}$), 3.96 (1H, m, H-6$_b^{Glc}$), 3.99 (1H, m, H-4$^{Gal1}$), 4.11 (1H, m, H-4$^{Gal2}$), 4.15 – 4.22 (3H, m, H-3$^{GalN}$, CH of Fmoc, C*H*HAr), 4.27 – 4.33 (3H, m, H-1$^{Glc}$, CH*H*Ar x 2), 4.33 – 4.70 (m, C*H*HAr), 4.74 (1H, d, *J* = 11.6 Hz, C*H*HAr), 4.81 (1H, d, *J* = 8.3 Hz, H-1$^{GalN}$), 4.92 (1H, d, *J* = 11.6 Hz, C*H*HAr), 4.96 (1H, s, H-1$^{Gal1}$), 5.01 (1H, s, H-2$^{Gal1}$), 5.04 (1H, s, H-2$^{Gal2}$), 5.06 (1H, dd, *J* = 8.4, 9.4 Hz, H-2$^{Glc}$), 5.19 (1H, d, *J* = 7.8 Hz, NH), 5.31 (1H, s, H-1$^{Gal2}$), 5.46 (1H, s, H-4$^{GalN}$), 5.90 (1H, br s, H-phosphonate), 7.03 – 7.79 (83H, m, Ar-H). $^{13}$C NMR (189 MHz, $CDCl_3$) δ 108.7, 108.6, 103.6, 103.0, 86.1, 85.8, 85.7, 83.8, 83.7, 80.8, 79.6, 79.5, 79.2, 79.1, 77.7, 77.4, 77.2, 76.7, 76.5, 76.3, 76.1, 75.8, 75.7, 74.7, 74.4, 74.3, 73.5, 72.9, 72.7, 71.1, 70.4, 69.9, 69.8, 67.9, 57.2, 53.0, 50.0, 49.3, 48.8, 40.4, 40.3, 40.2. ESI-MS: [M + H]$^-$ $C_{155}H_{169}N_2O_{37}P$ calcd. 2681.1142 found 2681.1072.

**2-Acetamido-2-deoxy-3,6-di-*O*-benzyl-*β*-D-galactopyranosyl-(1→6)-2-*O*-levulinoyl-3,5-di-*O*-benzyl-*β*-D-galactofuranosyl-(1→3)-4,6-di-*O*-benzyl-2-*O*-levulinoyl-*β*-D-glucopyranosyl-(1→6)-2-*O*-acetyl-3,5-di-*O*-benzyl-*β*-D-galactofuranosyl-(1→1)-5-[5-benzyl(benzyloxycarbonyl)aminopentyl]-phosphoryl-2,3,4-tri-*O*-benzyl-D-ribose (22)**

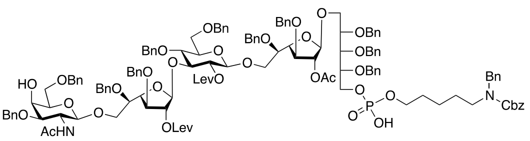


To a solution of pentasaccharide **22a** (500 mg, 0.18 mmol) in $CH_2Cl_2$ (10 mL) was added $Et_3N$ (0.5 mL) and the reaction mixture was left stirring at RT overnight. The mixture was concentrated, absorbed on silica gel, and chromatographed using $CH_3OH$:$CH_2Cl_2$ (1:9) to give the desired product as a white solid in quantitative yield.
(440 mg, quant). $^1$H NMR (750 MHz, $CDCl_3$): δ 1.15 – 1.39 (4H, br m, $CH_2$ x 2 linker), 1.65 (3H, s, NHAc), 1.84 (3H, s, OAc), 1.93 (3H, s, OLev), 2.09 (3H, s, OLev), 2.29 – 2.60 (8H, m, C*H*H of OLev), 2.87 – 3.04 (4H, m, $CH_2$ x 2 linker), 3.36 – 3.44 (2H, m, H-5$^{Glc}$, H-2$^{GalN}$), 3.53 – 3.67 (7H, m, H-6$^{Gal1}$, H-6$^{Gal2}$, H-5$^{Gal2}$, H-5$_a^{Rib}$, H-6$_a^{GalN}$), 3.70 (1H, m, H-6$_a^{Glc}$), 3.71 (1H, m, H-5$^{Gal1}$), 3.74 (1H, m, H-3$^{Gal1}$), 3.77 (3H, m, H-5$^{GalN}$, H-3$^{Gal2}$, H$^{Rib}$), 3.81 (1H, m, H-6$_b^{GalN}$), 3.84 (1H, m, H-5$_b^{Rib}$), 3.87 – 3.93 (4H, m, H-3$^{Glc}$, H-4$^{Glc}$, H-1$^{Rib}$), 3.96 (1H, m, H-6$_b^{Glc}$), 3.99 (1H, m, H-4$^{Gal1}$), 4.11 (1H, m, H-4$^{Gal2}$), 4.15 – 4.22 (2H, m, H-3$^{GalN}$, C*H*HAr), 4.27 – 4.33 (5H,

m, H-1$^{Glc}$, $CH_2$NBnCbz, CH*H*Ar x 2), 4.33 – 4.70 (m, C*H*HAr), 4.74 (1H, d, *J* = 11.6 Hz, C*H*HAr), 4.83 (1H, br d, H-1$^{GalN}$), 4.92 (1H, d, *J* = 11.6 Hz, C*H*HAr), 4.96 (1H, s, H-1$^{Gal1}$), 5.01 (1H, s, H-2$^{Gal1}$), 5.03 (2H, m, $CH_2$ of Cbz), 5.04 (1H, s, H-2$^{Gal2}$), 5.06 (1H, dd, *J* = 8.4, 9.4 Hz, H-2$^{Glc}$), 5.25 (1H, br d, NH), 5.31 (1H, s, H-1$^{Gal2}$), 7.03 – 7.79 (75H, m, Ar-H). $^{13}$C NMR (189 MHz, $CDCl_3$) δ 108.7, 108.5, 103.5, 102.9, 85.9, 85.8, 85.7, 83.8, 80.9, 79.7, 79.5, 79.1, 77.7, 77.2, 76.7, 76.5, 76.4, 76.3, 76.1, 76.0, 75.9, 75.7, 75.5, 74.8, 74.7, 74.3, 74.2, 72.8, 71.6, 71.5, 71.4, 71.1, 69.8, 68.2, 67.7, 56.3, 52.8, 49.9, 48.8, 40.4, 40.3, 40.2. ESI-MS: [M + H]$^-$ $C_{140}H_{159}N_2O_{35}P$ calcd. 2549.0461 found 2459.0378.

**Compound 23**

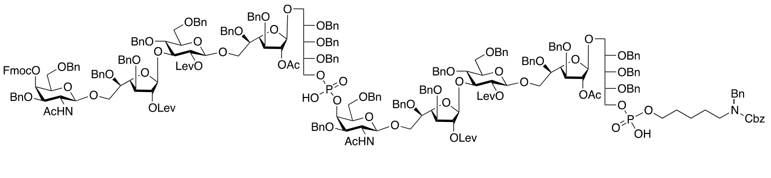

To a solution of the phosphonate **13** (330 mg, 0.14 mmol) and alcohol **22** (295 mg, 0.12 mmol) in pyridine (5 mL) was added PivCl (88 μL, 0.72 mmol), and the reaction mixture was left stirring at that temperature for 2 h. $I_2$ (538 mg, 2.1 mmol) and $H_2O$ (0.5 mL) were added, and the reaction mixture was additionally stirred at RT for 2 h. The reaction mixture was neutralized with sat. $Na_2S_2O_3$, concentrated *in vacuo*, re-dissolved in $CH_2Cl_2$, and washed with water. The organic phase was dried and concentrated *in vacuo* to give the residue which was further chromatographed to give the desired product as a white solid (271 mg, 46%). Purification conditions: 1) 100% $CH_3CN$; 2) $CH_2Cl_2$:$CH_3CN$ (1:1); 3) 3:47:50 $CH_3OH$:$CH_2Cl_2$:$CH_3CN$; 4) $CH_3OH$:$CH_2Cl_2$ (1:9). $^1$H NMR (750 MHz, $CDCl_3$): δ 1.15 – 1.39 (4H, br m, $CH_2$ x 2 linker), 1.65 (6H, s, NHAc x 2), 1.84 (6H, s, OAc x 2), 1.93 (6H, s, OLev x 2), 2.09 (6H, s, OLev x 2), 2.29 – 2.60 (16H, m, C*H*H of OLev), 2.87 – 3.04 (4H, m, $CH_2$ x 2 linker), 3.36 – 3.44 (4H, m, 2 x H-5$^{Glc}$, 2 x H-2$^{GalN}$), 3.53 – 3.67 (14H, m, 2 x H-6$^{Gal1}$, 2 x H-6$^{Gal2}$, 2 x H-5$^{Gal2}$, 2 x H-5$_a^{Rib}$, H-6$_a^{GalN}$), 3.70 (2H, m, 2 x H-6$_a^{Glc}$), 3.71 (2H, m, 2 x H-5$^{Gal1}$), 3.74 (2H, m, 2 x H-3$^{Gal1}$), 3.77 (6H, m, 2 x H-5$^{GalN}$, 2 x H-3$^{Gal2}$, 2 x H$^{Rib}$), 3.81 (2H, m, 2 x H-6$_b^{GalN}$), 3.84 (2H, m, 2 x H-5$_b^{Rib}$), 3.87 – 3.93 (8H, m, 2 x H-3$^{Glc}$, 2 x H-4$^{Glc}$, H-1$^{Rib}$), 3.96 (2H, m, 2 x H-6$_b^{Glc}$), 3.99 (2H, m, 2 x H-4$^{Gal1}$), 4.11 (2H, m, 2 x H-4$^{Gal2}$), 4.15 – 4.22 (4H, m, 2 x H-3$^{GalN}$, CH of Fmoc, 1 x C*H*HAr), 4.27 – 4.33 (10H, m, 2 x H-1$^{Glc}$, $CH_2$NBnCbz, CH*H*Ar x 2), 4.33 – 4.70 (m, C*H*HAr), 4.74 (1H, d, *J* = 11.6 Hz, C*H*HAr), 4.83 (2H, br d, 2 x H-1$^{GalN}$), 4.92 (1H, d, *J* = 11.6 Hz, C*H*HAr), 4.96 (2H, s, 2 x H-1$^{Gal1}$), 5.01 (2H, s, 2 x H-2$^{Gal1}$), 5.03 (2H, m, $CH_2$ of Cbz), 5.04 (2H, s, 2 x H-2$^{Gal2}$), 5.06 (2H, dd, 2 x H-2$^{Glc}$), 5.22 (2H, br d, 2 x NH), 5.31 (2H, s, 2 x H-1$^{Gal2}$), 5.46 (2H, s, 2 x H-4$^{GalN}$), 7.03 – 7.79 (128H, m, Ar-H). $^{13}$C NMR (189 MHz, $CDCl_3$) δ 108.7, 108.5, 103.0, 86.1, 85.7, 83.7, 79.9, 79.5, 79.3, 79.1, 77.7, 77.5, 77.1, 76.7, 76.5, 76.3, 76.2, 76.1, 75.8, 75.7, 74.7, 74.5, 74.4, 74.3, 73.5, 72.9, 72.7, 71.1, 70.4, 69.6, 68.2, 57.2, 52.9, 52.7, 49.3, 40.3, 40.2. ESI-MS: [M + H]$^-$ $C_{275}H_{303}N_3O_{69}P_2$ calcd. 2406.4884 found 2406.4820.

**Compound 24**

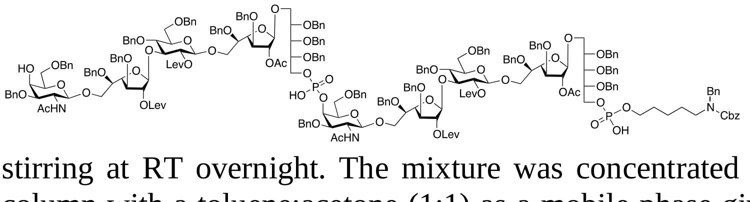

To a solution of decasaccharide **23** (271 mg, 0.056 mmol) in $CH_2Cl_2$ (5 mL) was added $Et_3N$ (0.5 mL) and the reaction mixture was left stirring at RT overnight. The mixture was concentrated and passed through a Biogel SX1 column with a toluene:acetone (1:1) as a mobile phase giving the desired product as a white solid. (250 mg, 96%). $^1$H NMR (750 MHz, $CDCl_3$): δ 1.15 – 1.39 (4H, br m, $CH_2$ x 2 linker), 1.65 (6H, s, NHAc x 2), 1.84 (6H, s, OAc x 2), 1.93 (6H, s, OLev x 2), 2.09 (6H, s, OLev x 2),

2.29 – 2.60 (16H, m, C*H*H of OLev), 2.87 – 3.04 (4H, m, $CH_2$ x 2 linker), 3.36 – 3.44 (4H, m, 2 x H-5$^{Glc}$, 2 x H-2$^{GalN}$), 3.53 – 3.67 (14H, m, 2 x H-6$^{Gal1}$, 2 x H-6$^{Gal2}$, 2 x H-5$^{Gal2}$, 2 x H-5$_a^{Rib}$, H-6$_a^{GalN}$), 3.70 (2H, m, 2 x H-6$_a^{Glc}$), 3.71 (2H, m, 2 x H-5$^{Gal1}$), 3.74 (2H, m, 2 x H-3$^{Gal1}$), 3.77 (6H, m, 2 x H-5$^{GalN}$, 2 x H-3$^{Gal2}$, 2 x H$^{Rib}$), 3.81 (2H, m, 2 x H-6$_b^{GalN}$), 3.84 (2H, m, 2 x H-5$_b^{Rib}$), 3.87 – 3.93 (8H, m, 2 x H-3$^{Glc}$, 2 x H-4$^{Glc}$, H-1$^{Rib}$), 3.96 (2H, m, 2 x H-6$_b^{Glc}$), 3.99 (2H, m, 2 x H-4$^{Gal1}$), 4.11 (2H, m, 2 x H-4$^{Gal2}$), 4.15 – 4.22 (3H, m, 2 x H-3$^{GalN}$, 1 x C*H*HAr), 4.27 – 4.33 (10H, m, 2 x H-1$^{Glc}$, $CH_2$NBnCbz, CH*H*Ar x 2), 4.33 – 4.70 (m, C*H*HAr), 4.74 (1H, d, *J* = 11.6 Hz, C*H*HAr), 4.83 (2H, br d, 2 x H-1$^{GalN}$), 4.92 (1H, d, *J* = 11.6 Hz, C*H*HAr), 4.96 (2H, s, 2 x H-1$^{Gal1}$), 5.01 (2H, s, 2 x H-2$^{Gal1}$), 5.03 (2H, m, $CH_2$ of Cbz), 5.04 (2H, s, 2 x H-2$^{Gal2}$), 5.06 (2H, dd, 2 x H-2$^{Glc}$), 5.22 (2H, br d, 2 x NH), 5.31 (2H, s, 2 x H-1$^{Gal2}$), 5.46 (2H, s, 2 x H-4$^{GalN}$), 7.03 – 7.79 (120H, m, Ar-H). $^{13}$C NMR (189 MHz, $CDCl_3$) δ 108.6, 108.4, 103.6, 102.9, 85.9, 85.8, 85.7, 83.9, 83.8, 81.3, 80.7, 79.6, 79.4, 79.3, 79.1, 77.7, 77.2, 76.7, 76.5, 76.4, 76.3, 76.1, 76.0, 75.7, 75.5, 75.1, 74.8, 74.7, 74.4, 74.2, 73.2, 72.8, 71.6, 71.1, 69.9, 69.8, 68.2, 68.1, 67.7, 62.2, 56.4, 55.2, 53.0, 52.8, 49.7, 48.8, 47.8, 40.4, 40.3, 40.2. ESI-MS: [M + H]$^-$ $C_{260}H_{293}N_3NaO_{67}P_2$ calcd. 2295.4544 found 2295.4433.

**Compound 25**

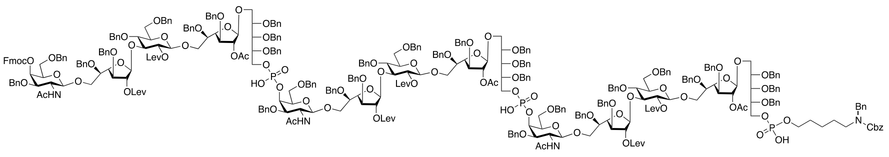

To a solution of the phosphonate **13** (110 mg, 0.047 mmol) and decasaccharide **24** (144 mg, 0.03 mmol) in pyridine (5 mL) was added PivCl (40 µL, 0.72 mmol), and the reaction mixture was left stirring at that temperature for 4 h. $I_2$ (80 mg, 0.313 mmol) and $H_2O$ (0.5 mL) were added, and the reaction mixture was additionally stirred at RT for 2 h. The reaction mixture was neutralized with sat. $Na_2S_2O_3$, concentrated *in vacuo,* re-dissolved in $CH_2Cl_2$, and washed with water. The organic phase was dried and concentrated *in vacuo* to give the residue which was further chromatographed to give the desired product as a white solid (89 mg, 41%). Purification conditions: 1) 100% $CH_3CN$; 2) $CH_2Cl_2$:$CH_3CN$ (1:1); 3) 2:48:50 $CH_3OH$:$CH_2Cl_2$:$CH_3CN$; 4) 1:4:5 $CH_3OH$:$CH_2Cl_2$:$CH_3CN$. $^1$H NMR (750 MHz, $CDCl_3$): δ 1.15 – 1.39 (4H, br m, $CH_2$ x 2 linker), 1.65 (9H, s, NHAc x 3), 1.84 (9H, s, OAc x 3), 1.93 (9H, s, OLev x 3), 2.09 (9H, s, OLev x 3), 2.29 – 2.60 (24H, m, C*H*H of OLev), 2.87 – 3.04 (4H, m, $CH_2$ x 2 linker), 3.36 – 3.44 (6H, m, 3 x H-5$^{Glc}$, 3 x H-2$^{GalN}$), 3.53 – 3.67 (13H, m, 3 x H-6$^{Gal1}$, 3 x H-6$^{Gal2}$, 3 x H-5$^{Gal2}$, 3 x H-5$_a^{Rib}$, H-6$_a^{GalN}$), 3.70 (3H, m, 3 x H-6$_a^{Glc}$), 3.71 (3H, m, 3 x H-5$^{Gal1}$), 3.74 (3H, m, 3 x H-3$^{Gal1}$), 3.77 (9H, m, 3 x H-5$^{GalN}$, 3 x H-3$^{Gal2}$, 3 x H$^{Rib}$), 3.81 (3H, m, 3 x H-6$_b^{GalN}$), 3.84 (3H, m, 3 x H-5$_b^{Rib}$), 3.87 – 3.93 (7H, m, 3 x H-3$^{Glc}$, 3 x H-4$^{Glc}$, H-1$^{Rib}$), 3.96 (3H, m, 3 x H-6$_b^{Glc}$), 3.99 (3H, m, 3 x H-4$^{Gal1}$), 4.11 (3H, m, 3 x H-4$^{Gal2}$), 4.15 – 4.22 (5H, m, 3 x H-3$^{GalN}$, CH of Fmoc, 1 x C*H*HAr), 4.27 – 4.33 (16H, m, 3 x H-1$^{Glc}$, $CH_2$NBnCbz, CH*H*Ar x 2), 4.33 – 4.70 (m, C*H*HAr), 4.74 (1H, d, C*H*HAr), 4.83 (3H, br d, 3 x H-1$^{GalN}$), 4.92 (1H, d, C*H*HAr), 4.96 (3H, s, 3 x H-1$^{Gal1}$), 5.01 (3H, s, 3 x H-2$^{Gal1}$), 5.03 (2H, m, $CH_2$ of Cbz), 5.04 (3H, s, 3 x H-2$^{Gal2}$), 5.06 (3H, dd, 3 x H-2$^{Glc}$), 5.22 (3H, br d, 3 x NH), 5.31 (3H, s, 3 x H-1$^{Gal2}$), 5.46 (3H, s, 3 x H-4$^{GalN}$), 7.03 – 7.79 (183H, m, Ar-H). $^{13}$C NMR (189 MHz, $CDCl_3$) δ 108.7, 108.5, 103.0, 86.1, 85.7, 83.7, 79.9, 79.5, 79.3, 79.1, 77.7, 77.5, 77.1, 76.7, 76.5, 76.3, 76.2, 76.1, 75.8, 75.7, 74.7, 74.5, 74.4, 74.3, 73.5, 72.9, 72.7, 71.1, 70.4, 69.6, 68.2, 57.2, 52.9, 52.7, 49.3, 40.3, 40.2. MALDI-MS: [M + Na]$^+$ $C_{395}H_{437}N_4NaO_{101}P_3$ calcd. 6967.8293 found 6967.1736.

### Compound 26

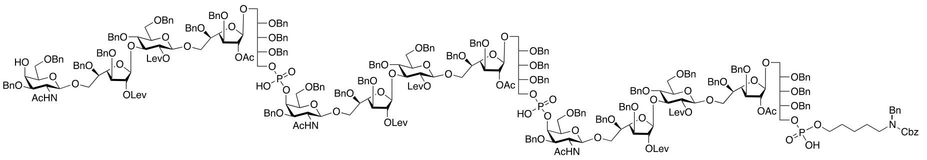

To a solution of pentadecasaccharide **25** (89 mg, 0.013 mmol) in $CH_2Cl_2$ (5 mL) was added $Et_3N$ (0.5 mL) and the reaction mixture was left stirring at RT overnight. The mixture was concentrated and passed through a Biogel SX1 column with a toluene:acetone (1:1) as a mobile phase giving the desired product as a white solid. (80 mg, 92%). $^{1}$H NMR (750 MHz, $CDCl_3$): δ 1.15 – 1.39 (4H, br m, $CH_2$ x 2 linker), 1.65 (9H, s, NHAc x 3), 1.84 (9H, s, OAc x 3), 1.93 (9H, s, OLev x 3), 2.09 (9H, s, OLev x 3), 2.29 – 2.60 (24H, m, C*H*H of OLev), 2.87 – 3.04 (4H, m, $CH_2$ x 2 linker), 3.36 – 3.44 (6H, m, 3 x H-5$^{Glc}$, 3 x H-2$^{GalN}$), 3.53 – 3.67 (13H, m, 3 x H-6$^{Gal1}$, 3 x H-6$^{Gal2}$, 3 x H-5$^{Gal2}$, 3 x H-5$_a^{Rib}$, H-6$_a^{GalN}$), 3.70 (3H, m, 3 x H-6$_a^{Glc}$), 3.71 (3H, m, 3 x H-5$^{Gal1}$), 3.74 (3H, m, 3 x H-3$^{Gal1}$), 3.77 (9H, m, 3 x H-5$^{GalN}$, 3 x H-3$^{Gal2}$, 3 x H$^{Rib}$), 3.81 (3H, m, 3 x H-6$_b^{GalN}$), 3.84 (3H, m, 3 x H-5$_b^{Rib}$), 3.87 – 3.93 (7H, m, 3 x H-3$^{Glc}$, 3 x H-4$^{Glc}$, H-1$^{Rib}$), 3.96 (3H, m, 3 x H-6$_b^{Glc}$), 3.99 (3H, m, 3 x H-4$^{Gal1}$), 4.11 (3H, m, 3 x H-4$^{Gal2}$), 4.15 – 4.22 (4H, m, 3 x H-3$^{GalN}$, 1 x C*H*HAr), 4.27 – 4.33 (16H, m, 3 x H-1$^{Glc}$, $CH_2$NBnCbz, CH*H*Ar x 2), 4.33 – 4.70 (m, C*H*HAr), 4.74 (1H, d, C*H*HAr), 4.83 (3H, br d, 3 x H-1$^{GalN}$), 4.92 (1H, d, C*H*HAr), 4.96 (3H, s, 3 x H-1$^{Gal1}$), 5.01 (3H, s, 3 x H-2$^{Gal1}$), 5.03 (2H, m, $CH_2$ of Cbz), 5.04 (3H, s, 3 x H-2$^{Gal2}$), 5.06 (3H, dd, 3 x H-2$^{Glc}$), 5.22 (3H, br d, 3 x NH), 5.31 (3H, s, 3 x H-1$^{Gal2}$), 5.46 (3H, s, 3 x H-4$^{GalN}$), 7.03 – 7.79 (175H, m, Ar-H). $^{13}$C NMR (189 MHz, $CDCl_3$) δ 108.6, 108.5, 103.6, 102.9, 85.9, 85.8, 85.8, 85.7, 83.8, 79.8, 79.7, 79.6, 79.5, 79.5, 79.1, 77.7, 77.2, 77.2, 76.7, 76.5, 76.5, 76.3, 76.1, 76.1, 75.7, 75.5, 74.7, 74.7, 74.3, 74.2, 74.2, 72.8, 72.8, 71.6, 71.5, 69.6, 68.2, 56.4, 52.7, 40.4, 40.3, 40. ESI-MS: $[M + H]^-$ $C_{380}H_{427}N_4O_{99}P_3$ calcd. 2240.9328 found 2240.8775.

**Compound 1**

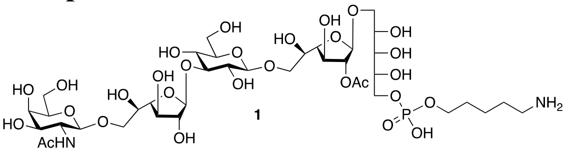


Pentasaccharide **22** (120 mg, 0.048 mmol) was dissolved in $CH_2Cl_2$:$CH_3OH$ (1:1, 10 mL), after which solid $N_2H_4$*HOAc (120 mg) and the reaction mixture was left stirring at RT overnight. The mixture was concentrated *in vacuo*, and passed through Biogel SX1. This material was dissolved in $CH_3OH$:$H_2O$ (4:1, 10 mL), after which AcOH (100 μL) was added, followed by the addition of the Degussa type $Pd(OH)_2$ (20% w/w) (50 mg). The reaction mixture was then hydrogenated overnight, filtered through Celite, and concentrated to afford the target pentasaccharide as a white solid. (40 mg, 78% over two steps). ESI-MS: $[M + H]^-$ $C_{38}H_{69}N_2O_{29}P$ calcd. 1048.3724 found 1048.3716.

$^1$H (750 MHz, $D_2O$): δ (ppm)

| | H-1 | H-2 | H-3 | H-4 | H-5 | H-6 |
|---|---|---|---|---|---|---|
| Rib | 3.84/3.74 | 4.03 | 3.78 | 3.94 | 4.14 | n/a |
| Gal1 | 5.19 | 4.97, d, *J* = 2.05Hz | 4.26<br>dd, *J* = 5.6, 1.9 Hz | 4.10 | 3.80 | n/a |
| Glc | 4.56<br>d, *J* = 8.4 Hz | 3.47 | 3.66 | 3.49 | 3.50 | n/a |
| Gal2 | 5.29<br>d, *J* = 1.9 Hz | 4.18 | 4.08 | 3.95 | 3.72 | n/a |
| GalN | 4.50<br>d, *J* = 8.8 Hz | 3.92 | 3.75 | n/a | n/a | n/a |

$^{13}$C (187 MHz, $D_2O$): δ (ppm)

| | C-1 | C-2 | C-3 | C-4 | C-5 | C-6 |
|---|---|---|---|---|---|---|
| Rib | 68.3 | 70.2 | 71.5 | 69.5 | 67.2 | n/a |
| Gal1 | 105.3 | 83.5 | 75.5 | 83.4 | 71.3 | n/a |
| Glc | 105.8 | 73.6 | 82.6 | 68.5 | 76.0 | n/a |
| Gal2 | 108.6 | 81.4 | 76.4 | 84.0 | 71.3 | n/a |
| GalN | 105.5 | 55.8 | n/a | n/a | n/a | n/a |

**Compound 2**

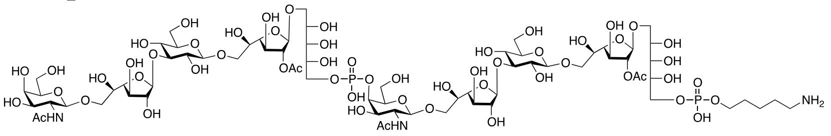

Decasaccharide **24** (100 mg, 0.021 mmol) was dissolved in $CH_2Cl_2$:$CH_3OH$ (1:1, 5 mL), after which solid $N_2H_4$*HOAc (100 mg) and the reaction mixture was left stirring at RT overnight. The mixture was concentrated *in vacuo*, and passed through Biogel SX1. This material was dissolved in $CH_3OH$:$H_2O$ (4:1, 10 mL), after which AcOH (100 µL) was added, followed by the addition of the Degussa type $Pd(OH)_2$ (20% w/w) (50 mg). The reaction mixture was then hydrogenated overnight, filtered through Celite, and concentrated to afford the target Pentasaccharide as a white solid. (34 mg, 82% over two steps). ESI-MS: $[M + H]^-$ $C_{71}H_{125}N_3O_{57}P_2$ calcd. 996.8225 found 996.8202.

$^1$H (750 MHz, $D_2O$): δ (ppm)

| | H-1 | H-2 | H-3 | H-4 | H-5 | H-6 |
|---|---|---|---|---|---|---|
| Rib | 3.84/3.74 | 4.03 | 3.78 | 3.94 | 4.14 | n/a |
| Gal1 | 5.19 | 4.97 | 4.25 | 4.10 | 3.80 | n/a |
| Glc | 4.56<br>d, *J* = 7.9 Hz | 3.47 | 3.66 | 3.49 | 3.50 | n/a |
| Gal2 | 5.29 | 4.18 | 4.08 | 3.95 | 3.72 | n/a |
| GalN-1 | 4.56 | 3.97 | 3.85 | 4.52<br>dd, *J* = 9.9, 3.8 Hz | n/a | n/a |
| GalN-2 | 4.50<br>d, *J* = 8.4 Hz | 3.92 | 3.75 | n/a | n/a | n/a |

$^{13}$C (187 MHz, $D_2O$): δ (ppm)

| | C-1 | C-2 | C-3 | C-4 | C-5 | C-6 |
|---|---|---|---|---|---|---|
| Rib | 68.3 | 70.2 | 71.5 | 69.5 | 67.2 | n/a |
| Gal1 | 105.3 | 83.5 | 75.5 | 83.4 | 71.3 | n/a |
| Glc | 105.8 | 73.6 | 82.6 | 68.5 | 76.0 | n/a |
| Gal2 | 108.6 | 81.4 | 76.4 | 84.0 | 71.3 | n/a |
| GalN | 105.5 | 55.8 | n/a | n/a | n/a | n/a |
| GalN-2 | 105.7 | 55.8 | n/a | 76.2 | n/a | n/a |

**Compound 3**

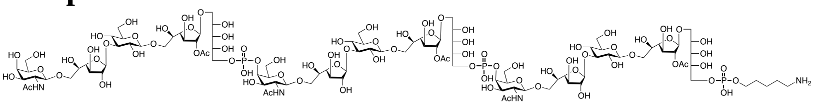

Pentadecasaccharide **26** (131 mg, 0.019 mmol) was dissolved in $CH_2Cl_2$:$CH_3OH$ (1:1, 5 mL), after which solid $N_2H_4$*HOAc (100 mg) and the reaction mixture was left stirring at RT overnight. The mixture was concentrated *in vacuo*, and passed through Biogel SX1. This material was dissolved in $CH_3OH$:$H_2O$ (4:1, 10 mL), after which AcOH (100 µL) was added, followed by the addition of the Degussa type $Pd(OH)_2$ (20% w/w) (50 mg). The reaction mixture was then hydrogenated overnight, filtered through Celite, and concentrated to afford the target pentasaccharide as a white solid. (34 mg, 82% over two steps). ESI-MS: $[M + H]^-$ $C_{104}H_{181}N_4O_{85}P_3$ calcd. 979.6392 found 979.6513.

$^1H$ (750 MHz, $D_2O$): δ (ppm)

| | H-1 | H-2 | H-3 | H-4 | H-5 | H-6 |
|---|---|---|---|---|---|---|
| Rib | 3.84/3.74 | 4.03 | 3.78 | 3.94 | 4.14 | n/a |
| Gal1 | 5.19 | 4.97 | 4.25 | 4.10 | 3.80 | n/a |
| Glc | 4.56<br>d, *J* = 7.9 Hz | 3.47 | 3.66 | 3.49 | 3.50 | n/a |
| Gal2 | 5.29 | 4.18 | 4.08 | 3.95 | 3.72 | n/a |
| GalN-1 | 4.56 | 3.97 | 3.85 | 4.52<br>dd, *J* = 9.9, 3.8 Hz | n/a | n/a |
| GalN-2 | 4.56 | 3.97 | 3.85 | 4.52<br>dd, *J* = 9.9, 3.8 Hz | n/a | n/a |
| GalN-3 | 4.50<br>*J* = 8.4 Hz | 3.92 | 3.75 | n/a | n/a | n/a |

$^{13}C$ (187 MHz, $D_2O$): δ (ppm)

| | C-1 | C-2 | C-3 | C-4 | C-5 | C-6 |
|---|---|---|---|---|---|---|
| Rib | 68.3 | 70.2 | 71.5 | 69.5 | 67.2 | n/a |
| Gal1 | 105.3 | 83.5 | 75.5 | 83.4 | 71.3 | n/a |
| Glc | 105.8 | 73.6 | 82.6 | 68.5 | 76.0 | n/a |
| Gal2 | 108.6 | 81.4 | 76.4 | 84.0 | 71.3 | n/a |
| GalN-1 | 105.7 | 55.8 | n/a | 76.2 | n/a | n/a |
| GalN-2 | 105.7 | 55.8 | n/a | 76.2 | n/a | n/a |
| GalN-3 | 105.5 | 55.8 | n/a | n/a | n/a | n/a |

**Microarray Printing Procedure.** All oligosaccharides were printed on NHS-ester activated glass slides (NEXTERION® Slide H, Schott Inc.) using a Scienion sciFLEXARRAYER S3 non-contact microarray equipped with a Scienion PDC80 nozzle (Scienion Inc.). Individual samples were dissolved in sodium phosphate buffer (50 µL, 0.225 M, pH 8.5) at a concentration of 100 µM and were printed in replicates of 10 with spot volume ~ 400 pL, at 20°C and 50% humidity. Each slide has 24 subarrays in a 3x8 layout. After printing, slides were incubated in a humidity chamber for 8 h and then blocked for 30 min with a 5mM ethanolamine in a Tris buffer (pH 8,5, 50 mM) at 40°C. Blocked slides were rinsed with DI water, spun dry, and kept in a desiccator at room temperature for future use. The printed glass slide was pre-blocked with a solution of 1x TSM binding buffer (20 mM Tris·HCl, pH 7.4, 150 mM NaCl, 2 mM $CaCl_2$, and 2 mM $MgCl_2$, 0.05% Tween-20, 1% BSA) for 90 min and the blocking solution was discarded. The protein or sera samples were diluted in different ratios in TMS binding buffer, and the solution was incubated on the microarray for 60 min at room temperature. The slide was sequentially washed with TSM wash buffer (20 mM Tris·HCl, pH 7.4, 150 mM NaCl, 2 mM $CaCl_2$, and 2 mM $MgCl_2$, 0.05% Tween-20), TSM buffer (20 mM Tris·HCl, pH 7.4, 150 mM NaCl, 2 mM $CaCl_2$, and 2 mM $MgCl_2$) and water.

**Microarray Binding Studies with Factor Sera 29b and 35a**. Factor sera 29b and 35a were obtained from SSI Diagnostica. The slides were incubated with the anti-sera at different dilutions for 1 h, followed by washing and re-incubation with a premixed solution of biotinylated Goat-anti-Rabbit antibody (Southern Biotech, 4041-08, 10 µg/mL in TSM binding buffer) and Streptavidin-Alexa Fluor 647 conjugate (Invitrogen, S21347, 10 µg/mL in TSM binding buffer) for 30 min. The slide was scanned after washing and drying. The slides were scanned using a GenePix 4000B microarray scanner (Molecular Devices) at the appropriate excitation wavelength with a resolution of 5µM. Various gains and PMT values were employed in the scanning to ensure that all the signals were within the liner range of the scanner's detector and there was no saturation of signals. The image was analyzed using GenePix Pro 7 software (version 7.2.29.2, Molecular Devices). The data was analyzed with an Excel macro (https://doi.org/ 10.5281/zenodo.5146251) to provide the results. The highest and the lowest value of the total fluorescence intensity of the six replicates spots were removed, and the four values in the middle were used to provide the mean value and standard deviation.

**Microarray Binding Studies with Ficolin-2**. The slides were incubated with ficolin-2 (R&D Systems, 2428-FC-050, 2, 20 and 100 µg/mL in TMS binding buffer) for 1 h, followed by washing and re-incubation with a solution of AlexaFluor® 647 conjugated anti-His antibody (BioLegend #652513,10 µg/mL) for 30 min. The slide was scanned after washing and drying.

### References


(1) Tadano, K.; Maeda, H.; Hoshino, M.; Iimura, Y.; Suami, T. A Novel Transformation of Four Aldoses to Some Optically Pure Pseudohexopyranoses and a Pseudopentofuranose, Carbocyclic Analogs of Hexopyranoses and Pentofuranose. Synthesis of Derivatives of (1S,2S,3R,4S,5S)-, (1S,2S,3R,4R,5S)-, (1R,2R,3R,4R,5S)-, (1S,2S,3R,4S,5R)-2,3,4,5-tetrahydroxy-1-(hydroxymethyl)cyclohexanes and (1S,2S,3S,4S)-2,3,4-trihydroxy-1(hydroxymethyl)cyclopentane. *J. Org. Chem.* **1987**, *52* (10), 1946-1956.

(2) Wu, Y.; Xiong, D.-C.; Chen, S.-C.; Wang, Y.-S.; Ye, X.-S. Total Synthesis of Mycobacterial Arabinogalactan Containing 92 Monosaccharide Units. *Nat. Commun.* **2017**, *8* (1), 14851.

(3) Ellervik, U.; Grundberg, H.; Magnusson, G. Synthesis of Lactam and Acetamido Analogues of Sialyl Lewis X Tetrasaccharide and Lewis X Trisaccharide. *J. Org. Chem.* **1998**, *63* (25), 9323-9338.

(4) Lu, S.-R.; Lai, Y.-H.; Chen, J.-H.; Liu, C.-Y.; Mong, K.-K. T. Dimethylformamide: An Unusual Glycosylation Modulator. *Angew. Chem. Int. Ed.* **2011**, *50* (32), 7315-7320.

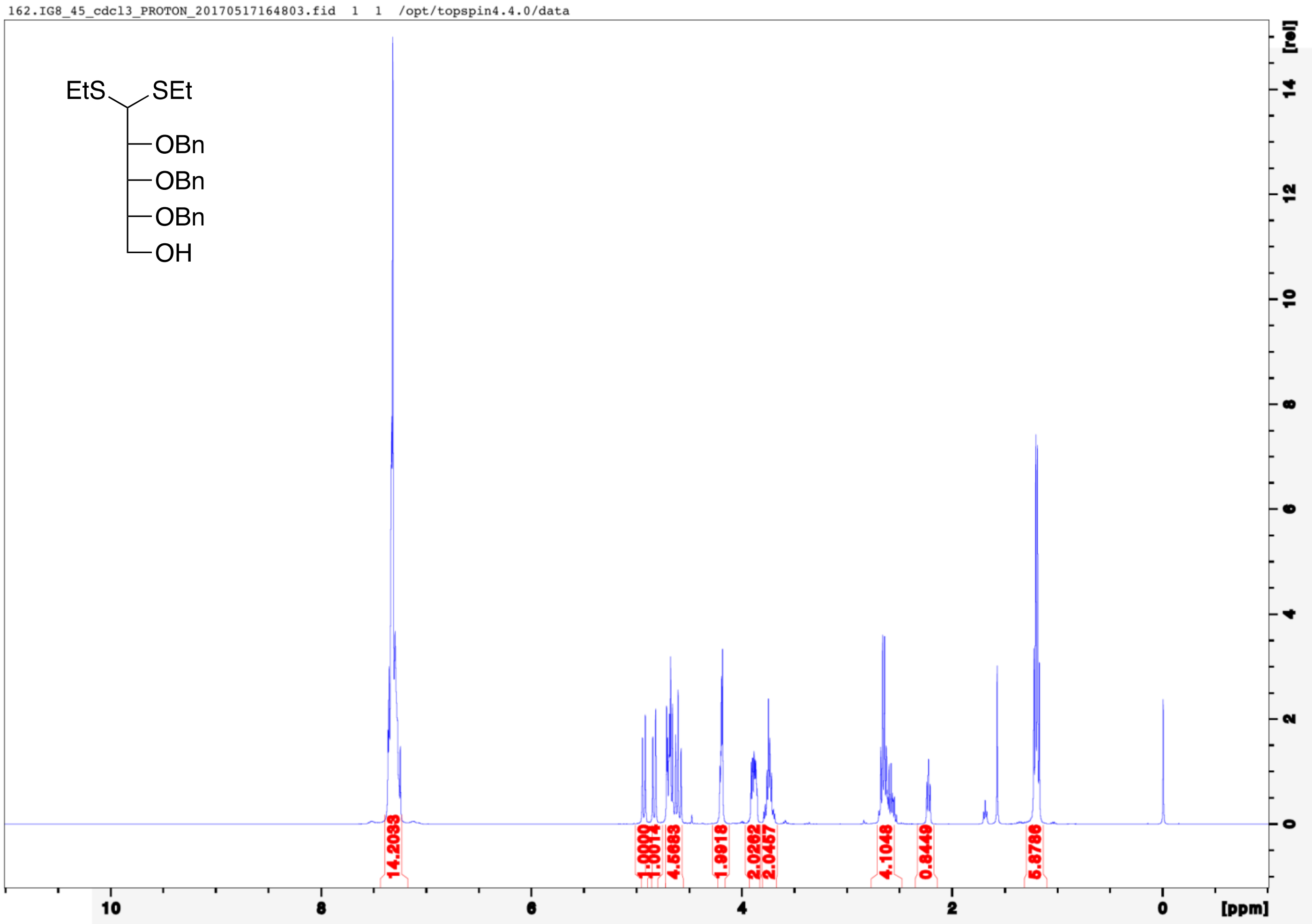

162.IG8_45_cdcl3_PROTON_20170517164803.fid 1 1 /opt/topspin4.4.0/data
EtS
SEt
OBn
OBn
OBn
OH
14.2033
1.0000
1.0014
4.5683
1.9918
2.0262
2.0457
4.1048
0.8449
5.8786
10
8
6
4
2
0
[ppm]
[rel]

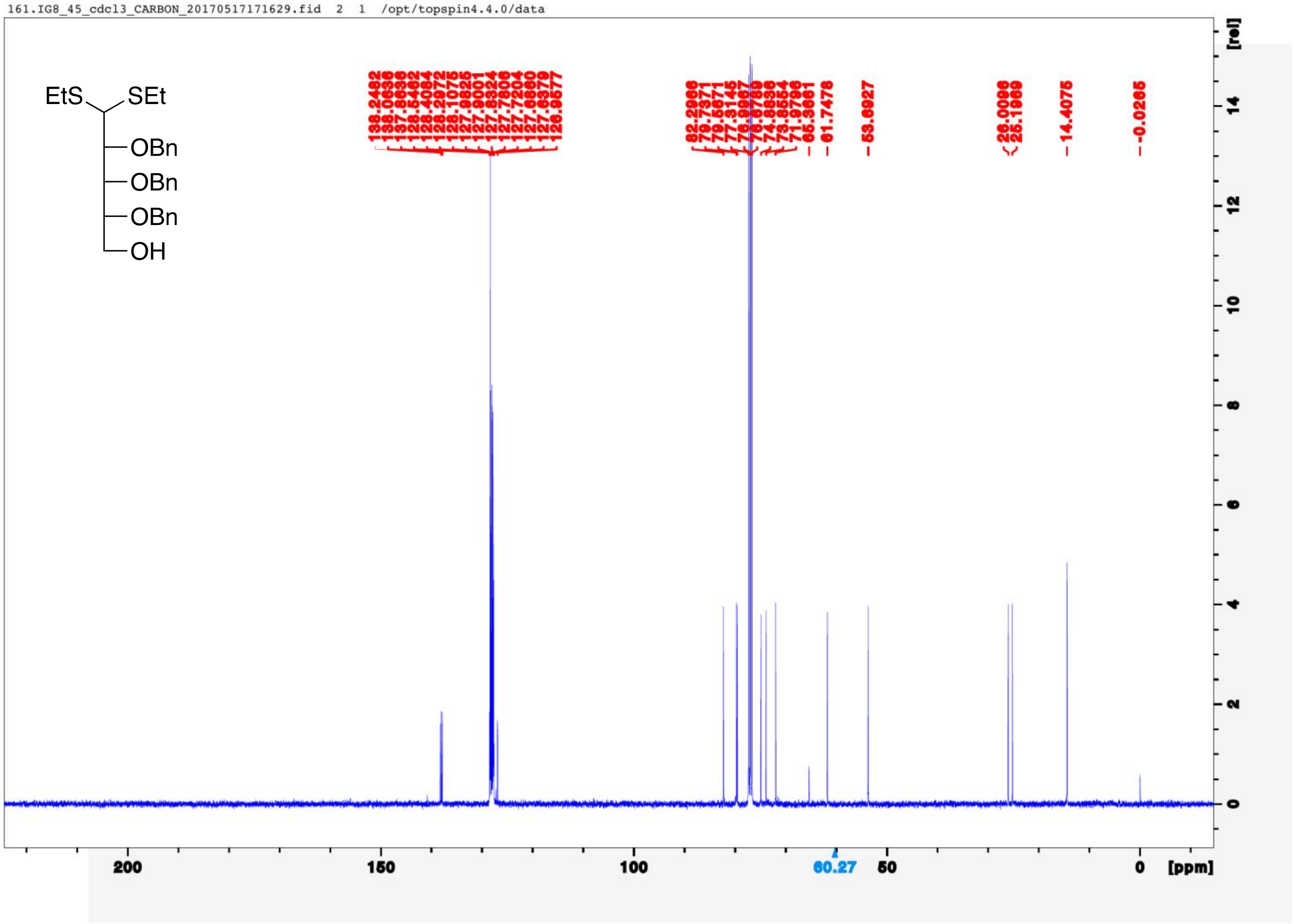
161.IG8_45_cdcl3_CARBON_20170517171629.fid 2 1 /opt/topspin4.4.0/data
EtS
SEt
OBn
OBn
OBn
OH
138.2482
138.0636
137.8636
128.5462
128.4084
128.2972
128.1075
127.9825
127.9001
127.8324
127.7806
127.7204
127.6860
127.6379
126.9577
82.2966
79.7371
77.5671
77.3145
76.9967
76.6789
74.8836
73.8554
71.9796
65.3661
61.7478
53.6927
26.0096
25.1969
14.4075
-0.0265
[rel]
14
12
10
8
6
4
2
0
200
150
100
60.27
50
0
[ppm]

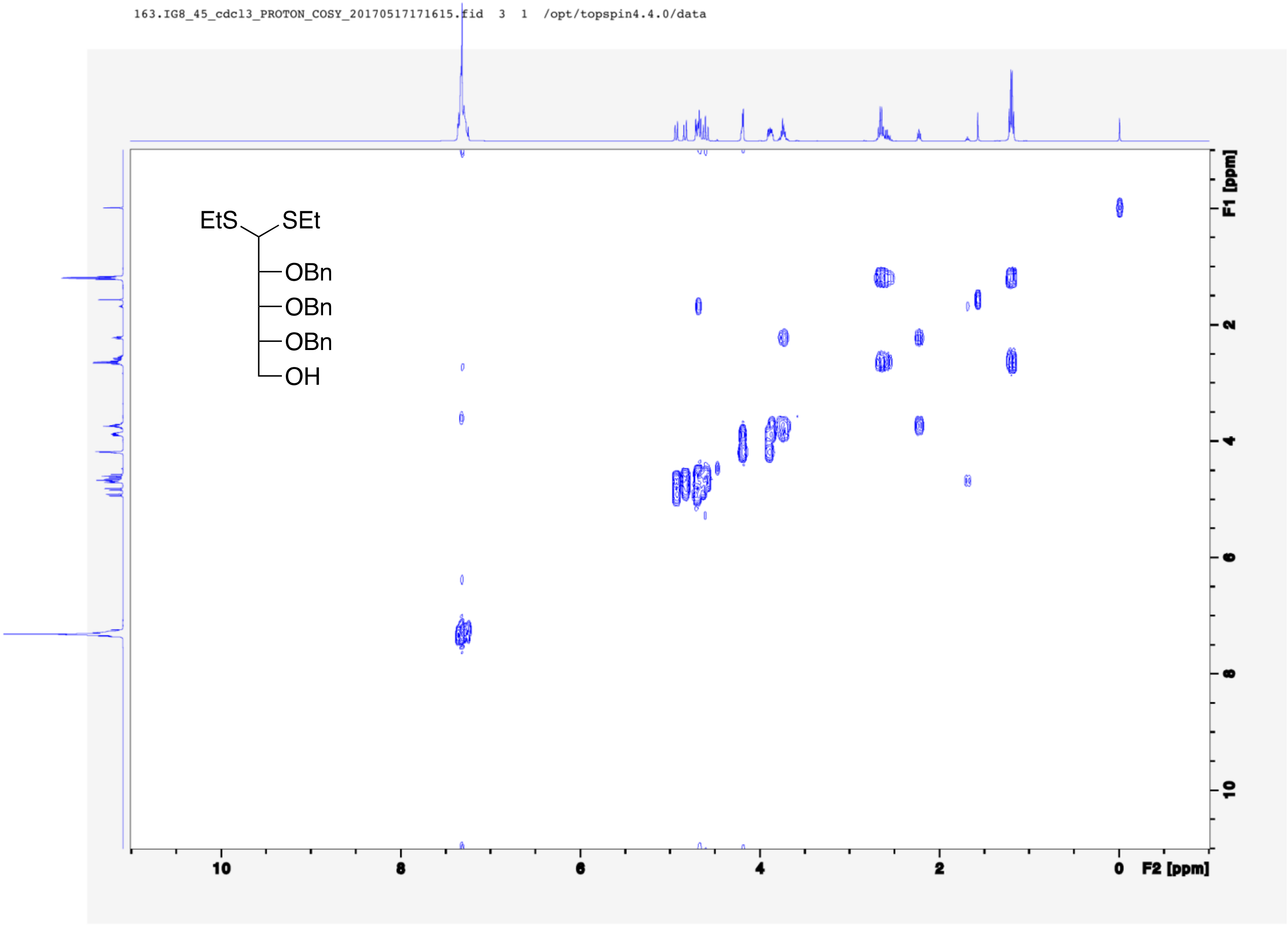
163.IG8_45_cdcl3_PROTON_COSY_20170517171615.fid 3 1 /opt/topspin4.4.0/data
EtS
SEt
OBn
OBn
OBn
OH
F1 [ppm]
F2 [ppm]
10
8
6
4
2
0

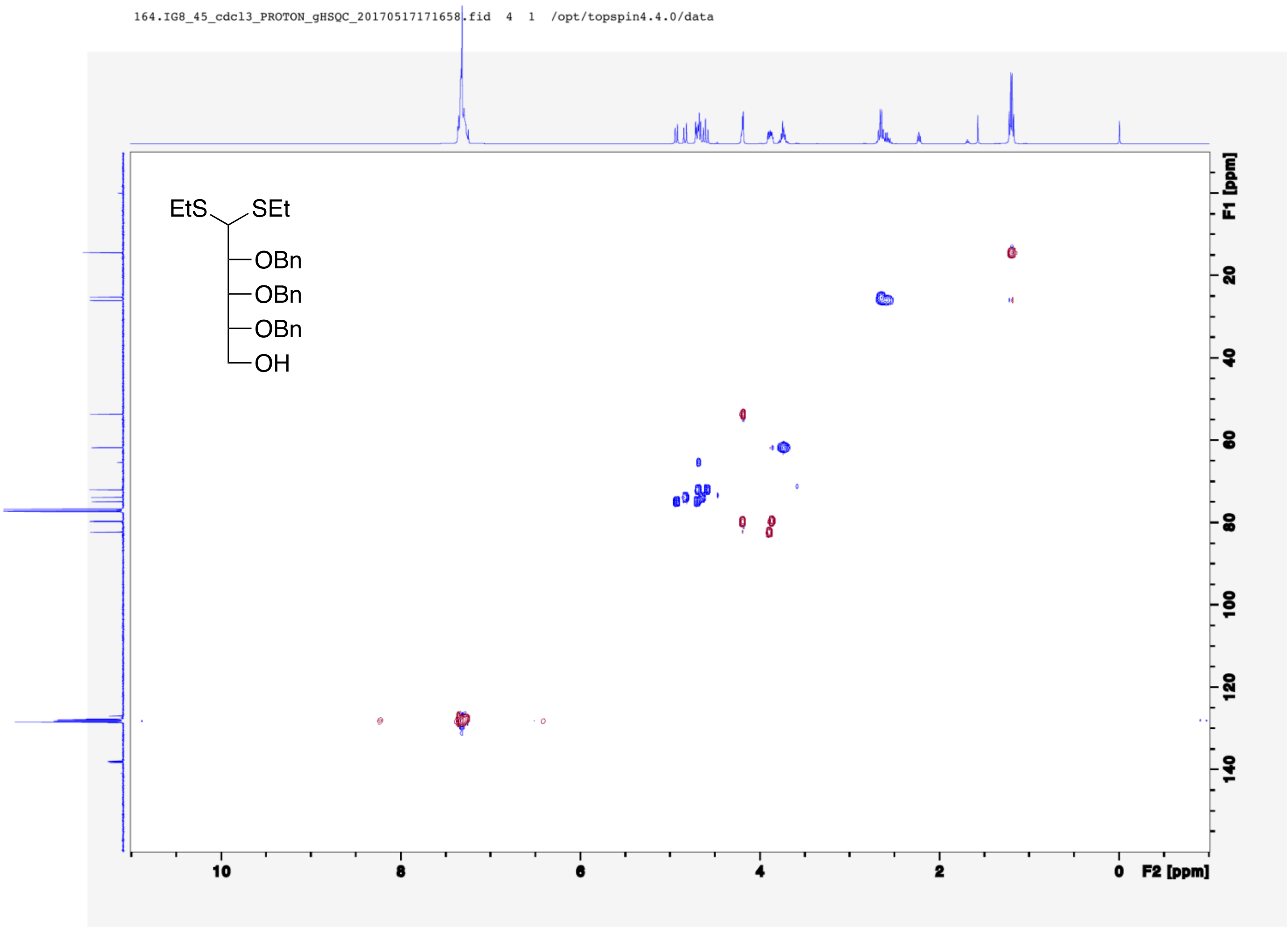
164.IG8_45_cdcl3_PROTON_gHSQC_20170517171658.fid 4 1 /opt/topspin4.4.0/data
EtS SEt
OBn
OBn
OBn
OH
F1 [ppm]
20
40
60
80
100
120
140
10
8
6
4
2
0
F2 [ppm]

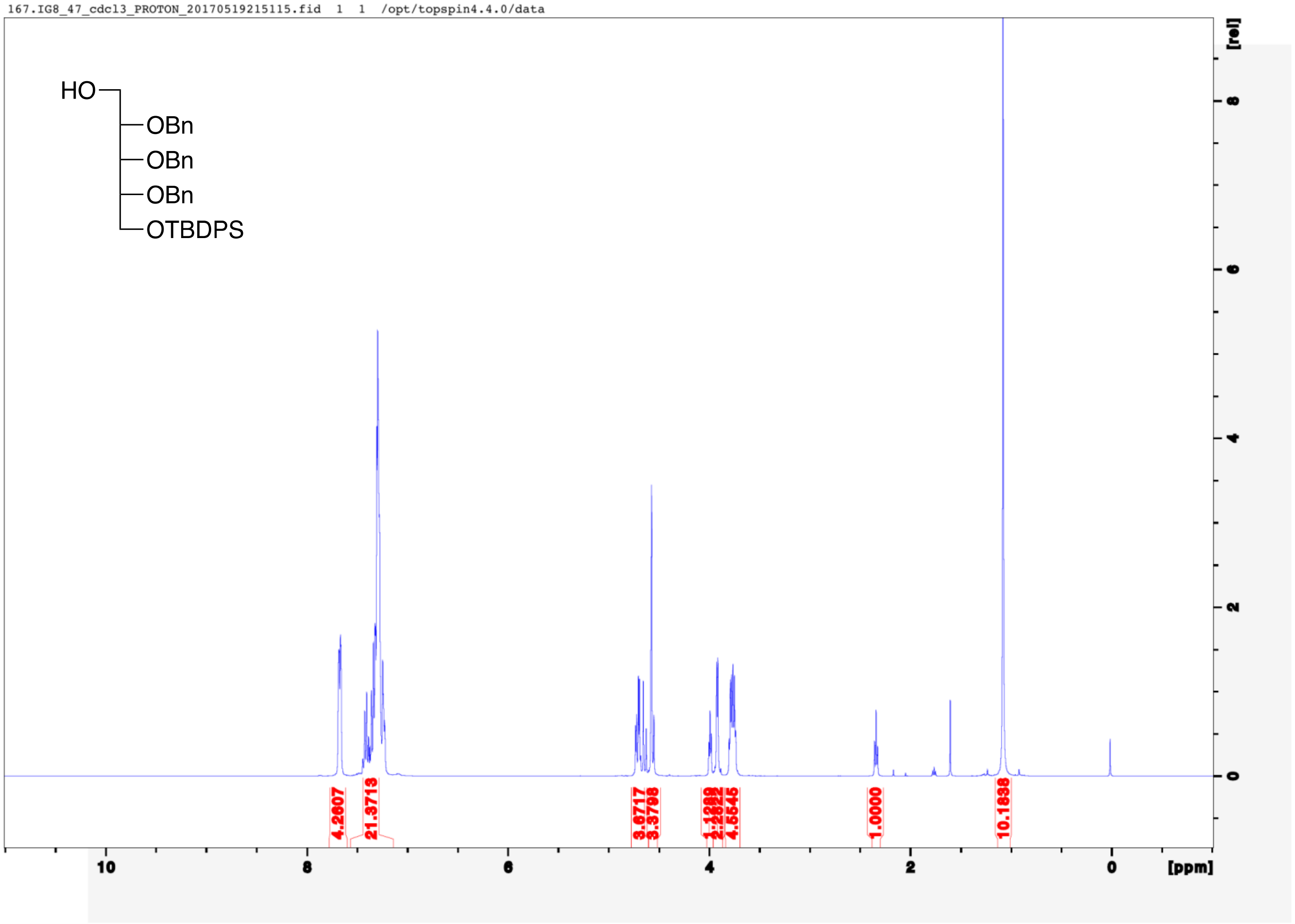
167.IG8_47_cdcl3_PROTON_20170519215115.fid 1 1 /opt/topspin4.4.0/data
HO
OBn
OBn
OBn
OTBDPS
[rel]
4.2607
21.3713
3.6717
3.3798
4.5545
1.0000
10.1838
10
8
6
4
2
0
[ppm]

165.IG8_47_cdcl3_CARBON_20170519215300.fid 1 1 /opt/topspin4.4.0/data

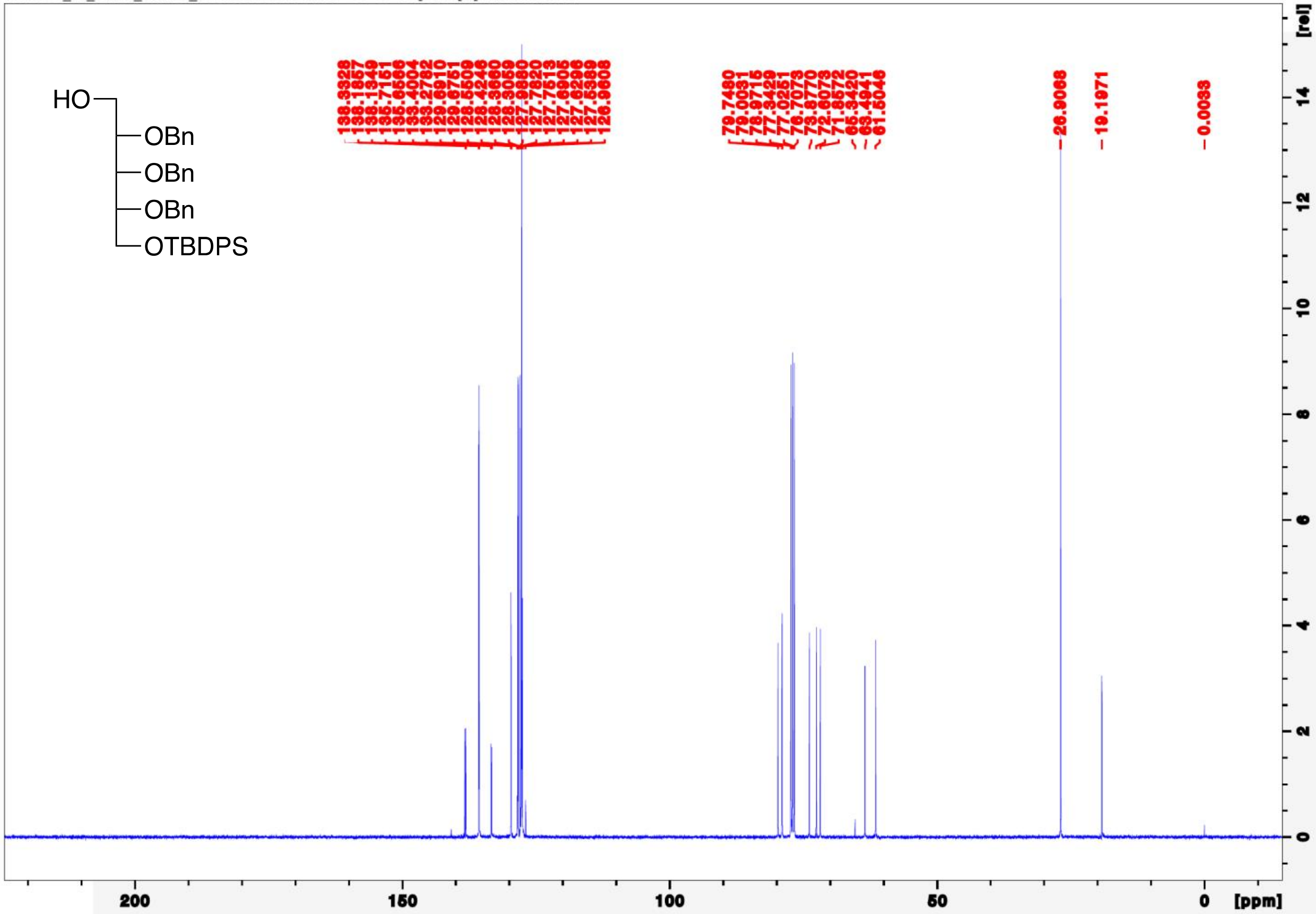

168.IG8_47_cdcl3_PROTON_gCOSY_20170518175052.fid 1 1 /opt/topspin4.4.0/data

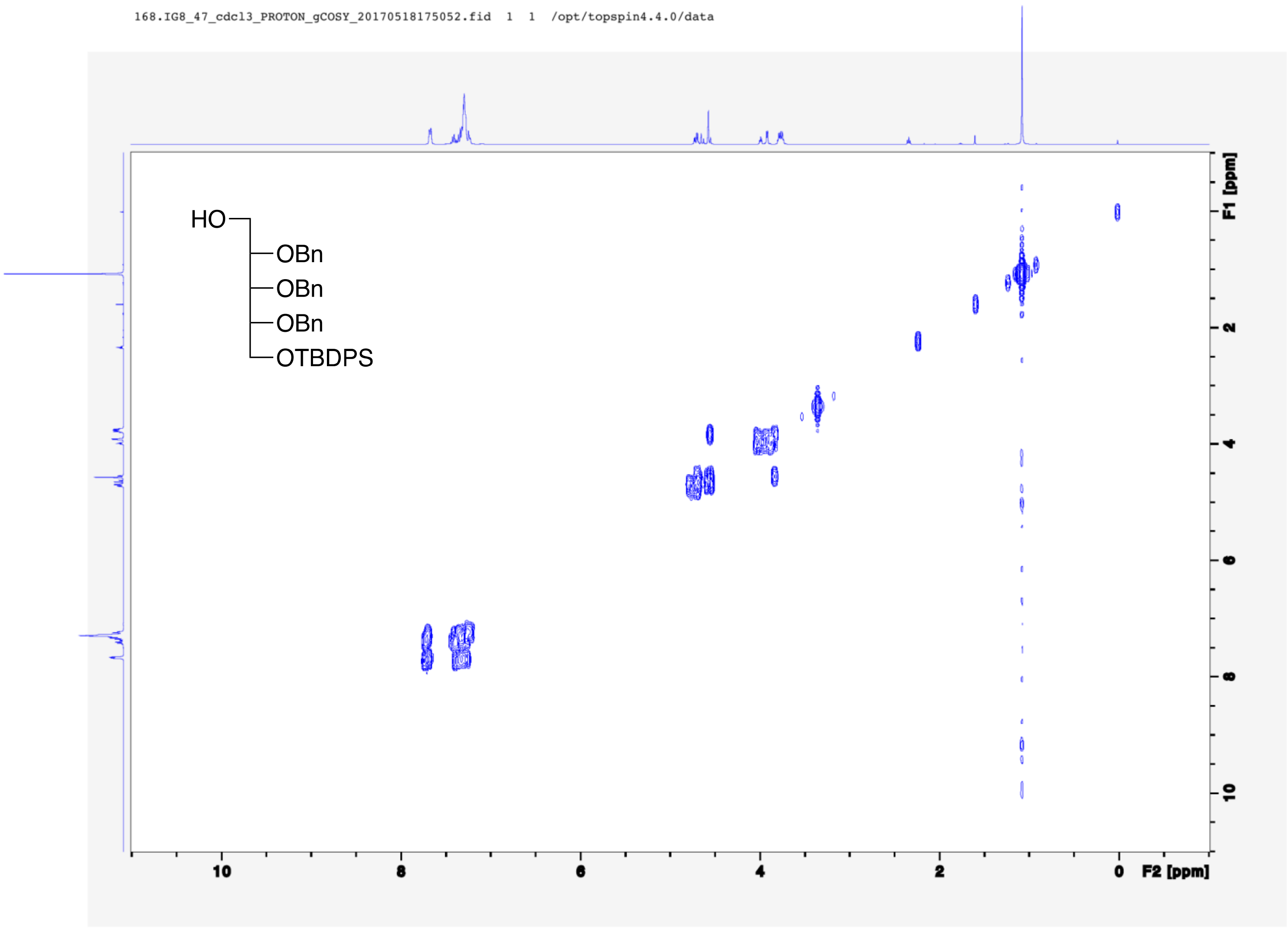

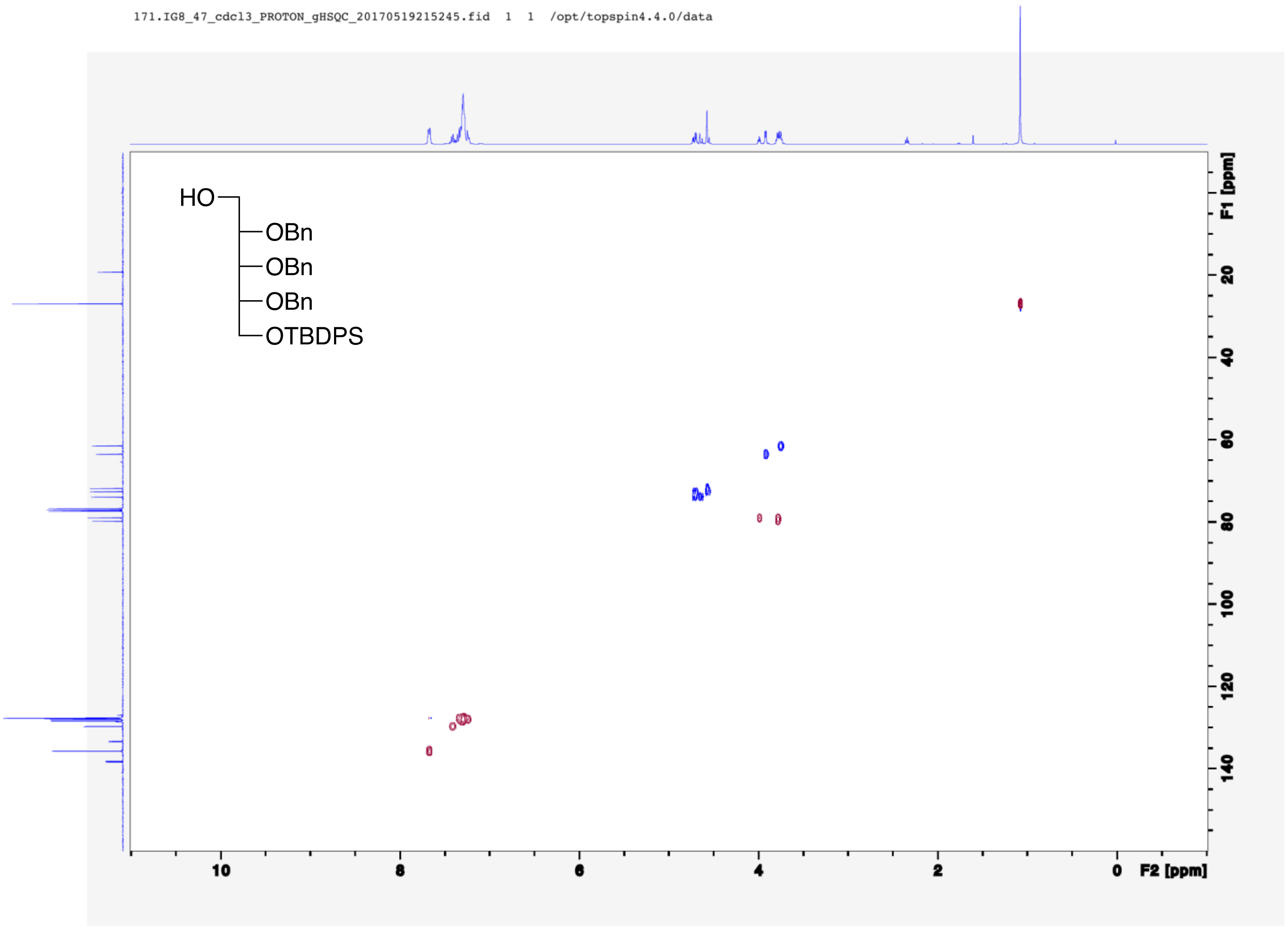
171.IG8_47_cdcl3_PROTON_gHSQC_20170519215245.fid 1 1 /opt/topspin4.4.0/data
HO
OBn
OBn
OBn
OTBDPS
F1 [ppm]
20
40
60
80
100
120
140
10
8
6
4
2
0
F2 [ppm]

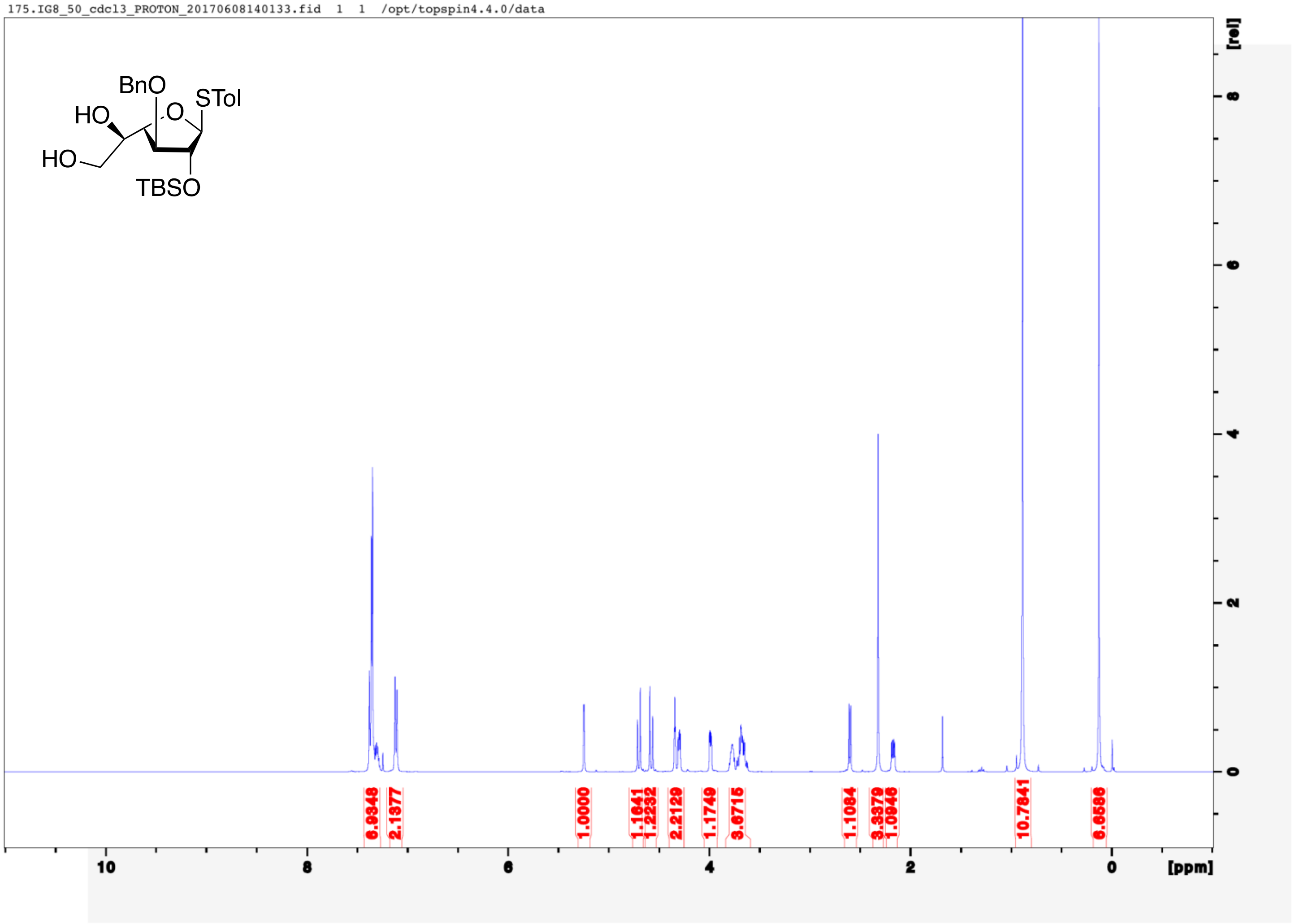
175.IG8_50_cdcl3_PROTON_20170608140133.fid 1 1 /opt/topspin4.4.0/data
BnO
STol
HO
O
HO
TBSO
6.9348
2.1377
1.0000
1.1641
1.2232
2.2129
1.1749
3.6715
1.1084
3.3379
1.0946
10.7841
6.6586
10
8
6
4
2
0
[ppm]
[rel]

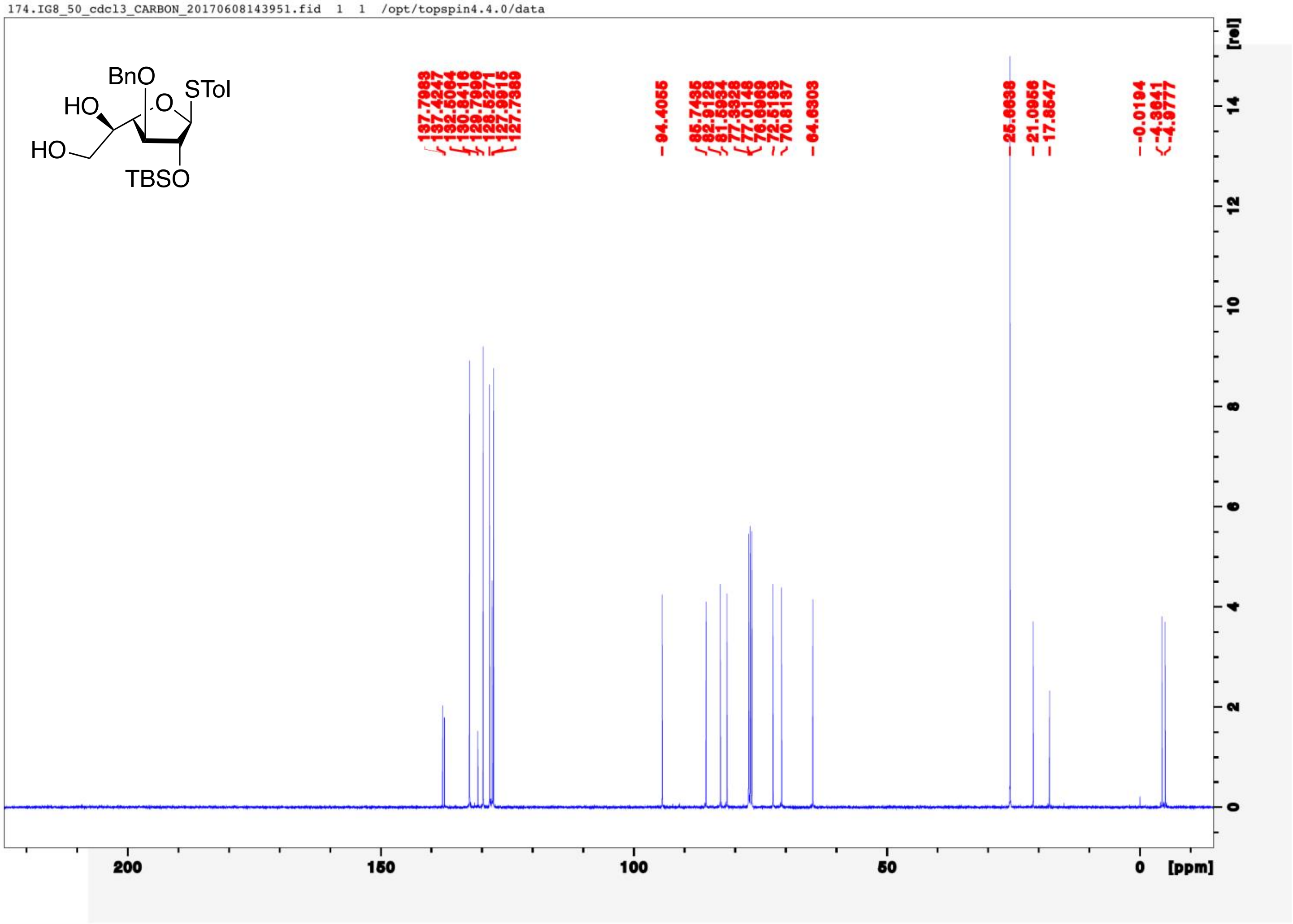
174.IG8_50_cdcl3_CARBON_20170608143951.fid 1 1 /opt/topspin4.4.0/data
BnO
STol
HO
O
HO
TBSO
137.7983
137.4247
132.5064
130.8416
129.7996
128.5271
127.9915
127.7389
94.4055
85.7435
82.9128
81.5934
77.3328
77.0148
76.6969
72.5193
70.8137
64.6303
25.6638
21.0956
17.8547
-0.0194
-4.3641
-4.3777
200
150
100
50
0
[ppm]
[rel]

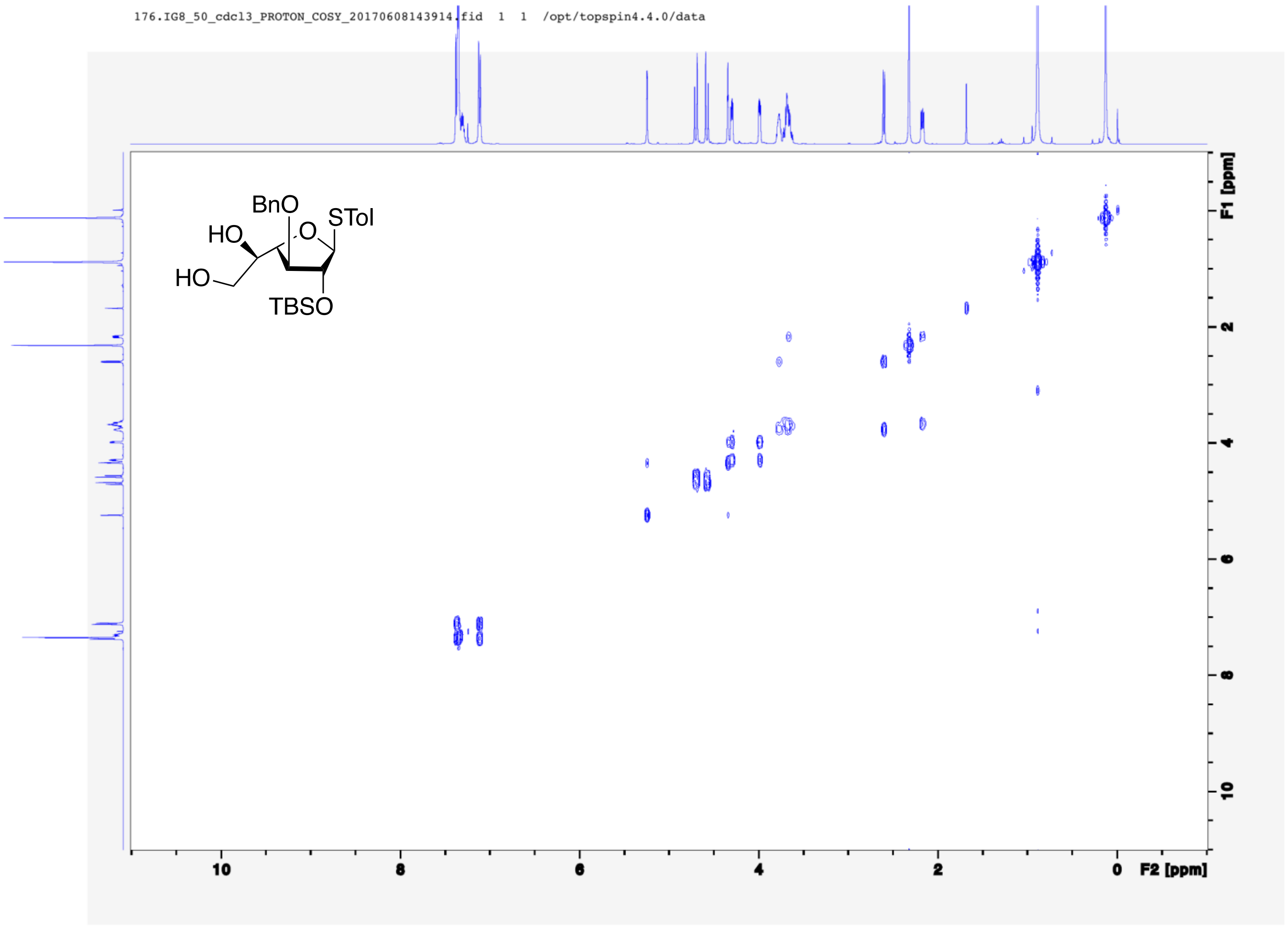
176.IG8_50_cdcl3_PROTON_COSY_20170608143914.fid 1 1 /opt/topspin4.4.0/data
BnO
STol
HO
O
HO
TBSO
F1 [ppm]
F2 [ppm]
10
8
6
4
2
0

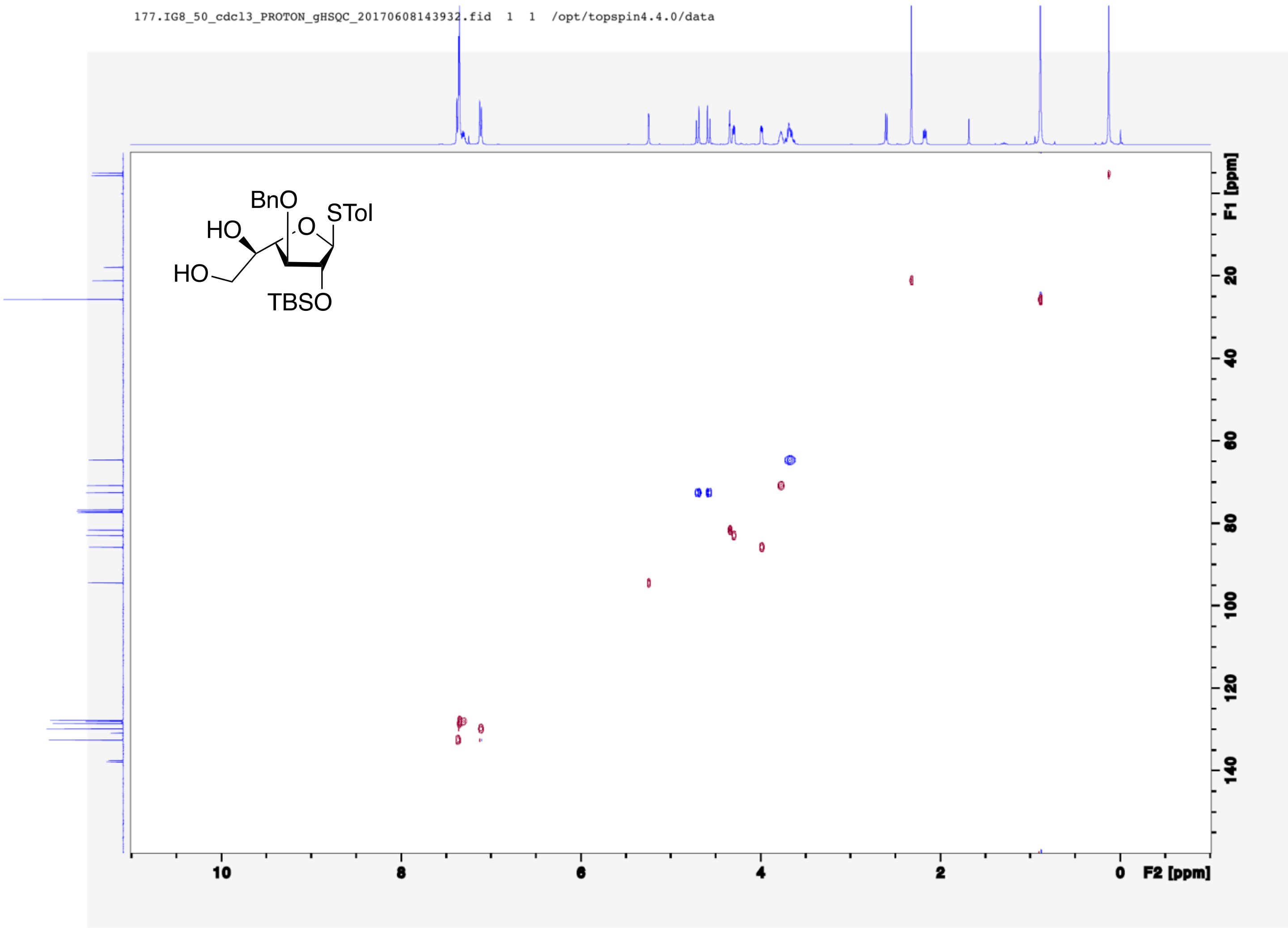
177.IG8_50_cdcl3_PROTON_gHSQC_20170608143932.fid 1 1 /opt/topspin4.4.0/data
BnO
HO
O
STol
HO
TBSO
F1 [ppm]
20
40
60
80
100
120
140
10
8
6
4
2
0
F2 [ppm]

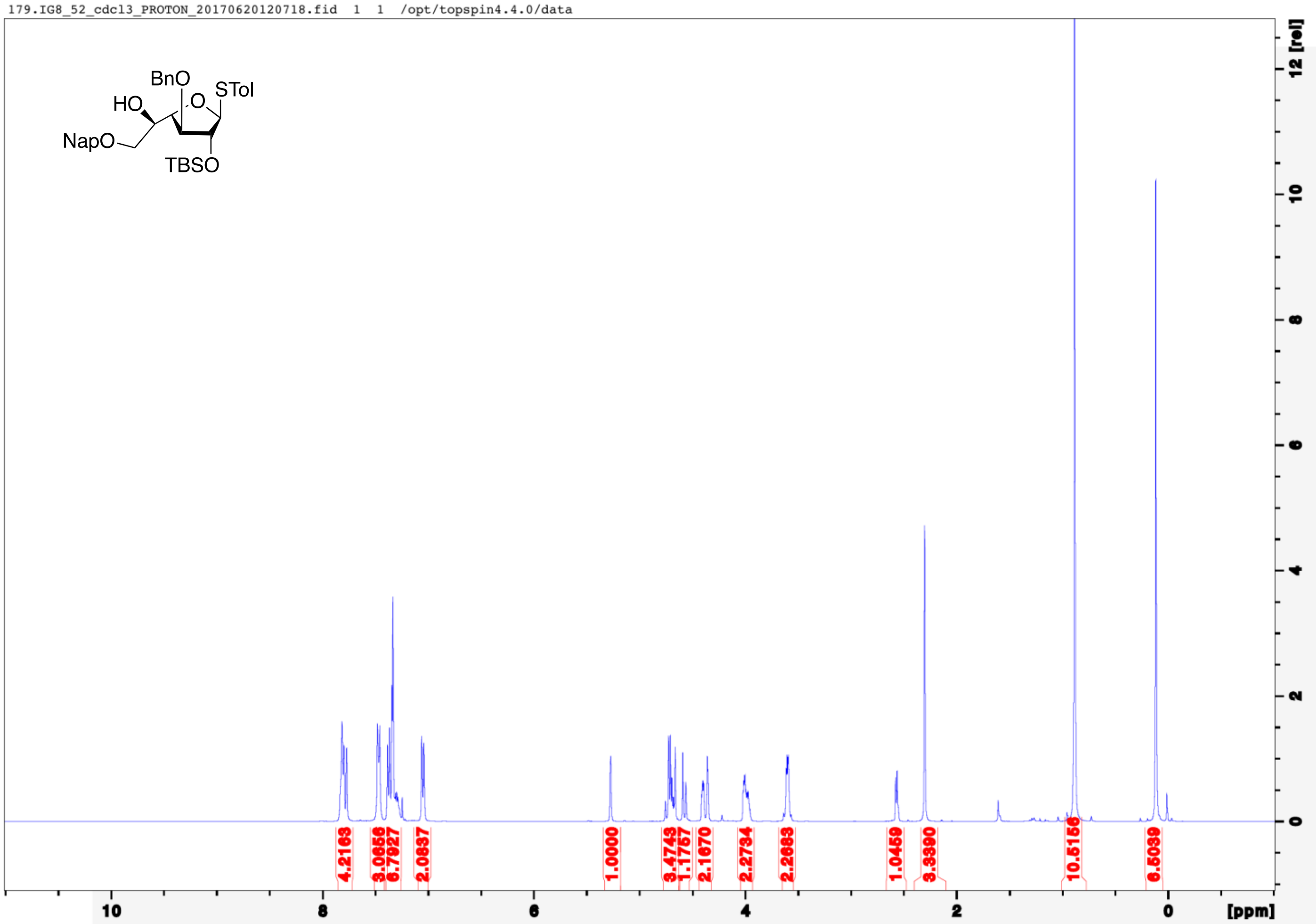

179.IG8_52_cdcl3_PROTON_20170620120718.fid 1 1 /opt/topspin4.4.0/data
BnO
HO
STol
NapO
TBSO
4.2163
3.0656
6.7927
2.0837
1.0000
3.4743
1.1757
2.1670
2.2734
2.2683
1.0459
3.3390
10.5156
6.5039
10
8
6
4
2
0
[ppm]
12 [rel]

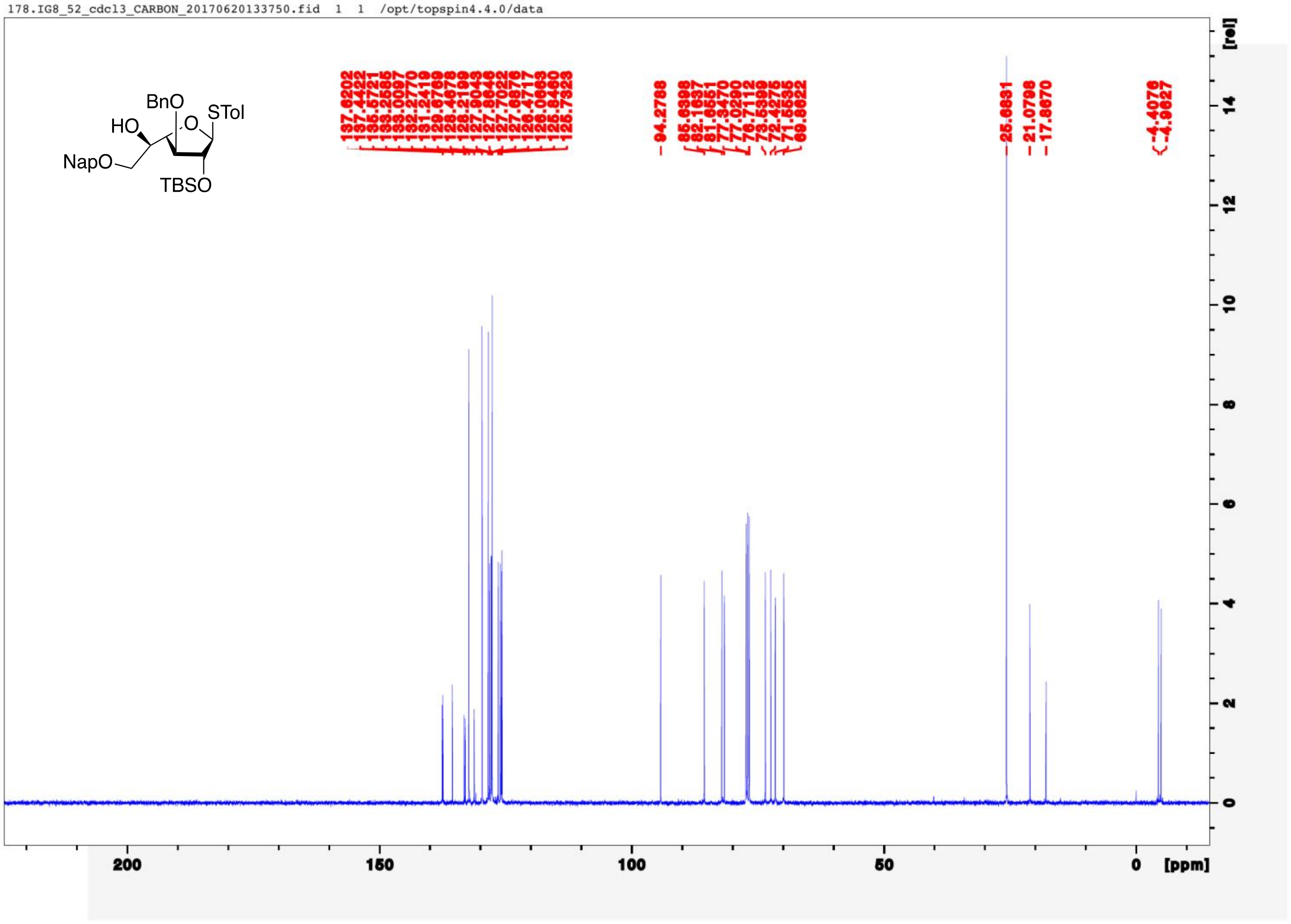
178.IG8_52_cdcl3_CARBON_20170620133750.fid 1 1 /opt/topspin4.4.0/data
BnO
STol
HO
NapO
TBSO
137.6202
137.4422
135.5721
133.2585
133.0097
132.2770
131.2419
129.6789
128.4678
128.2199
127.9043
127.8646
127.7022
127.6876
126.4717
126.0663
125.8460
125.7323
94.2788
85.6398
82.1637
81.6551
77.3470
77.0290
76.7112
73.5399
72.4275
71.5535
69.8622
25.8831
21.0798
17.8670
-4.4076
-4.9627
[rel]
[ppm]

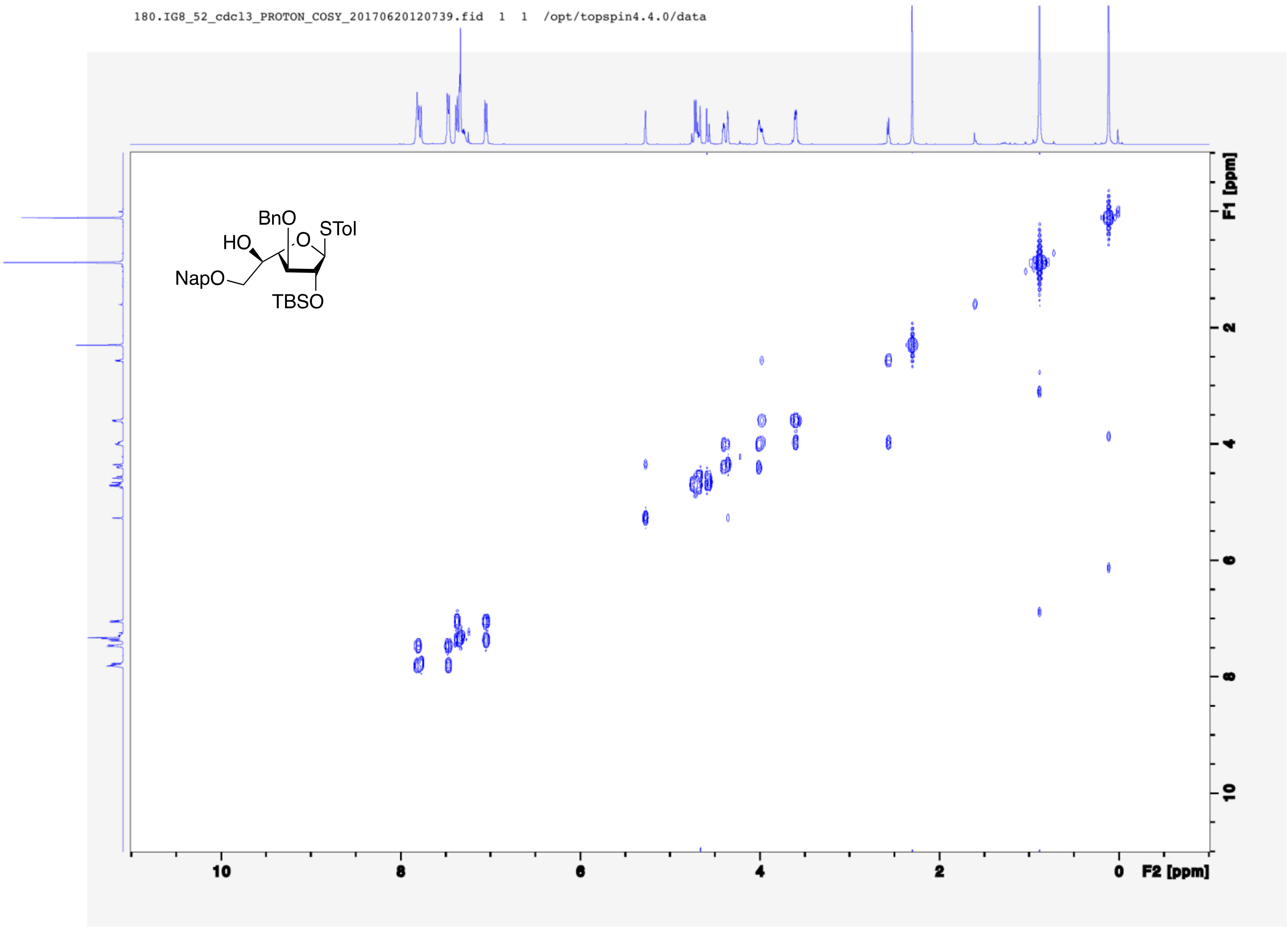

180.IG8_52_cdcl3_PROTON_COSY_20170620120739.fid 1 1 /opt/topspin4.4.0/data
BnO
HO
O
STol
NapO
TBSO
F1 [ppm]
F2 [ppm]
10
8
6
4
2
0

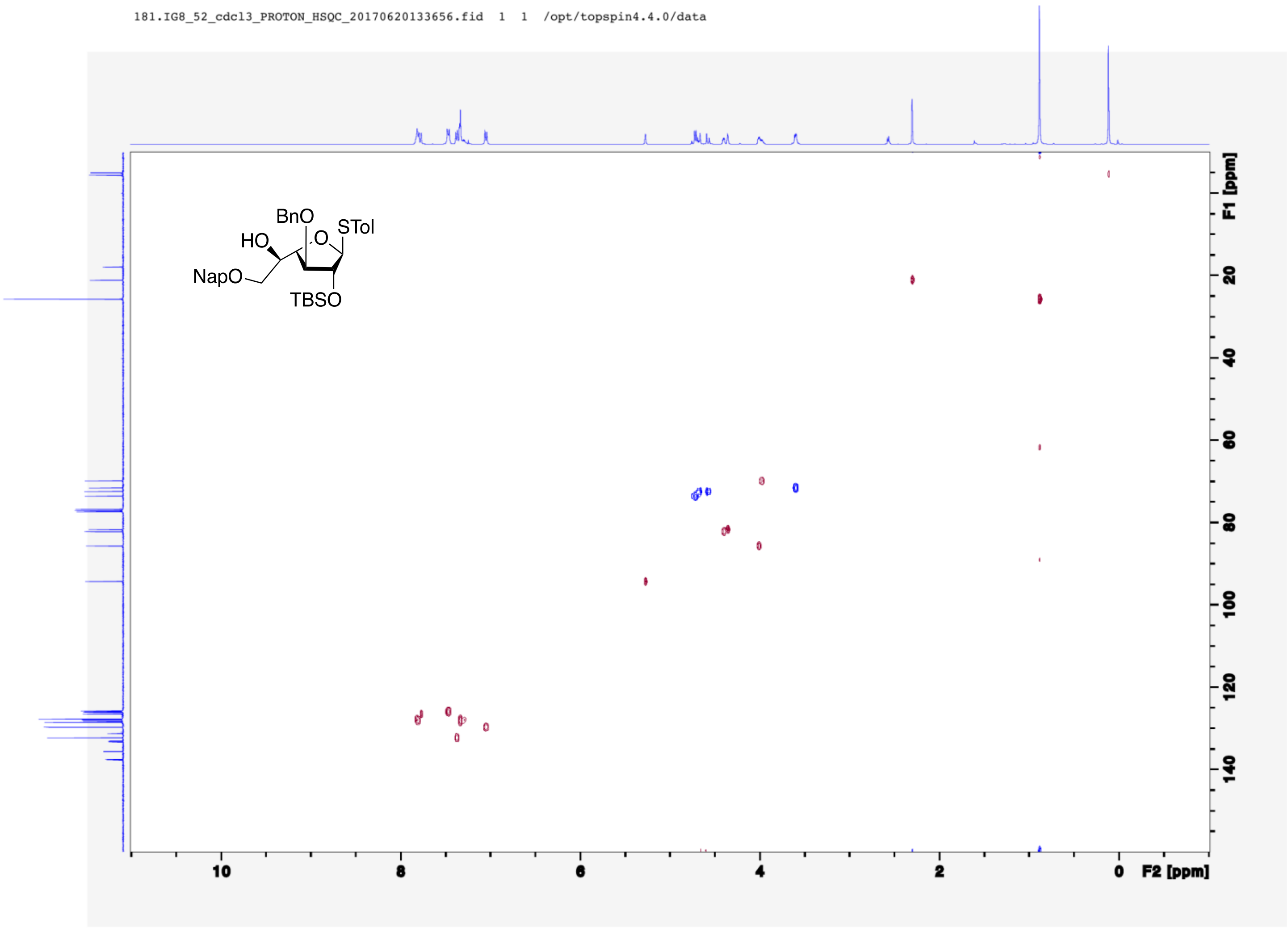
181.IG8_52_cdcl3_PROTON_HSQC_20170620133656.fid 1 1 /opt/topspin4.4.0/data
BnO
STol
HO
O
NapO
TBSO
F1 [ppm]
F2 [ppm]
10
8
6
4
2
0
20
40
60
80
100
120
140

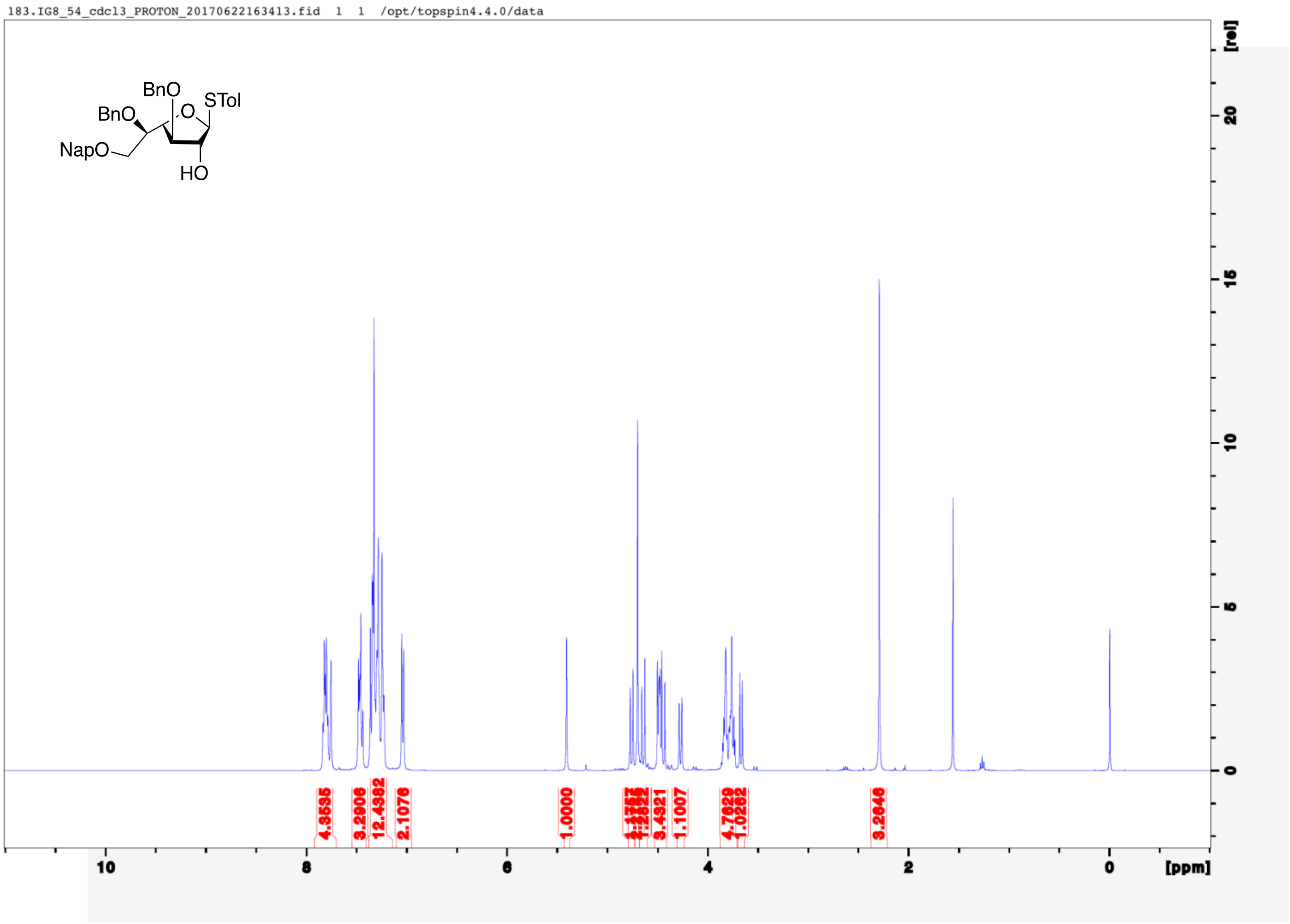
183.IG8_54_cdcl3_PROTON_20170622163413.fid 1 1 /opt/topspin4.4.0/data
BnO
BnO
O
STol
NapO
HO
[rel]
[ppm]

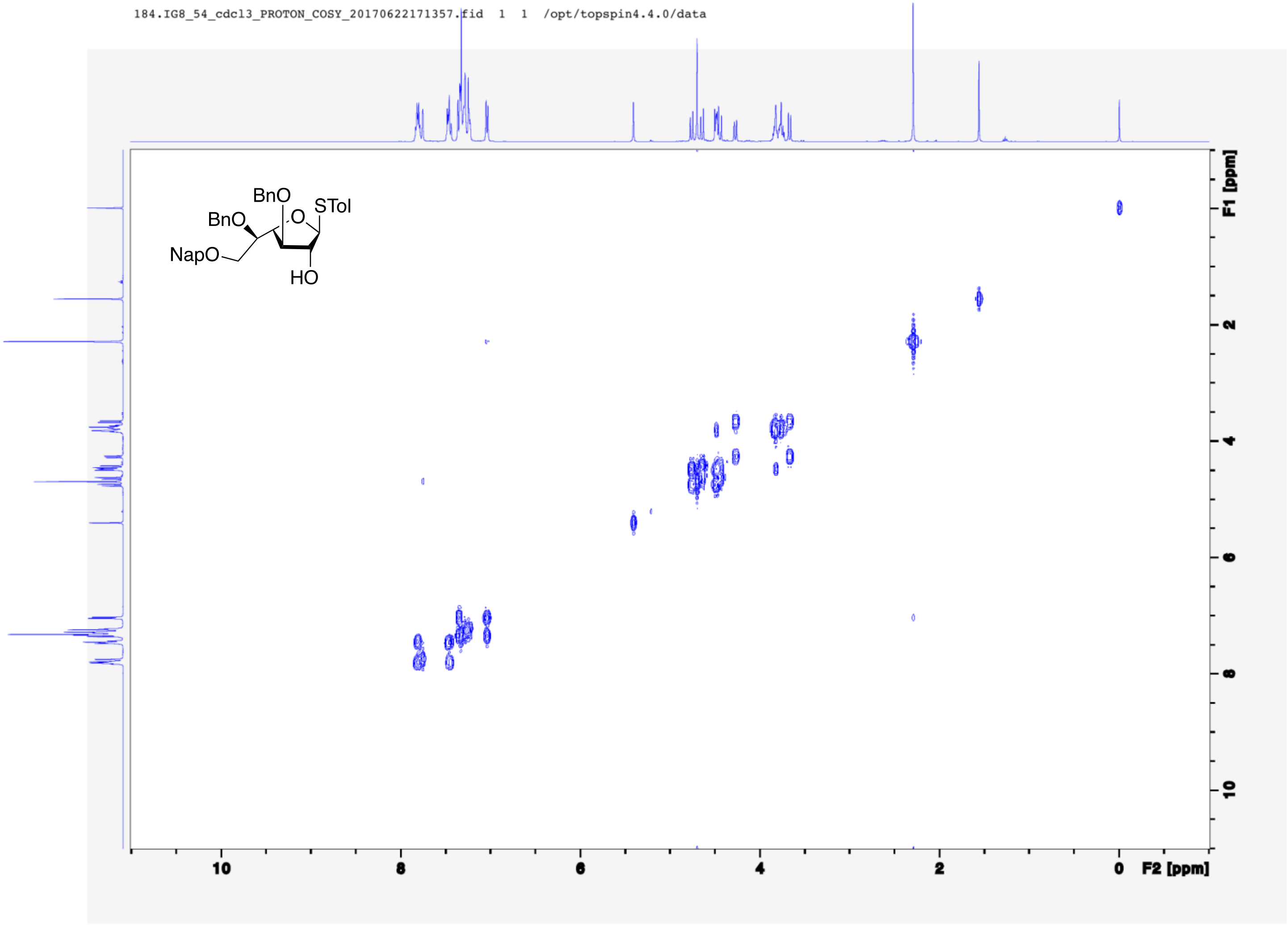
184.IG8_54_cdcl3_PROTON_COSY_20170622171357.fid 1 1 /opt/topspin4.4.0/data
BnO
BnO
O
STol
NapO
HO
F1 [ppm]
F2 [ppm]
10
8
6
4
2
0

182.IG8_54_cdcl3_CARBON_20170622192002.fid 1 1 /opt/topspin4.4.0/data

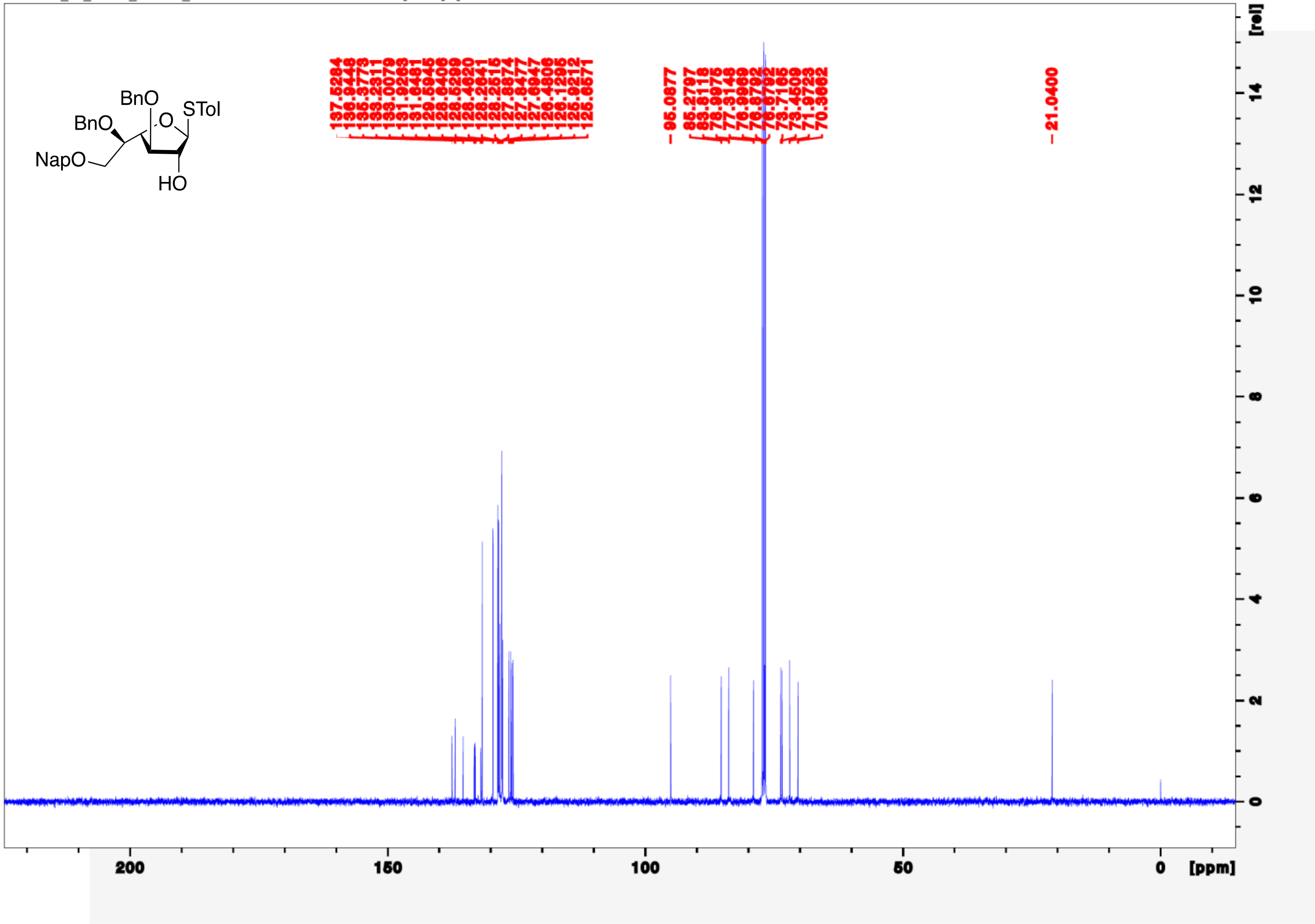

185.IG8_54_cdcl3_PROTON_gHSQC_20170622192034.fid 1 1 /opt/topspin4.4.0/data

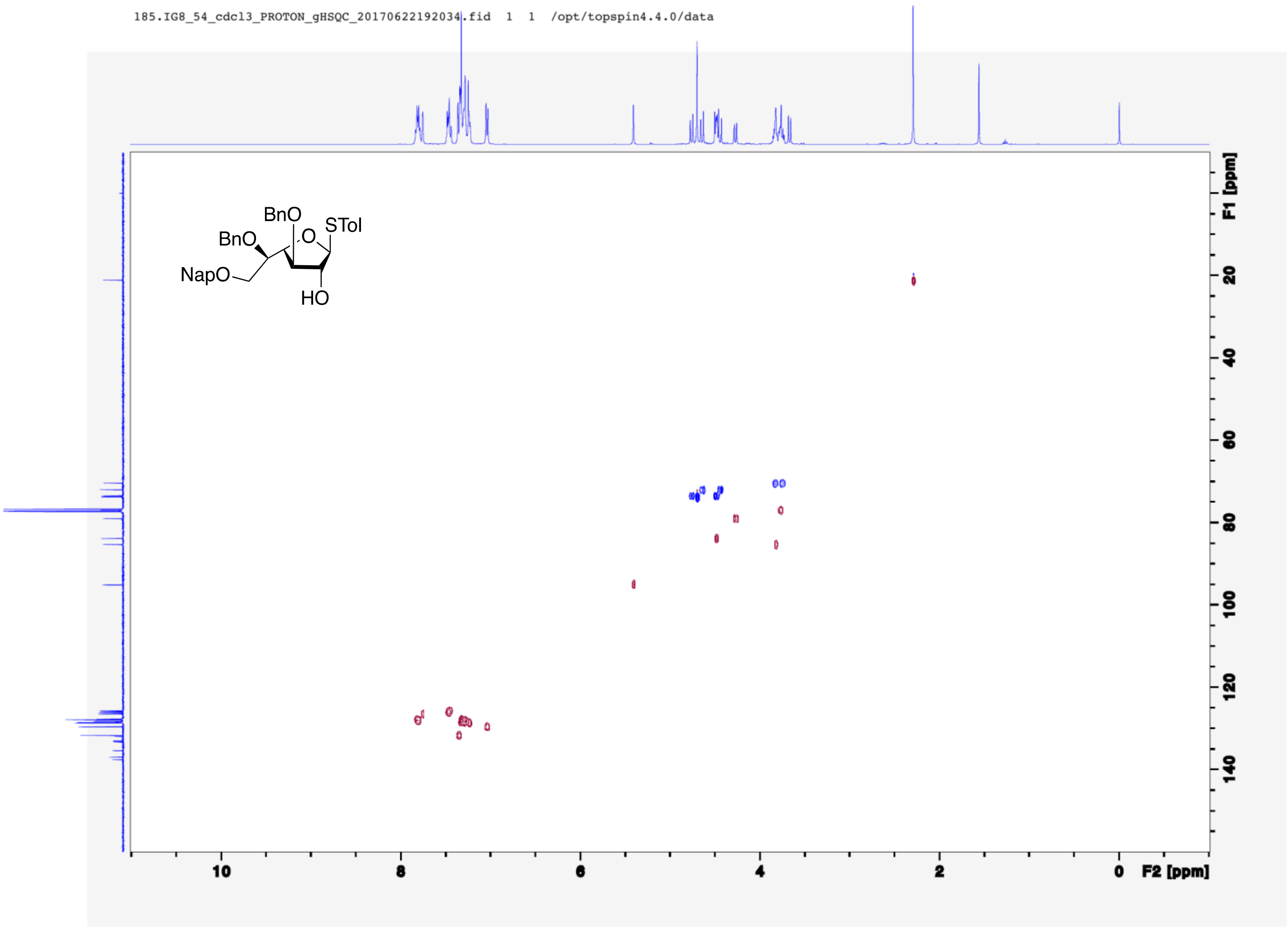

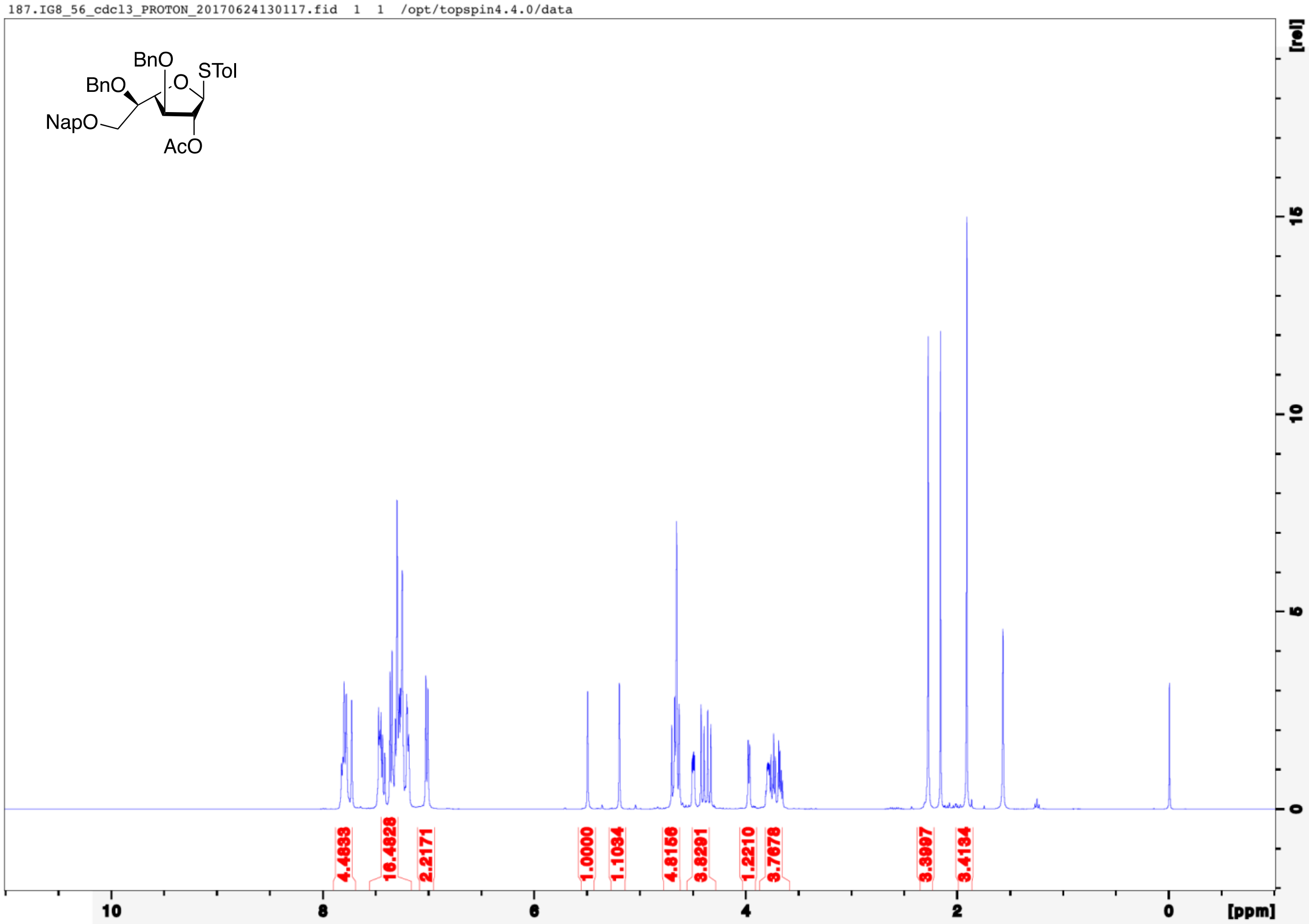

187.IG8_56_cdcl3_PROTON_20170624130117.fid 1 1 /opt/topspin4.4.0/data
BnO
STol
BnO
O
NapO
AcO
4.4933
16.4828
2.2171
1.0000
1.1034
4.8156
3.8291
1.2210
3.7678
3.3997
3.4134
10
8
6
4
2
0
[ppm]
[rel]
15
10
5
0

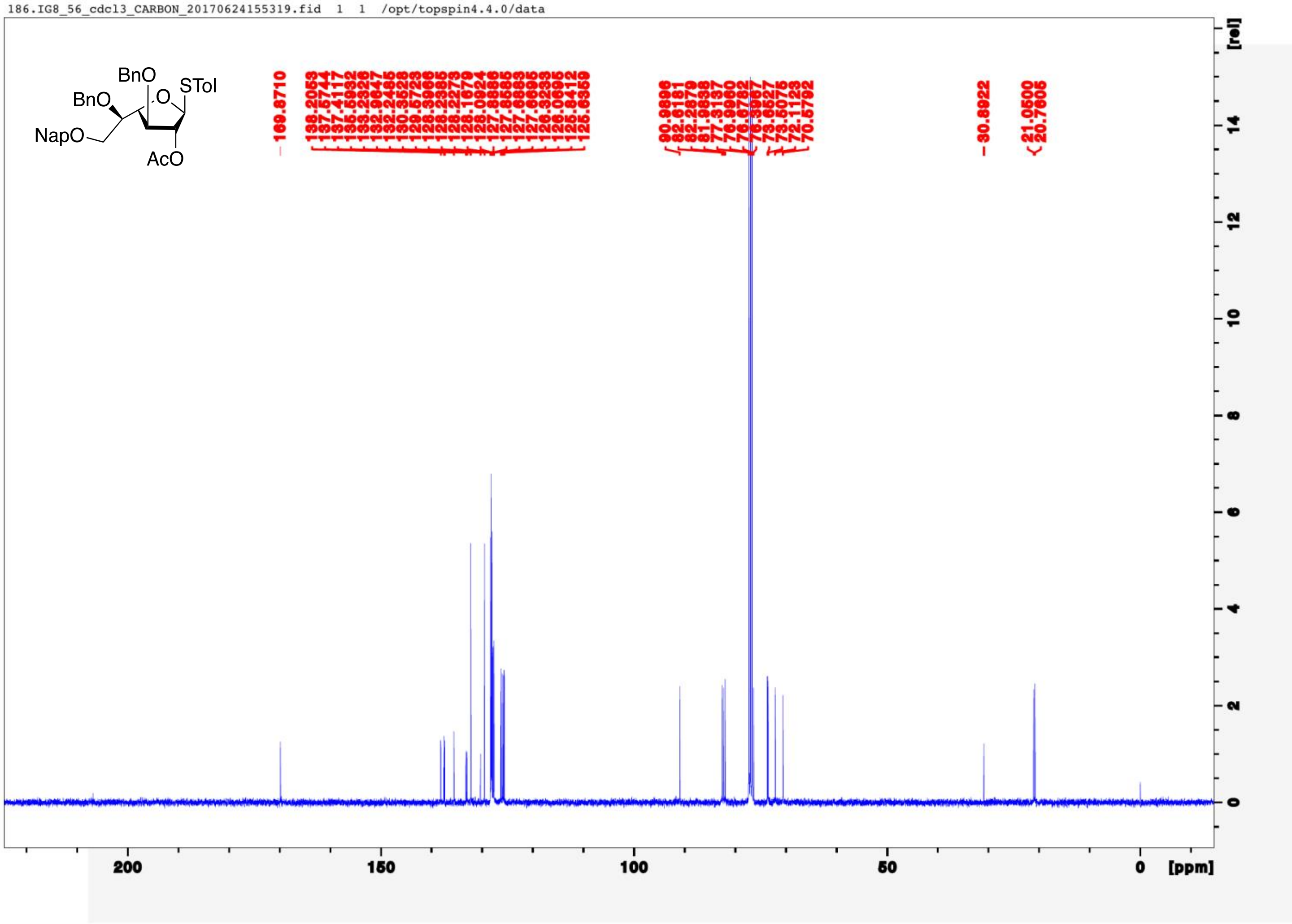
186.IG8_56_cdcl3_CARBON_20170624155319.fid 1 1 /opt/topspin4.4.0/data
BnO
BnO
STol
O
NapO
AcO
169.8710
138.2053
137.5744
137.4117
135.5932
133.2326
132.9647
132.2485
130.3528
129.5723
128.3966
128.2385
128.2273
128.1679
128.0924
127.8886
127.8585
127.6883
127.6695
126.3293
126.0695
125.8412
125.6359
90.9896
82.6181
82.2879
81.9838
77.3137
76.9960
76.6782
76.3967
73.6527
73.5075
72.1123
70.5792
30.8922
21.0500
20.7605
200
150
100
50
0
[ppm]
[rel]
14
12
10
8
6
4
2
0

188.IG8_56_cdcl3_PROTON_COSY_20170624155309.fid 1 1 /opt/topspin4.4.0/data

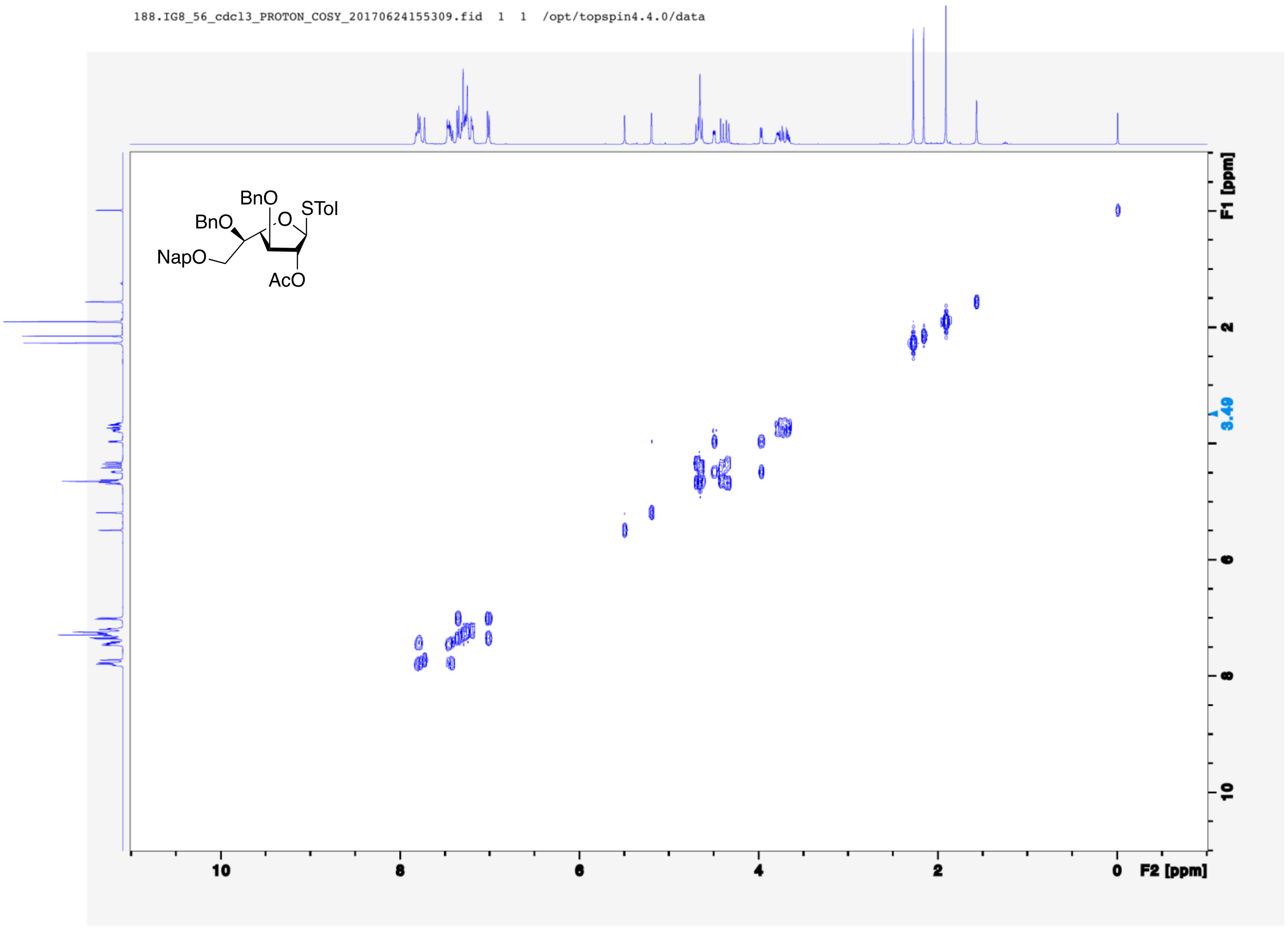

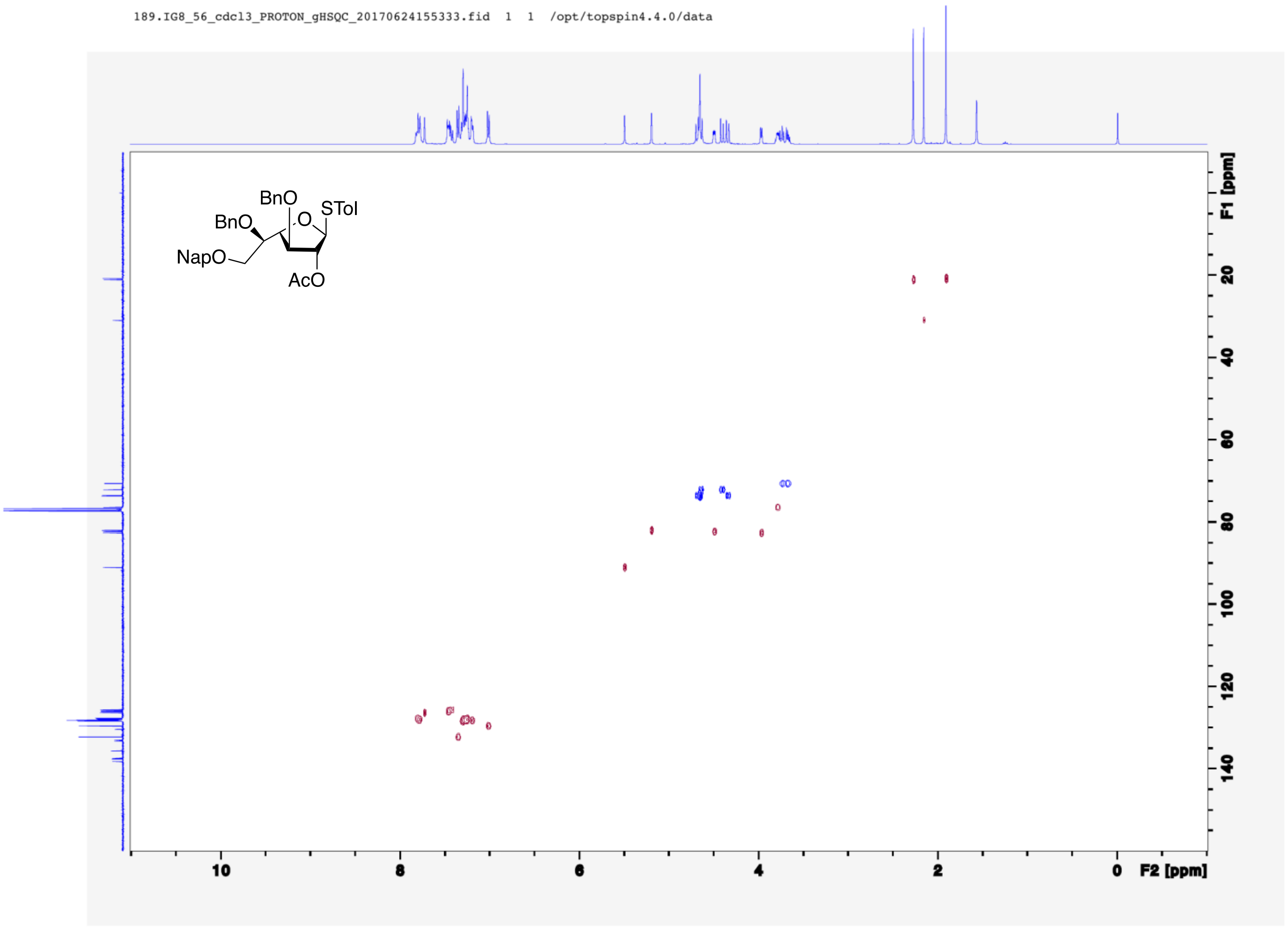
189.IG8_56_cdcl3_PROTON_gHSQC_20170624155333.fid 1 1 /opt/topspin4.4.0/data
BnO
BnO
O
STol
NapO
AcO
F1 [ppm]
20
40
60
80
100
120
140
10
8
6
4
2
0
F2 [ppm]

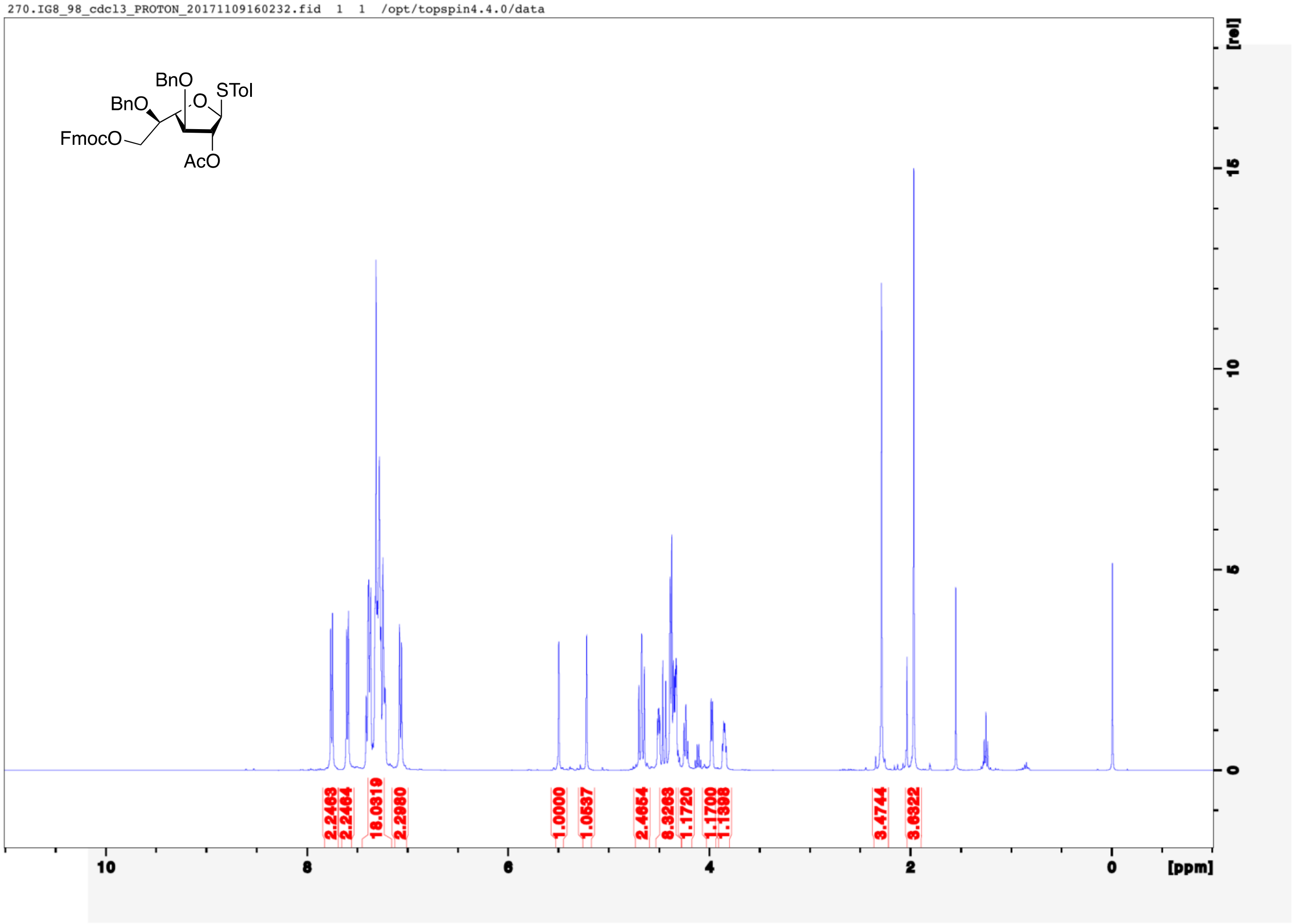

270.IG8_98_cdcl3_PROTON_20171109160232.fid 1 1 /opt/topspin4.4.0/data
BnO
BnO
O
STol
FmocO
AcO
[rel]
15
10
5
0
2.2463
2.2464
18.0319
2.2980
1.0000
1.0537
2.4654
8.3263
1.1720
1.1700
1.1398
3.4744
3.6322
10
8
6
4
2
0
[ppm]

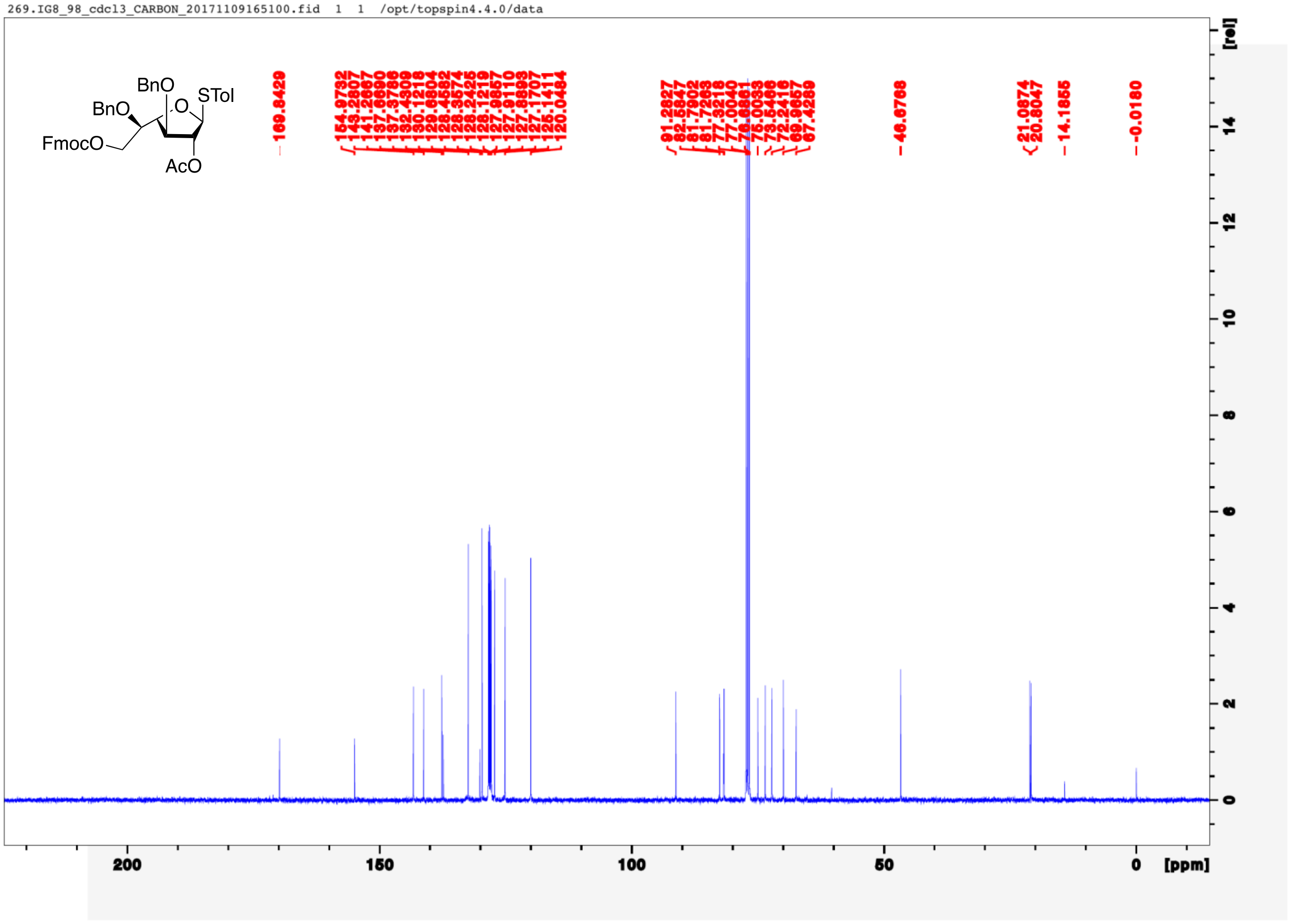
269.IG8_98_cdcl3_CARBON_20171109165100.fid 1 1 /opt/topspin4.4.0/data
BnO
BnO
O
STol
FmocO
AcO
169.8429
154.9732
143.2807
141.2667
137.6690
137.3786
132.4309
130.1218
129.6804
128.4582
128.3574
128.2425
128.1219
127.9857
127.9110
127.8893
127.1707
125.1411
120.0484
91.2827
82.5847
81.7902
81.7263
77.3218
77.0040
76.6861
75.0033
73.5466
72.2416
69.9657
67.4289
46.6768
21.0874
20.8047
14.1855
-0.0180
200
150
100
50
0
[ppm]
[rel]

271.IG8_98_cdcl3_PROTON_gCOSY_20171109164951.fid 1 1 /opt/topspin4.4.0/data

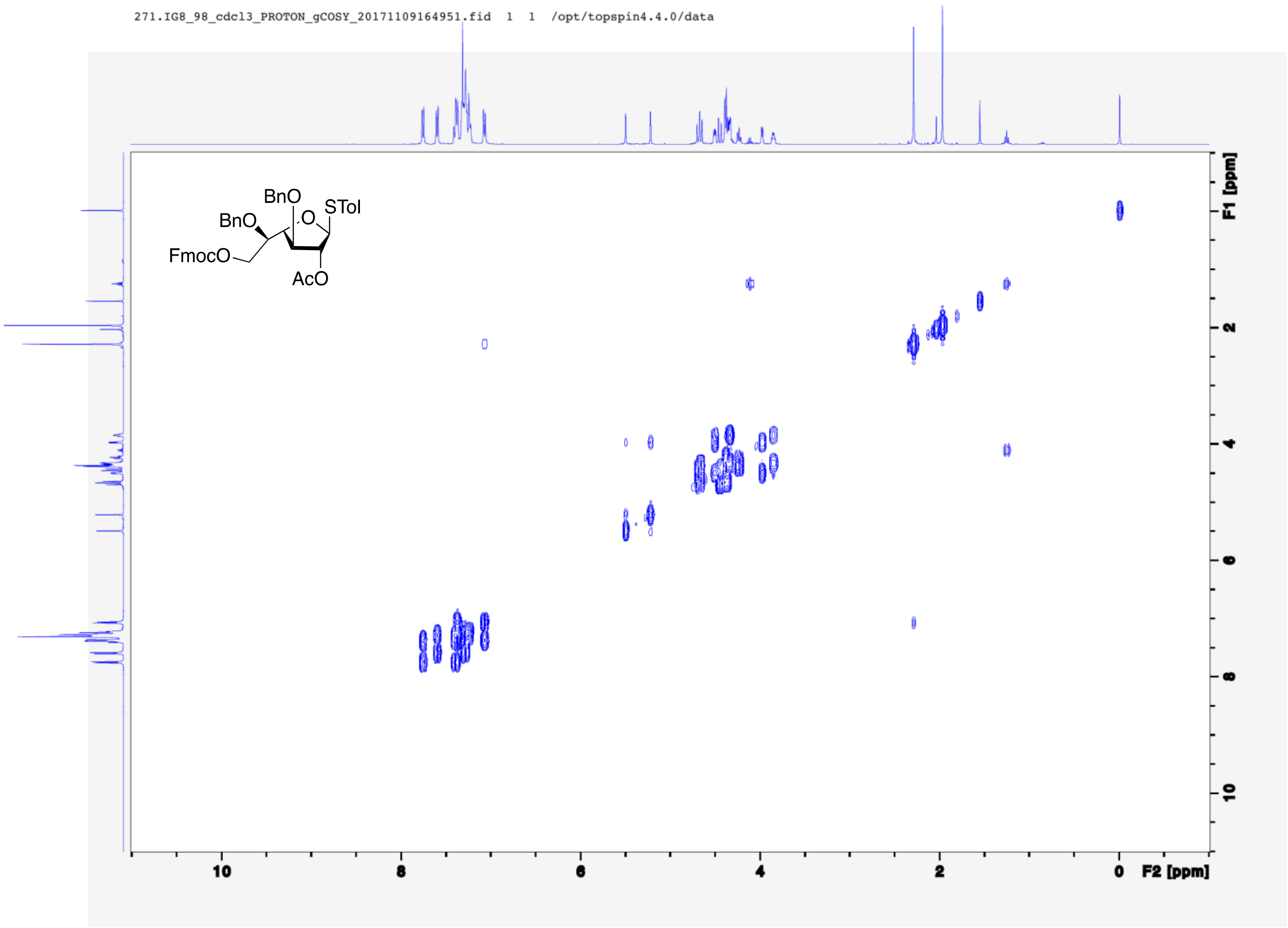

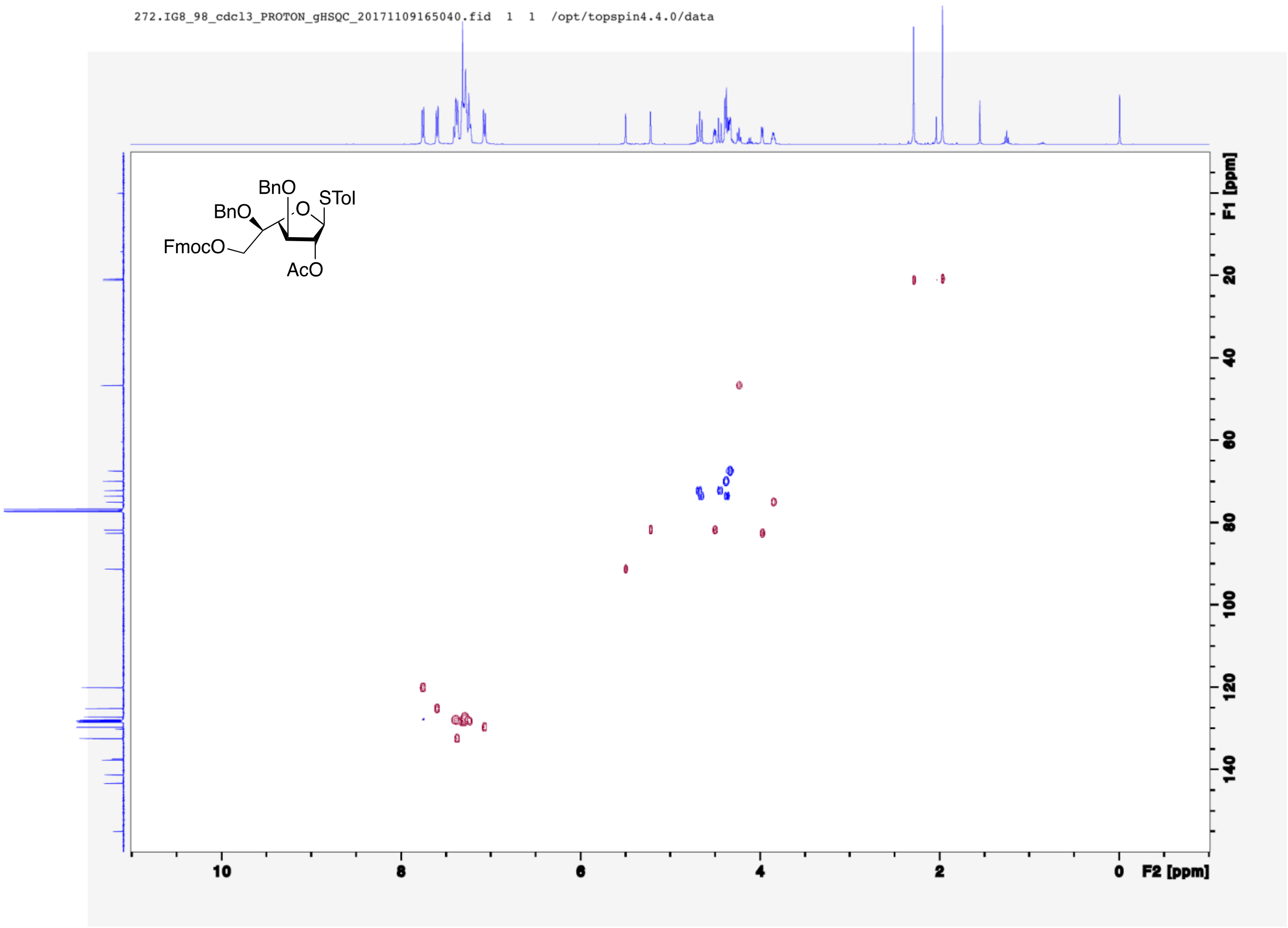
272.IG8_98_cdcl3_PROTON_gHSQC_20171109165040.fid 1 1 /opt/topspin4.4.0/data
BnO
BnO
STol
O
FmocO
AcO
F1 [ppm]
20
40
60
80
100
120
140
10
8
6
4
2
0
F2 [ppm]

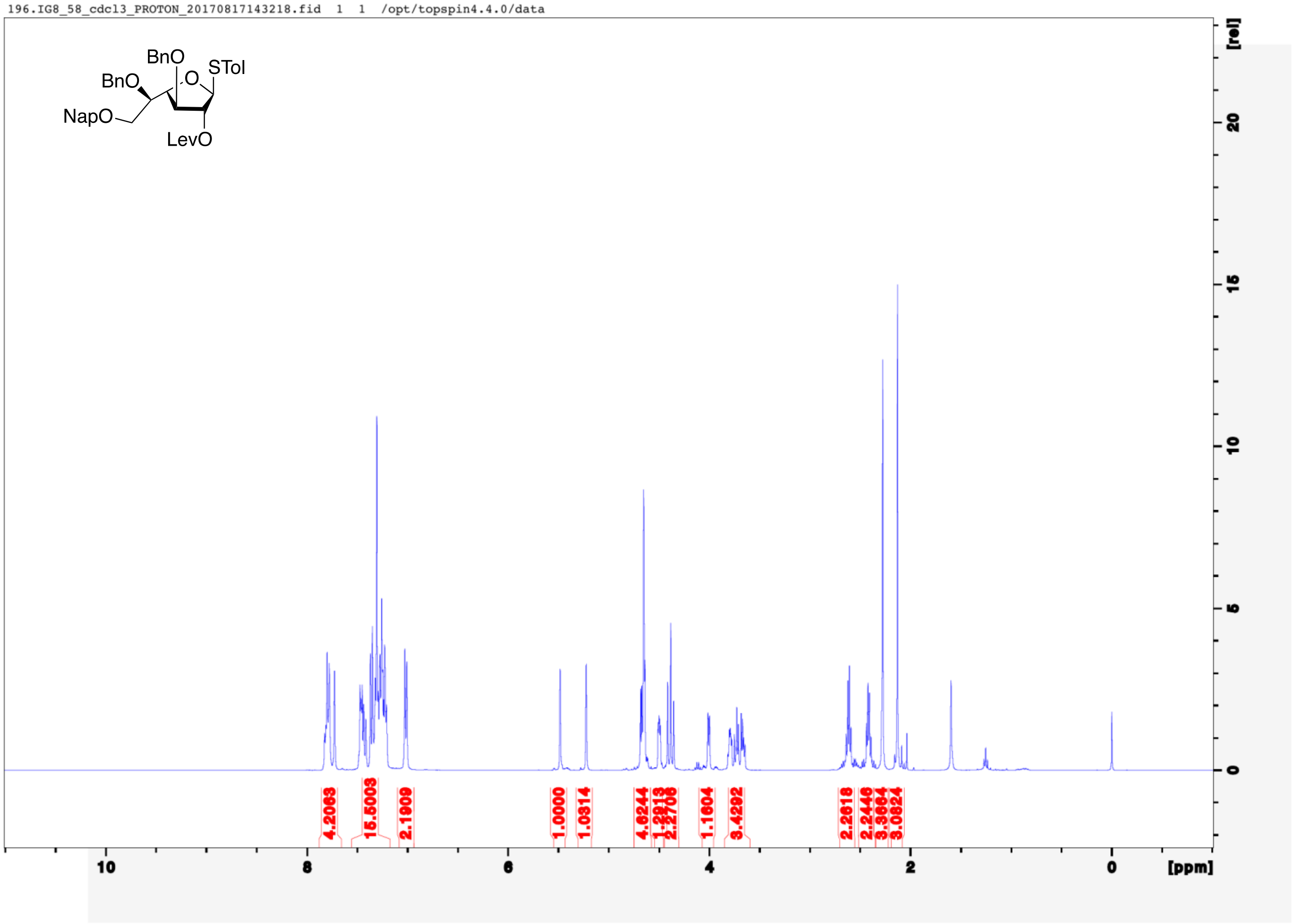
196.IG8_58_cdcl3_PROTON_20170817143218.fid 1 1 /opt/topspin4.4.0/data
BnO
BnO
O
STol
NapO
LevO
4.2063
15.5003
2.1909
1.0000
1.0314
4.6244
1.2913
2.2706
1.1604
3.4292
2.2618
2.2446
3.3664
3.0824
10
8
6
4
2
0
[ppm]
[rel]
20
15
10
5
0

194.IG8_58_cdcl3_CARBON_20170817142047.fid 1 1 /opt/topspin4.4.0/data

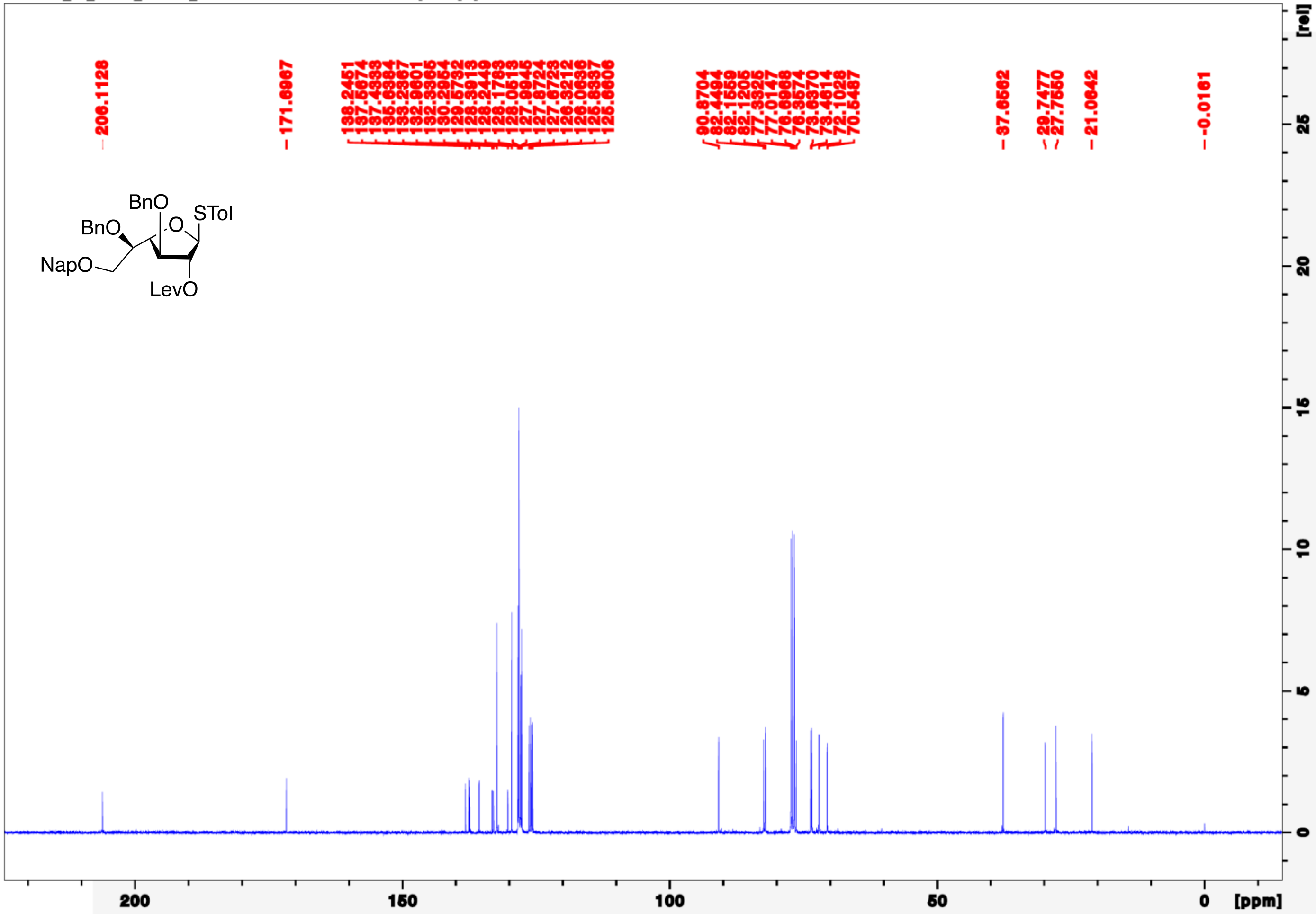

197.IG8_58_cdcl3_PROTON_COSY_20170817142028.fid 1 1 /opt/topspin4.4.0/data

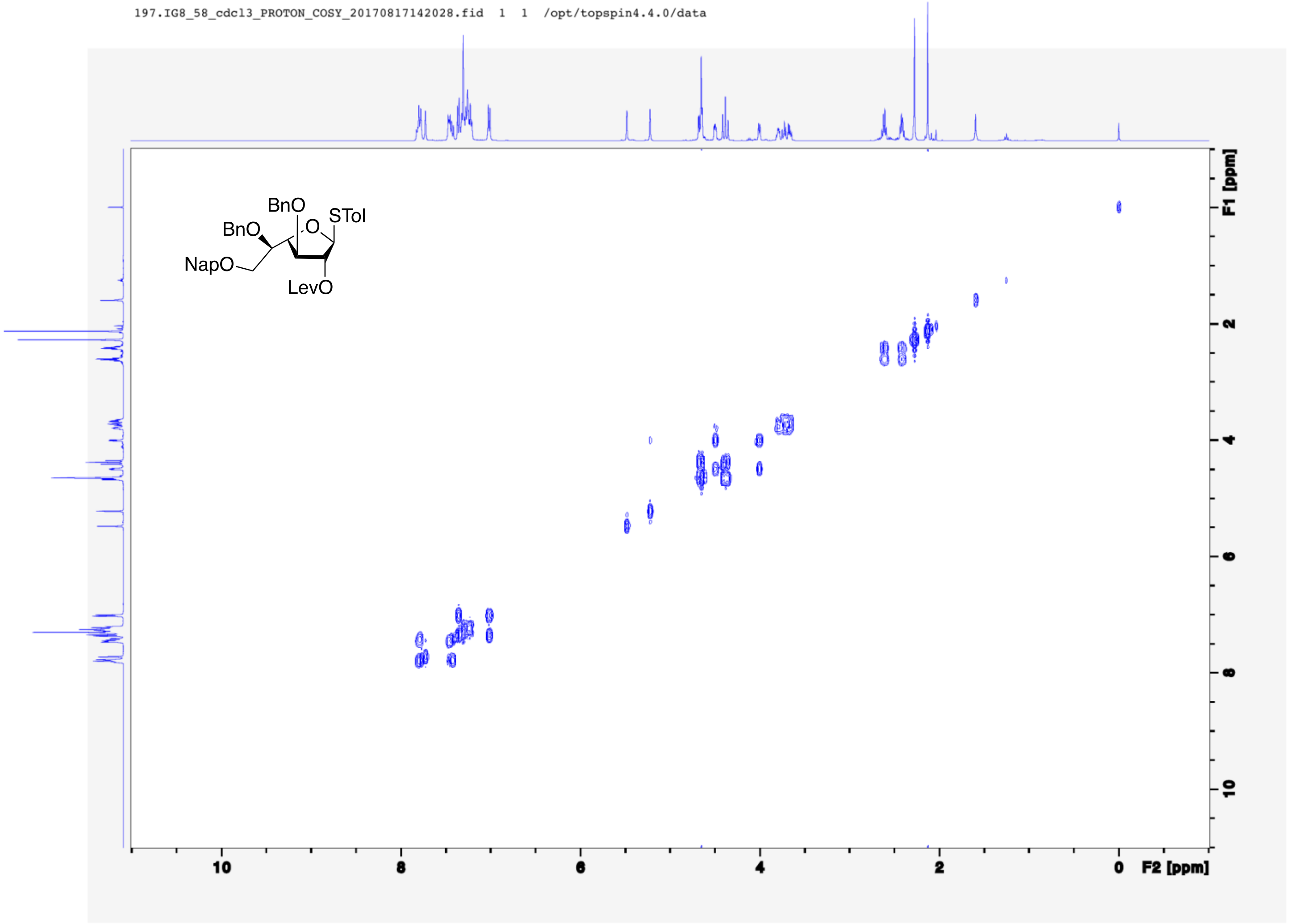

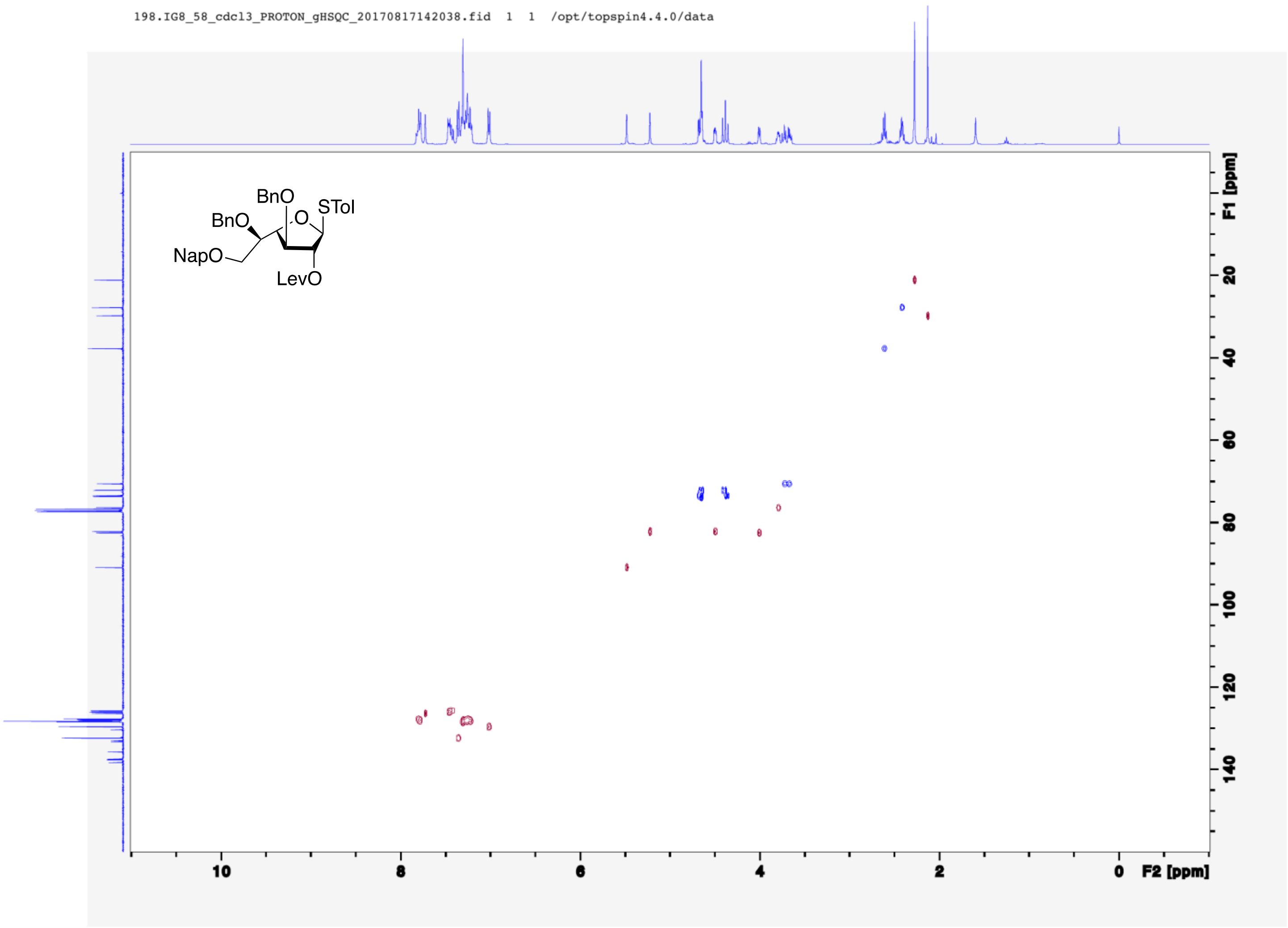
198.IG8_58_cdcl3_PROTON_gHSQC_20170817142038.fid 1 1 /opt/topspin4.4.0/data
BnO
BnO
O
STol
NapO
LevO
F1 [ppm]
20
40
60
80
100
120
140
10
8
6
4
2
0
F2 [ppm]

231.IG8_70_cdcl3_PROTON_20170818184419.fid 1 1 /opt/topspin4.4.0/data

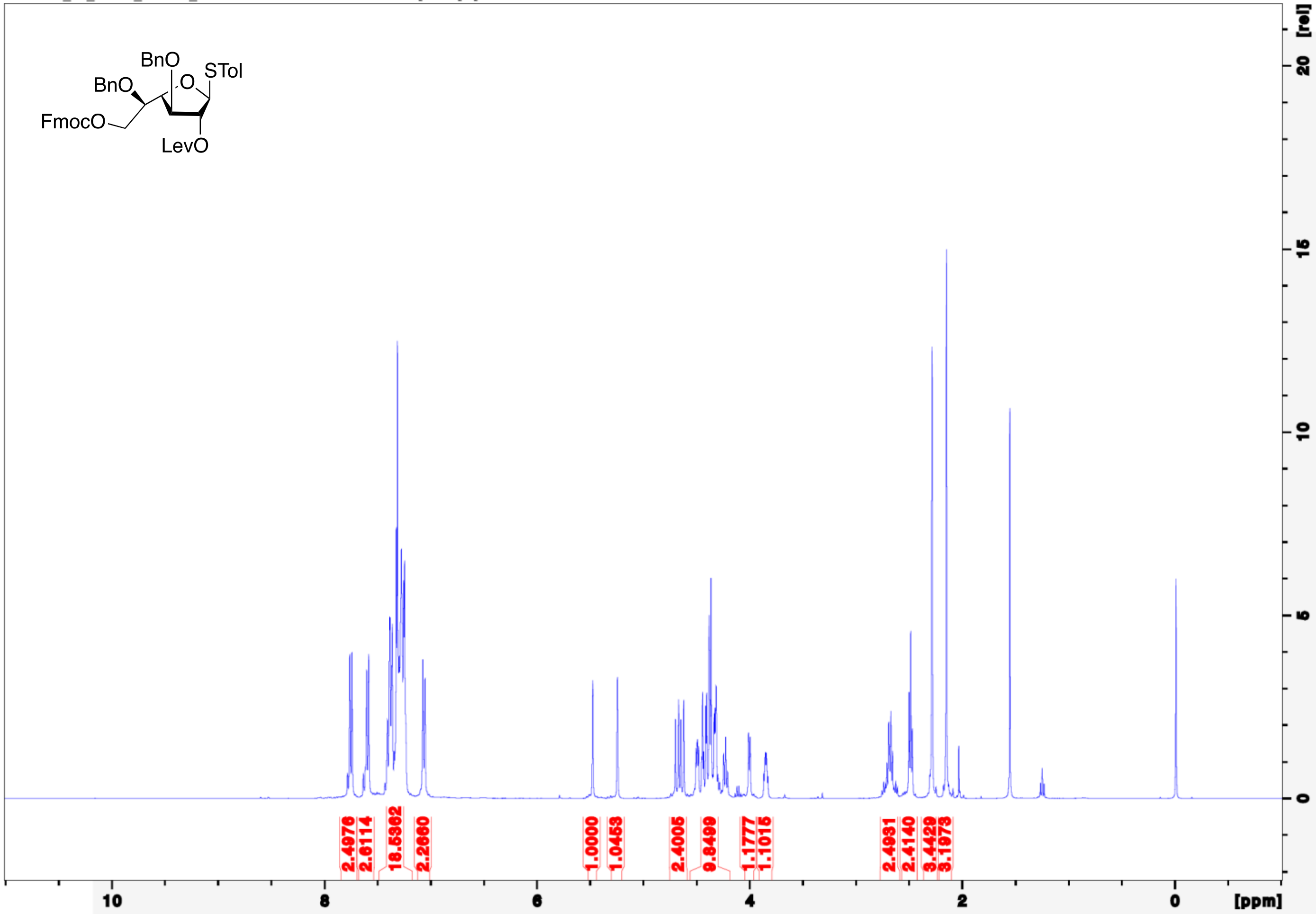

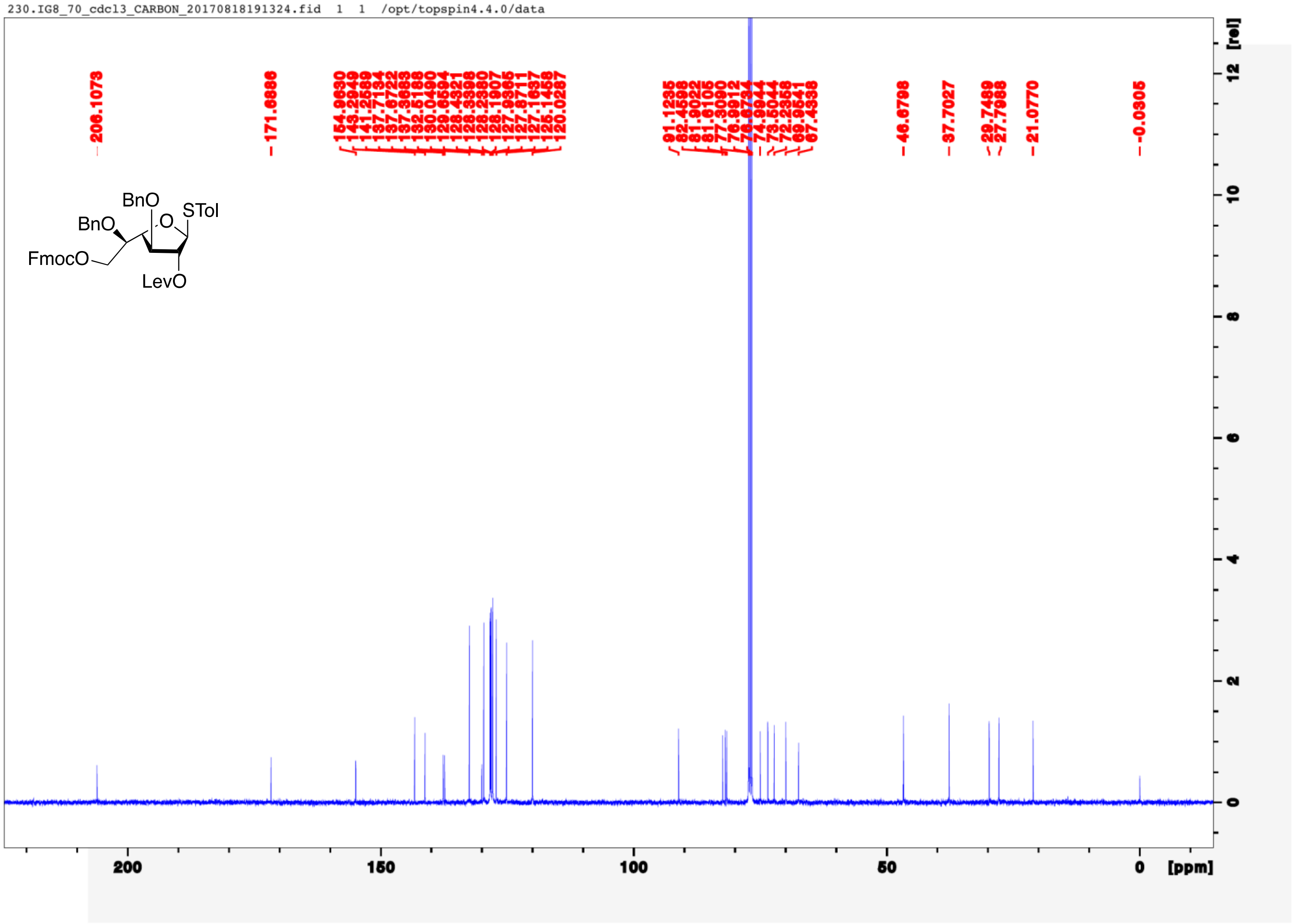
230.IG8_70_cdcl3_CARBON_20170818191324.fid 1 1 /opt/topspin4.4.0/data
206.1073
171.6886
154.9630
143.2949
141.2589
137.7134
137.6722
137.3683
132.5188
130.0490
129.6594
128.4321
128.3398
128.2380
128.1907
127.9365
127.8711
127.1637
125.1458
120.0287
91.1235
82.4598
81.9022
81.6105
77.3090
76.9912
76.6734
74.9944
73.5044
72.2358
69.9541
67.4338
46.6798
37.7027
29.7489
27.7988
21.0770
-0.0305
BnO
BnO
STol
O
FmocO
LevO
[rel]
12
10
8
6
4
2
0
200
150
100
50
0
[ppm]

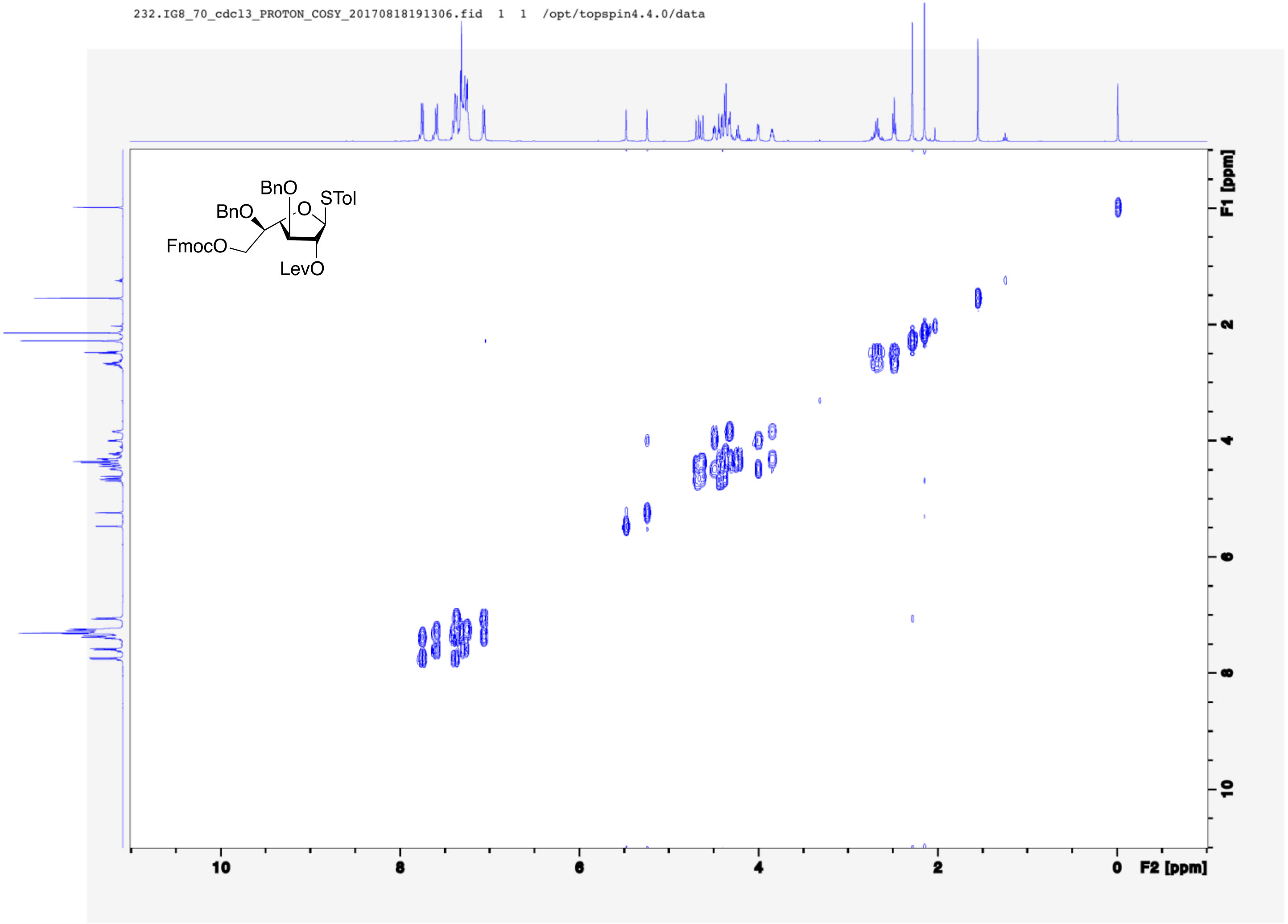
232.IG8_70_cdcl3_PROTON_COSY_20170818191306.fid 1 1 /opt/topspin4.4.0/data
BnO
BnO
STol
O
FmocO
LevO
F1 [ppm]
2
4
6
8
10
10
8
6
4
2
0
F2 [ppm]

233.IG8_70_cdcl3_PROTON_gHSQC_20170818191315.fid 1 1 /opt/topspin4.4.0/data

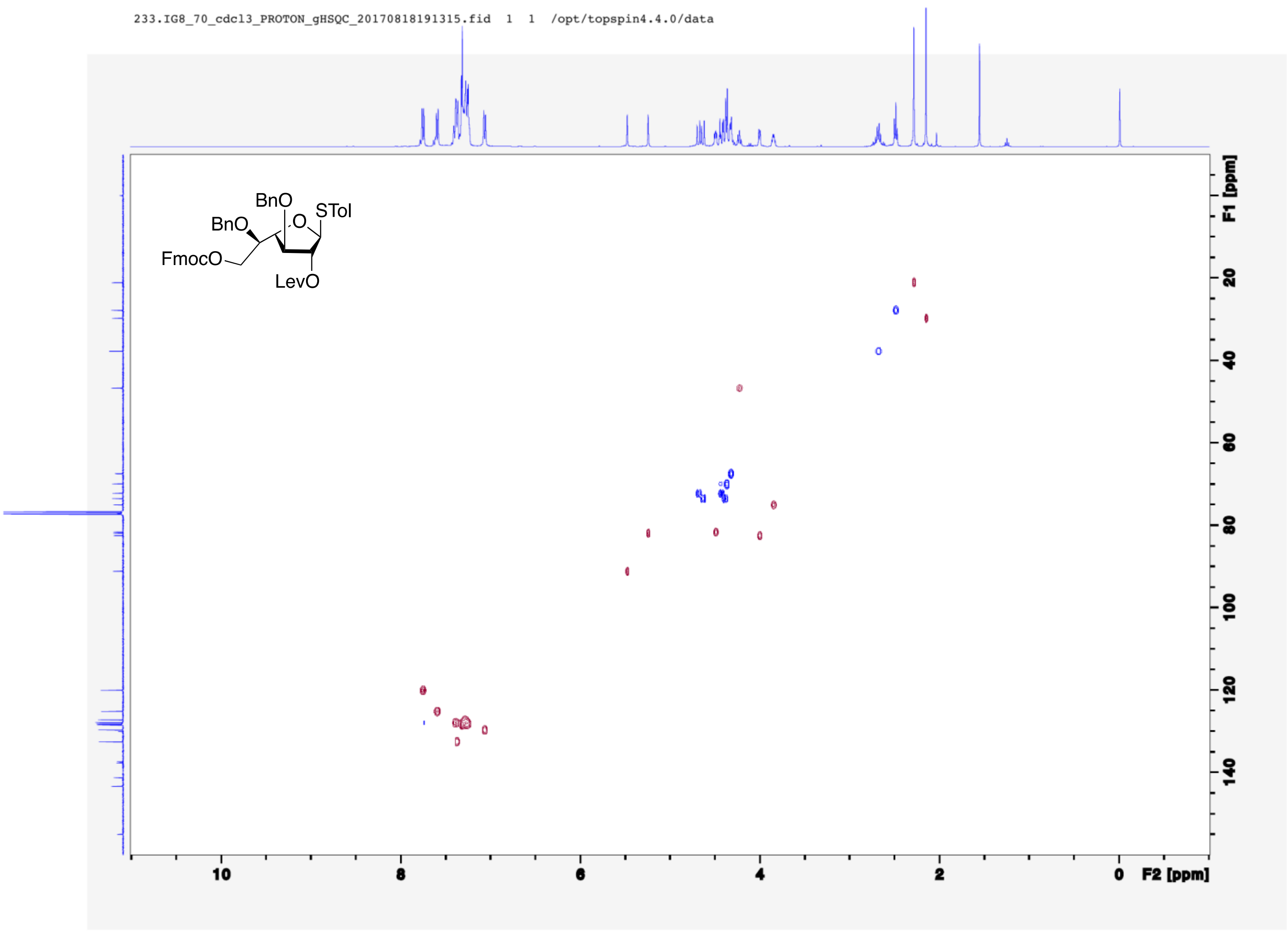

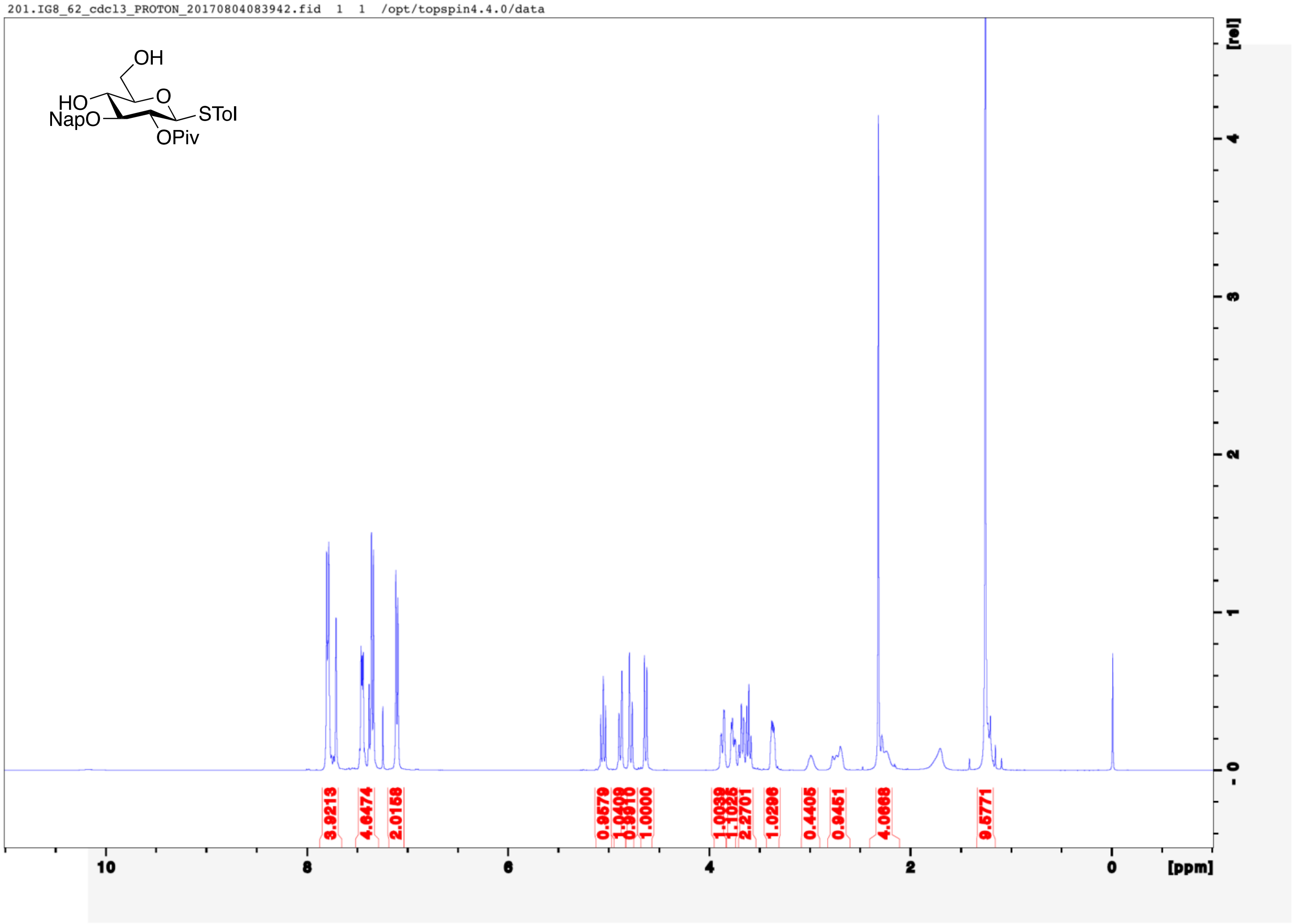
201.IG8_62_cdcl3_PROTON_20170804083942.fid 1 1 /opt/topspin4.4.0/data
OH
HO
O
NapO
STol
OPiv
3.9213
4.6474
2.0158
0.9579
1.0408
0.9810
1.0000
1.0039
1.1025
2.2701
1.0296
0.4405
0.9451
4.0668
9.5771
10
8
6
4
2
0
[ppm]
[rel]

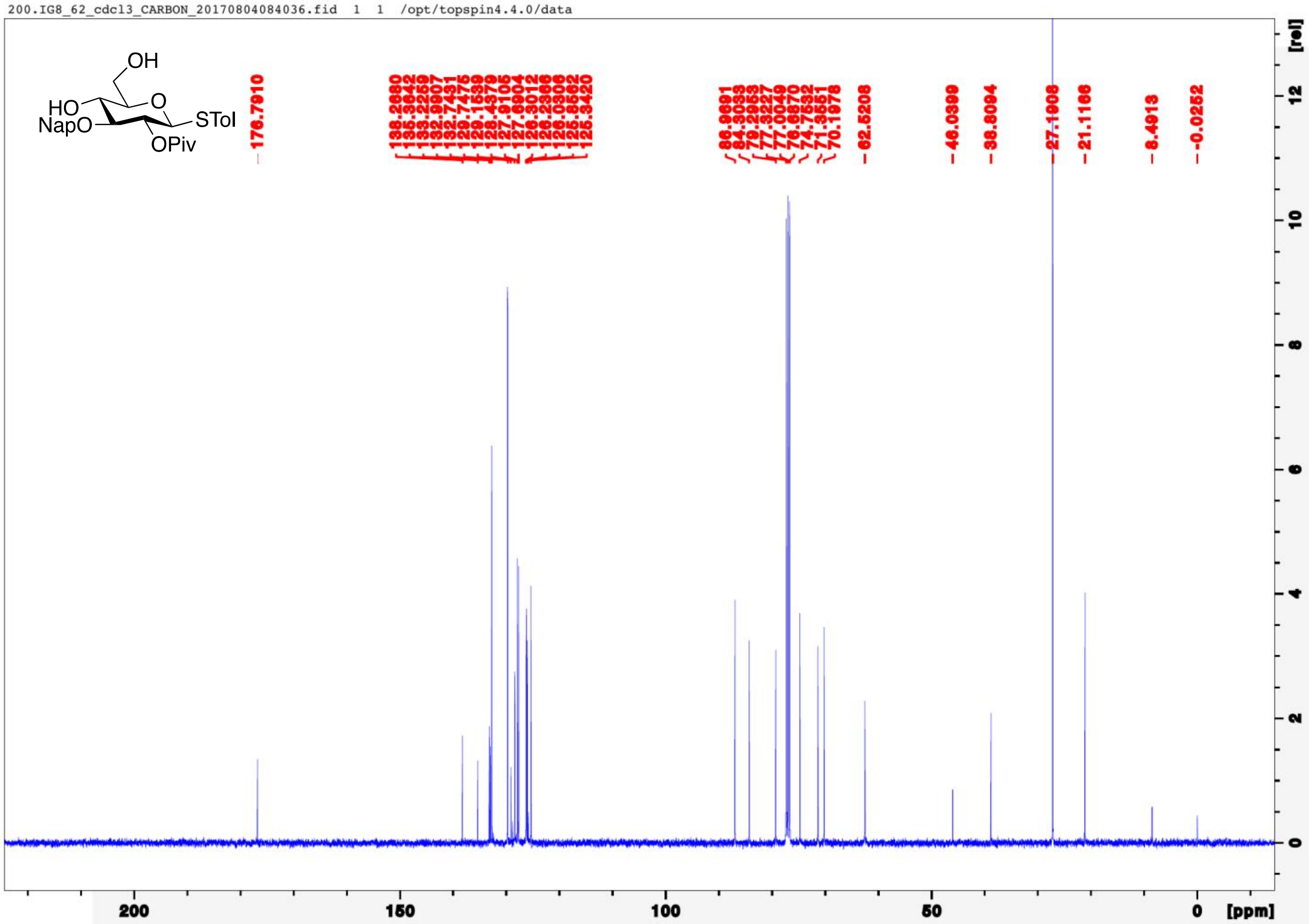
200.IG8_62_cdcl3_CARBON_20170804084036.fid 1 1 /opt/topspin4.4.0/data
OH
HO
O
NapO
STol
OPiv
176.7910
138.2680
135.3642
133.2259
132.9907
132.7431
129.7475
129.1539
128.4379
127.9105
127.6904
126.3012
126.2366
126.0306
125.8562
125.3420
85.9691
84.3033
79.2953
77.3227
77.0049
76.6870
74.7532
71.3551
70.1978
62.5208
46.0399
38.8094
27.1908
21.1166
8.4913
-0.0252
[rel]
[ppm]

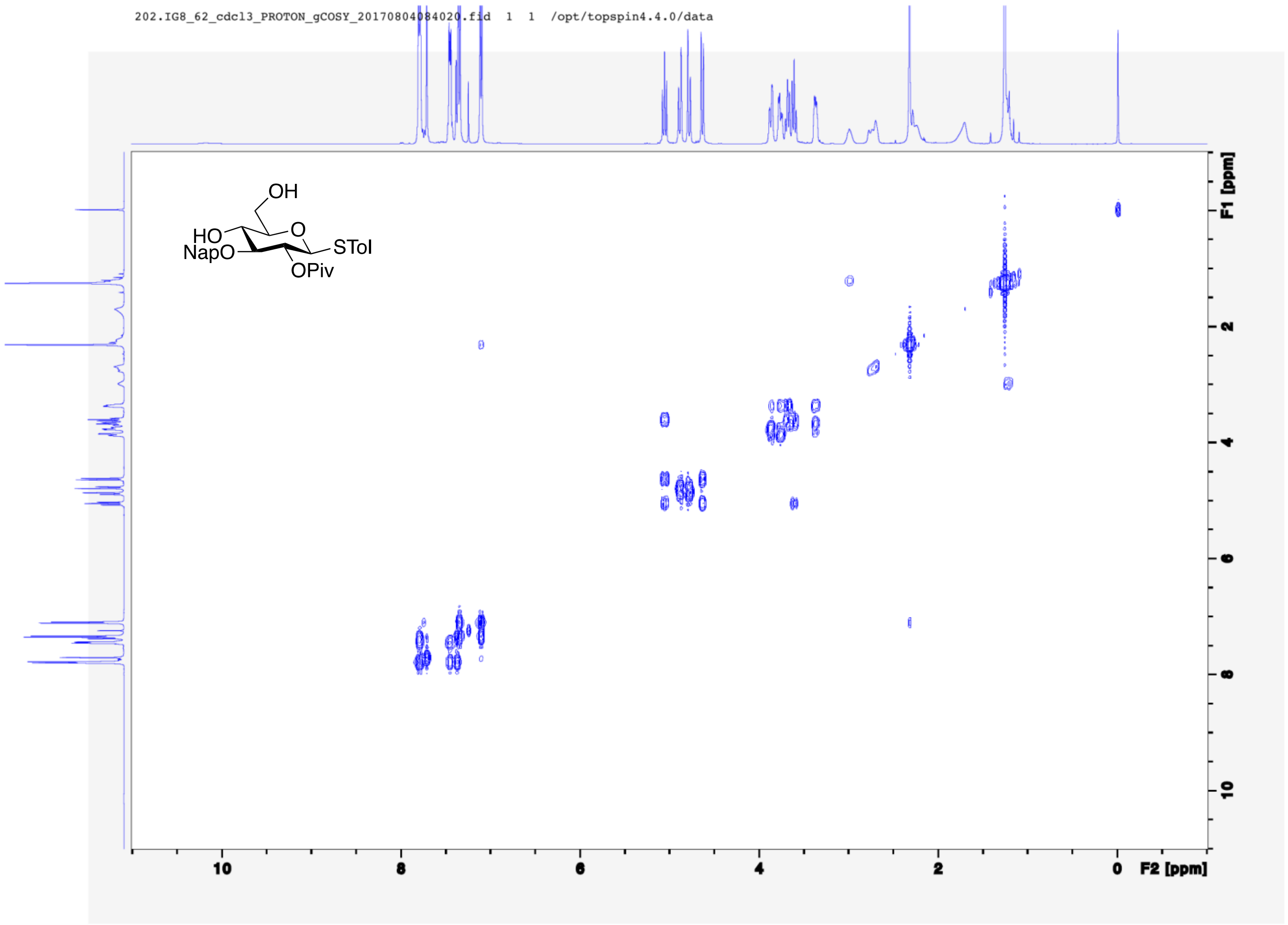
202.IG8_62_cdcl3_PROTON_gCOSY_20170804084020.fid 1 1 /opt/topspin4.4.0/data
OH
HO
O
NapO
STol
OPiv
F1 [ppm]
F2 [ppm]
10
8
6
4
2
0

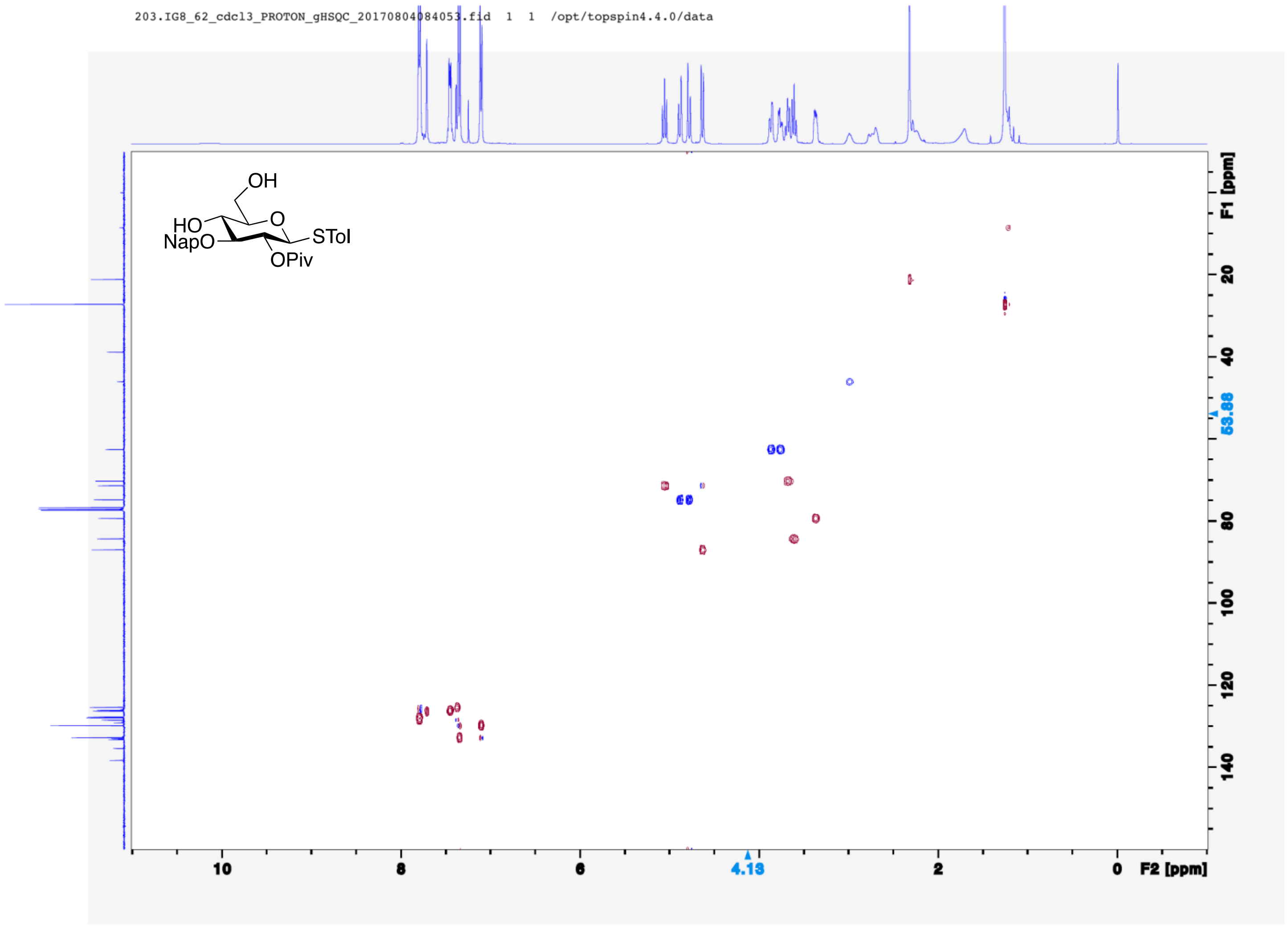
203.IG8_62_cdcl3_PROTON_gHSQC_20170804084053.fid 1 1 /opt/topspin4.4.0/data
OH
HO
O
NapO
STol
OPiv
F1 [ppm]
20
40
53.88
80
100
120
140
10
8
6
4.13
2
0
F2 [ppm]

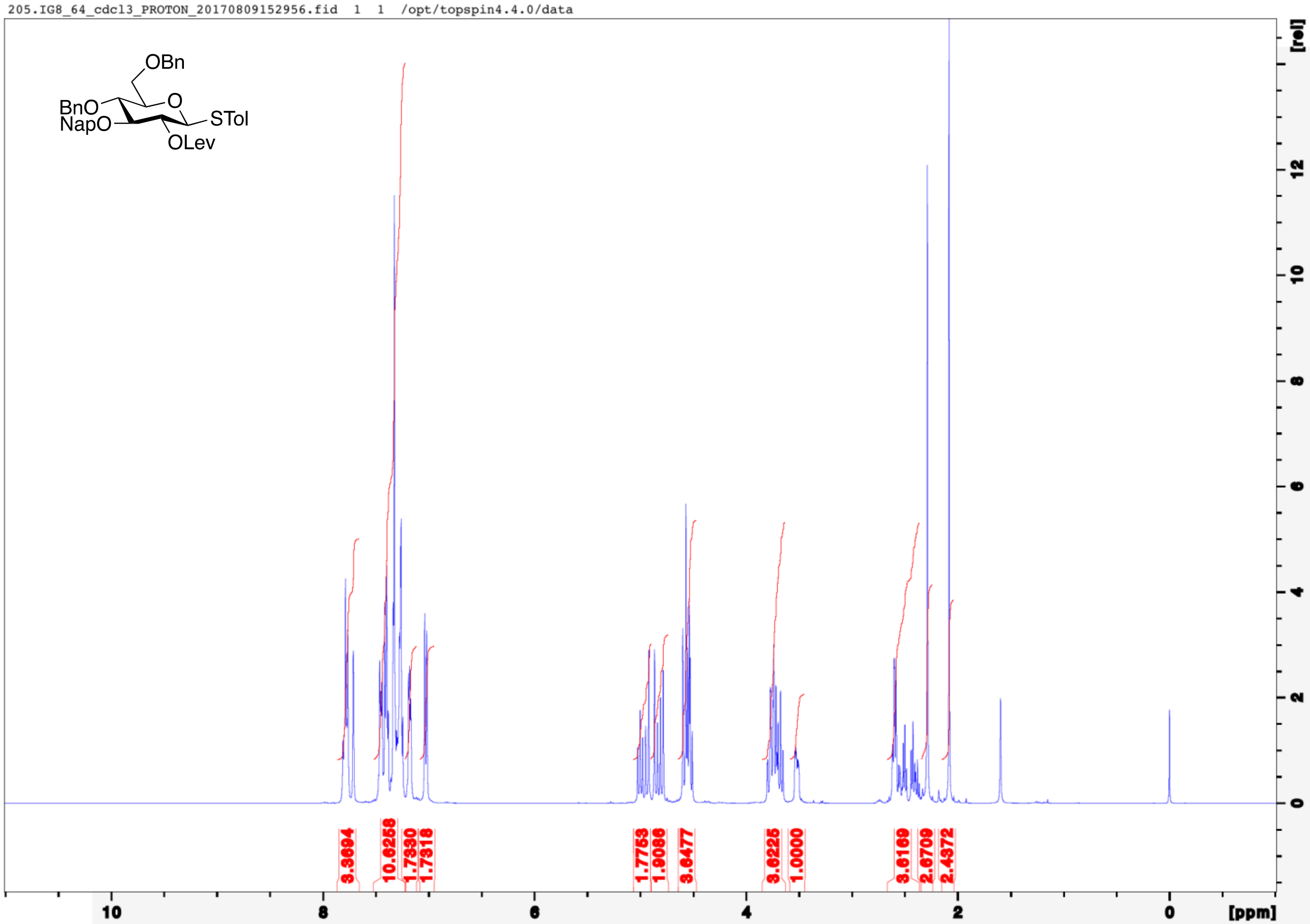

205.IG8_64_cdcl3_PROTON_20170809152956.fid 1 1 /opt/topspin4.4.0/data
OBn
BnO
NapO
O
STol
OLev
3.3604
10.6256
1.7330
1.7318
1.7753
1.9086
3.6477
3.6225
1.0000
3.6169
2.6709
2.4372
[rel]
[ppm]

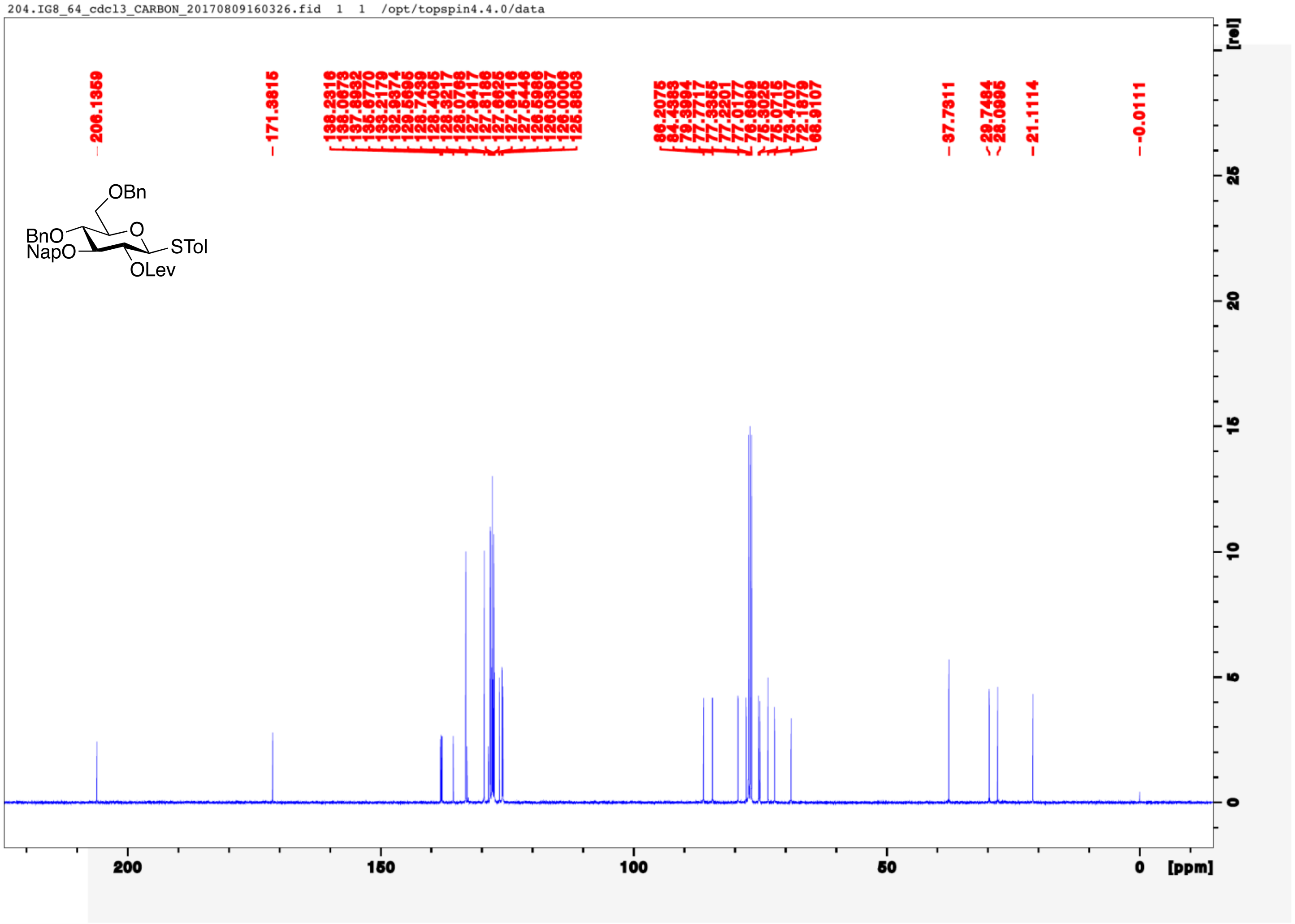
204.IG8_64_cdcl3_CARBON_20170809160326.fid 1 1 /opt/topspin4.4.0/data
206.1359
171.3815
138.2316
138.0673
137.8932
135.6770
133.2179
132.9374
129.5695
128.4095
128.7439
128.3217
128.0768
127.9418
127.8186
127.6625
127.6416
127.5446
126.5986
126.0397
126.0006
125.8803
86.2075
84.4383
79.3994
77.7717
77.3855
77.2201
77.0177
76.6999
75.3025
75.0715
73.4707
72.1879
68.9107
37.7311
29.7484
28.0896
21.1114
-0.0111
OBn
BnO
NapO
O
STol
OLev
[rel]
25
20
15
10
5
0
200
150
100
50
0
[ppm]

206.IG8_64_cdcl3_PROTON_COSY_20170809160256.fid 1 1 /opt/topspin4.4.0/data

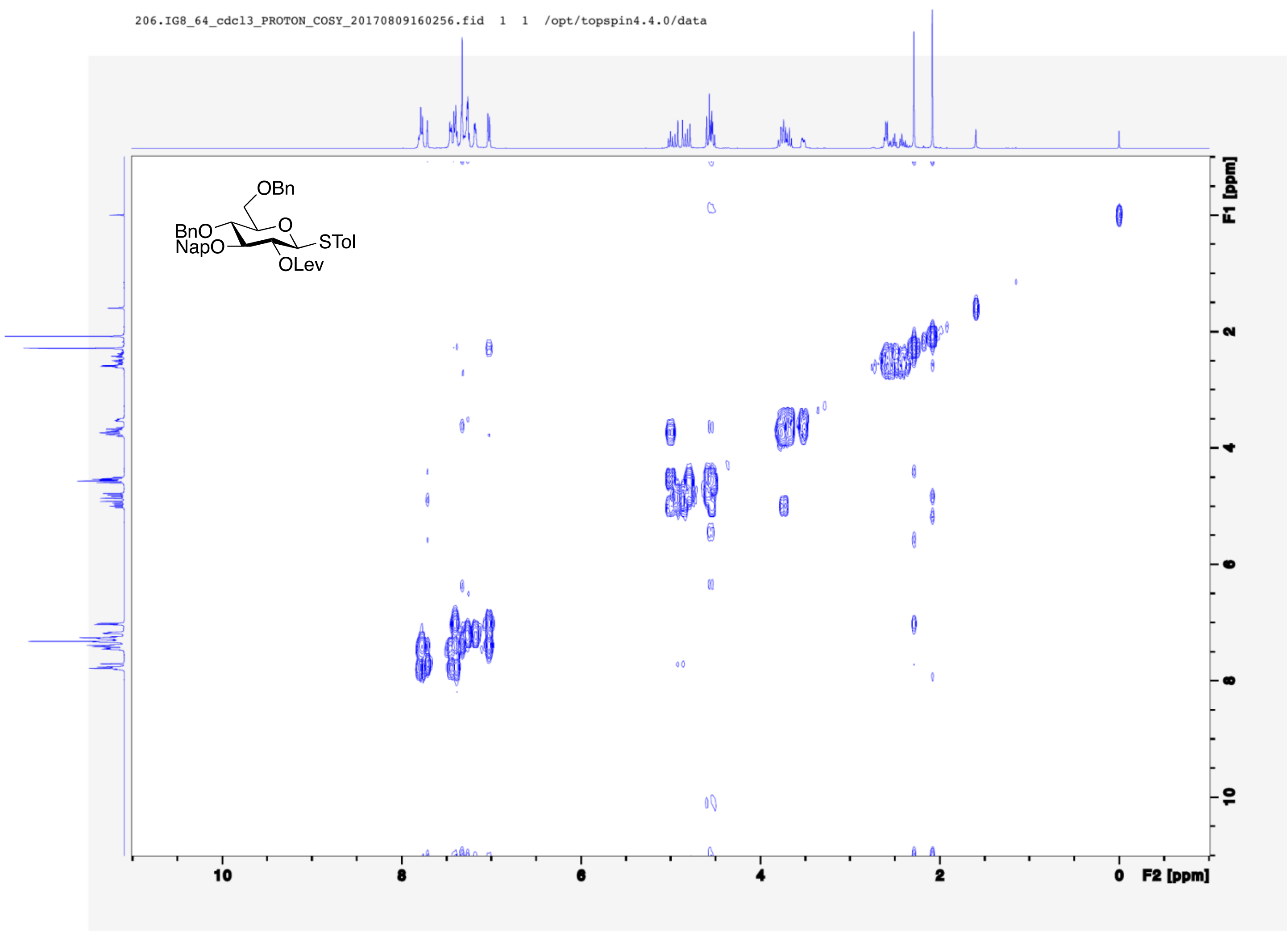

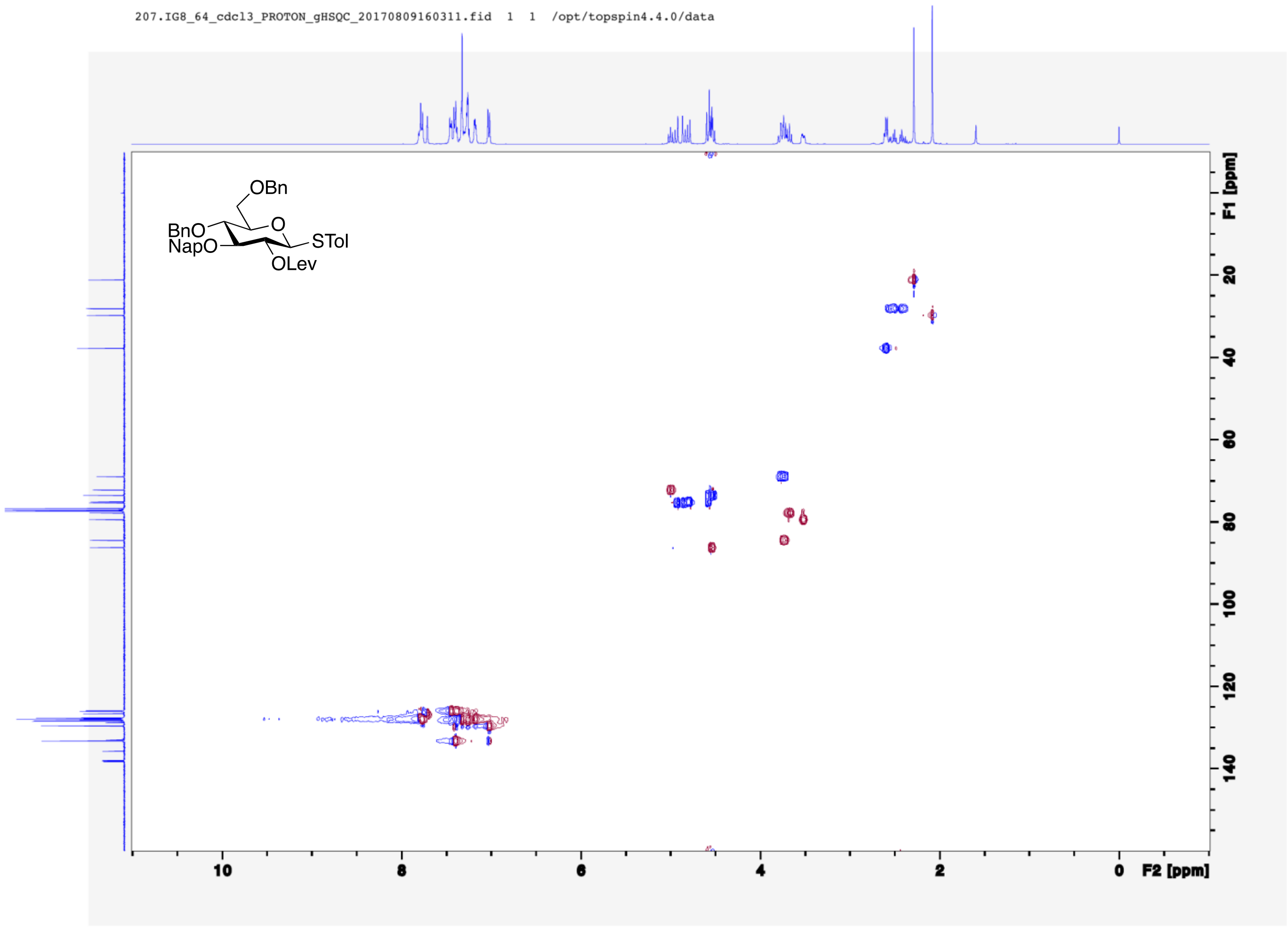
207.IG8_64_cdcl3_PROTON_gHSQC_20170809160311.fid 1 1 /opt/topspin4.4.0/data
OBn
O
BnO
NapO
STol
OLev
F1 [ppm]
20
40
60
80
100
120
140
10
8
6
4
2
0
F2 [ppm]

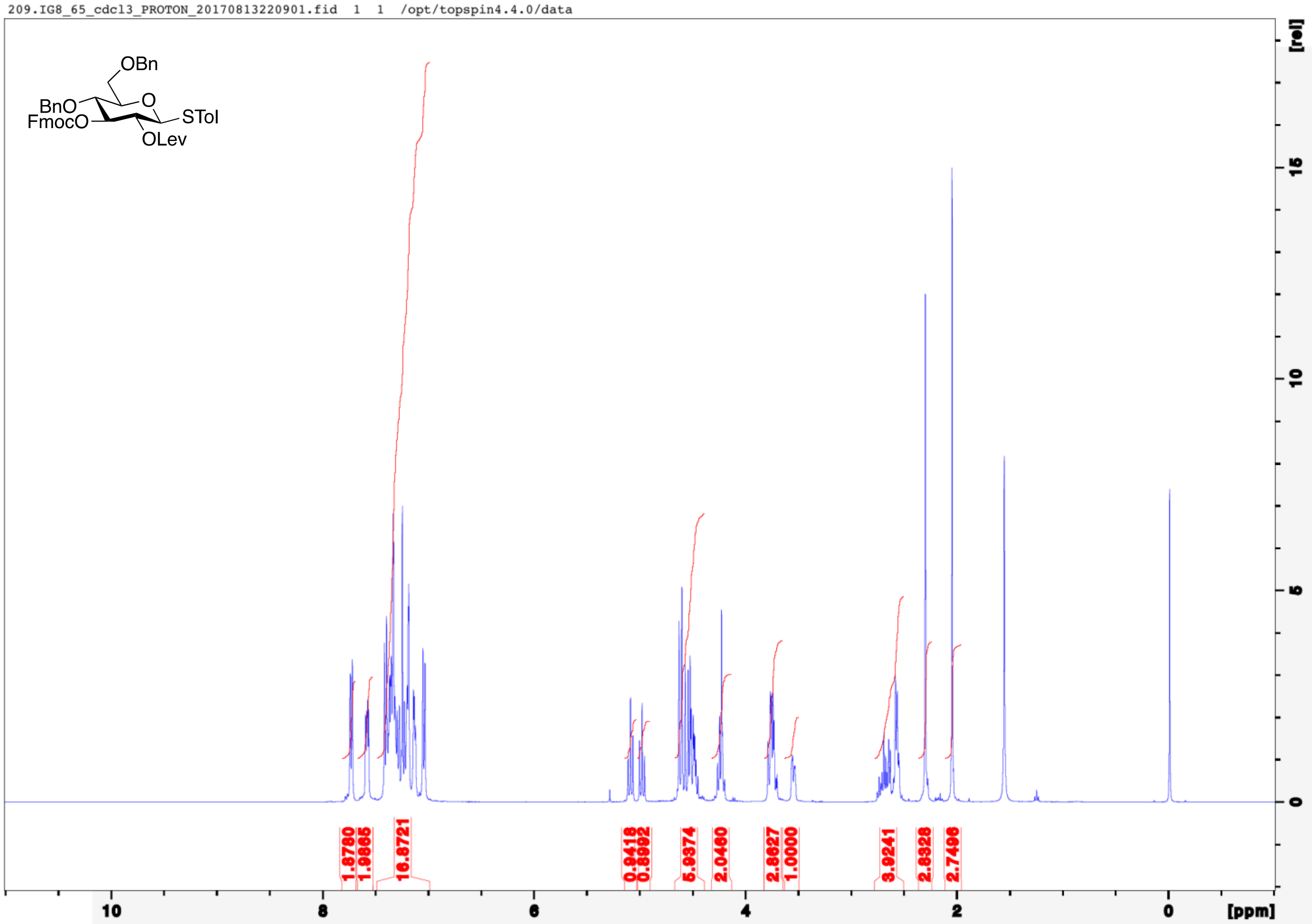
209.IG8_65_cdcl3_PROTON_20170813220901.fid 1 1 /opt/topspin4.4.0/data
OBn
BnO
O
FmocO
STol
OLev
[rel]
[ppm]

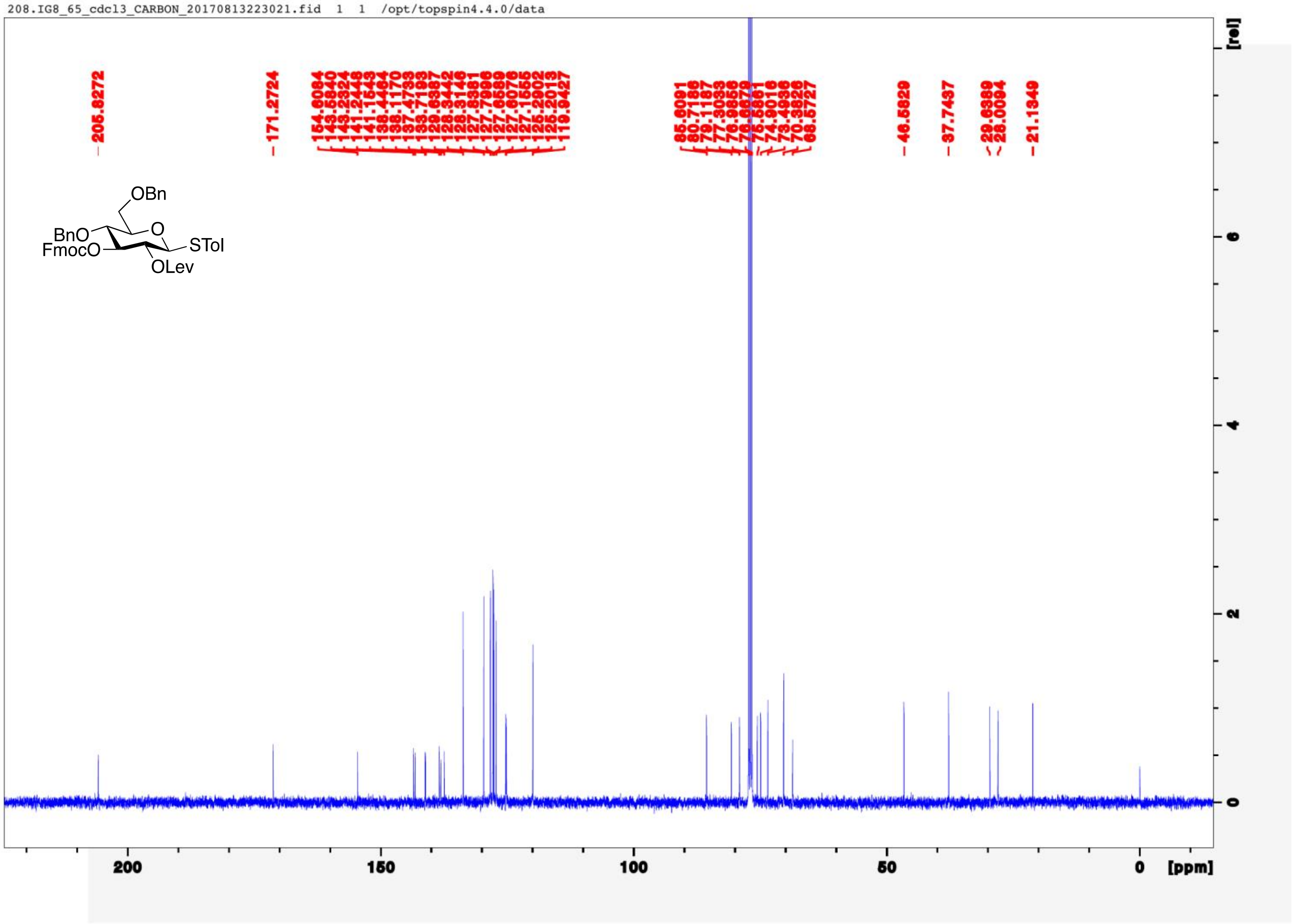
208.IG8_65_cdcl3_CARBON_20170813223021.fid 1 1 /opt/topspin4.4.0/data
OBn
BnO
O
FmocO
STol
OLev
[ppm]
[rel]

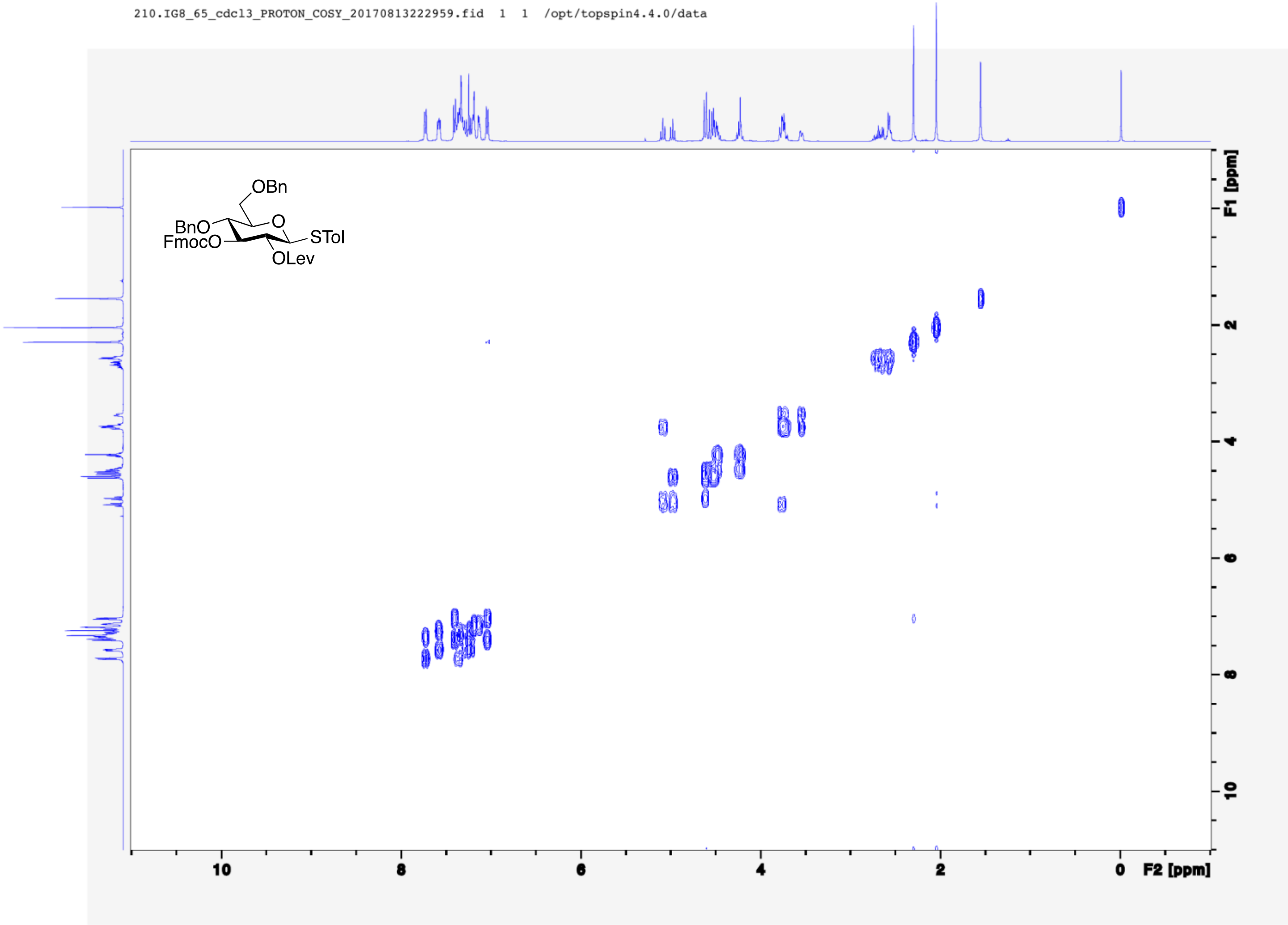
210.IG8_65_cdcl3_PROTON_COSY_20170813222959.fid 1 1 /opt/topspin4.4.0/data
OBn
BnO
FmocO
O
STol
OLev
F1 [ppm]
F2 [ppm]
10
8
6
4
2
0

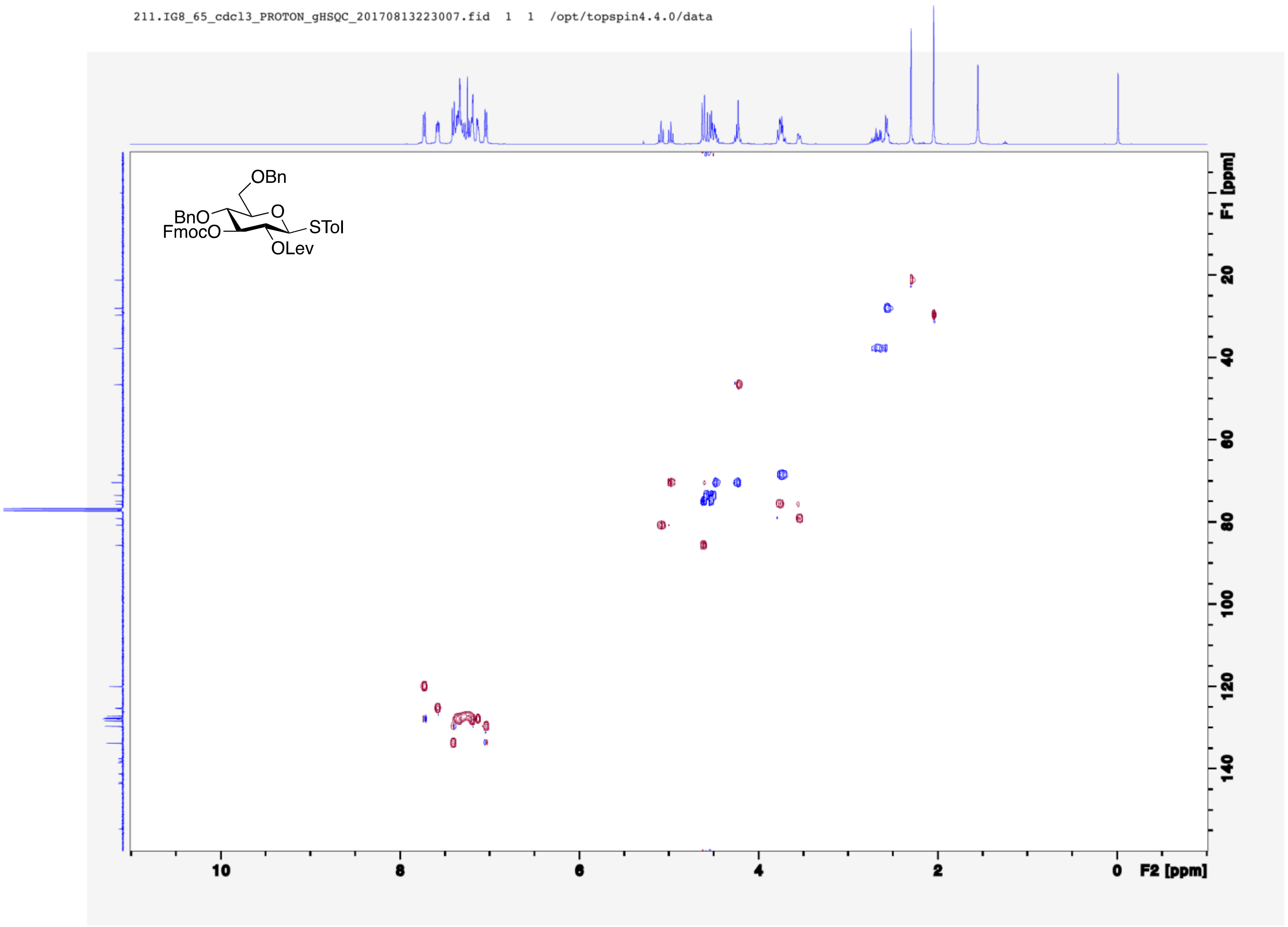
211.IG8_65_cdcl3_PROTON_gHSQC_20170813223007.fid 1 1 /opt/topspin4.4.0/data
OBn
O
BnO
FmocO
STol
OLev
F1 [ppm]
20
40
60
80
100
120
140
10
8
6
4
2
0
F2 [ppm]

257.IG8_92_cdcl3_PROTON_20170912115303.fid 1 1 /opt/topspin4.4.0/data

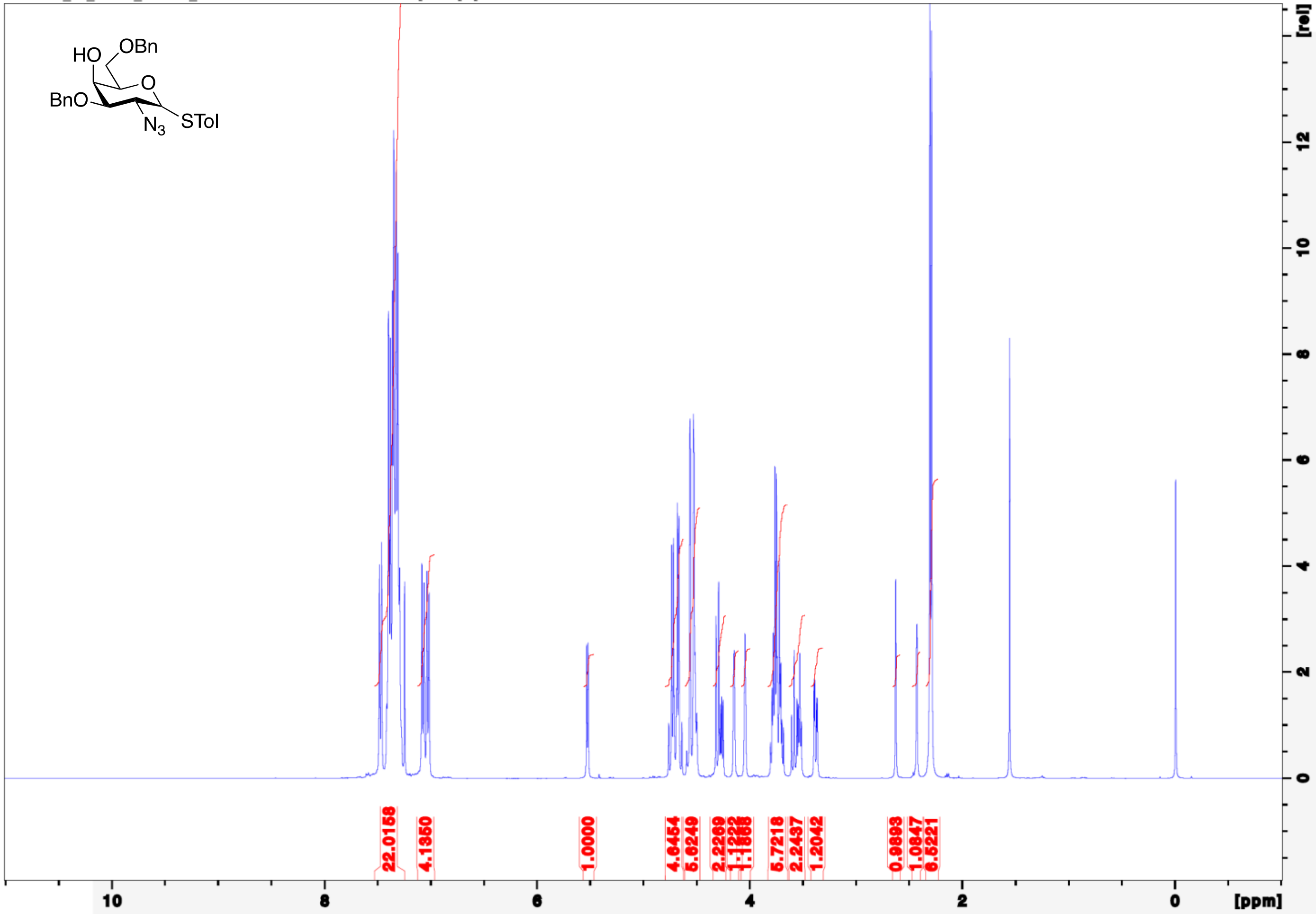

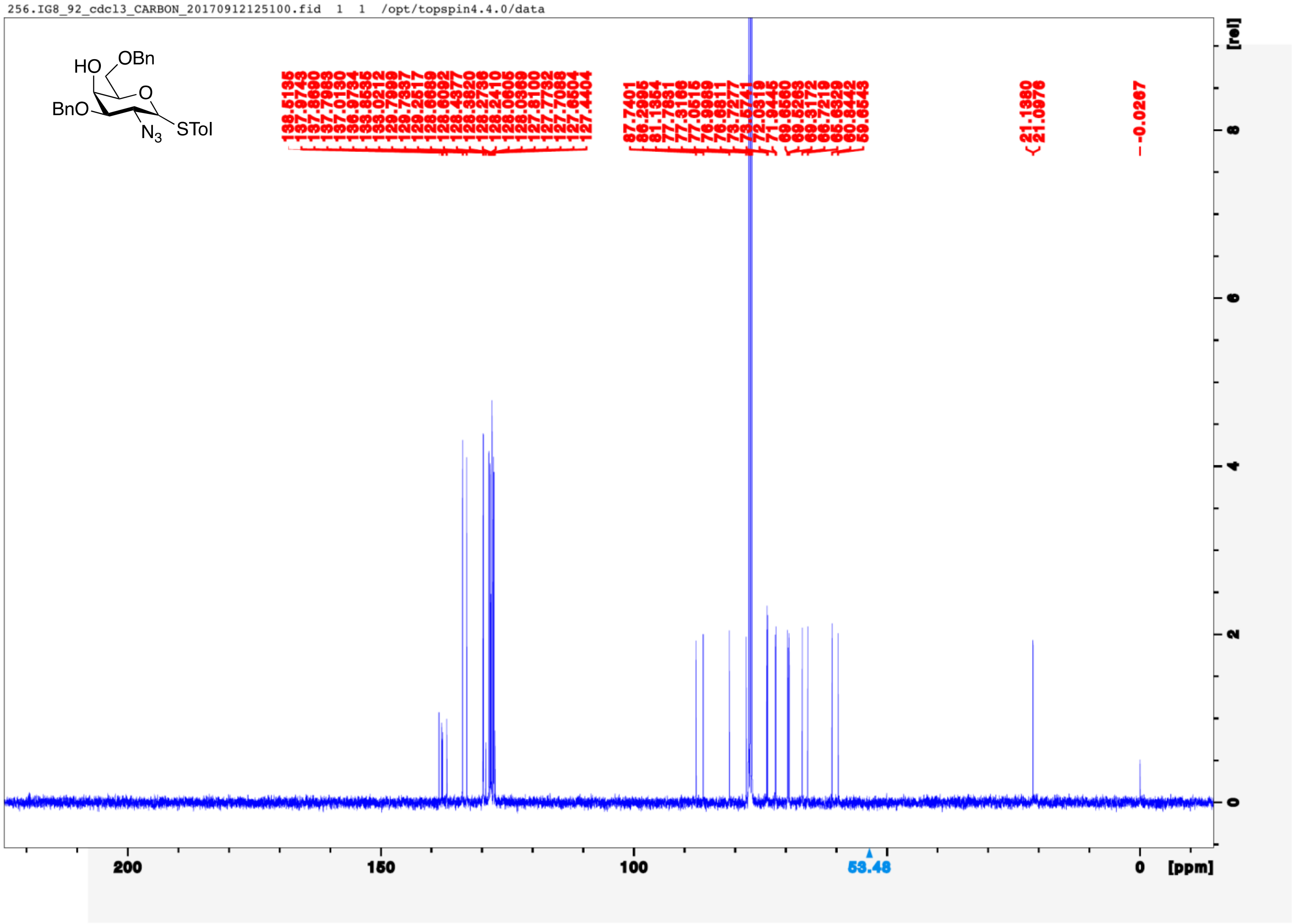
256.IG8_92_cdcl3_CARBON_20170912125100.fid 1 1 /opt/topspin4.4.0/data
HO
OBn
O
BnO
$N_3$
STol
21.1380
21.0976
-0.0267
53.48
200
150
100
0
[ppm]
[rel]
8
6
4
2
0

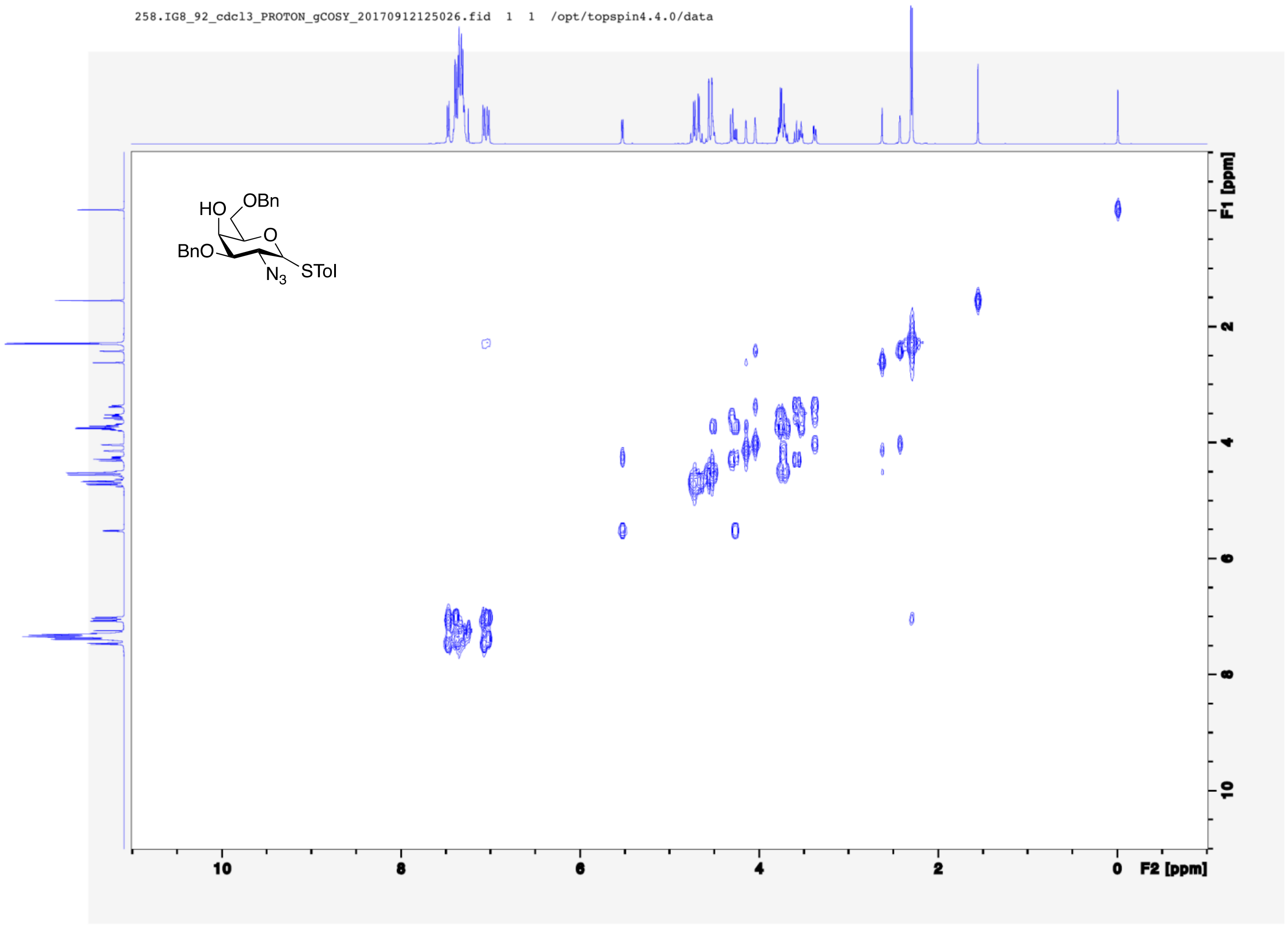
258.IG8_92_cdcl3_PROTON_gCOSY_20170912125026.fid 1 1 /opt/topspin4.4.0/data
HO
OBn
O
BnO
N3
STol
F1 [ppm]
F2 [ppm]
2
4
6
8
10
0

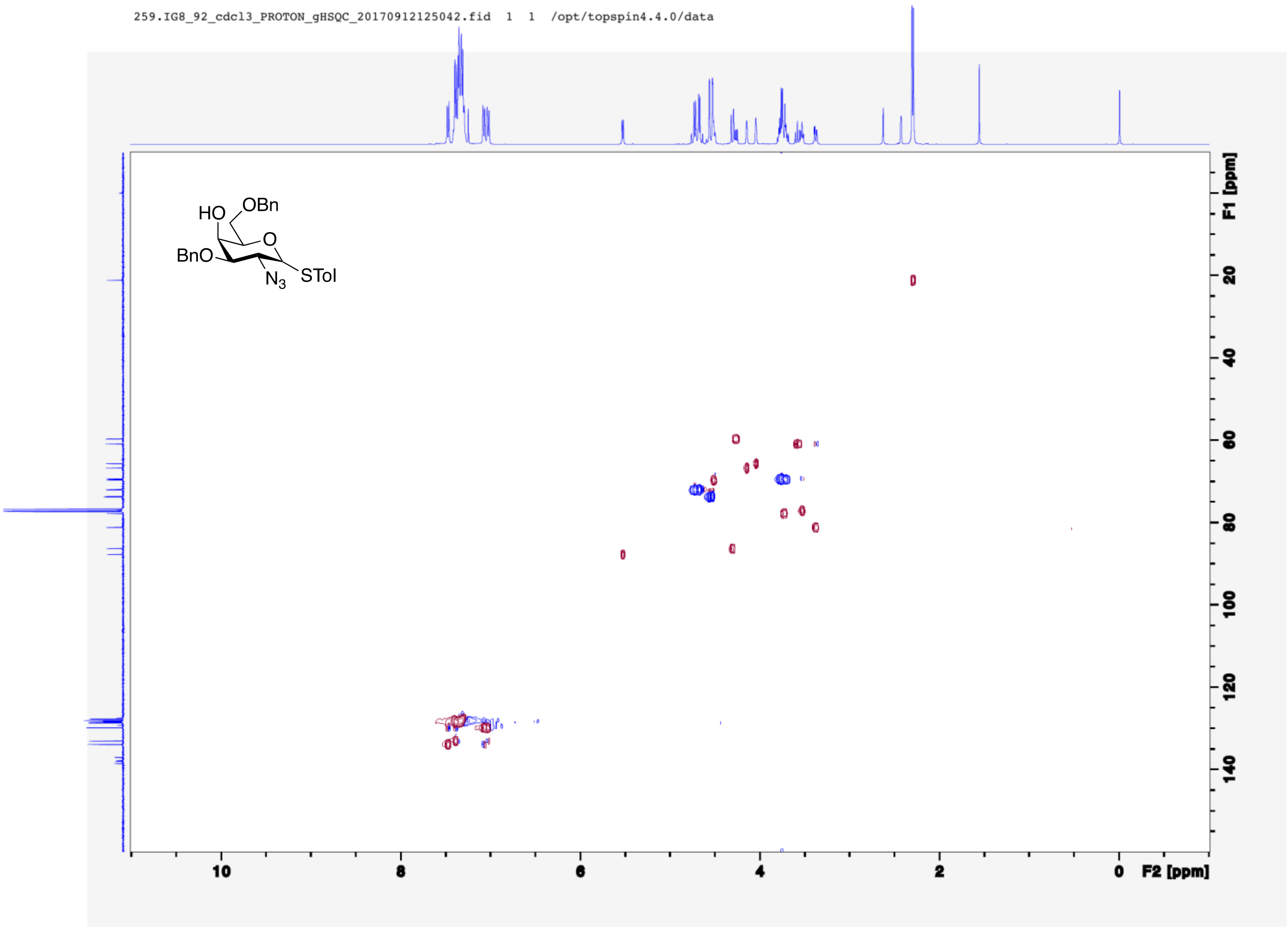
259.IG8_92_cdcl3_PROTON_gHSQC_20170912125042.fid 1 1 /opt/topspin4.4.0/data
HO
OBn
O
BnO
$N_3$
STol
F1 [ppm]
F2 [ppm]

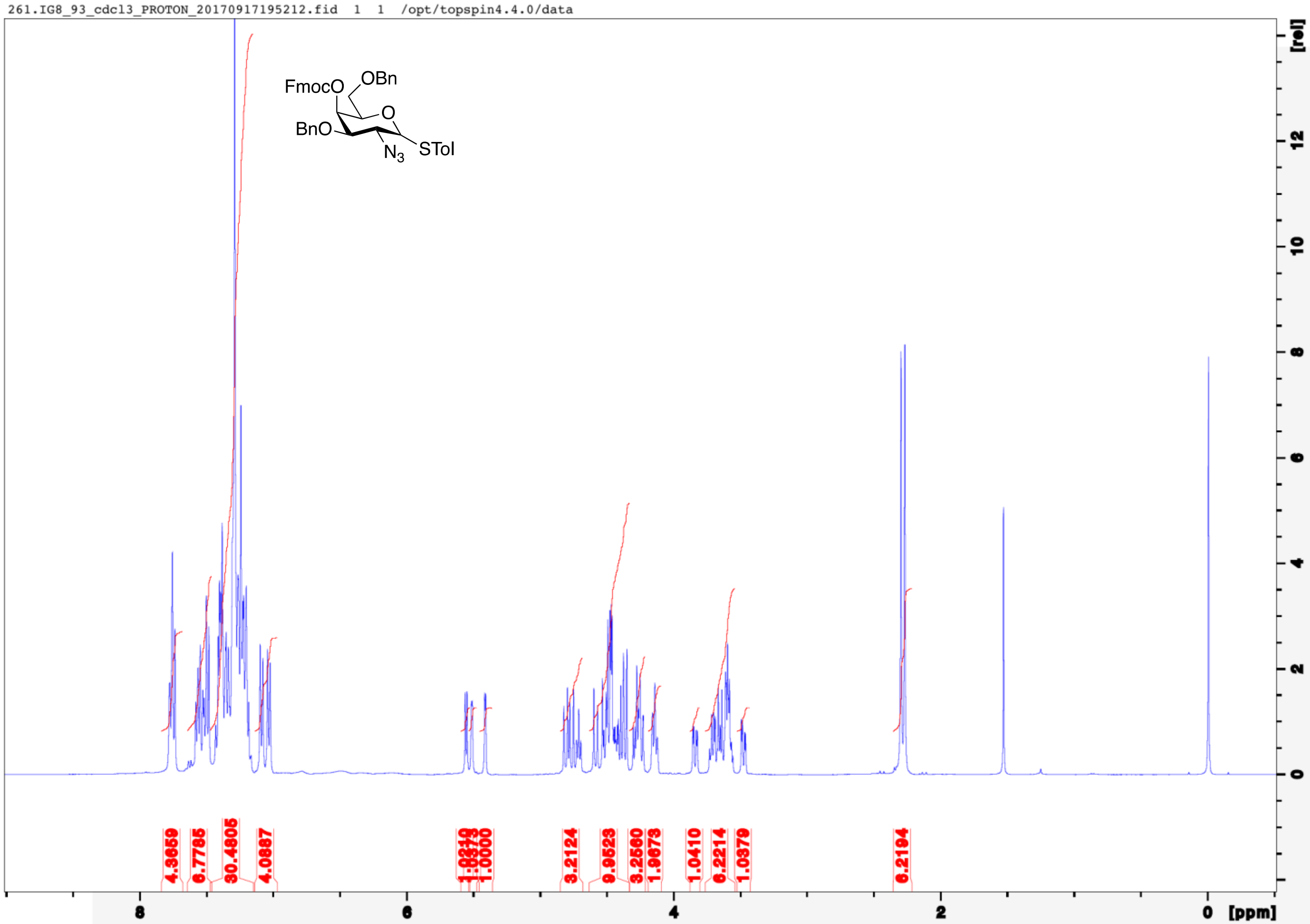
261.IG8_93_cdcl3_PROTON_20170917195212.fid 1 1 /opt/topspin4.4.0/data
FmocO
OBn
O
BnO
$N_3$
STol
[rel]
12
10
8
6
4
2
0
8
6
4
2
0
[ppm]

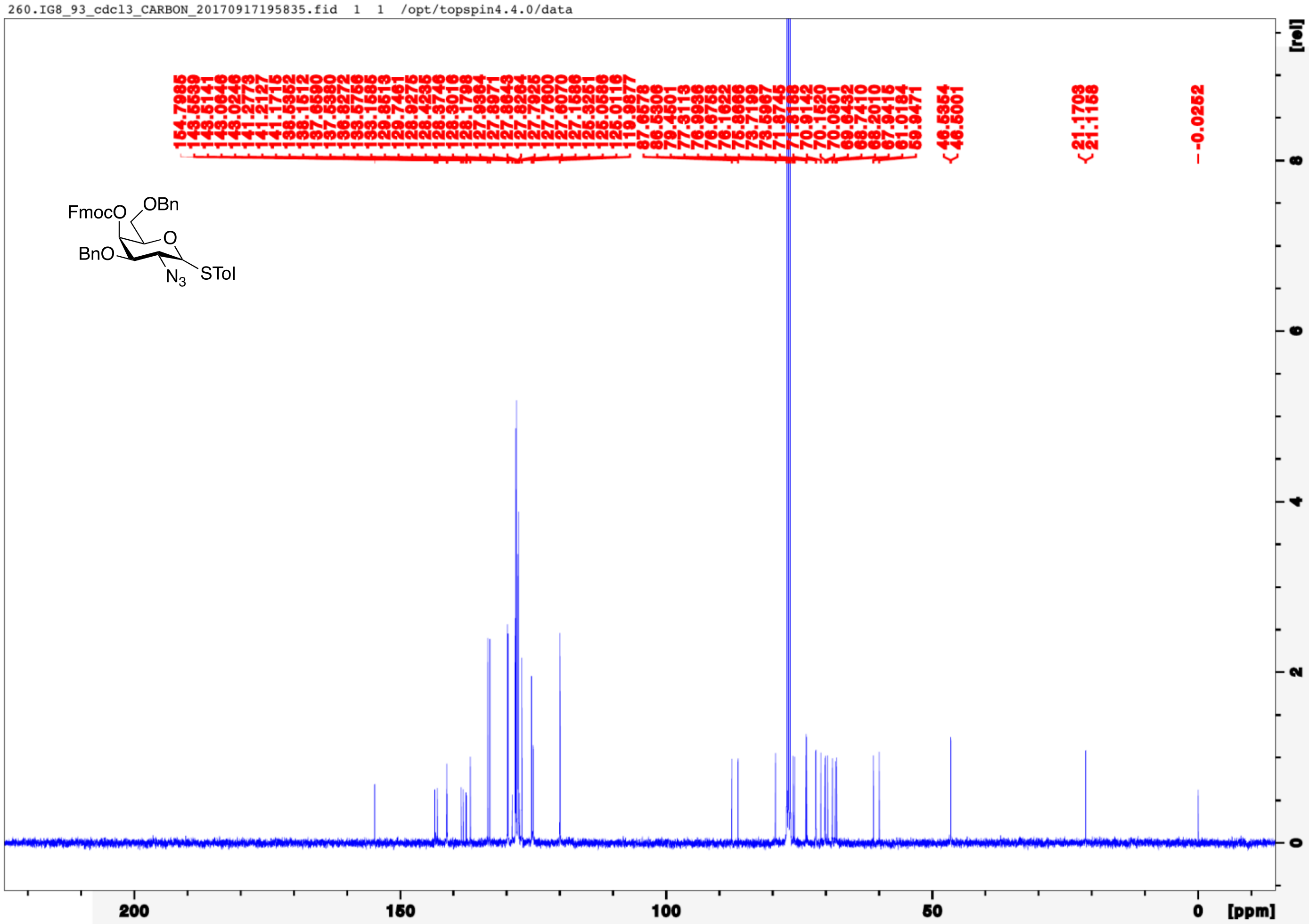
260.IG8_93_cdcl3_CARBON_20170917195835.fid 1 1 /opt/topspin4.4.0/data
FmocO
OBn
O
BnO
N3
STol
154.7985
143.5539
143.5141
143.0646
143.0246
141.2773
141.2127
141.1715
138.5352
138.1512
137.6590
137.5380
136.8272
133.5756
133.1565
129.8513
129.7461
128.9275
128.4235
128.3746
128.3016
128.1798
127.9364
127.8971
127.8643
127.8264
127.7925
127.7600
127.6070
127.1586
125.3251
125.0586
125.0116
119.9877
87.6578
86.5306
77.4501
77.3113
76.9936
76.6758
76.1622
75.8666
75.3799
73.7199
73.5967
71.8745
71.8188
70.9142
70.1520
70.0801
69.6452
68.7410
68.2010
67.9416
61.0184
59.9471
46.5354
46.5001
21.1703
21.1158
-0.0252
[rel]
8
6
4
2
0
200
150
100
50
0
[ppm]

262.IG8_93_cdcl3_PROTON_gCOSY_20170917195809.fid 1 1 /opt/topspin4.4.0/data

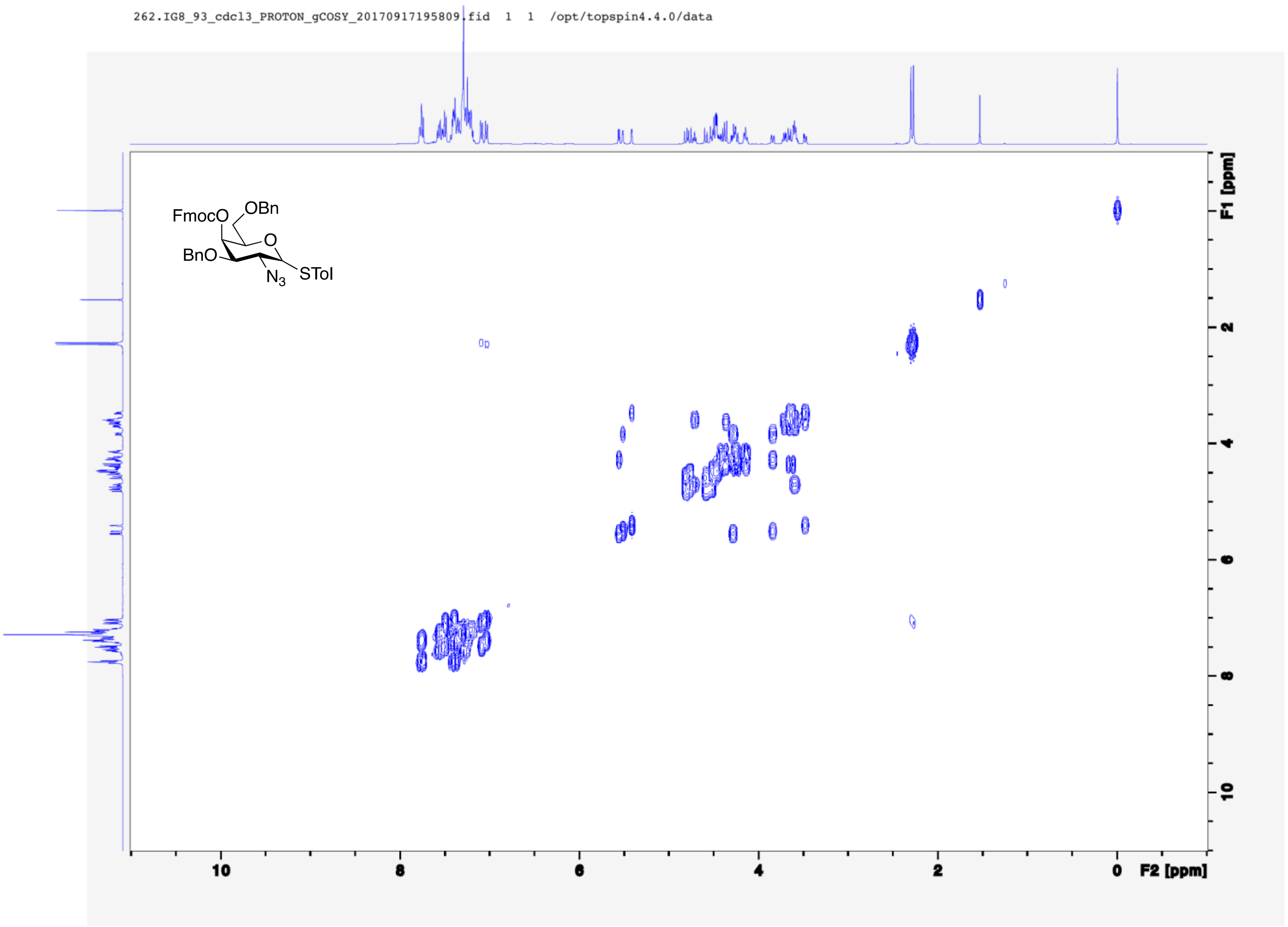

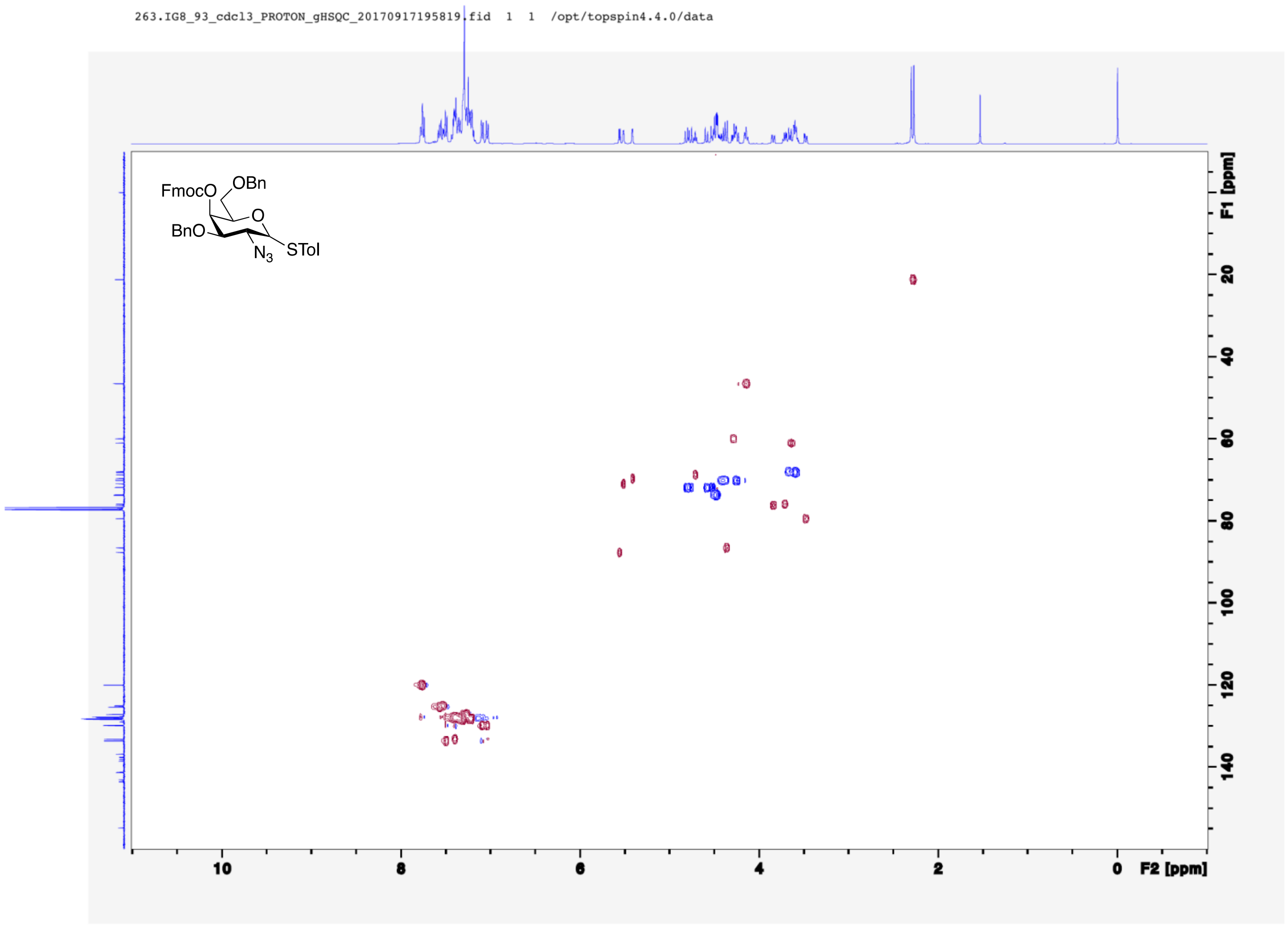
263.IG8_93_cdcl3_PROTON_gHSQC_20170917195819.fid 1 1 /opt/topspin4.4.0/data
FmocO
OBn
O
BnO
$N_3$
STol
F1 [ppm]
F2 [ppm]

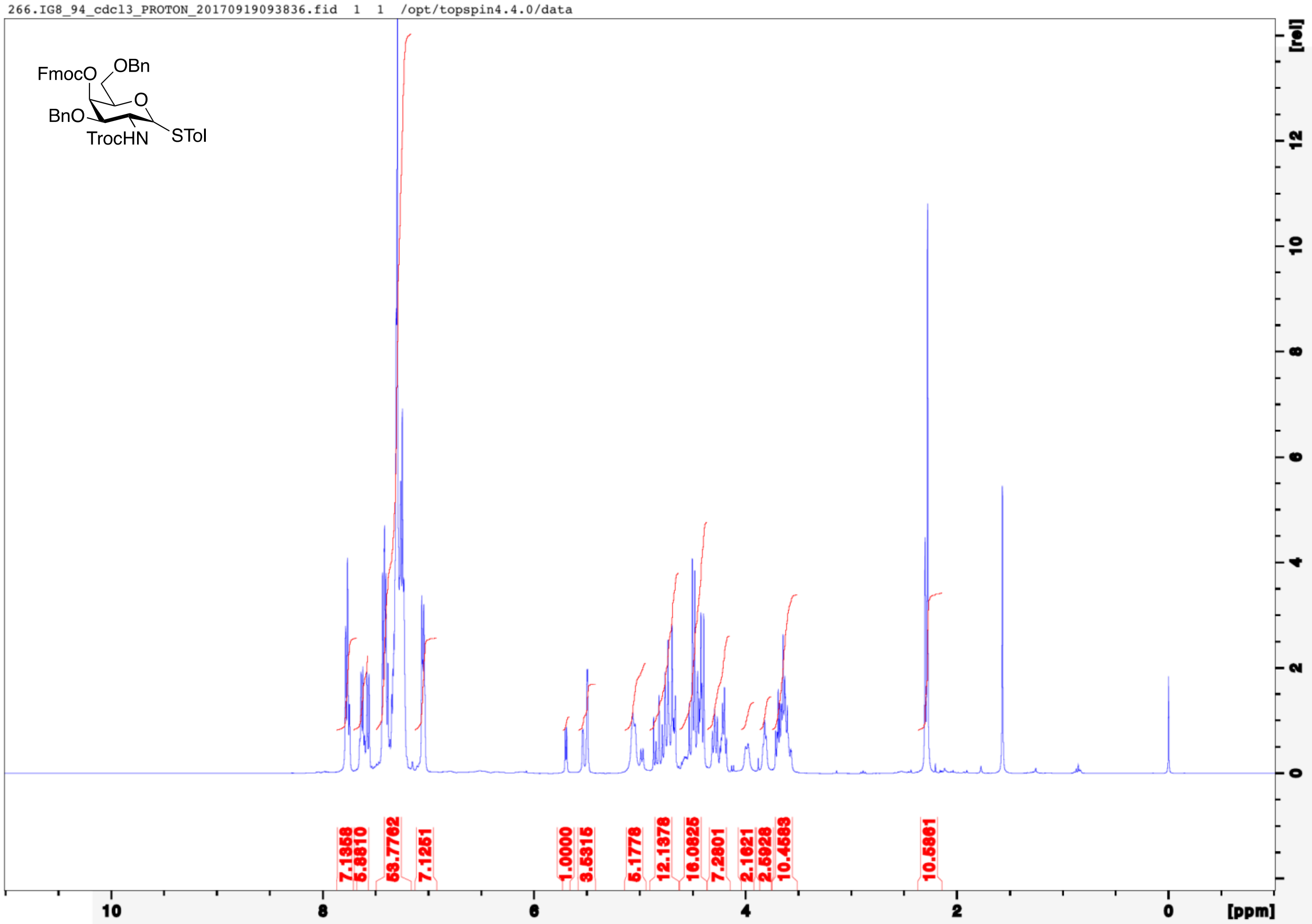

266.IG8_94_cdcl3_PROTON_20170919093836.fid 1 1 /opt/topspin4.4.0/data
FmocO
OBn
O
BnO
TrocHN
STol
7.1358
5.8810
53.7762
7.1251
1.0000
3.5315
5.1778
12.1376
16.0825
7.2801
2.1621
2.5928
10.4583
10.5861
10
8
6
4
2
0
[ppm]
[rel]
12
10
8
6
4
2
0

264.IG8_94_cdcl3_CARBON_20170919095009.fid 1 1 /opt/topspin4.4.0/data

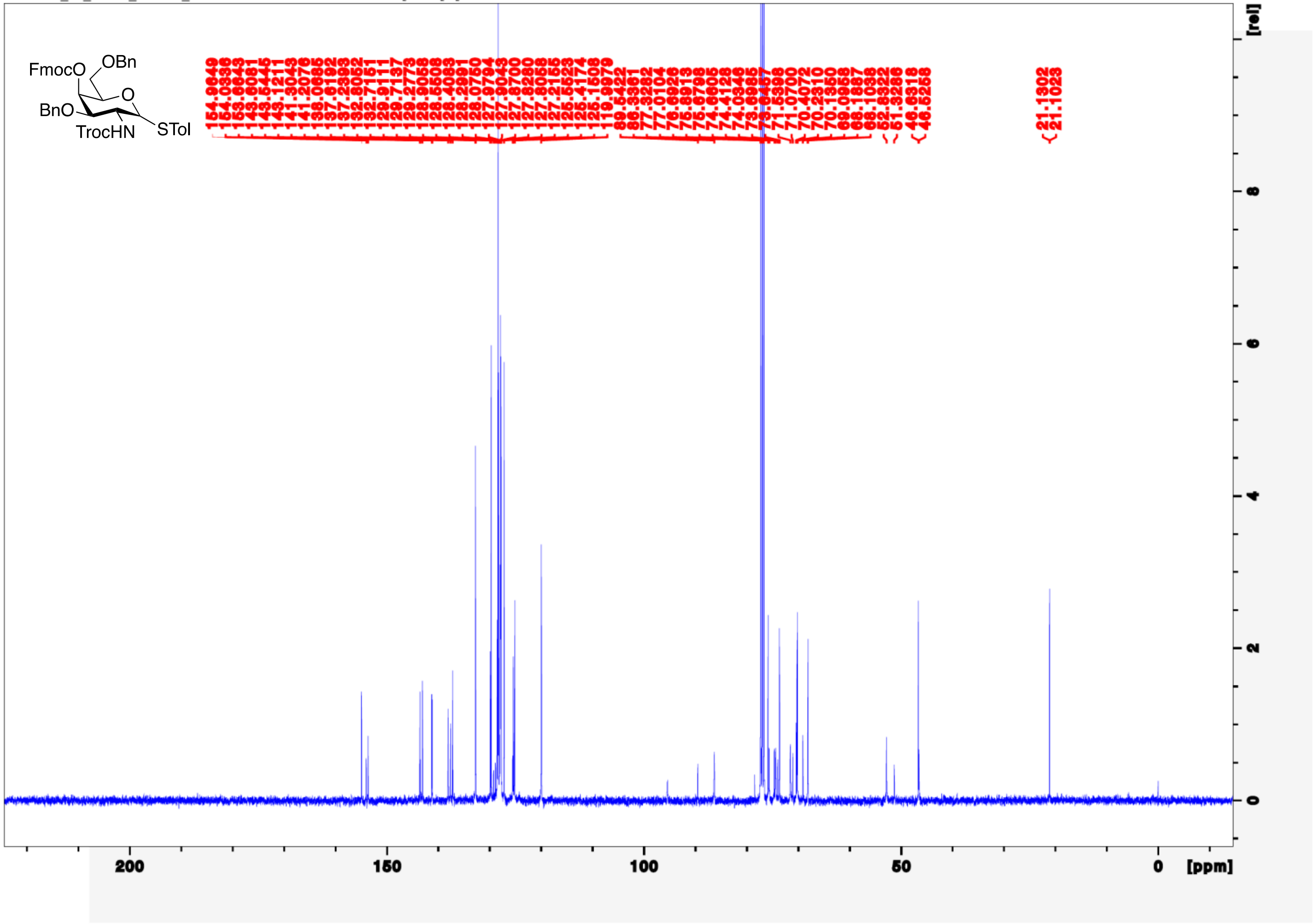

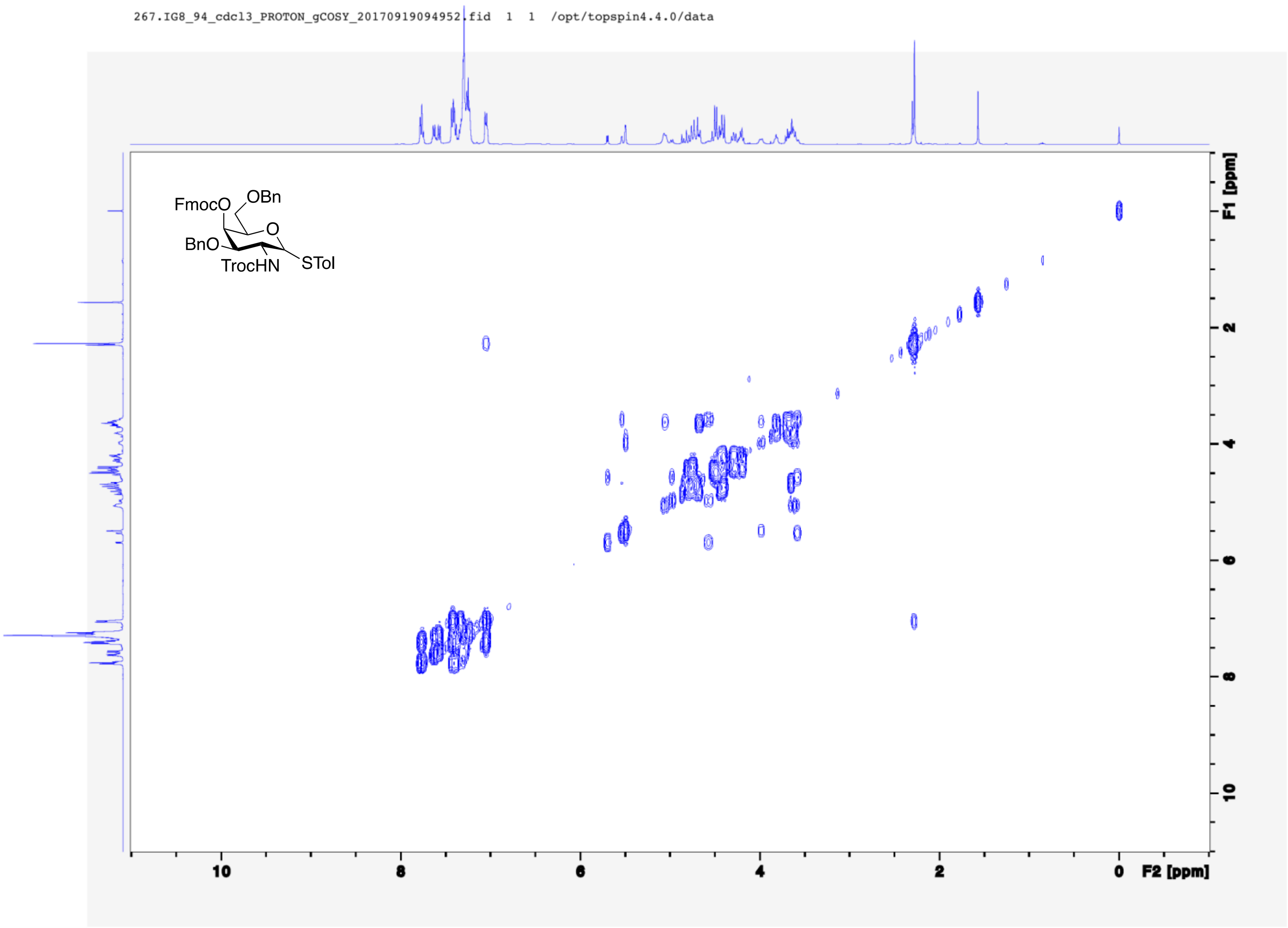

267.IG8_94_cdcl3_PROTON_gCOSY_20170919094952.fid 1 1 /opt/topspin4.4.0/data
FmocO
OBn
O
BnO
TrocHN
STol
F1 [ppm]
F2 [ppm]

268.IG8_94_cdcl3_PROTON_gHSQC_20170919094959.fid 1 1 /opt/topspin4.4.0/data

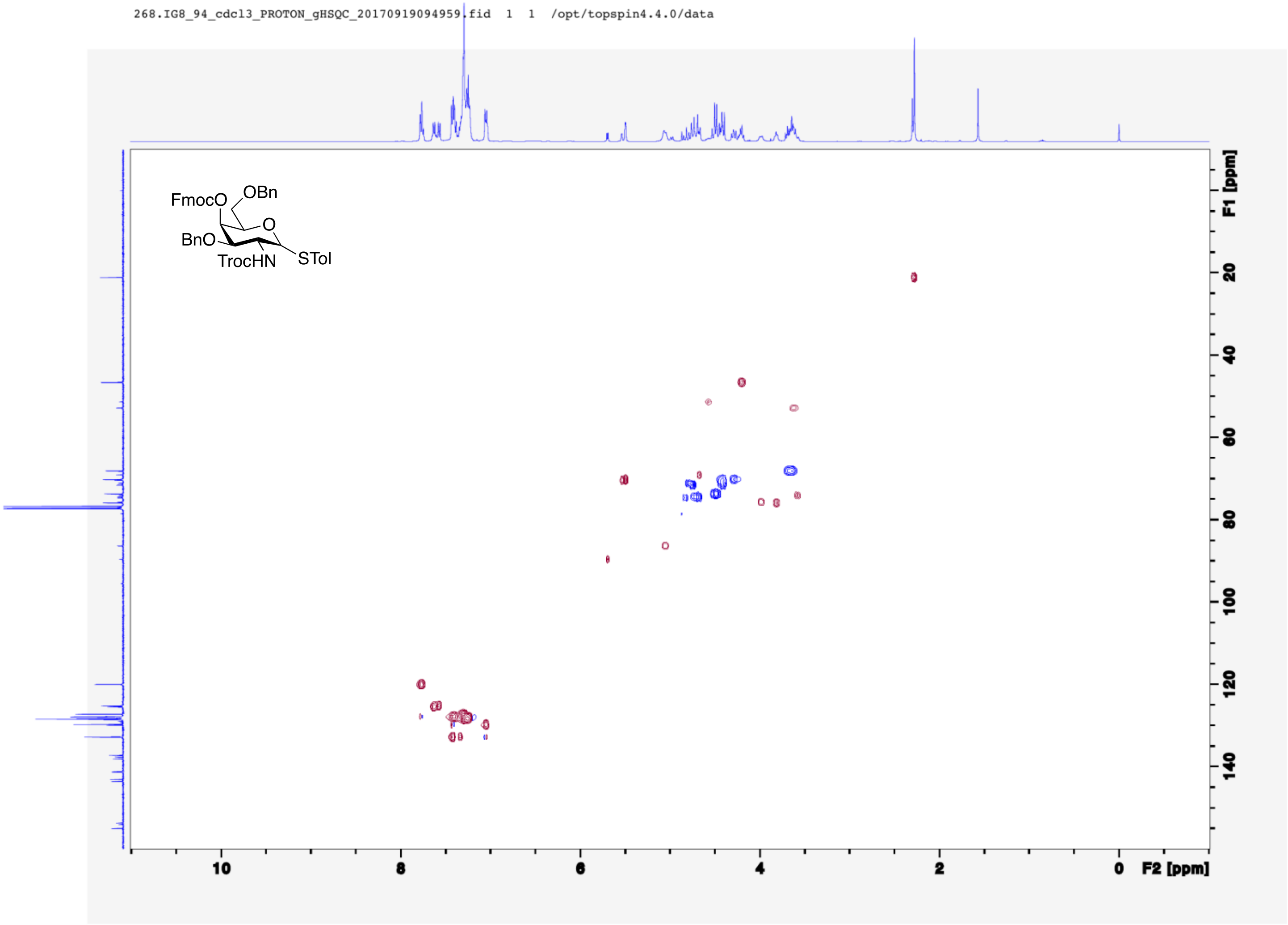

191.IG8_57_cdcl3_PROTON_20170624190225.fid 1 1 /opt/topspin4.4.0/data

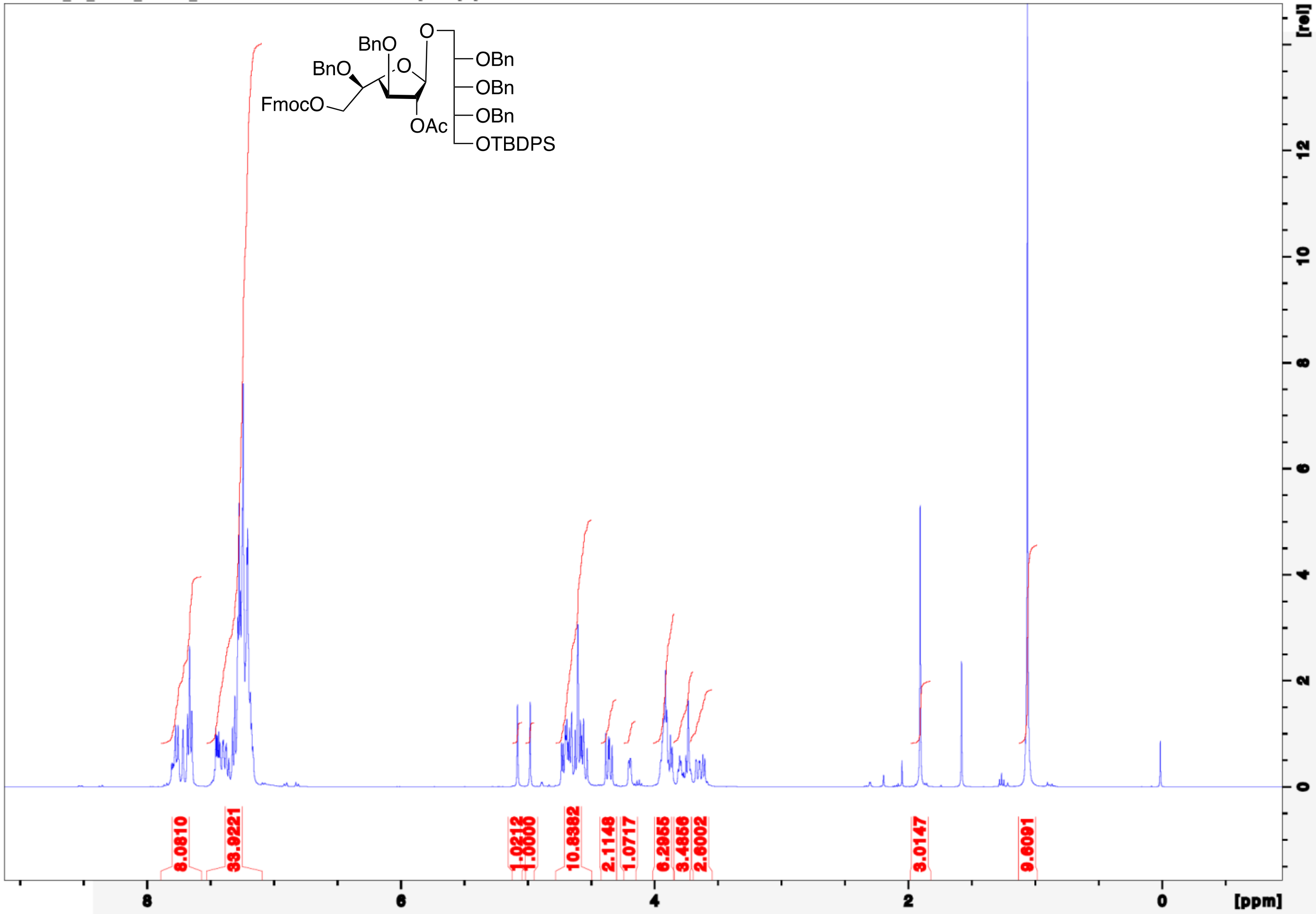

190.IG8_57_cdcl3_CARBON_20170624201520.fid 1 1 /opt/topspin4.4.0/data

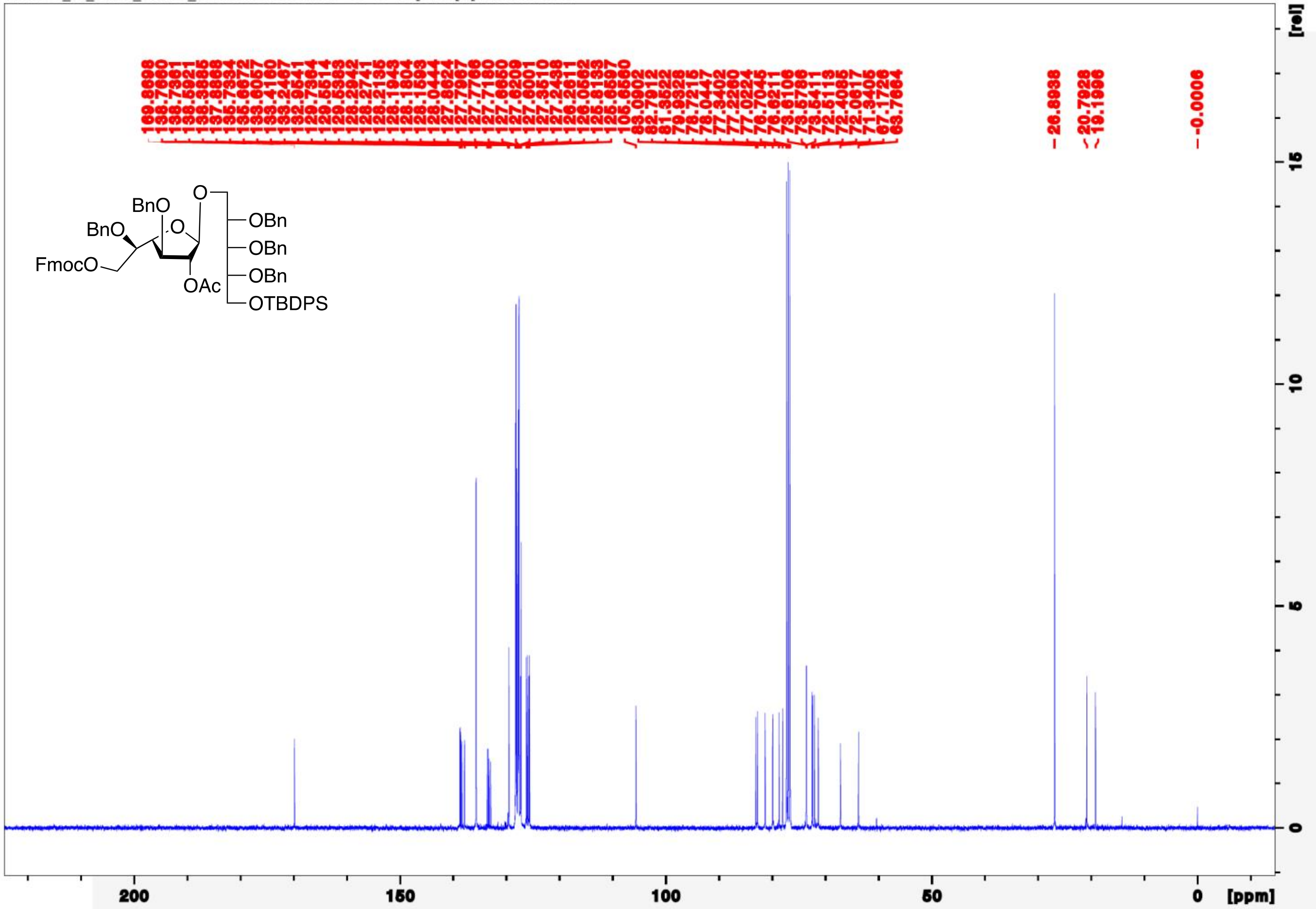

192.IG8_57_cdcl3_PROTON_COSY_20170624201451.fid 1 1 /opt/topspin4.4.0/data

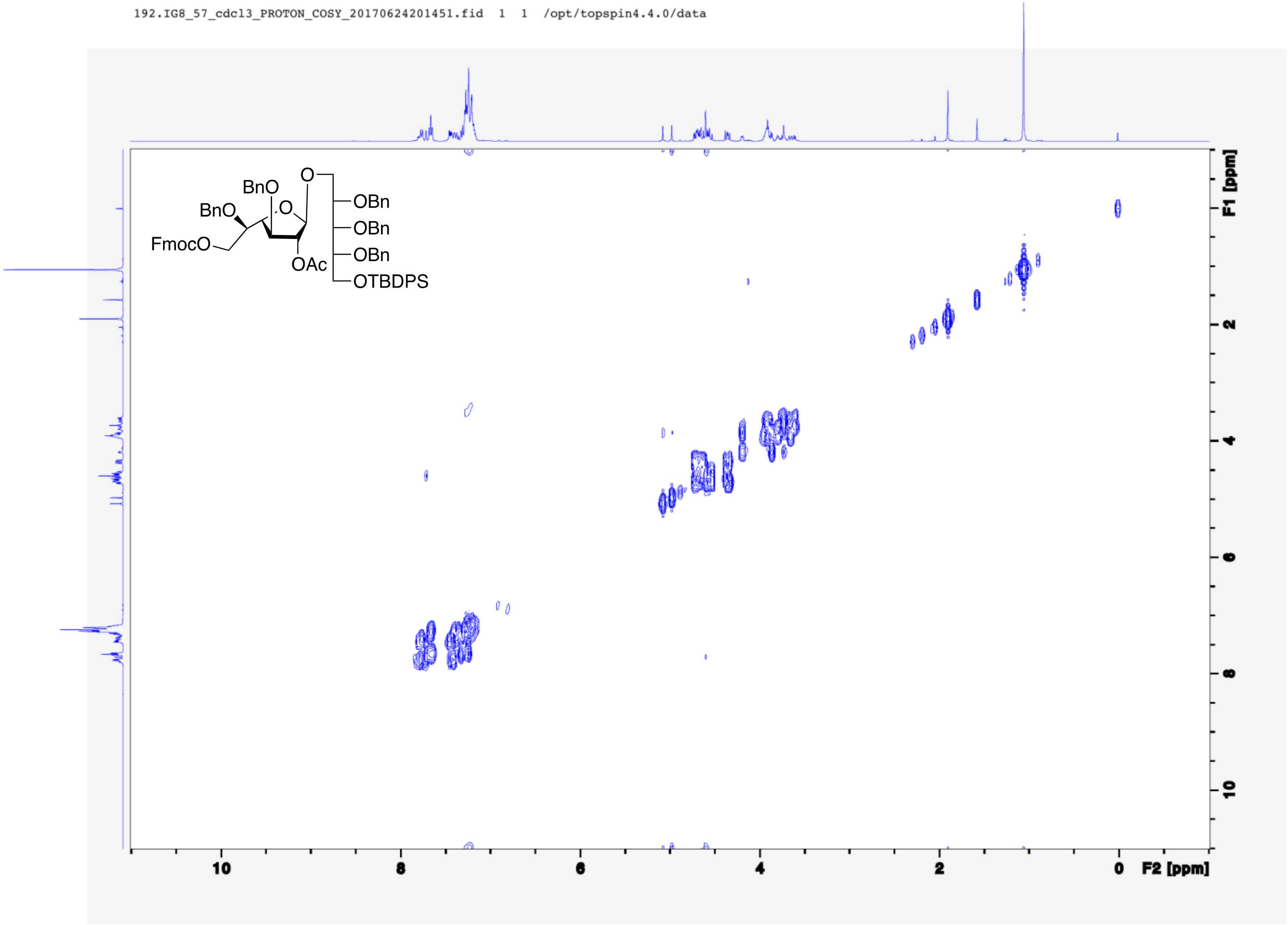

193.IG8_57_cdcl3_PROTON_gHSQC_20170624201503.fid 1 1 /opt/topspin4.4.0/data

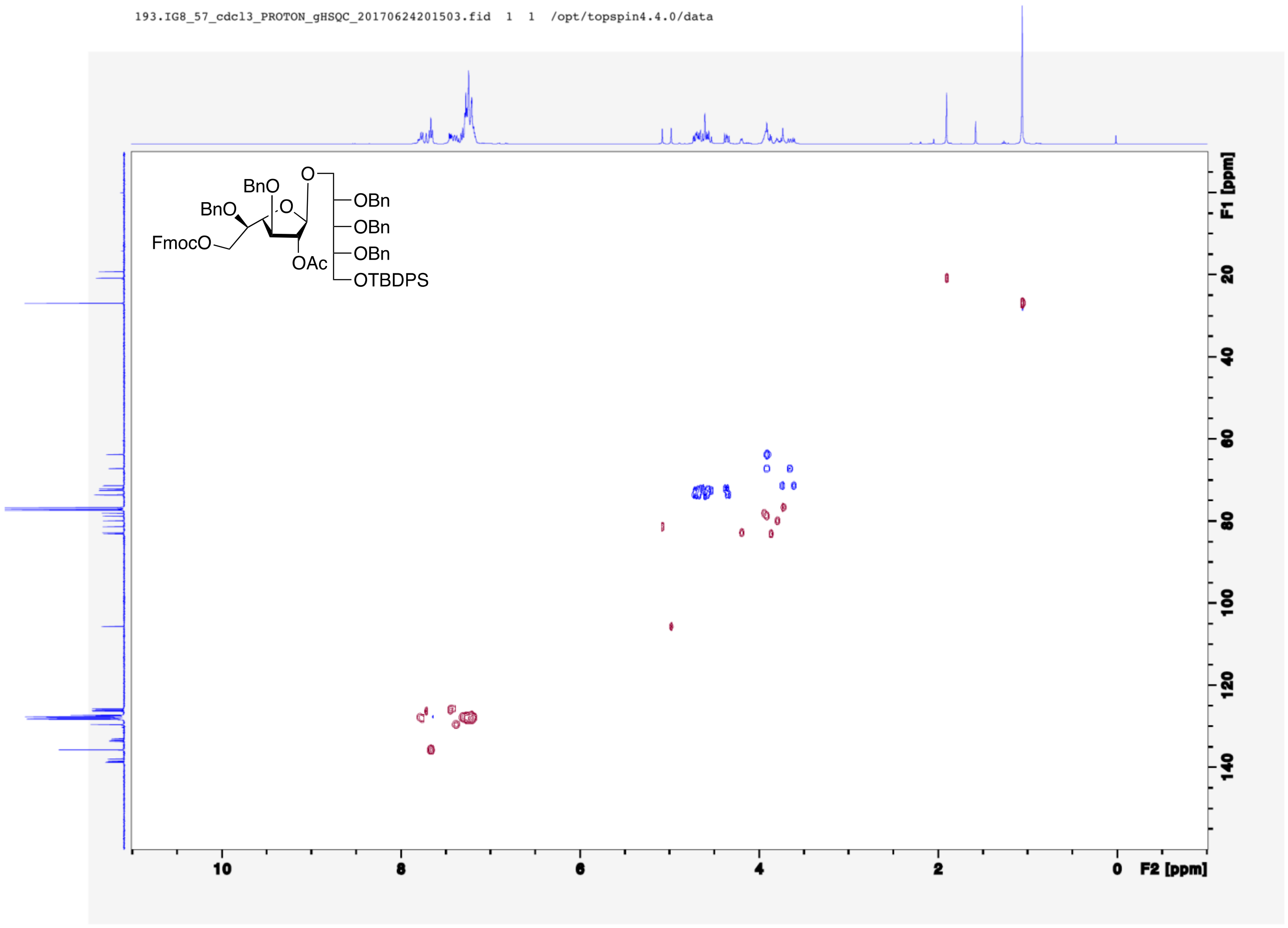

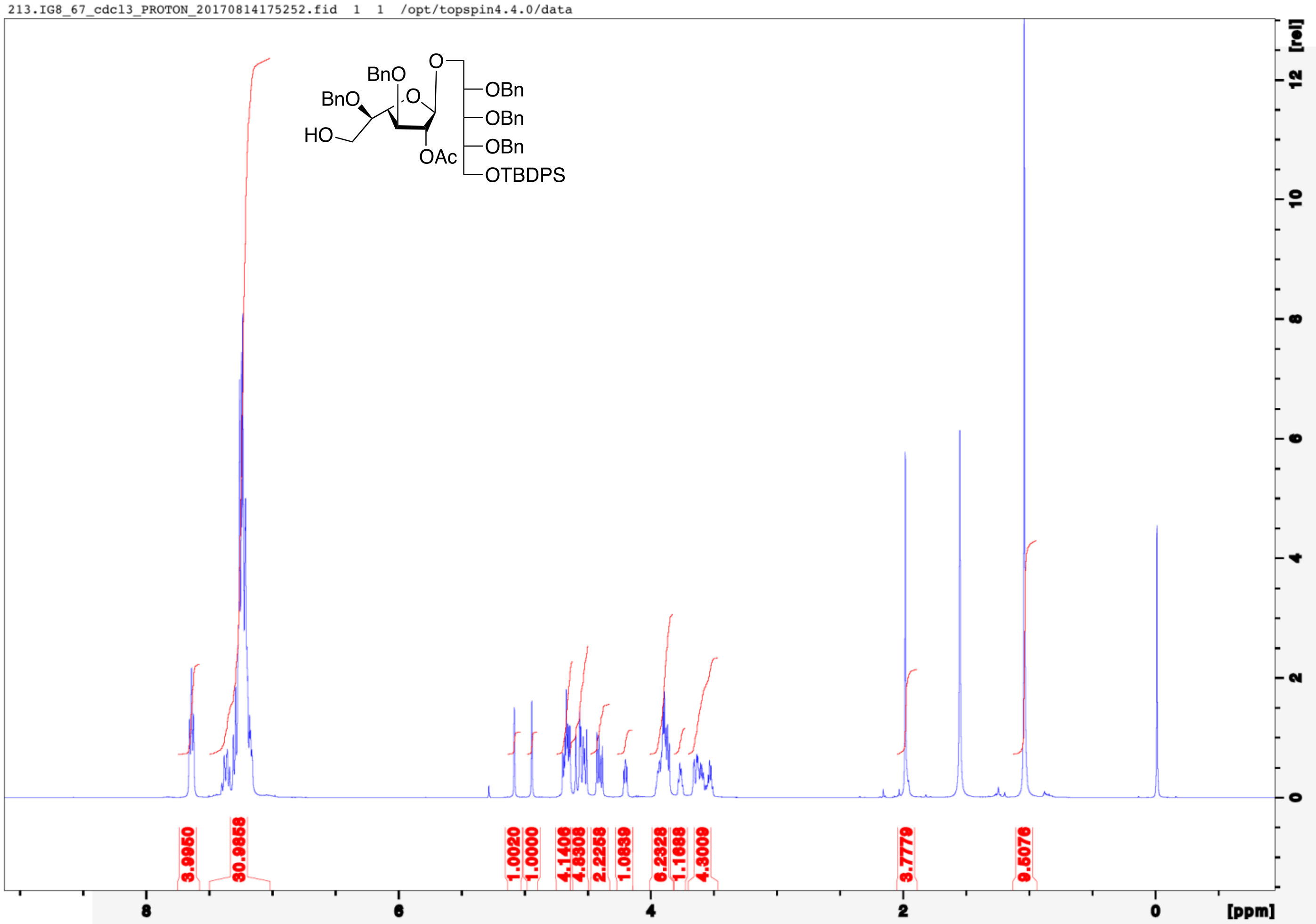
213.IG8_67_cdcl3_PROTON_20170814175252.fid 1 1 /opt/topspin4.4.0/data
BnO
BnO
HO
O
O
OAc
OBn
OBn
OBn
OTBDPS
3.9950
30.9858
1.0020
1.0000
4.1406
4.8308
2.2258
1.0839
6.2328
1.1688
4.3009
3.7779
9.5076
[rel]
[ppm]

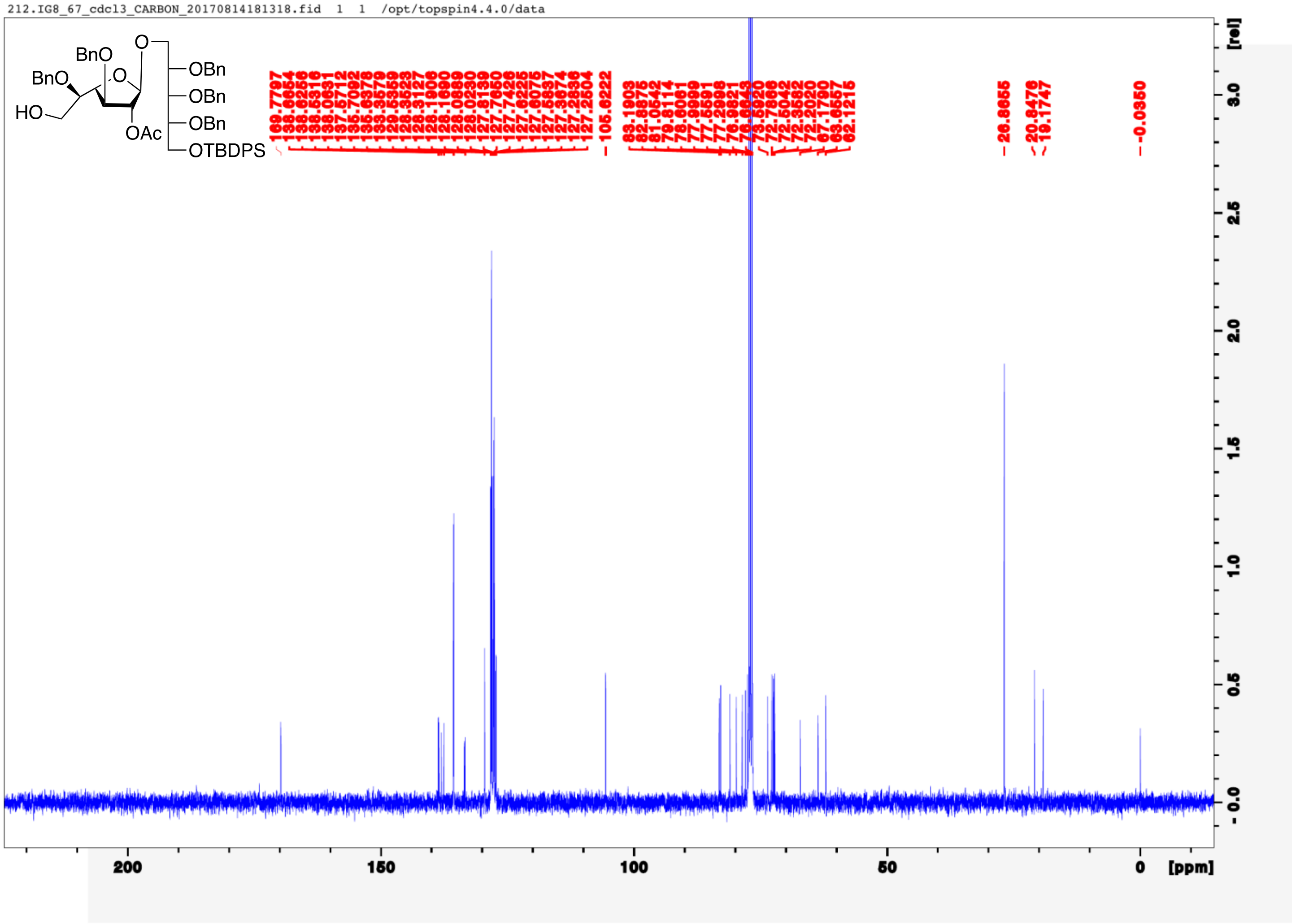
212.IG8_67_cdcl3_CARBON_20170814181318.fid 1 1 /opt/topspin4.4.0/data
BnO
BnO
HO
O
OAc
O
OBn
OBn
OBn
OTBDPS
[ppm]
[rel]

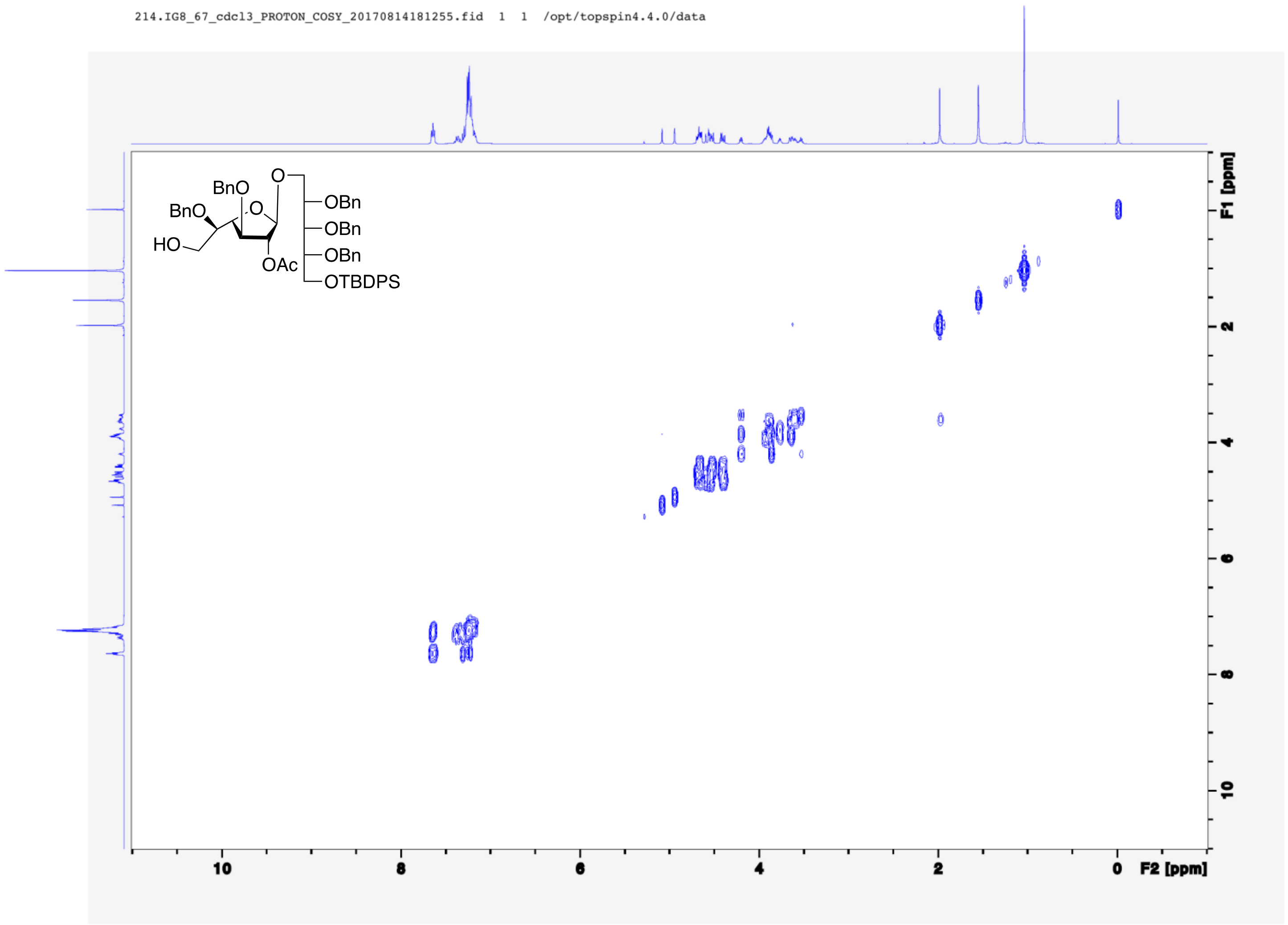
214.IG8_67_cdcl3_PROTON_COSY_20170814181255.fid 1 1 /opt/topspin4.4.0/data
BnO
BnO
HO
O
O
OAc
OBn
OBn
OBn
OTBDPS
F1 [ppm]
F2 [ppm]
10
8
6
4
2
0

215.IG8_67_cdcl3_PROTON_gHSQC_20170814181305.fid 1 1 /opt/topspin4.4.0/data

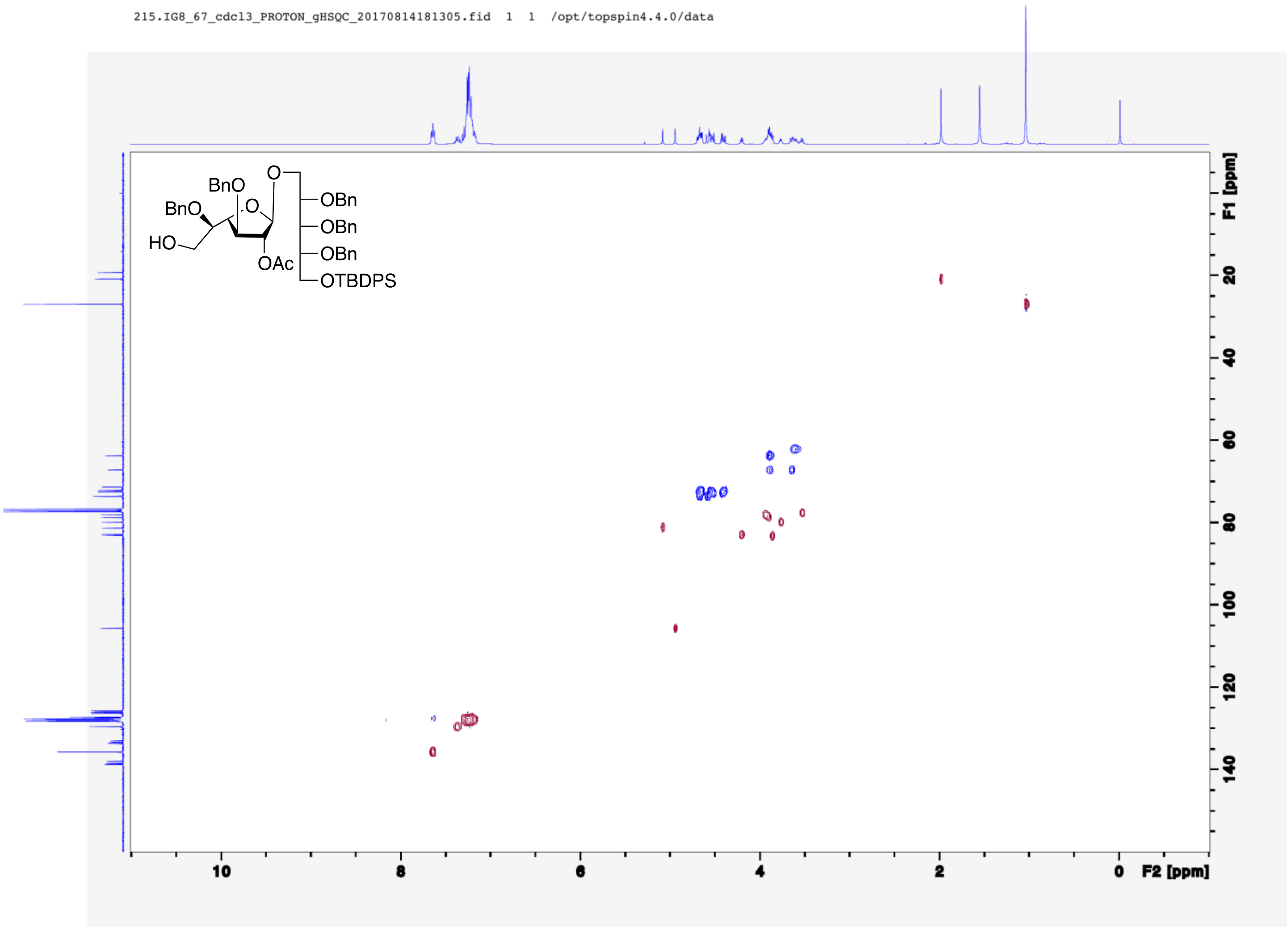

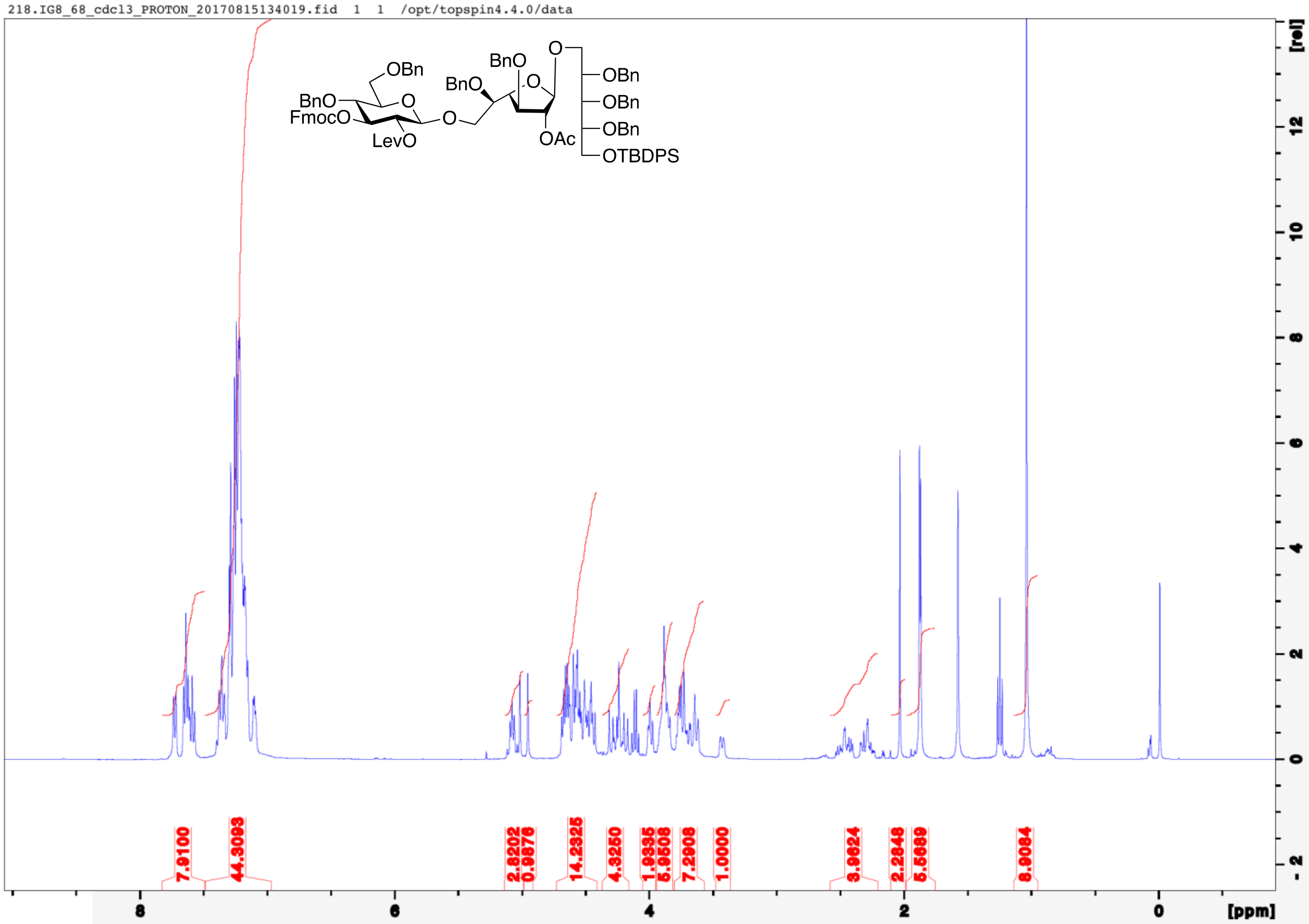
218.IG8_68_cdcl3_PROTON_20170815134019.fid 1 1 /opt/topspin4.4.0/data
OBn
BnO
FmocO
O
LevO
O
BnO
BnO
O
O
OAc
OBn
OBn
OBn
OTBDPS
7.9100
44.3093
2.8202
0.9876
14.2325
4.3250
1.9335
5.9508
7.2908
1.0000
3.9624
2.2848
5.5889
8.9084
[ppm]
[rel]

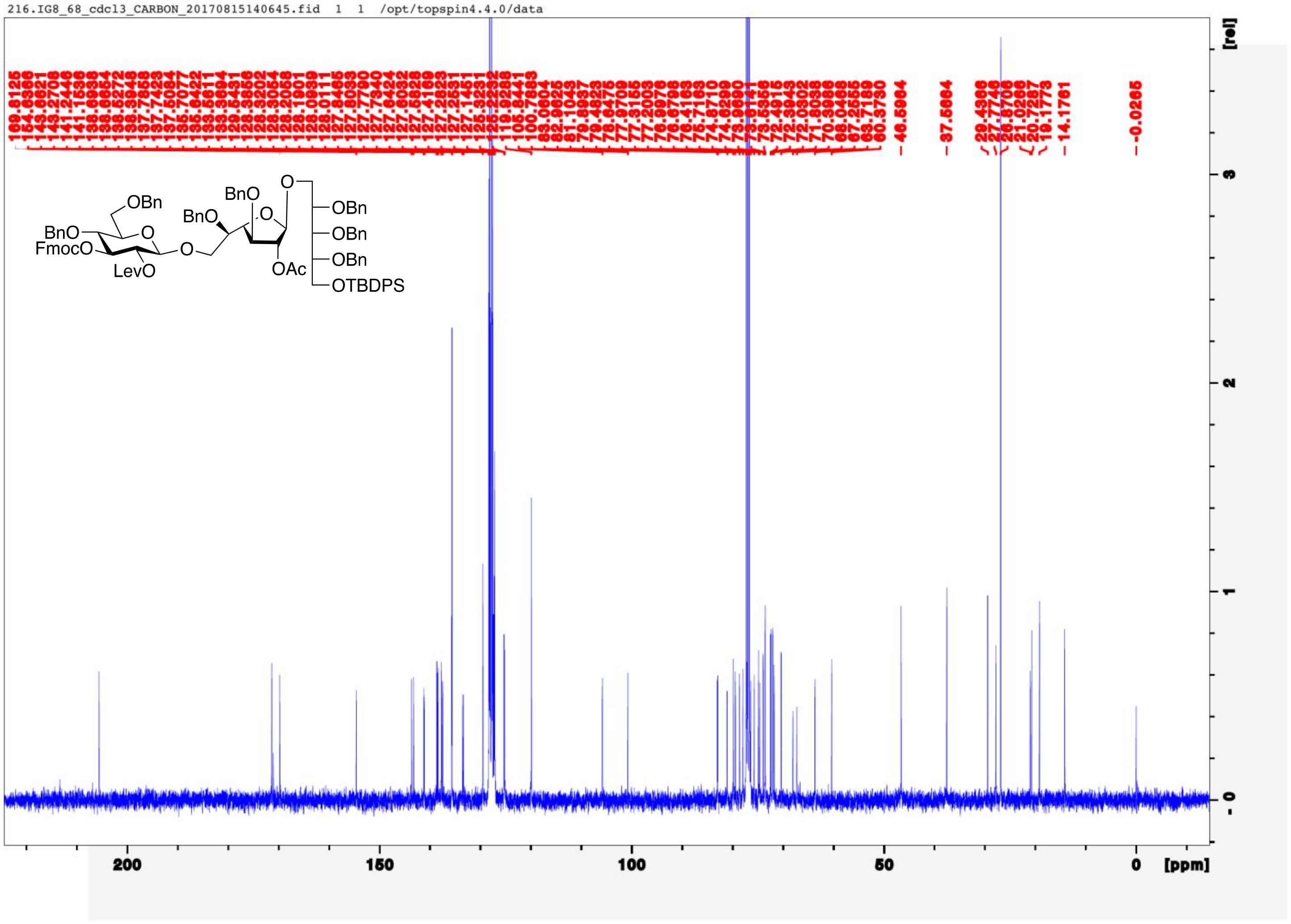
216.IG8_68_cdcl3_CARBON_20170815140645.fid 1 1 /opt/topspin4.4.0/data
OBn
BnO
FmocO
LevO
O
BnO
BnO
O
OAc
OBn
OBn
OBn
OTBDPS
[rel]
200
150
100
50
0
[ppm]

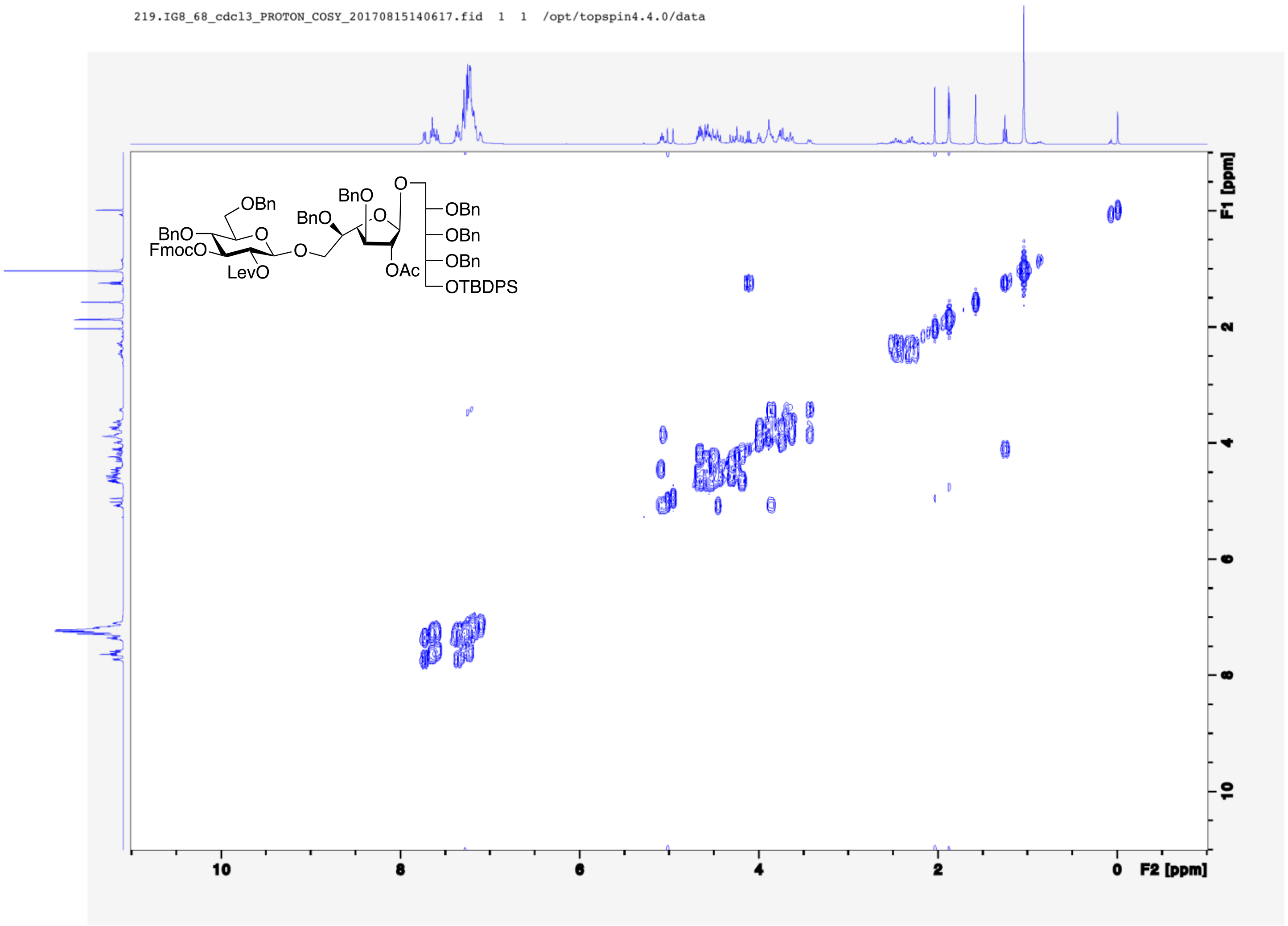
219.IG8_68_cdcl3_PROTON_COSY_20170815140617.fid 1 1 /opt/topspin4.4.0/data
OBn
BnO
FmocO
LevO
O
BnO
BnO
O
OAc
OBn
OBn
OBn
OTBDPS
F1 [ppm]
F2 [ppm]
10
8
6
4
2
0

220.IG8_68_cdcl3_PROTON_gHSQC_20170815140632.fid 1 1 /opt/topspin4.4.0/data

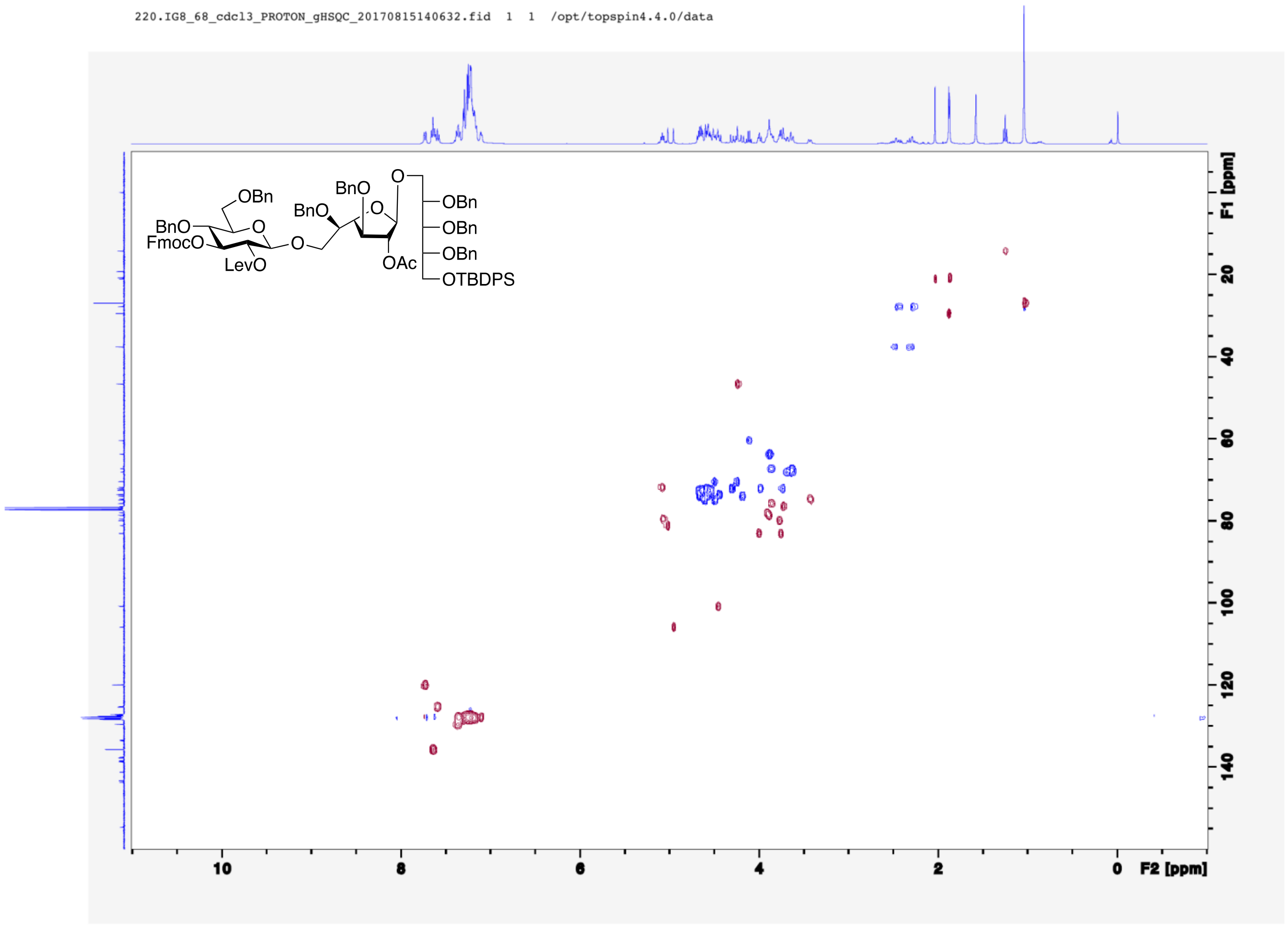

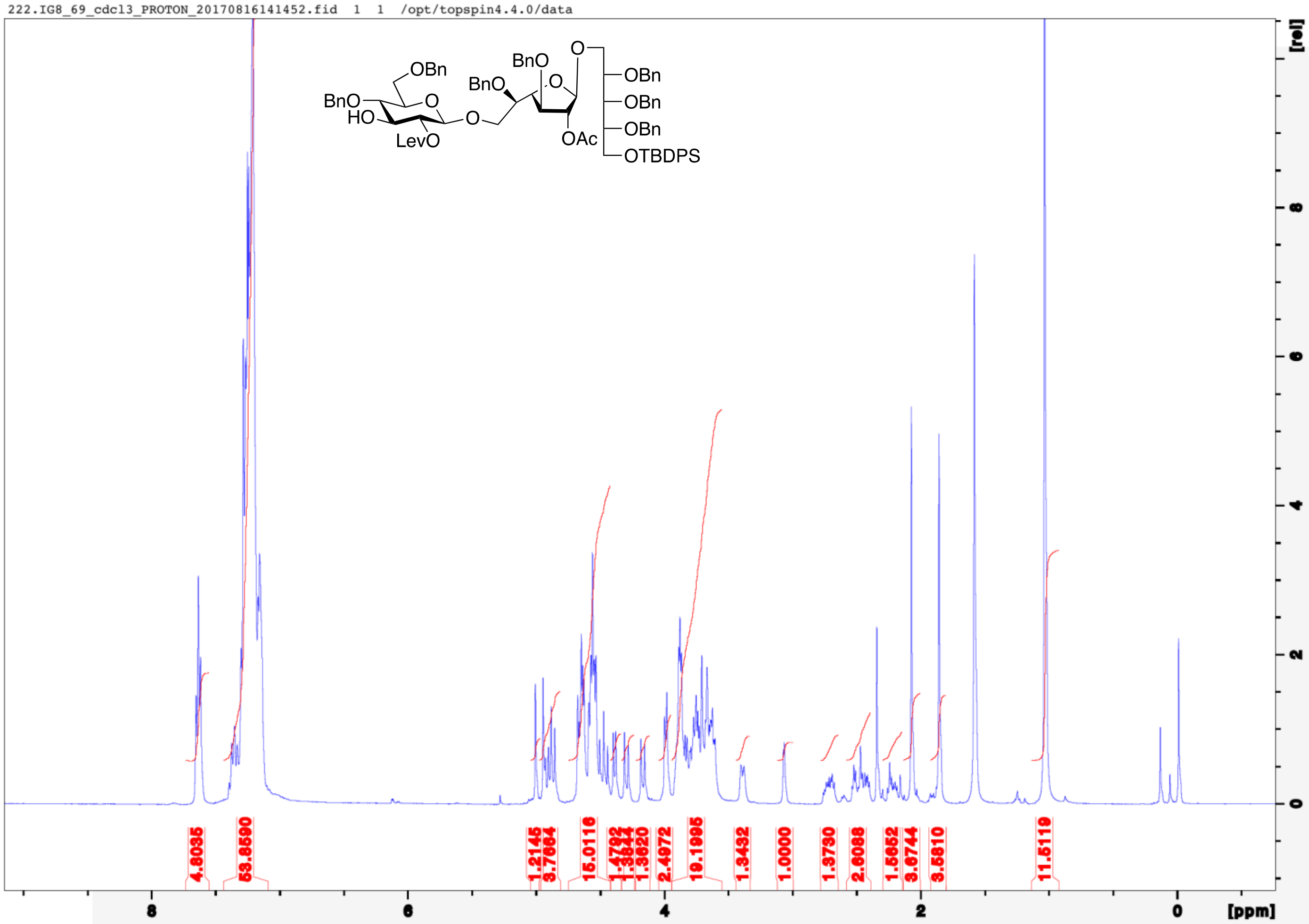
222.IG8_69_cdcl3_PROTON_20170816141452.fid 1 1 /opt/topspin4.4.0/data
OBn
BnO
HO
O
LevO
O
BnO
BnO
O
O
OAc
OBn
OBn
OBn
OTBDPS
4.8035
53.8590
1.2145
3.7684
15.0116
2.4972
19.1995
1.3432
1.0000
1.3730
2.6088
1.5652
3.6744
3.5810
11.5119
8
6
4
2
0
[ppm]
[rel]

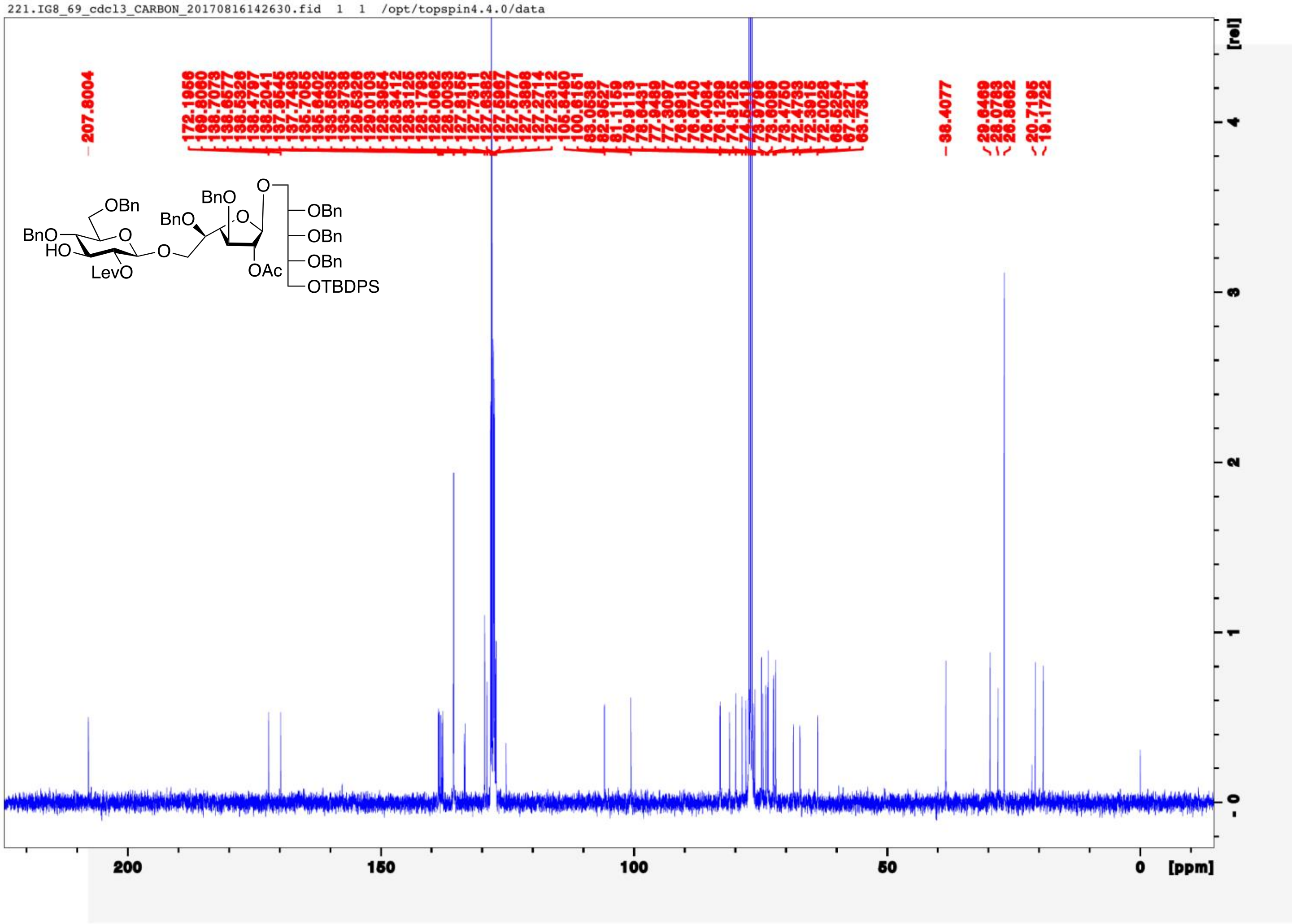
221.IG8_69_cdcl3_CARBON_20170816142630.fid 1 1 /opt/topspin4.4.0/data
BnO
HO
OBn
O
LevO
O
BnO
BnO
O
OAc
O
OBn
OBn
OBn
OTBDPS
200
150
100
50
0
[ppm]
[rel]

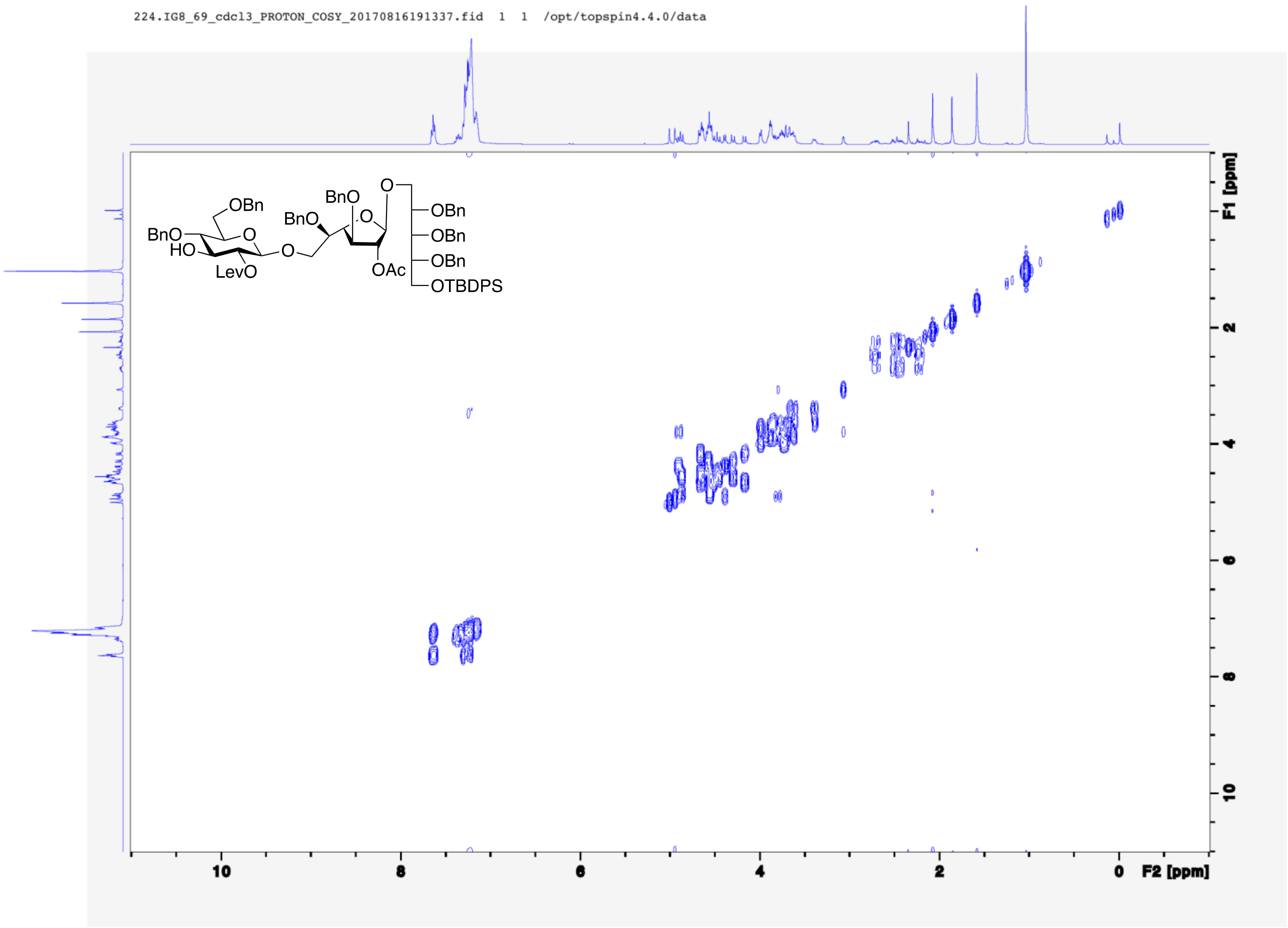
224.IG8_69_cdcl3_PROTON_COSY_20170816191337.fid 1 1 /opt/topspin4.4.0/data
OBn
BnO
HO
O
LevO
BnO
BnO
O
O
OAc
OBn
OBn
OBn
OTBDPS
F1 [ppm]
2
4
6
8
10
F2 [ppm]
10
8
6
4
2
0

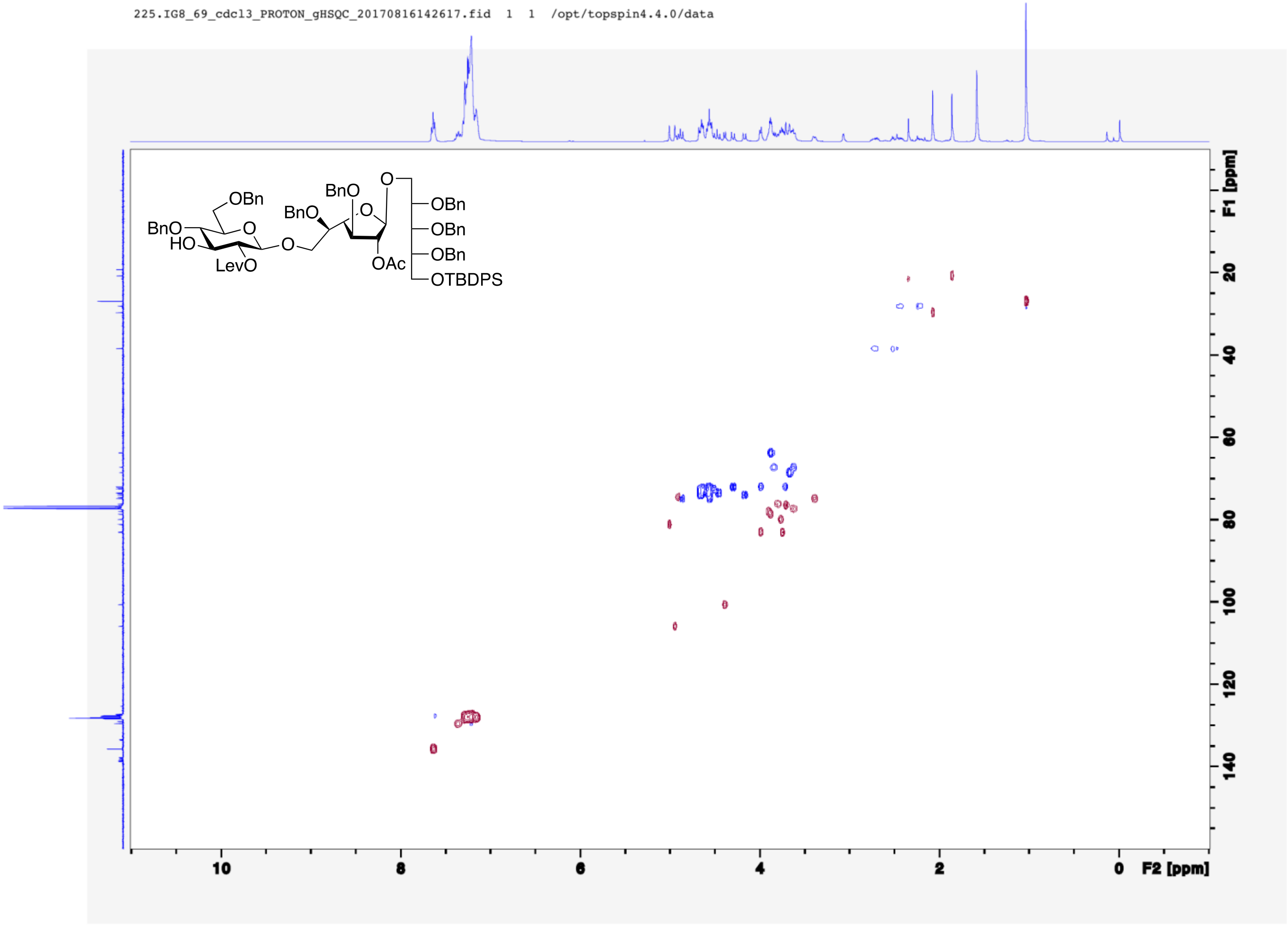
225.IG8_69_cdcl3_PROTON_gHSQC_20170816142617.fid 1 1 /opt/topspin4.4.0/data
OBn
BnO
BnO
O
HO
LevO
O
BnO
O
OAc
O
OBn
OBn
OBn
OTBDPS
F1 [ppm]
20
40
60
80
100
120
140
10
8
6
4
2
0
F2 [ppm]

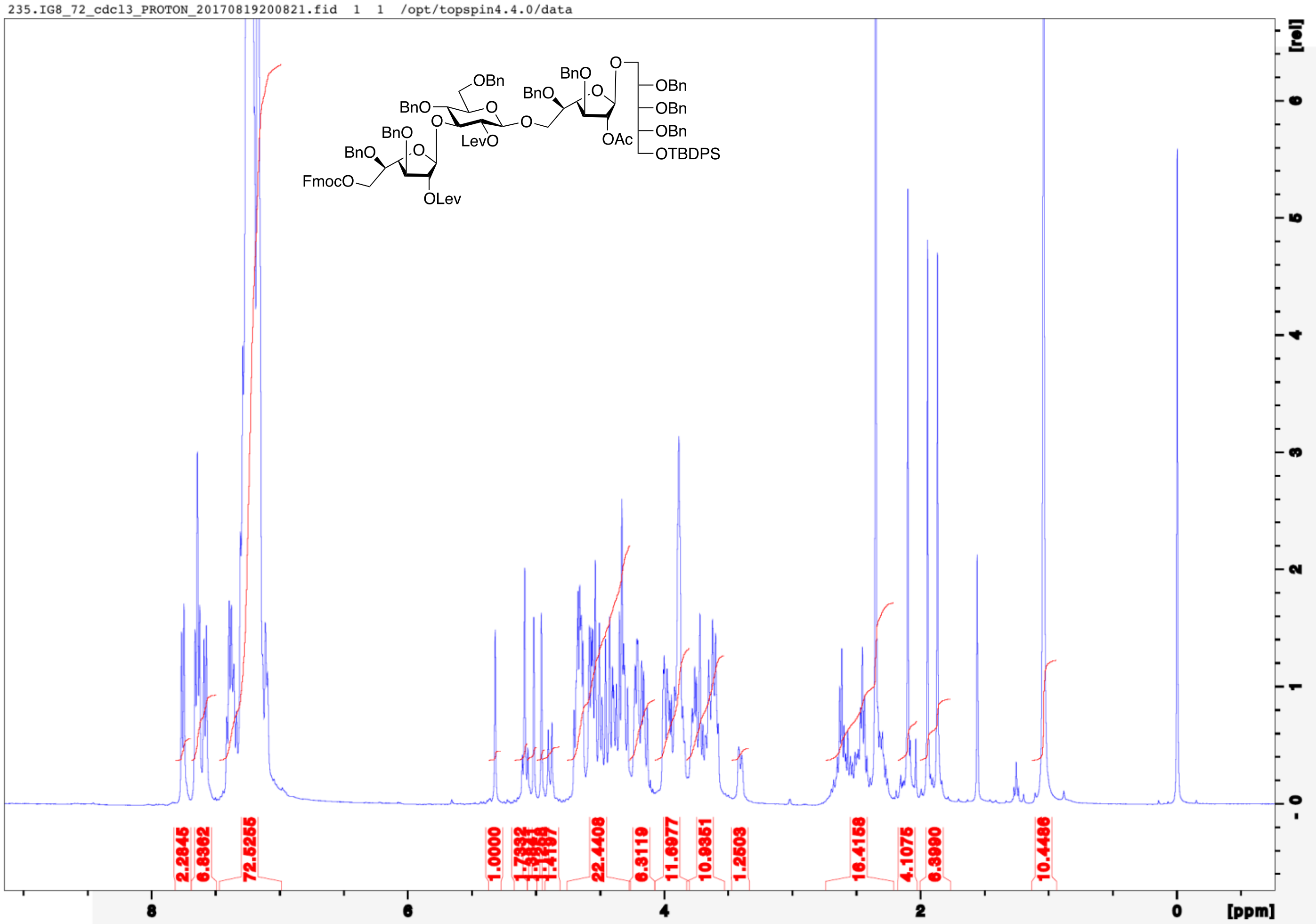
235.IG8_72_cdcl3_PROTON_20170819200821.fid 1 1 /opt/topspin4.4.0/data
OBn
BnO
BnO
O
O
O
LevO
BnO
O
BnO
O
OAc
OBn
OBn
OBn
OTBDPS
BnO
BnO
O
FmocO
OLev
2.2845
6.8362
72.5255
1.0000
22.4408
6.3119
11.6977
10.9351
1.2503
16.4158
4.1075
6.3980
10.4486
8
6
4
2
0
[ppm]
[rel]
6
5
4
3
2
1
0

234.IG8_72_cdcl3_CARBON_20170820192012.fid 1 1 /opt/topspin4.4.0/data

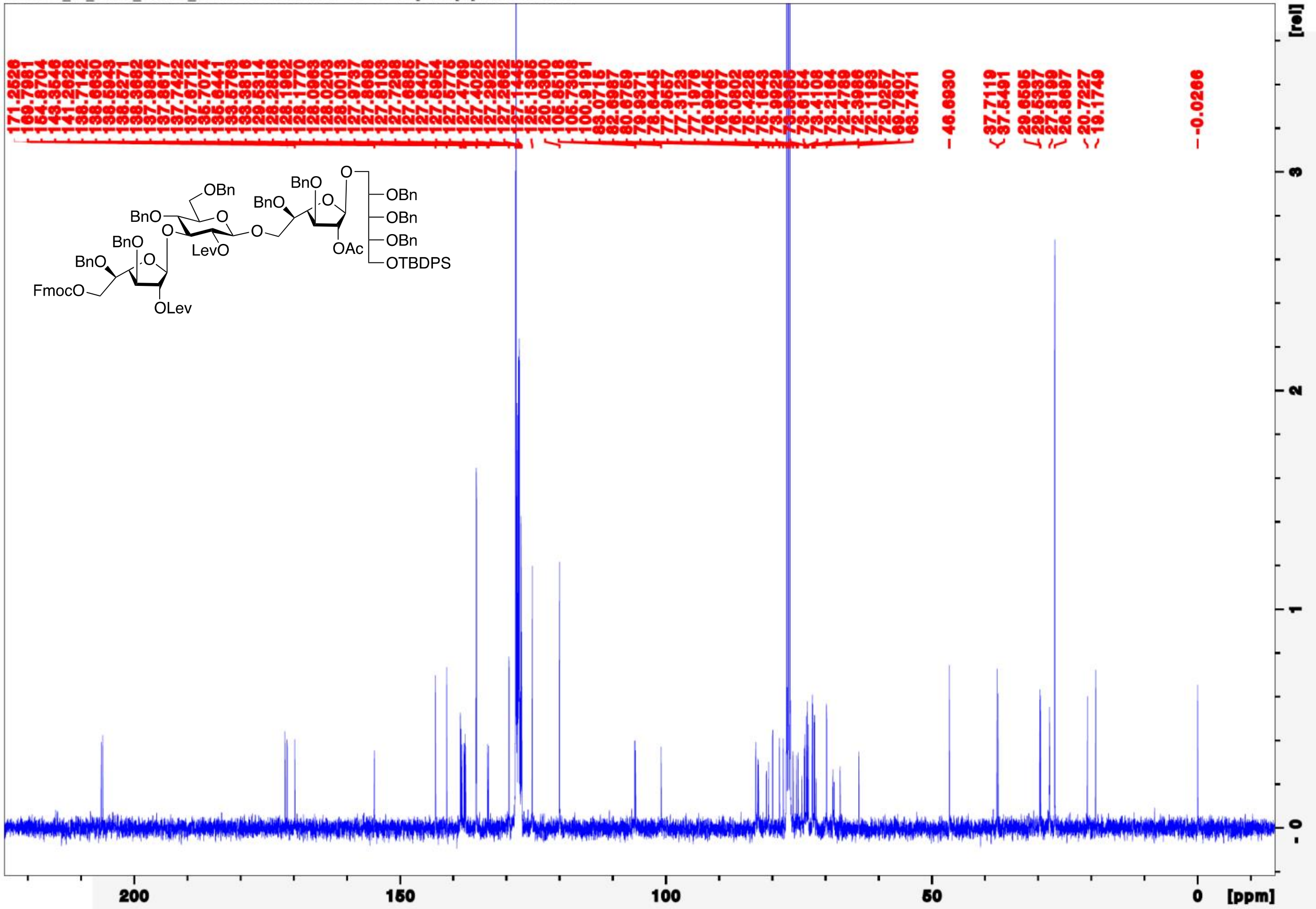

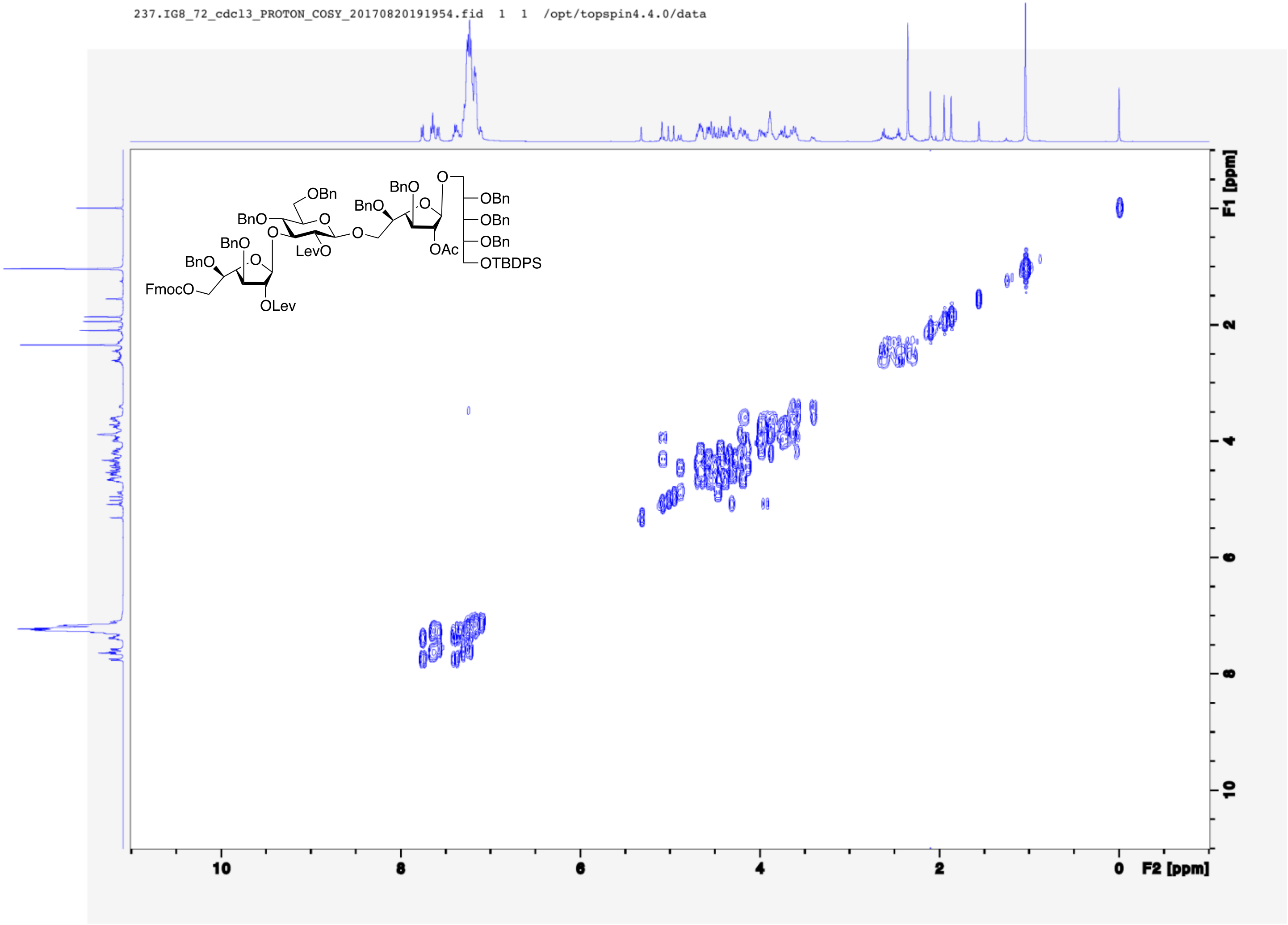
237.IG8_72_cdcl3_PROTON_COSY_20170820191954.fid 1 1 /opt/topspin4.4.0/data
OBn
BnO
O
O
BnO
O
BnO
BnO
OBn
OBn
OBn
OTBDPS
OAc
LevO
BnO
BnO
FmocO
OLev
F1 [ppm]
F2 [ppm]
10
8
6
4
2
0

238.IG8_72_cdcl3_PROTON_gHSQC_20170820192001.fid 1 1 /opt/topspin4.4.0/data

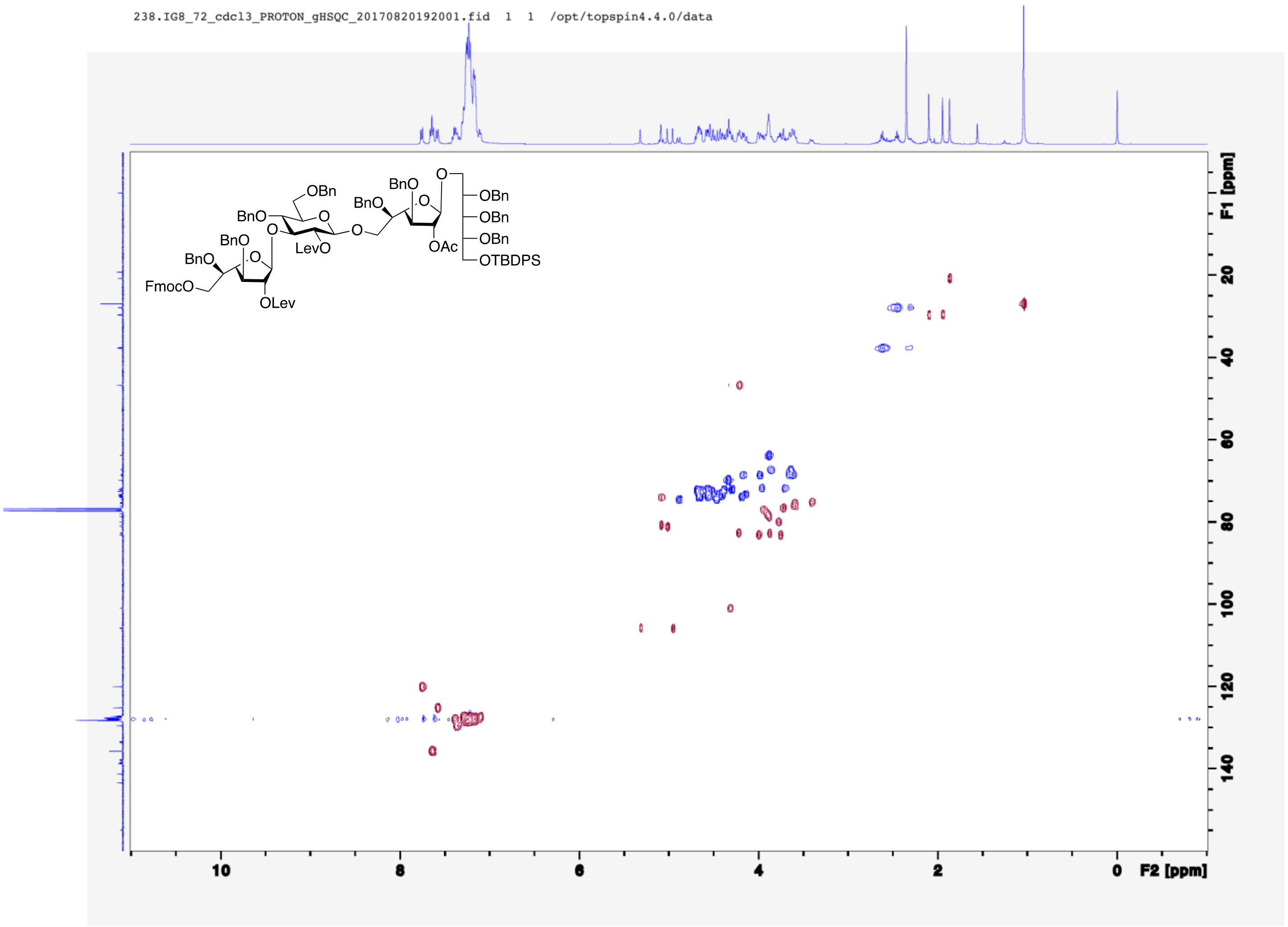

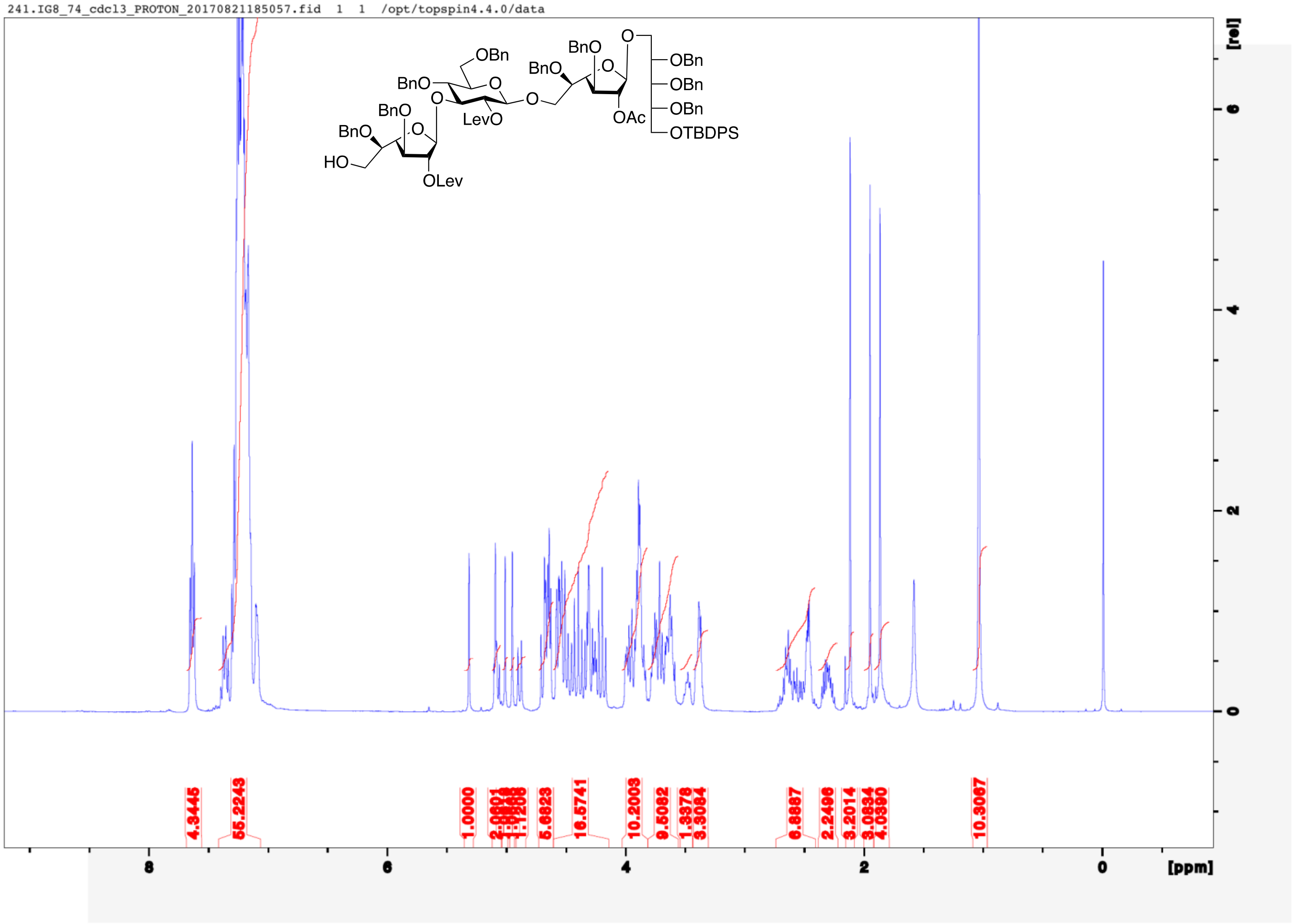
241.IG8_74_cdcl3_PROTON_20170821185057.fid 1 1 /opt/topspin4.4.0/data
OBn
BnO
O
BnO
BnO
O
O
O
BnO
LevO
O
OBn
OBn
OBn
OTBDPS
BnO
OAc
HO
OLev
[rel]
6
4
2
0
4.3445
55.2243
1.0000
5.6823
16.5741
10.2003
9.5082
3.3084
2.2496
3.2014
10.3067
8
6
4
2
0
[ppm]

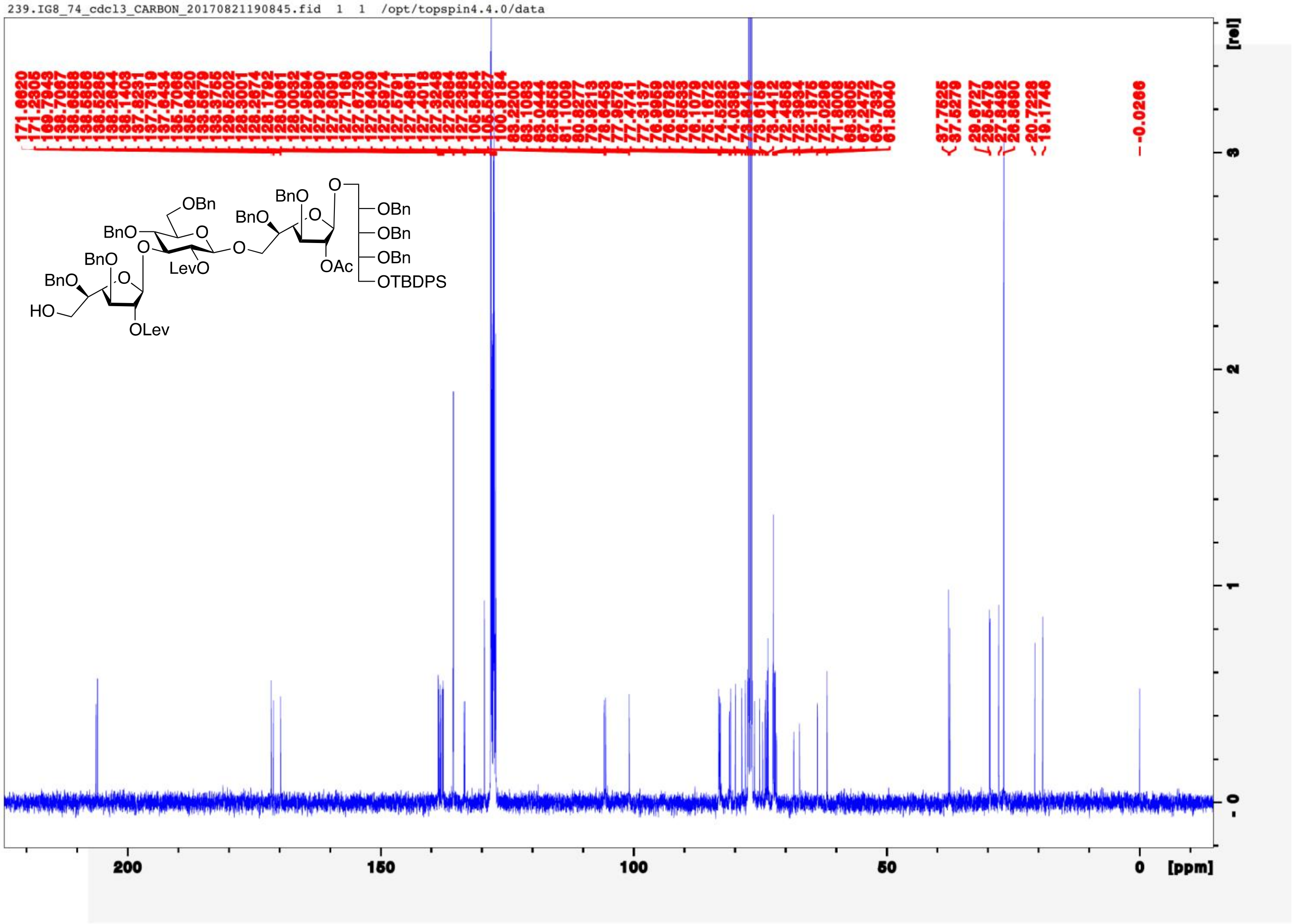
239.IG8_74_cdcl3_CARBON_20170821190845.fid 1 1 /opt/topspin4.4.0/data
OBn
BnO
BnO
O
O
LevO
O
BnO
BnO
HO
O
OLev
BnO
BnO
O
O
OAc
OBn
OBn
OBn
OTBDPS
200
150
100
50
0
[ppm]
[rel]

242.IG8_74_cdcl3_PROTON_COSY_20170821185124.fid 1 1 /opt/topspin4.4.0/data

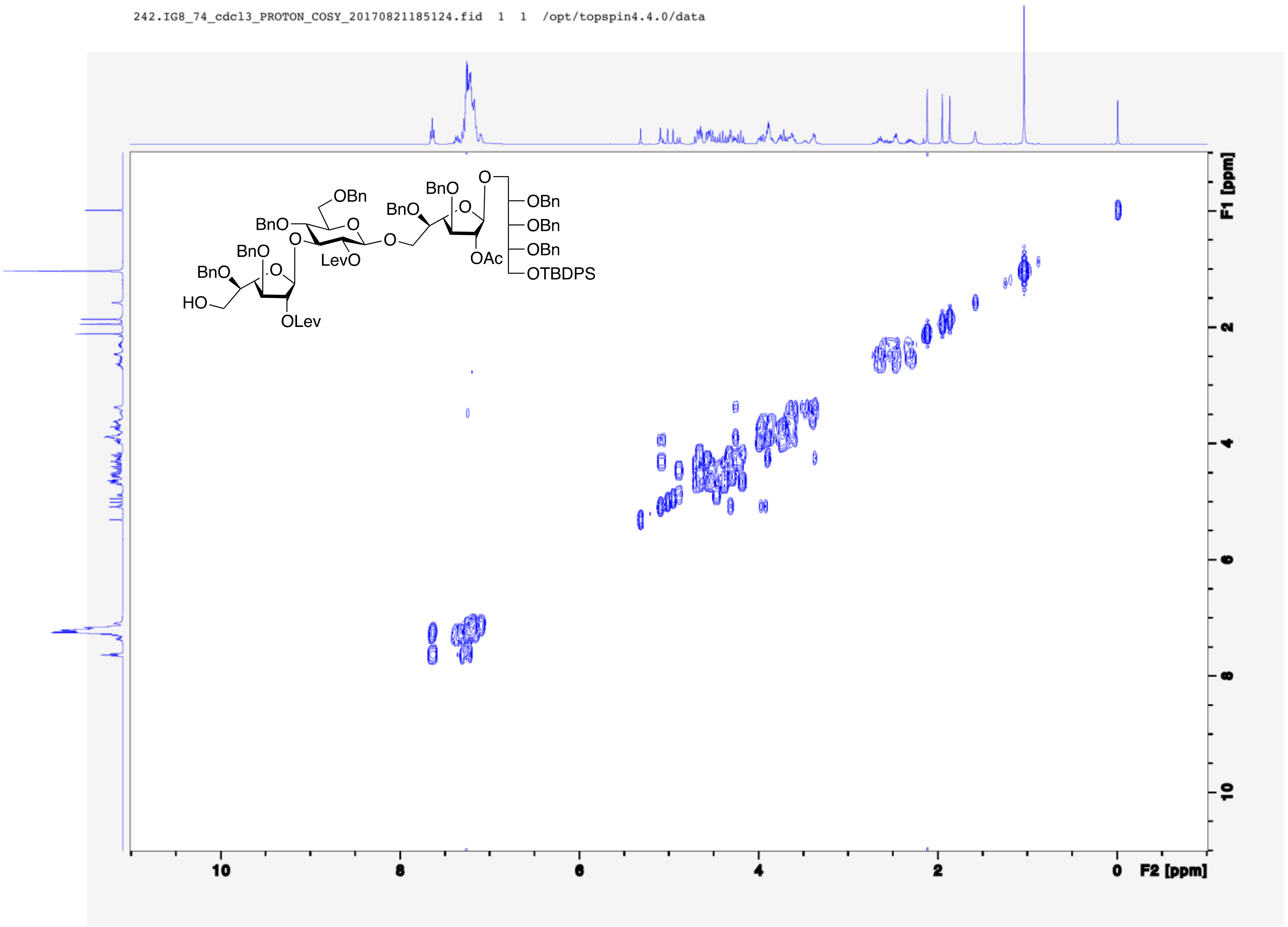

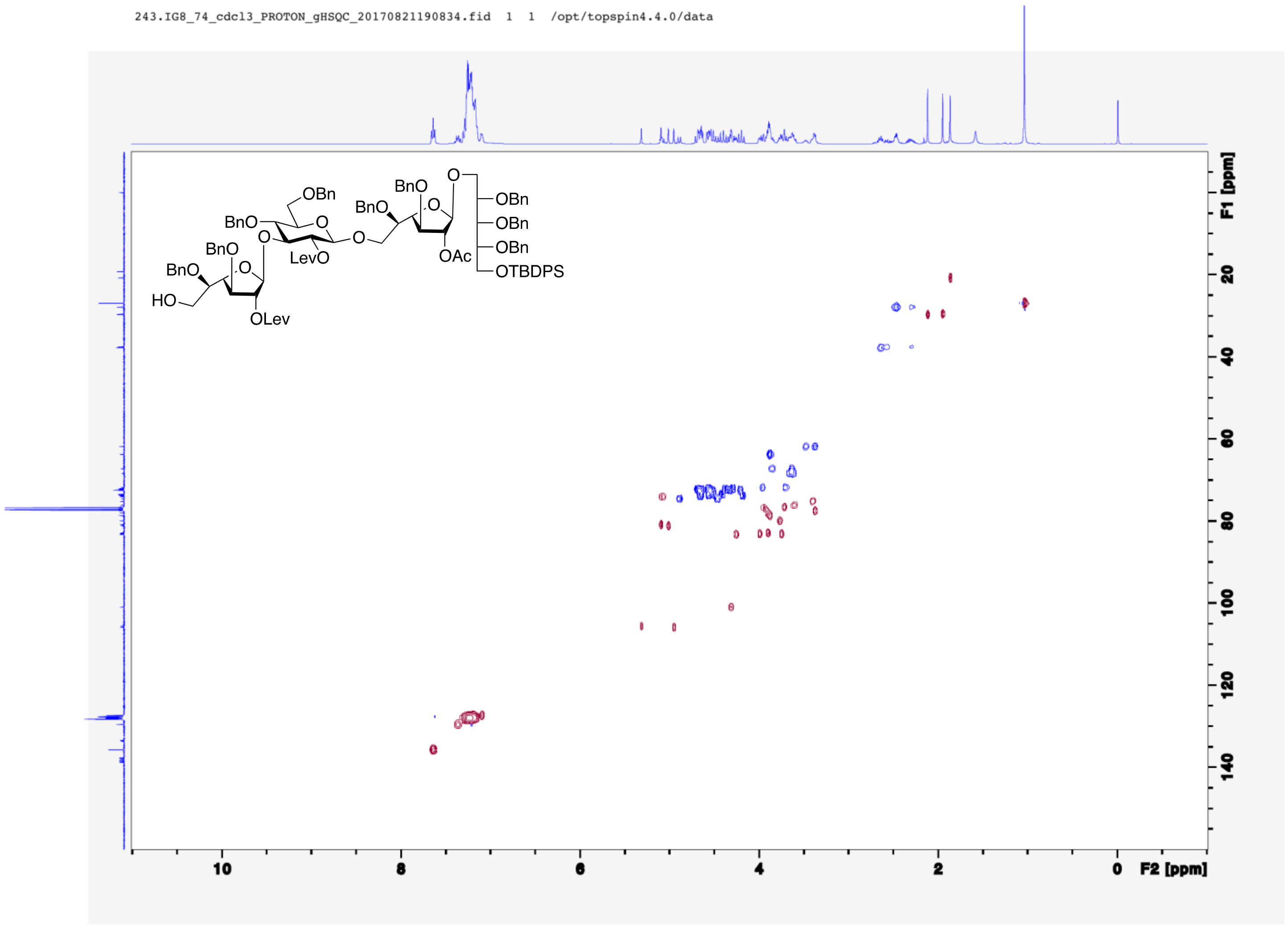
243.IG8_74_cdcl3_PROTON_gHSQC_20170821190834.fid 1 1 /opt/topspin4.4.0/data
OBn
BnO
BnO
O
OBn
OBn
OBn
OTBDPS
OAc
LevO
BnO
BnO
HO
OLev
F1 [ppm]
F2 [ppm]

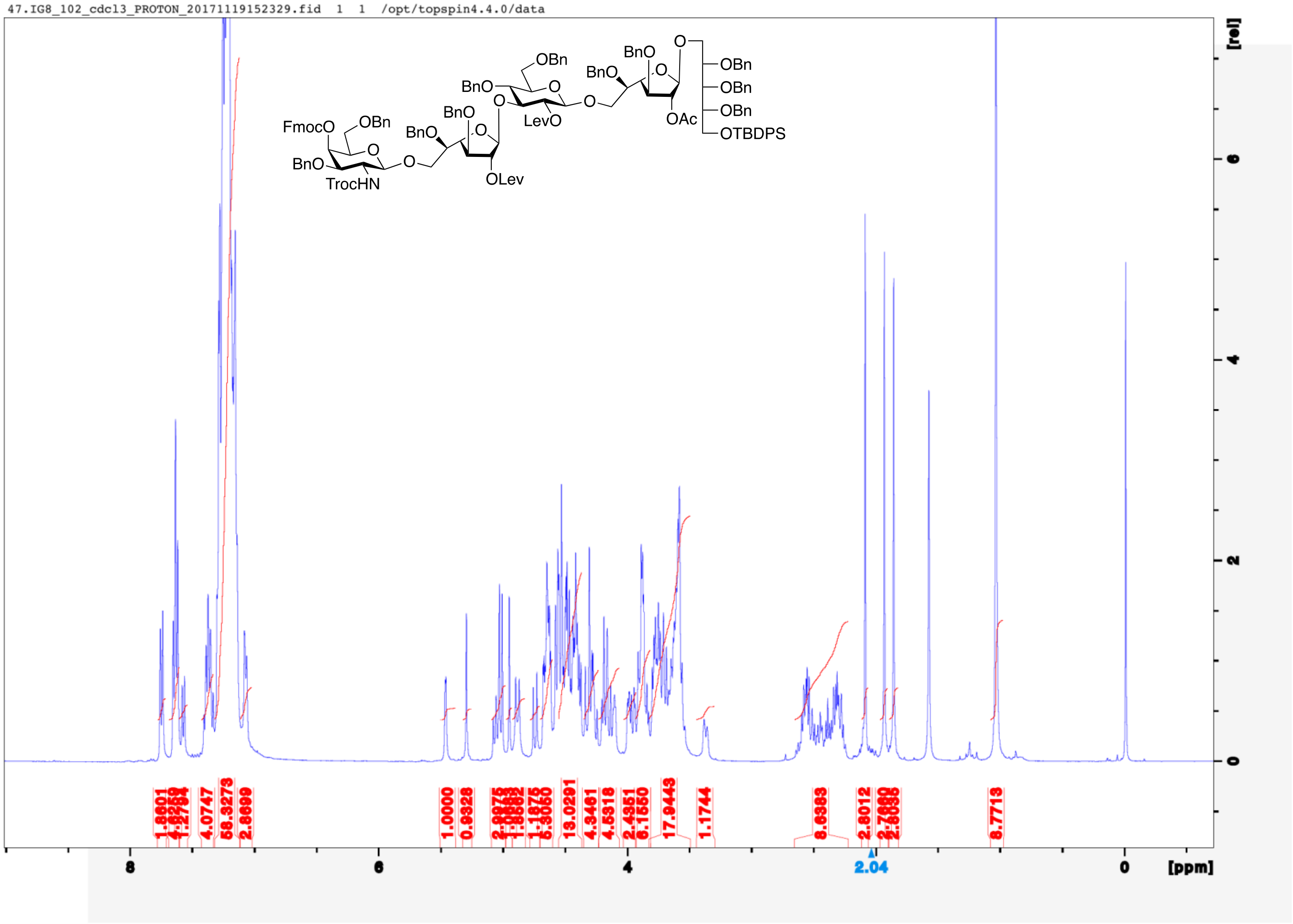
47.IG8_102_cdcl3_PROTON_20171119152329.fid 1 1 /opt/topspin4.4.0/data
FmocO
OBn
BnO
TrocHN
O
BnO
BnO
OLev
BnO
OBn
LevO
BnO
BnO
OAc
OBn
OBn
OBn
OTBDPS
[rel]
[ppm]
8
6
4
2.04
0

44.IG8_102_cdcl3_CARBON_20171119161045.fid 1 1 /opt/topspin4.4.0/data

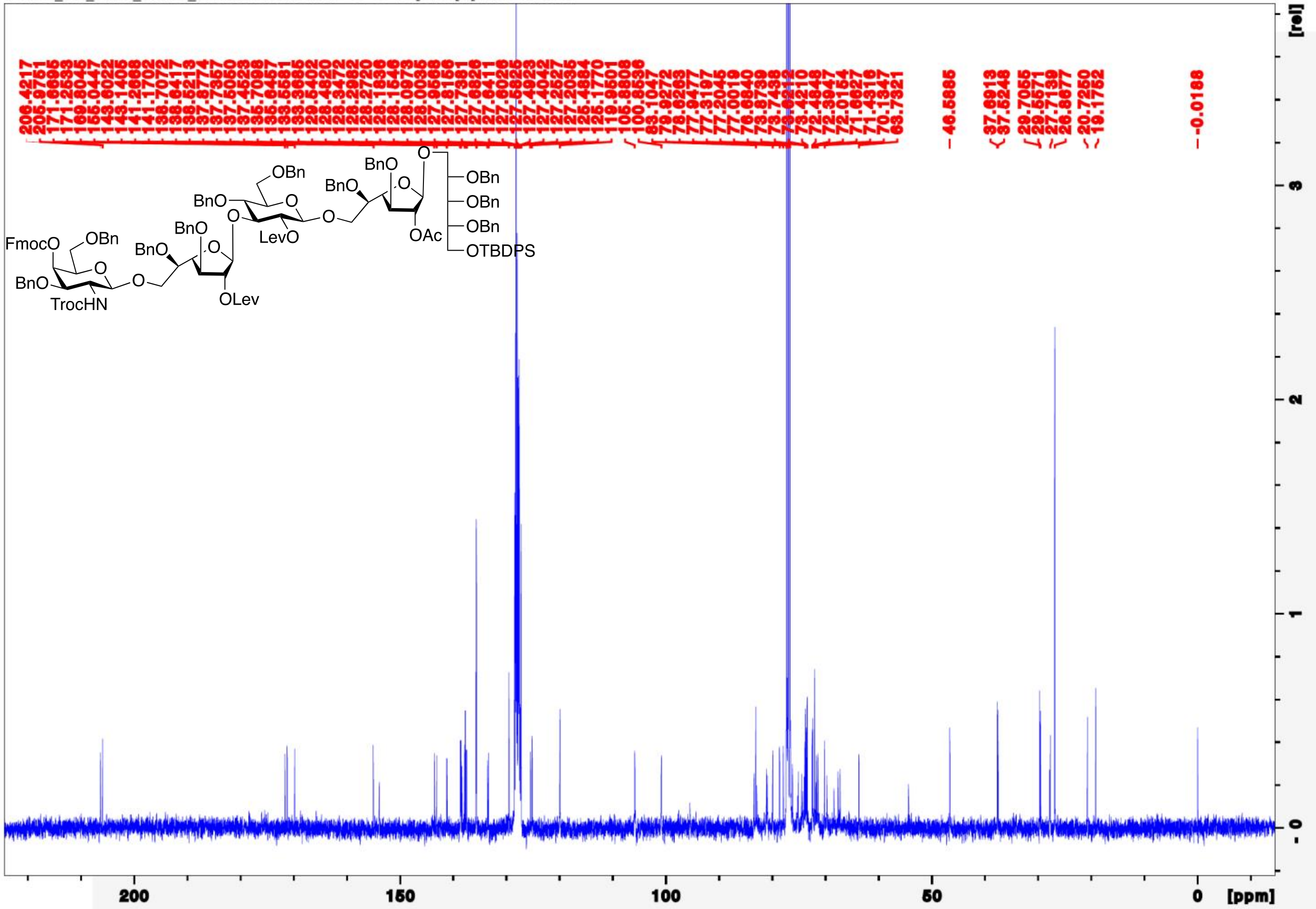

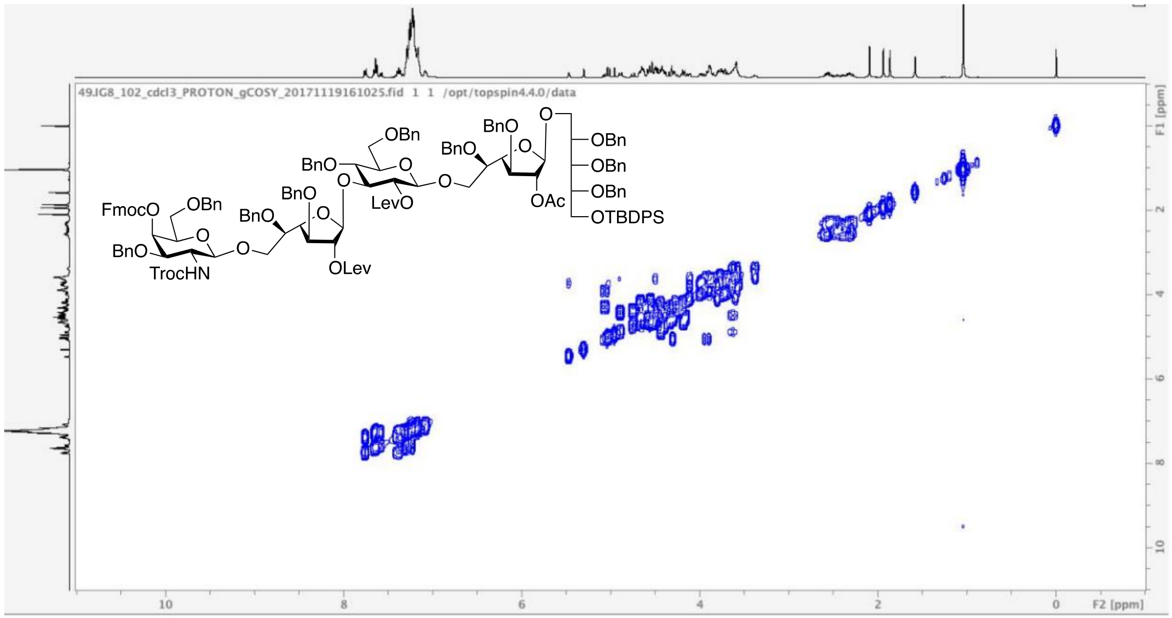
49.IG8_102_cdcl3_PROTON_gCOSY_20171119161025.fid 1 1 /opt/topspin4.4.0/data
FmocO
OBn
BnO
TrocHN
O
BnO
BnO
O
OLev
BnO
OBn
LevO
BnO
BnO
OAc
OBn
OBn
OBn
OTBDPS
F1 [ppm]
F2 [ppm]
10
8
6
4
2
0

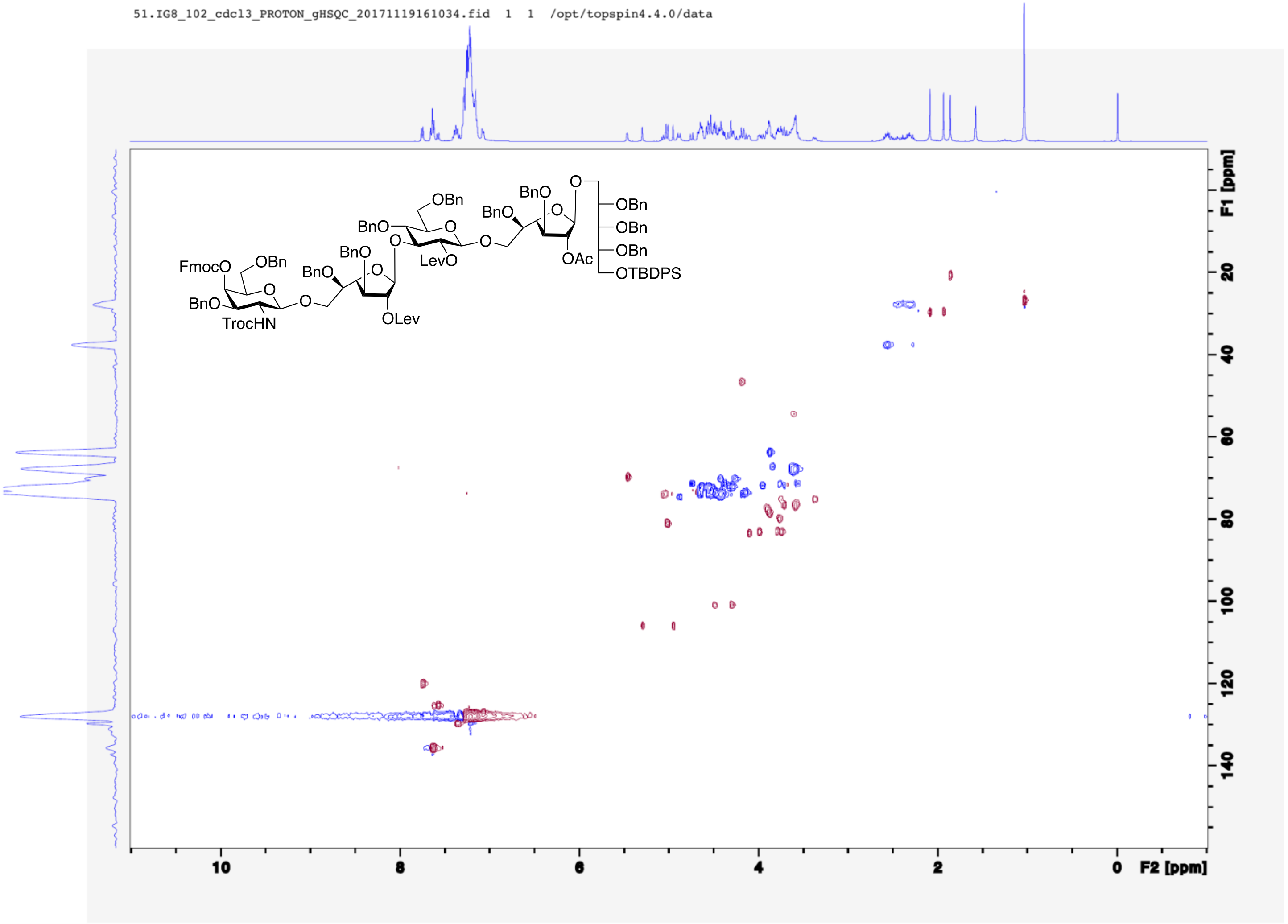
51.IG8_102_cdcl3_PROTON_gHSQC_20171119161034.fid 1 1 /opt/topspin4.4.0/data
FmocO
OBn
BnO
TrocHN
BnO
BnO
OLev
BnO
OBn
LevO
BnO
BnO
OAc
OBn
OBn
OBn
OTBDPS
F1 [ppm]
F2 [ppm]

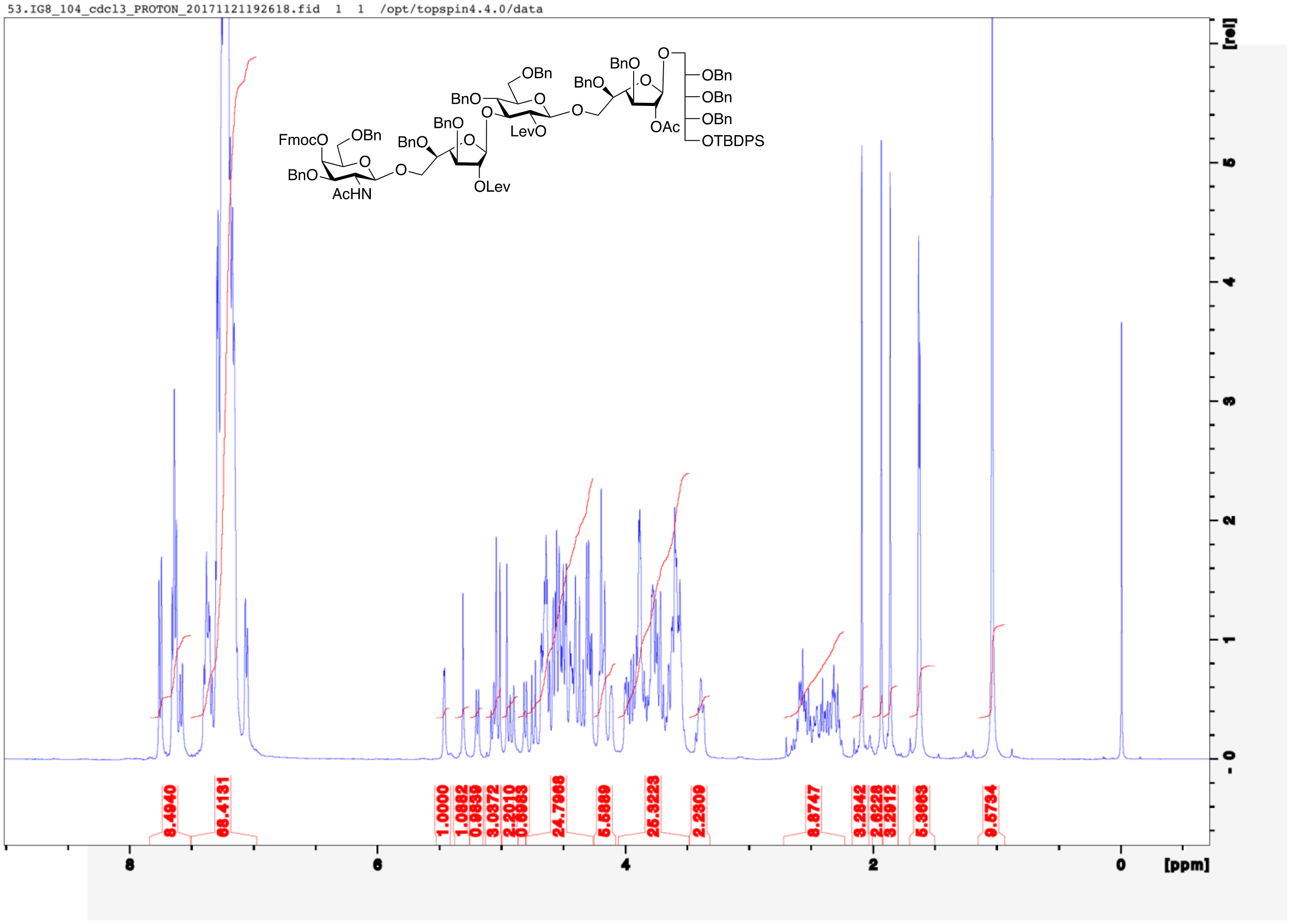

53.IG8_104_cdcl3_PROTON_20171121192618.fid 1 1 /opt/topspin4.4.0/data
FmocO
OBn
BnO
AcHN
BnO
BnO
OLev
BnO
OBn
LevO
BnO
BnO
OAc
OBn
OBn
OBn
OTBDPS
8.4940
68.4131
1.0000
1.0882
0.9839
3.0372
2.2010
0.6983
24.7068
5.5889
25.3223
2.2309
8.8747
3.2842
2.6228
3.2912
5.3663
9.5734
8
6
4
2
0
[ppm]
[rel]

52.IG8_104_cdcl3_CARBON_20171122162130.fid 1 1 /opt/topspin4.4.0/data

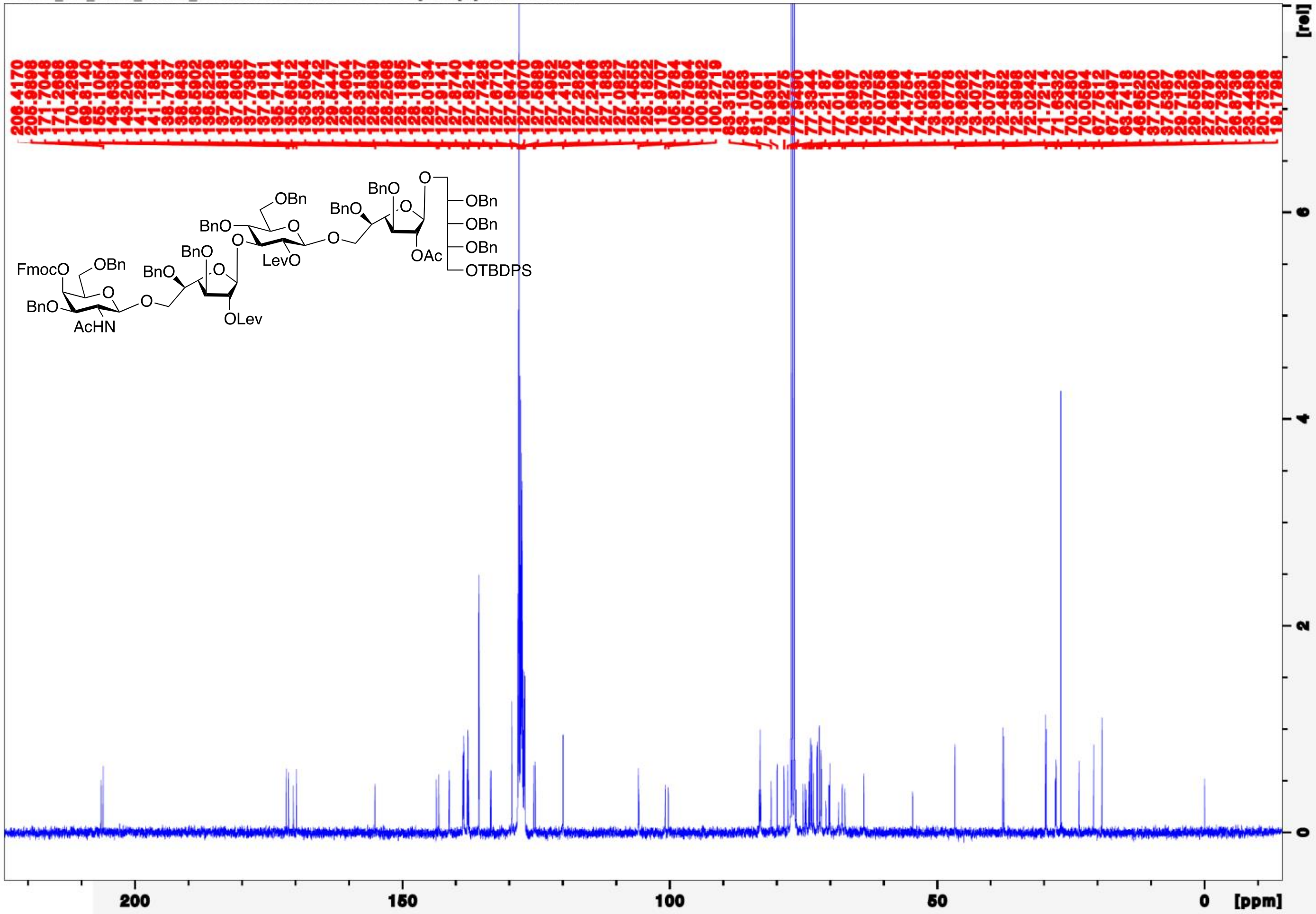

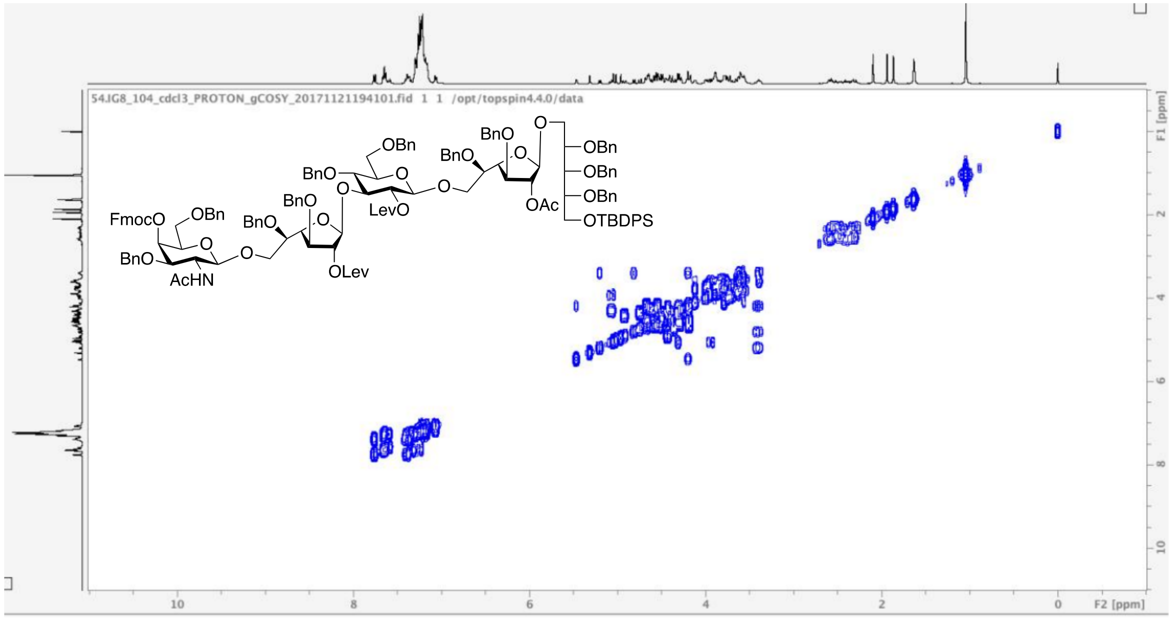
54JG8_104_cdcl3_PROTON_gCOSY_20171121194101.fid 1 1 /opt/topspin4.4.0/data
FmocO
OBn
BnO
AcHN
O
BnO
OLev
BnO
OBn
LevO
BnO
OAc
OBn
OBn
OBn
OTBDPS
F1 [ppm]
F2 [ppm]
10
8
6
4
2
0

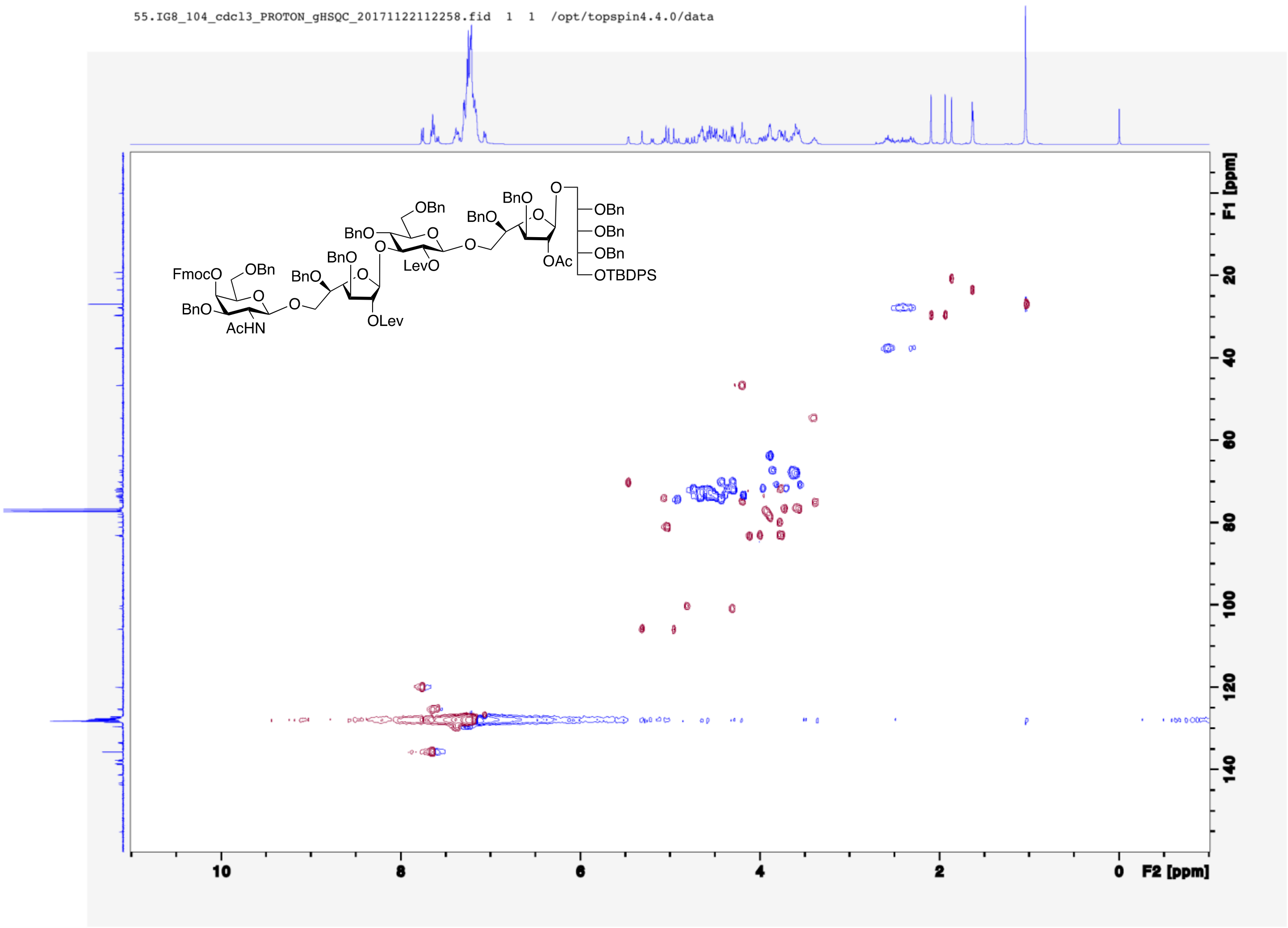
55.IG8_104_cdcl3_PROTON_gHSQC_20171122112258.fid 1 1 /opt/topspin4.4.0/data
F1 [ppm]
F2 [ppm]
FmocO
OBn
BnO
AcHN
OLev
LevO
OAc
OTBDPS

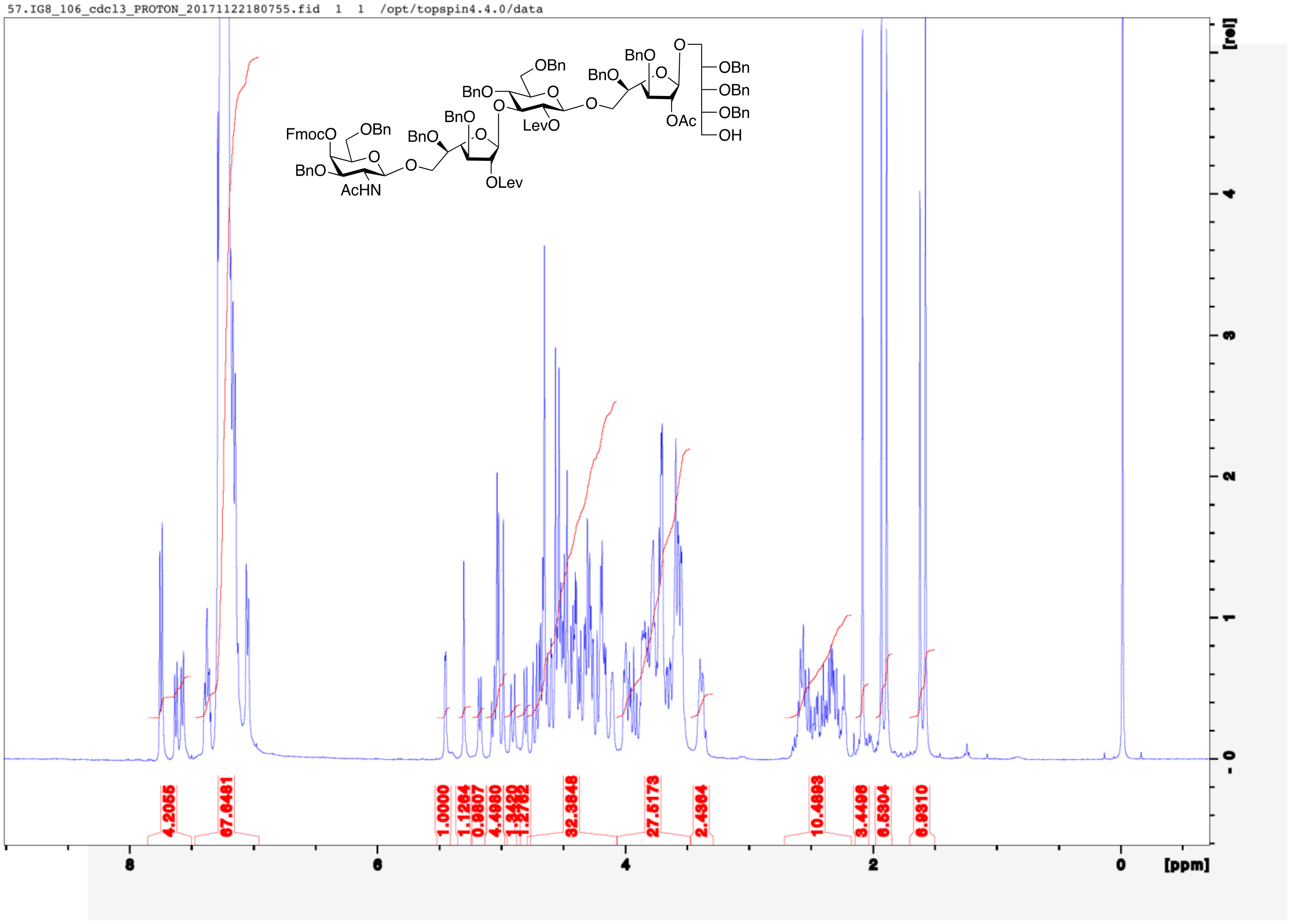
57.IG8_106_cdcl3_PROTON_20171122180755.fid 1 1 /opt/topspin4.4.0/data
FmocO
OBn
BnO
AcHN
O
BnO
BnO
OLev
BnO
OBn
LevO
BnO
BnO
OAc
OBn
OBn
OBn
OH
4.2055
67.6481
1.0000
1.1264
0.9807
4.4980
1.3420
1.2762
32.3848
27.5173
2.4364
10.4893
3.4496
6.5304
6.9310
8
6
4
2
0
[ppm]
[rel]

56.IG8_106_cdcl3_CARBON_20171122180845.fid 1 1 /opt/topspin4.4.0/data

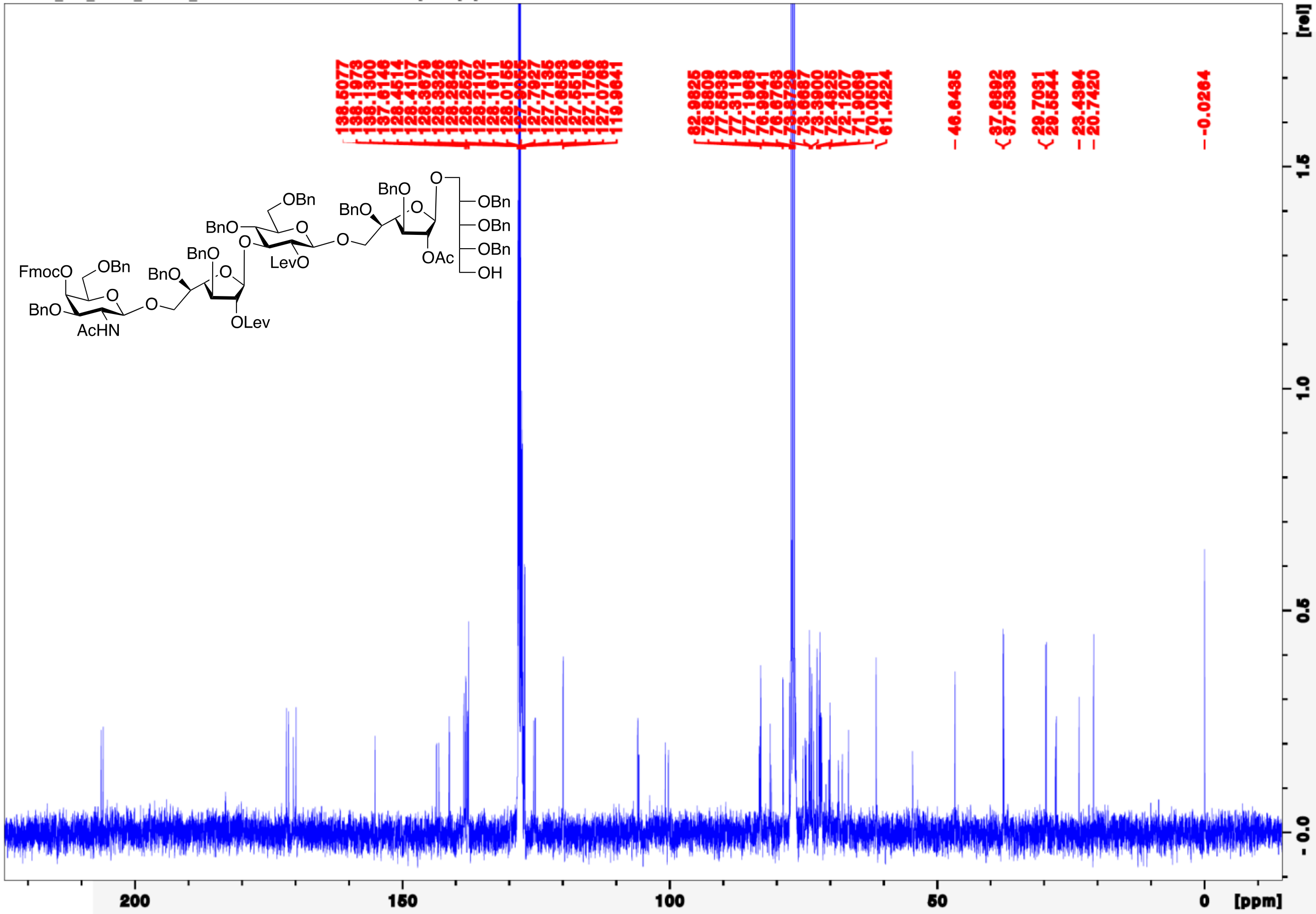

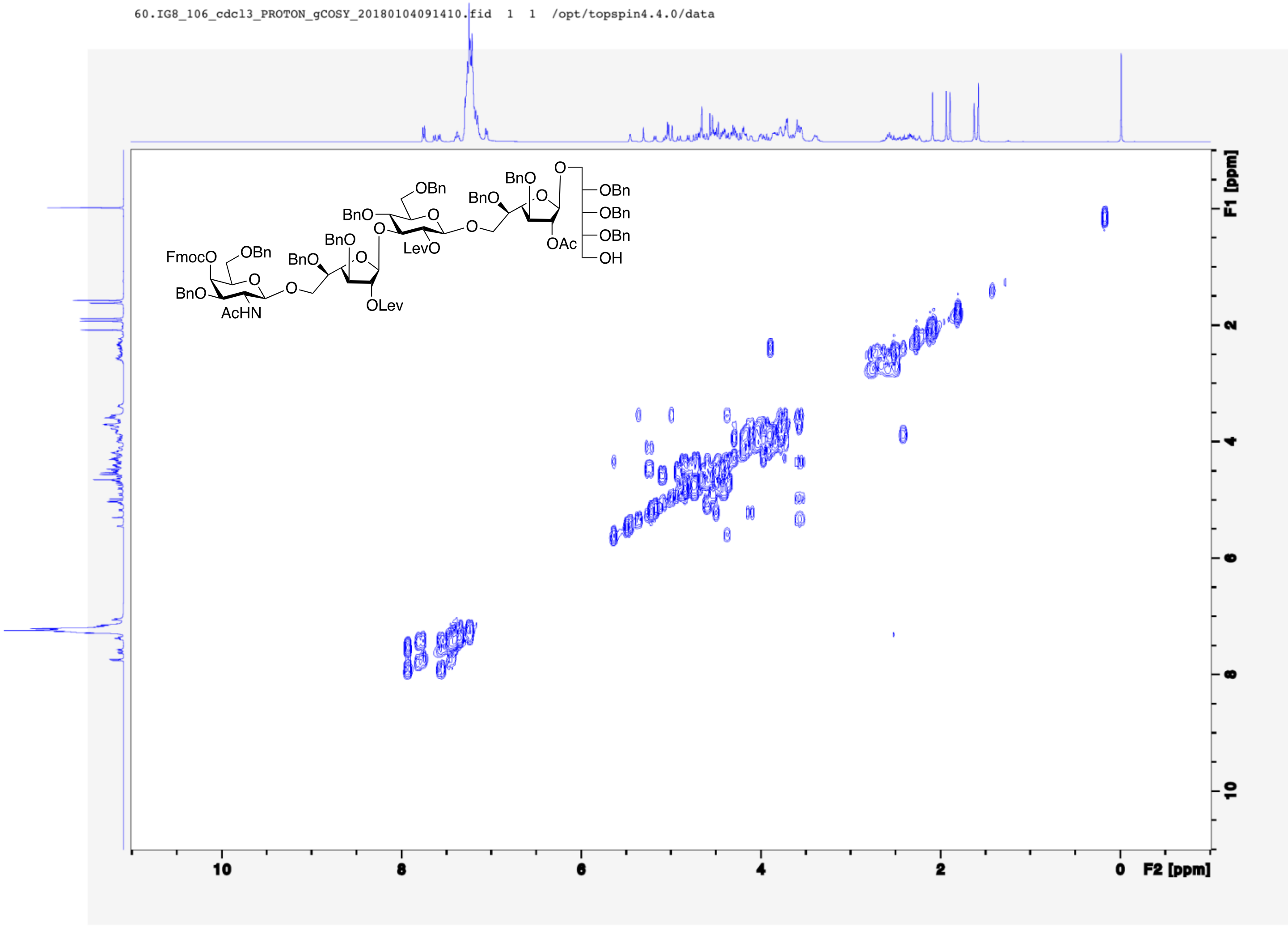
60.IG8_106_cdcl3_PROTON_gCOSY_20180104091410.fid 1 1 /opt/topspin4.4.0/data
FmocO
OBn
BnO
AcHN
O
BnO
OLev
LevO
OAc
OH
F1 [ppm]
F2 [ppm]

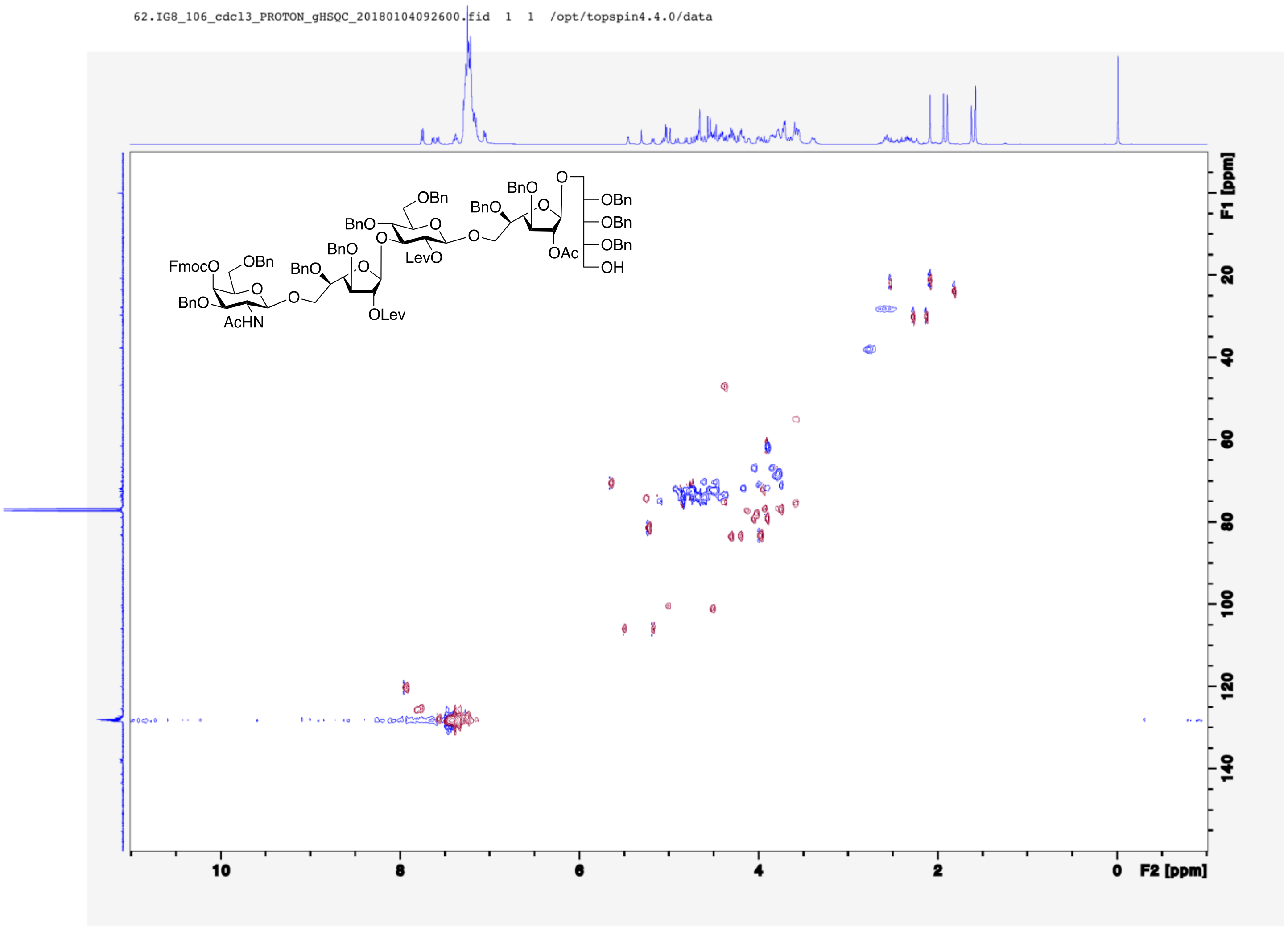
62.IG8_106_cdcl3_PROTON_gHSQC_20180104092600.fid 1 1 /opt/topspin4.4.0/data
FmocO
OBn
BnO
AcHN
O
BnO
OLev
BnO
LevO
OAc
OH
F1 [ppm]
F2 [ppm]

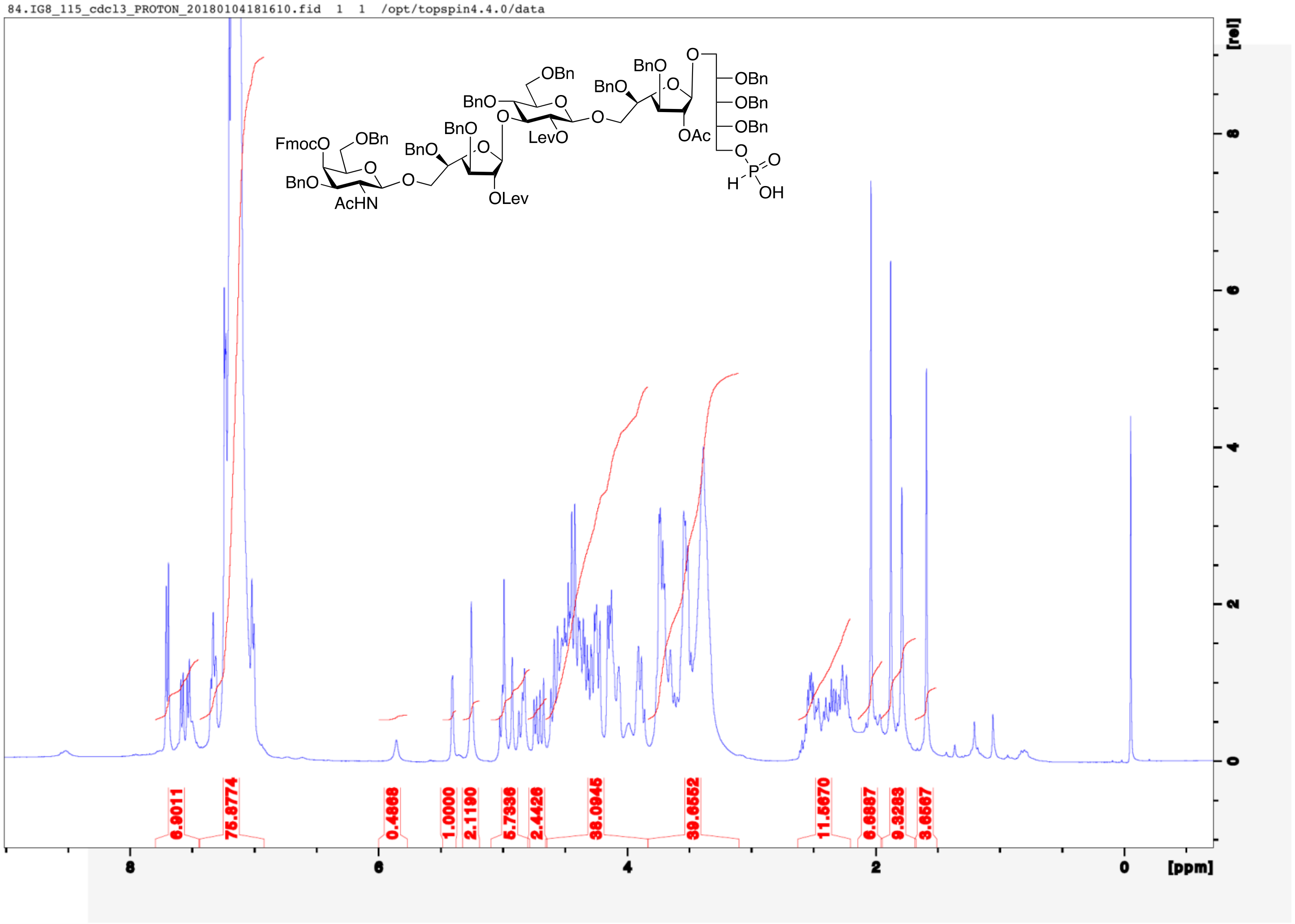
84.IG8_115_cdcl3_PROTON_20180104181610.fid 1 1 /opt/topspin4.4.0/data
FmocO
OBn
BnO
AcHN
BnO
BnO
OLev
BnO
OBn
LevO
BnO
BnO
OAc
OBn
OBn
OBn
H
P
OH
0.9011
75.8774
0.4866
1.0000
2.1190
5.7336
2.4426
38.0945
39.8552
11.5670
6.6887
9.3283
3.6567
8
6
4
2
0
[ppm]
[rel]

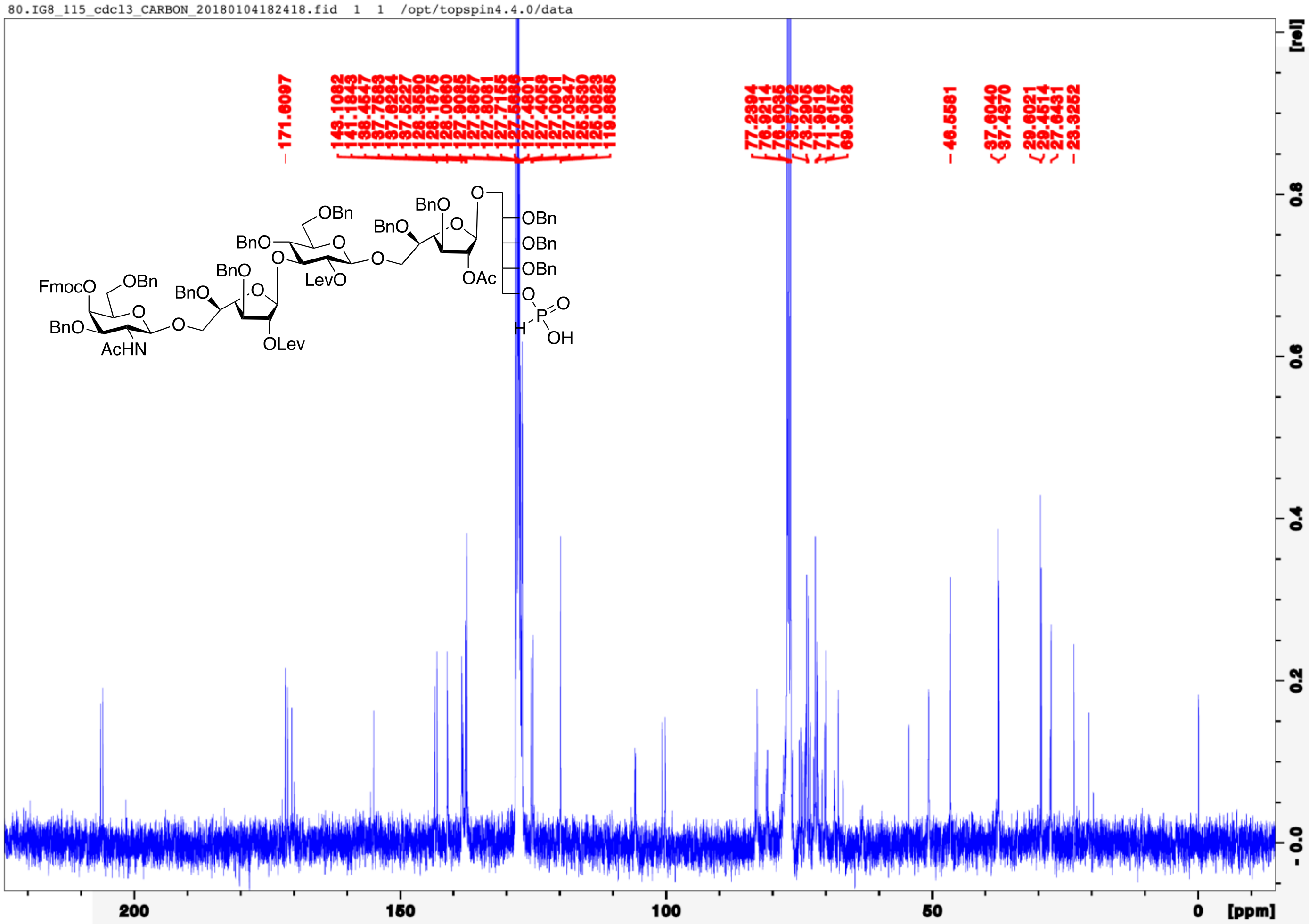
80.IG8_115_cdcl3_CARBON_20180104182418.fid 1 1 /opt/topspin4.4.0/data
171.6097
143.1082
141.1843
138.4547
137.7583
137.6284
137.5220
128.3590
128.1875
128.0660
127.9085
127.8657
127.8081
127.7155
127.5686
127.4801
127.4058
127.0901
127.0347
125.3530
125.0823
119.8685
77.2394
76.9214
76.6035
73.5702
73.2905
71.9516
71.6157
69.9628
46.5581
37.6040
37.4370
29.6021
29.4514
27.8431
23.3252
FmocO
OBn
BnO
AcHN
O
BnO
BnO
OLev
BnO
OBn
LevO
BnO
BnO
OAc
OBn
OBn
OBn
O
H
P
O
OH
[rel]
0.8
0.6
0.4
0.2
0.0
200
150
100
50
0
[ppm]

82.IG8_115_cdcl3_PHOSPHORUS_20180104181627.fid 1 1 /opt/topspin4.4.0/data

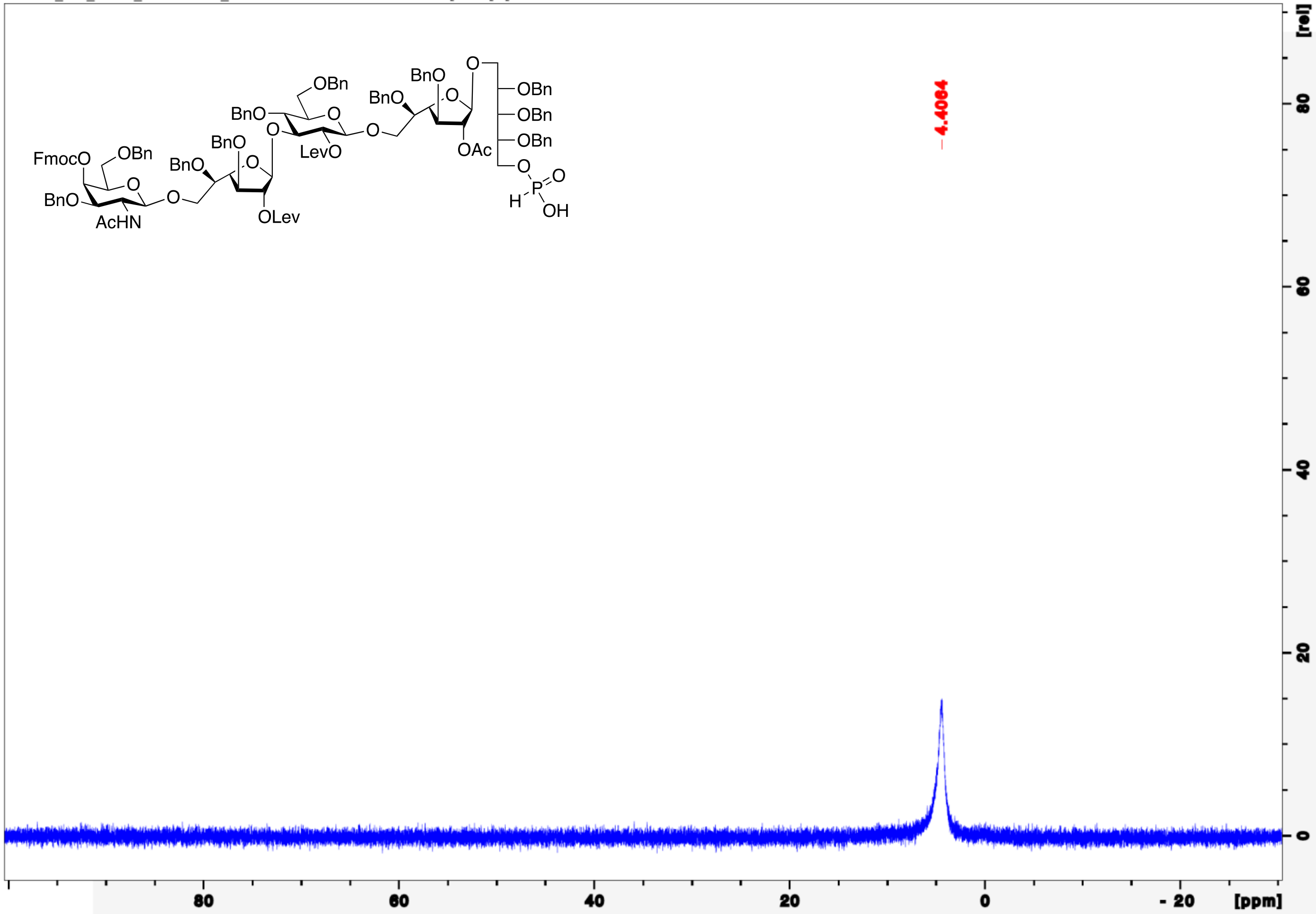

85.IG8_115_cdcl3_PROTON_gCOSY_20180104182314.fid 1 1 /opt/topspin4.4.0/data

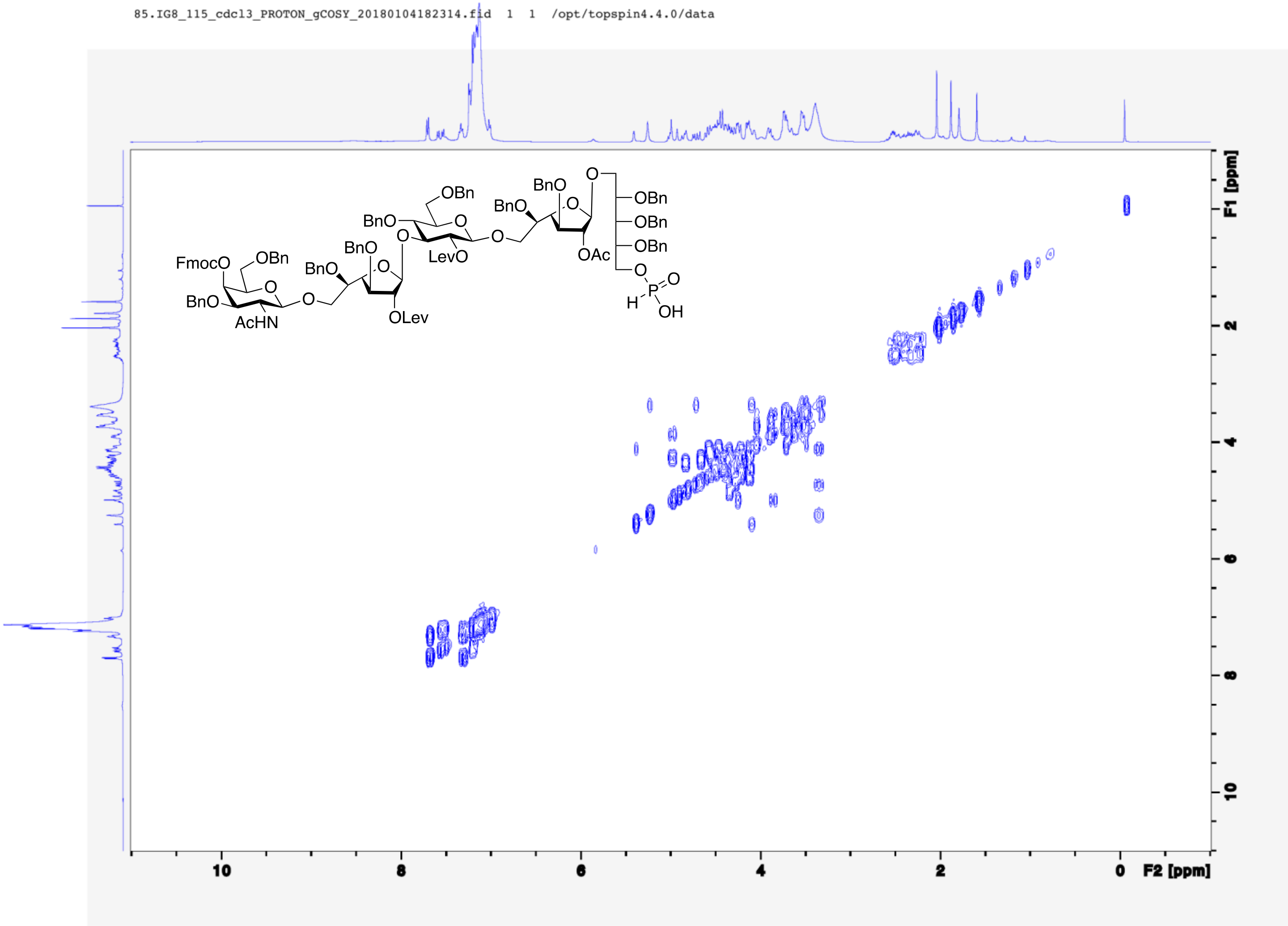

86.IG8_115_cdcl3_PROTON_gHSQC_20180104182337.fid 1 1 /opt/topspin4.4.0/data

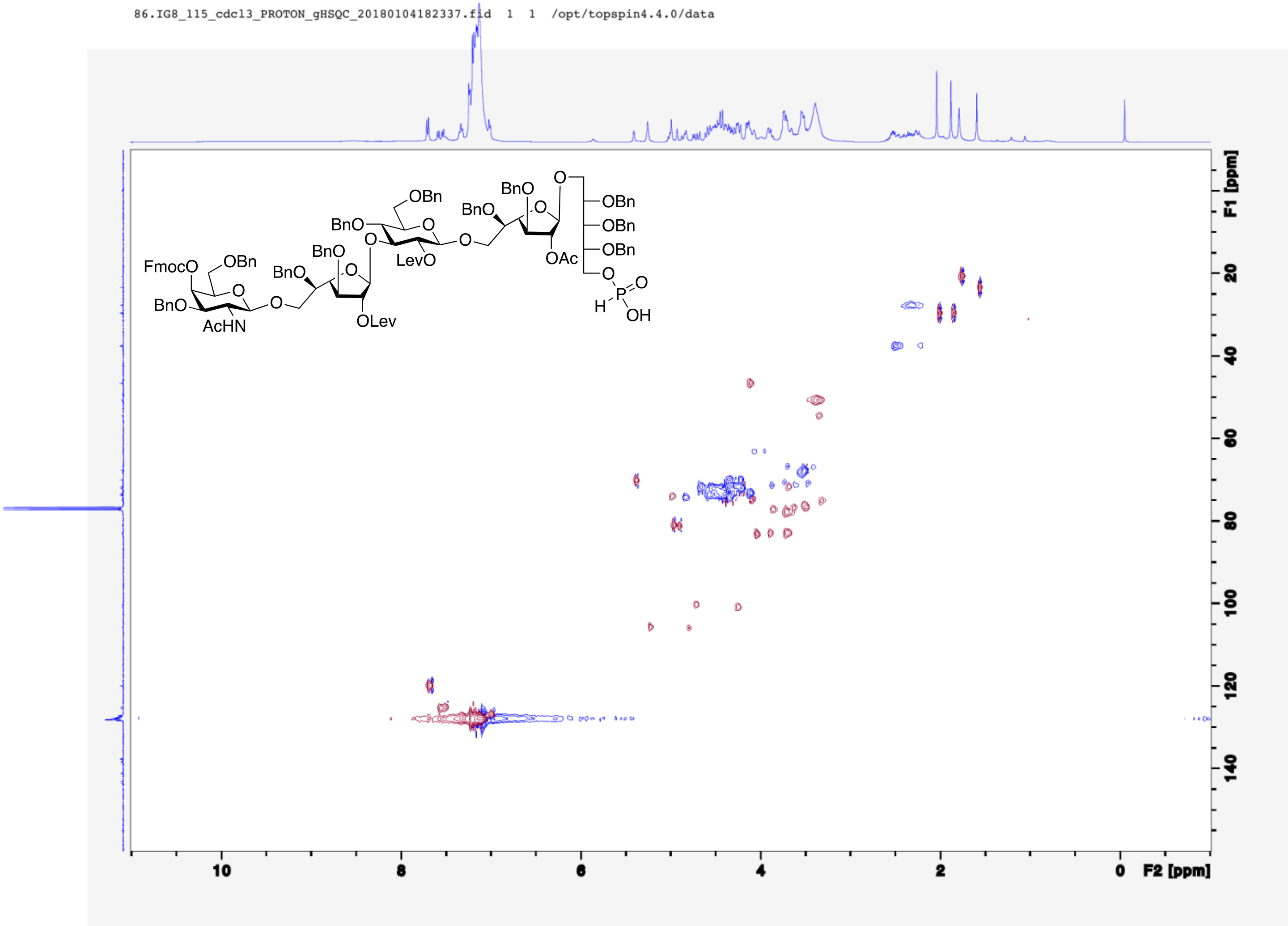

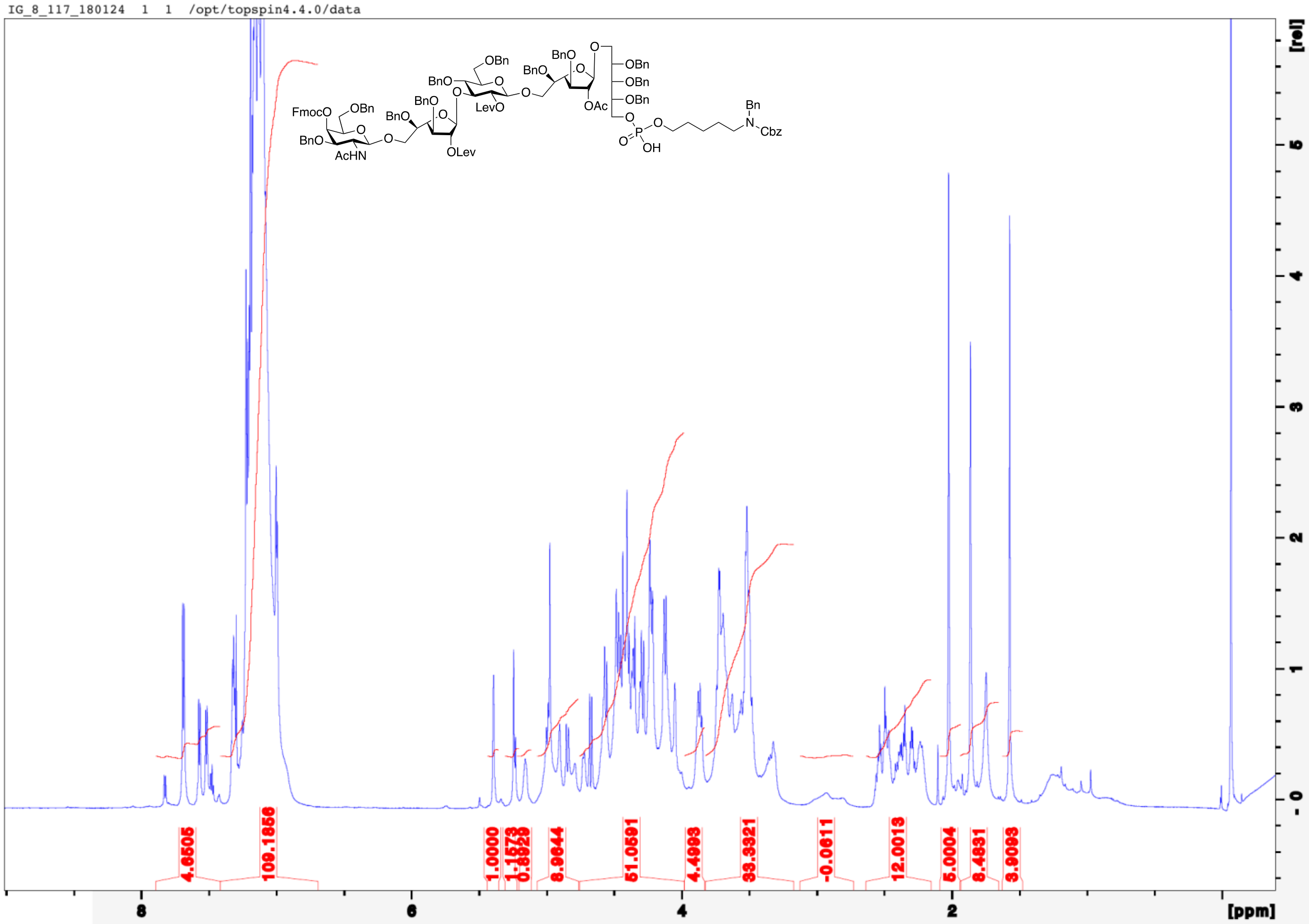
IG_8_117_180124 1 1 /opt/topspin4.4.0/data
FmocO
OBn
BnO
AcHN
BnO
OLev
LevO
OAc
Bn
N
Cbz
OH
4.6505
109.1856
1.0000
1.1573
0.8829
8.9644
51.0591
4.4993
33.3321
-0.0611
12.0013
5.0004
8.4931
3.9093
8
6
4
2
[ppm]
[rel]

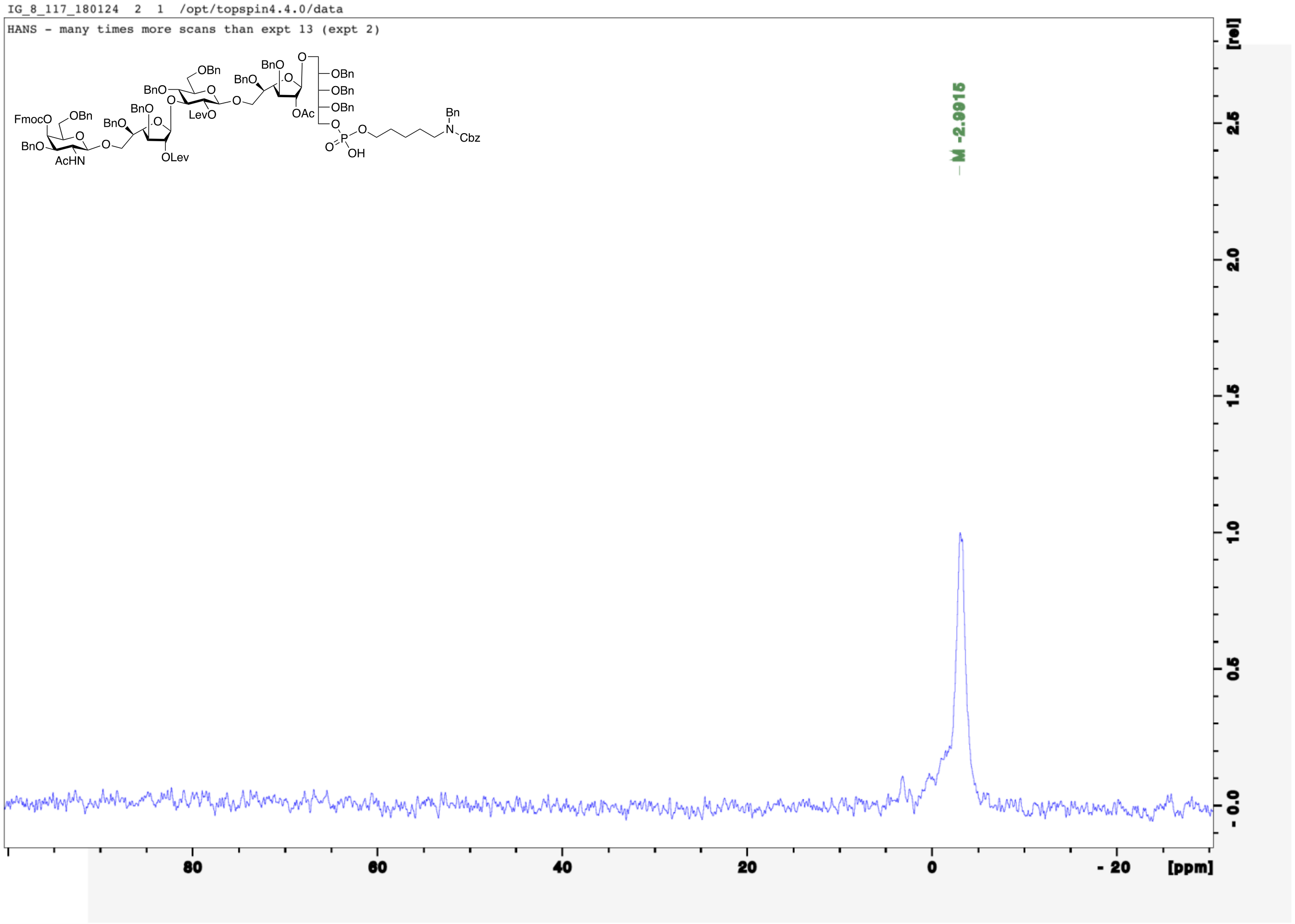
IG_8_117_180124 2 1 /opt/topspin4.4.0/data
HANS - many times more scans than expt 13 (expt 2)
-2.9915
[rel]
[ppm]

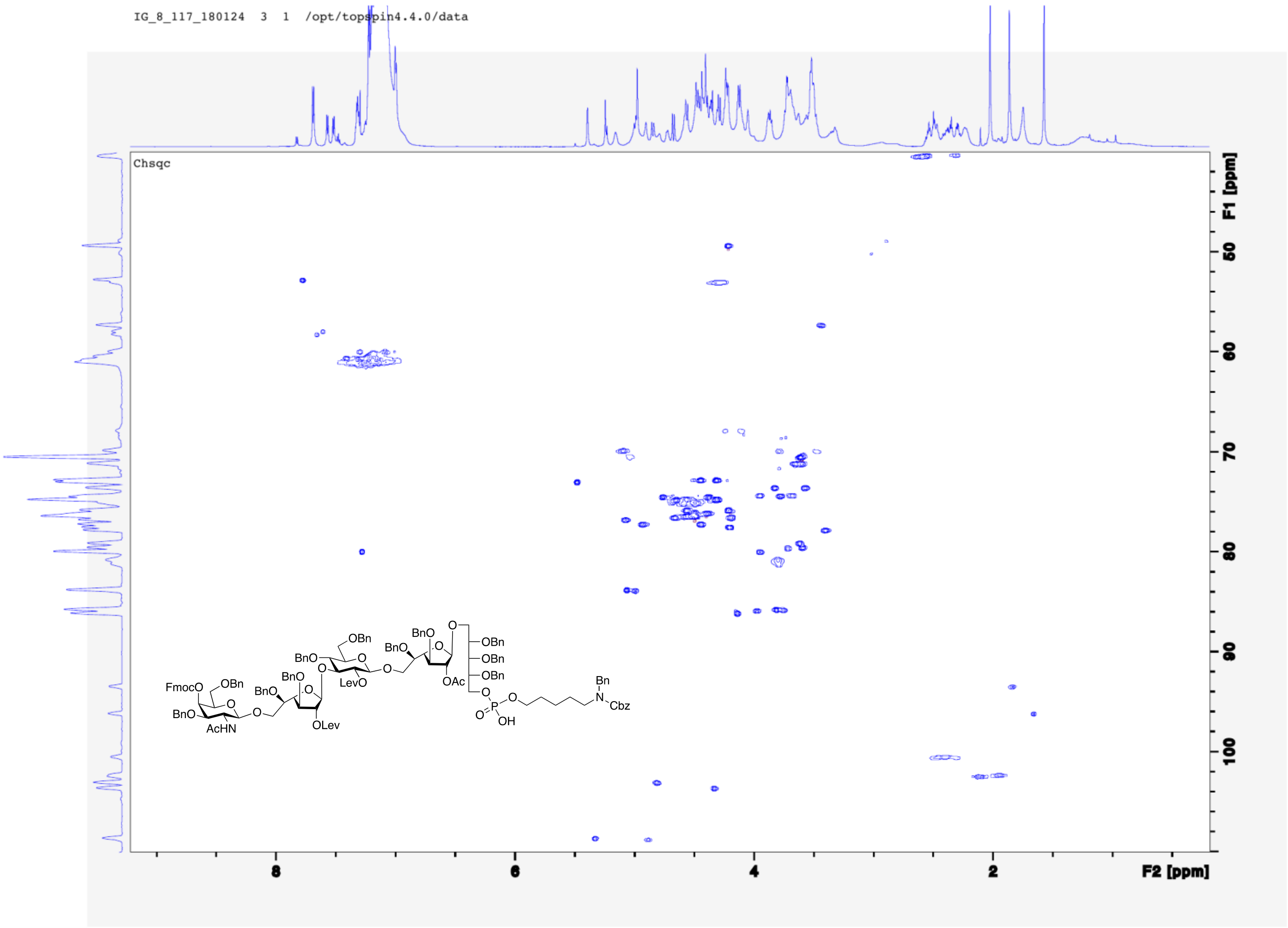
IG_8_117_180124 3 1 /opt/topspin4.4.0/data
Chsqc
F1 [ppm]
F2 [ppm]
FmocO
OBn
BnO
AcHN
OLev
LevO
OAc
OBn
BnO
Bn
N
Cbz
OH

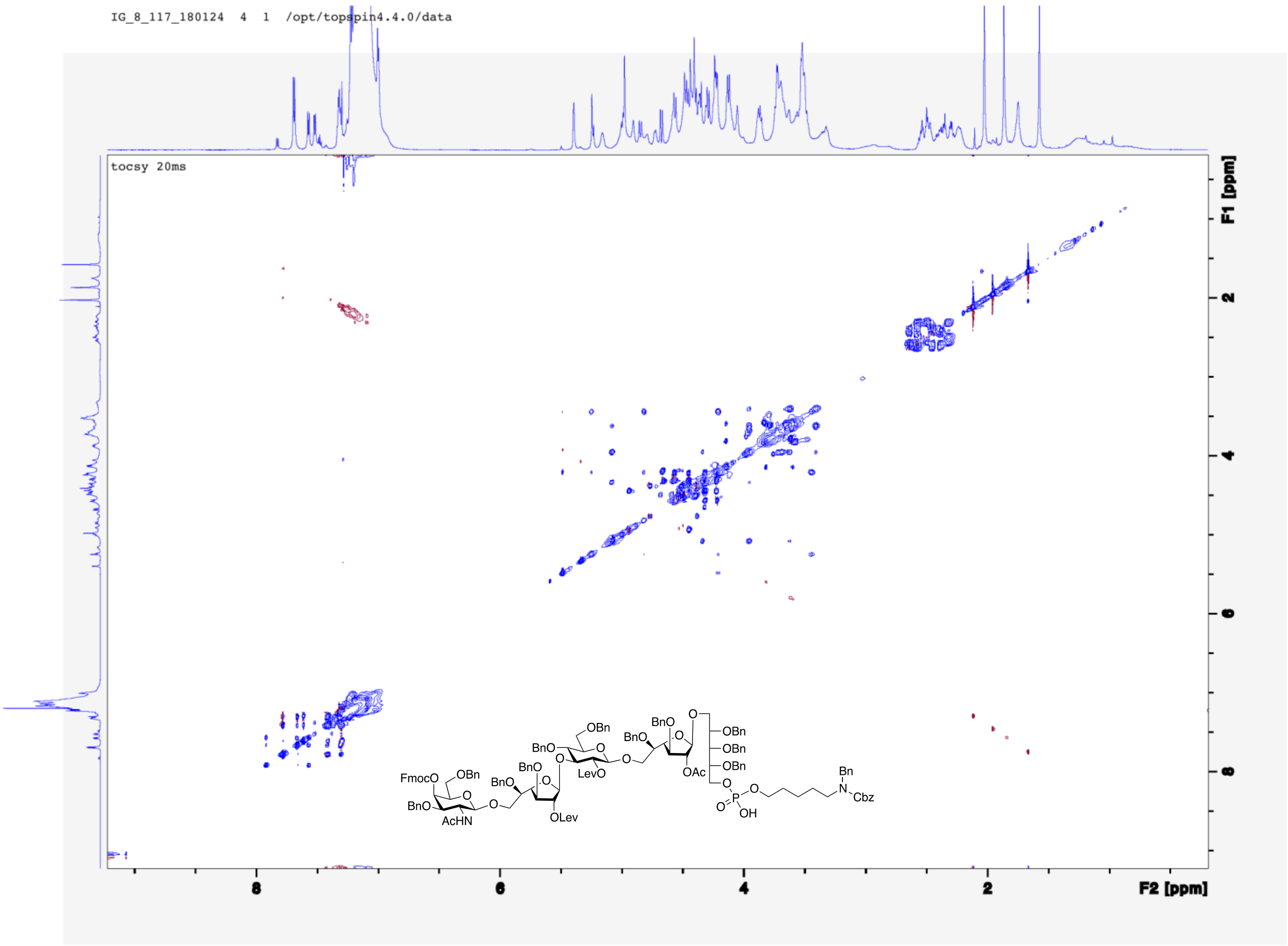
IG_8_117_180124 4 1 /opt/topspin4.4.0/data
tocsy 20ms
F1 [ppm]
F2 [ppm]
FmocO
OBn
BnO
AcHN
BnO
BnO
OLev
BnO
OBn
LevO
BnO
BnO
OAc
OBn
OBn
OBn
OH
Bn
N
Cbz

IG_8_119_180125 1 1 /opt/topspin4.4.0/data

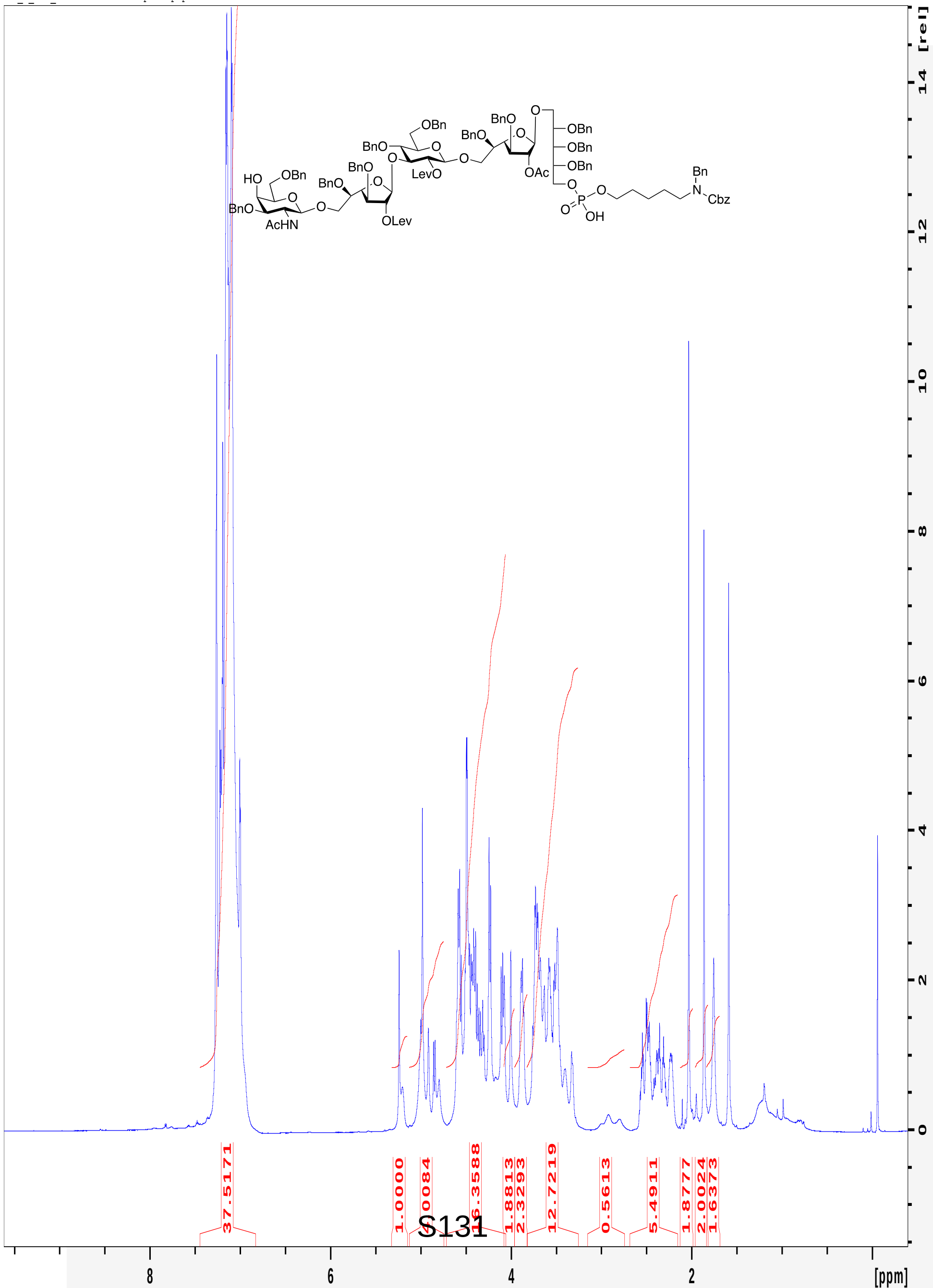

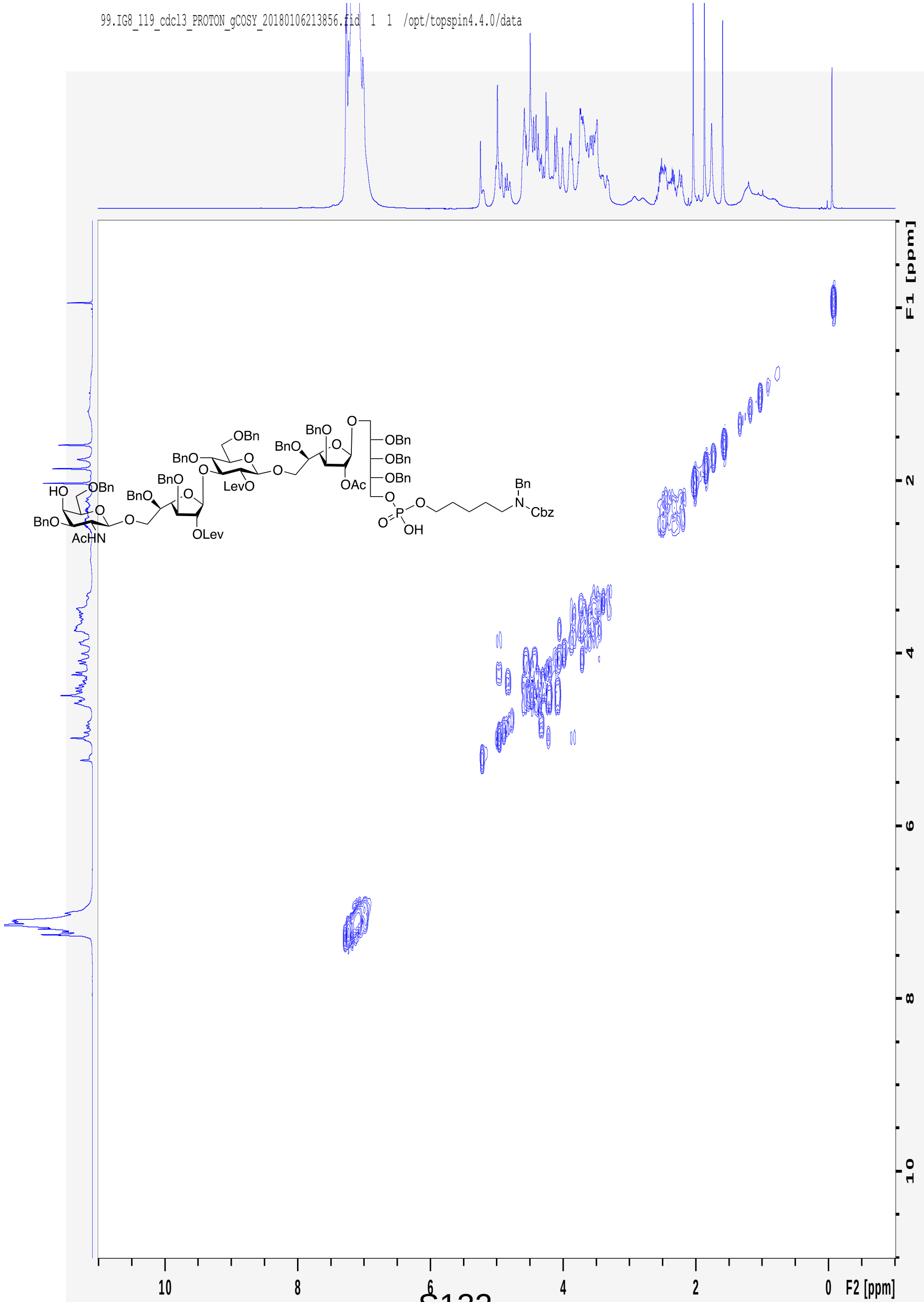
99.IG8_119_cdcl3_PROTON_gCOSY_20180106213856.fid 1 1 /opt/topspin4.4.0/data
F1 [ppm]
F2 [ppm]
HO
OBn
BnO
AcHN
O
BnO
BnO
OLev
BnO
OBn
LevO
BnO
BnO
OAc
OBn
OBn
OBn
O
P
OH
Bn
N
Cbz

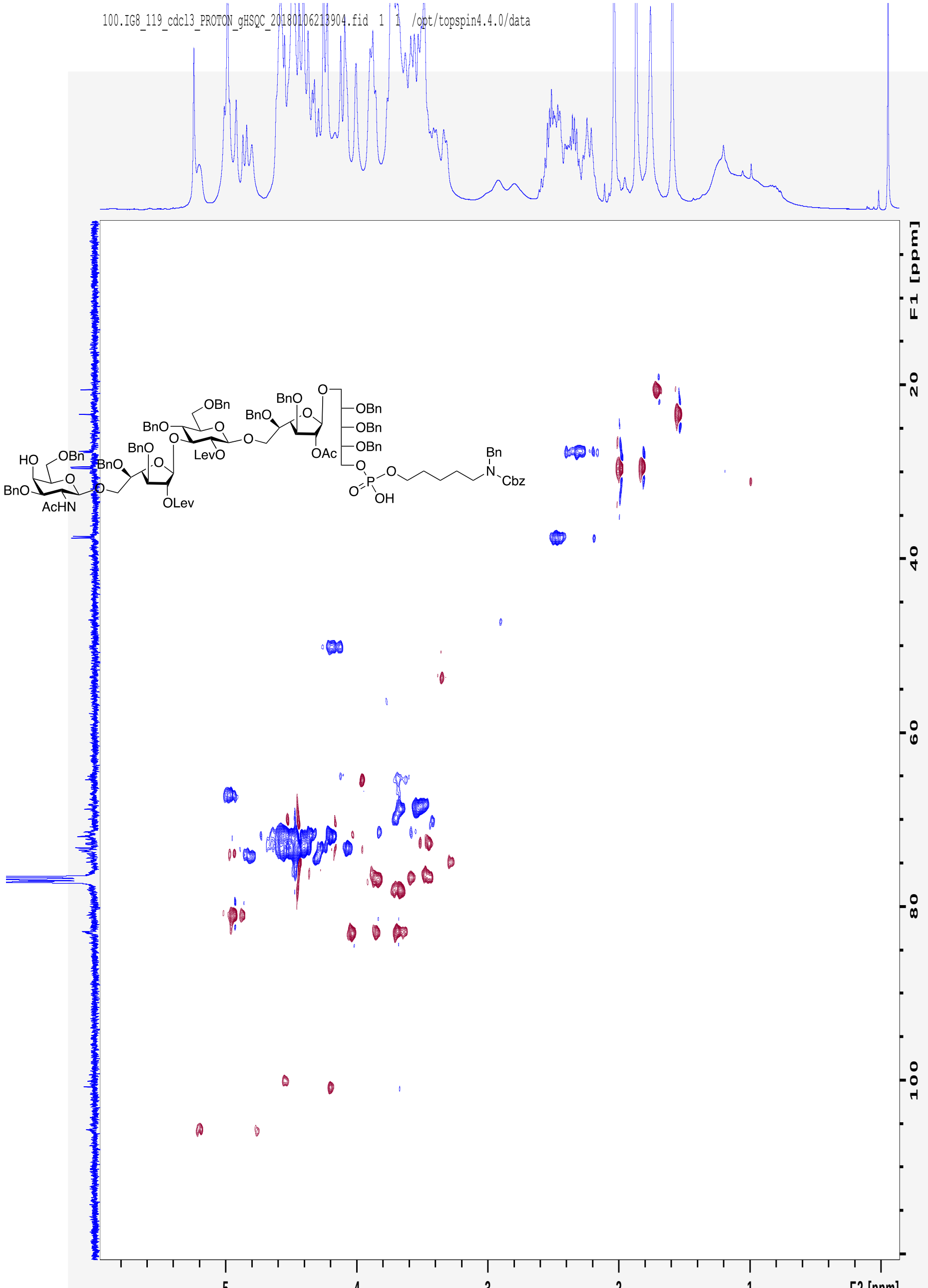
100.IG8_119_cdcl3_PROTON_gHSQC_20180106213904.fid 1 1 /opt/topspin4.4.0/data
F1 [ppm]
F2 [ppm]

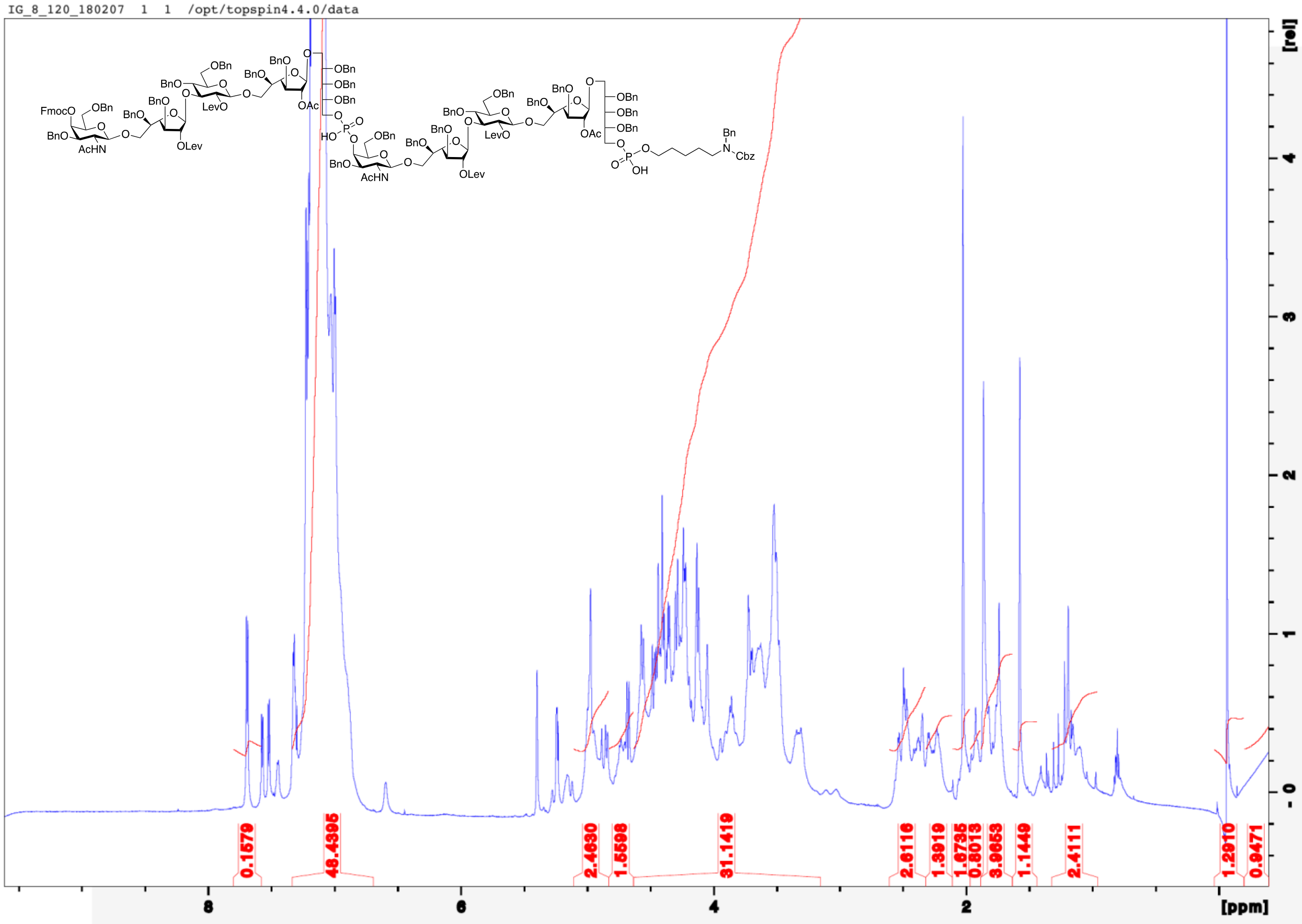
IG_8_120_180207 1 1 /opt/topspin4.4.0/data
[rel]
[ppm]
0.1579
48.4395
2.4630
1.5598
31.1419
2.6116
1.3919
1.6795
0.8013
3.9653
1.1449
2.4111
1.2910
0.9471

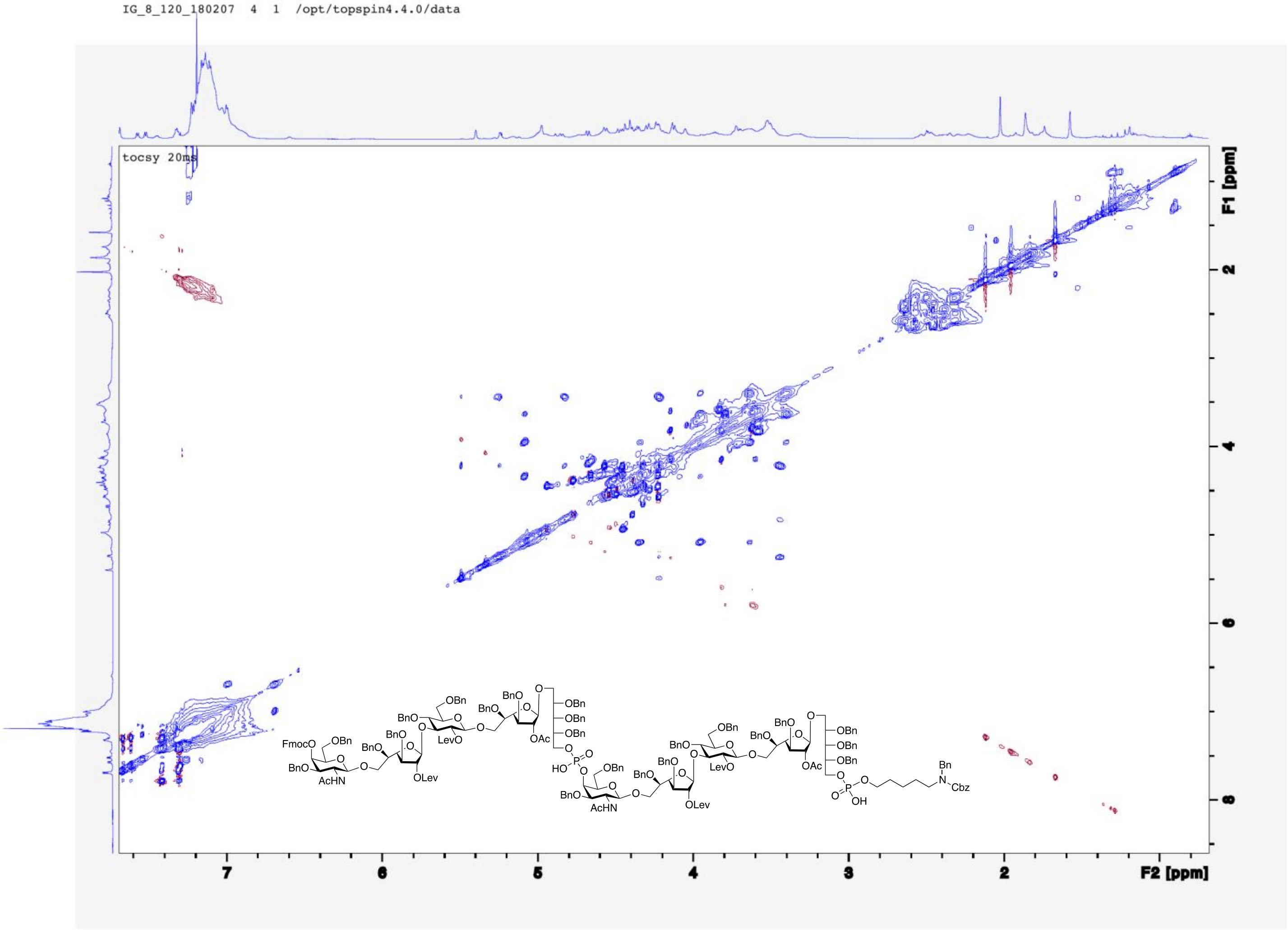

IG_8_120_180207 4 1 /opt/topspin4.4.0/data
tocsy 20ms
F1 [ppm]
F2 [ppm]
7
6
5
4
3
2
FmocO
OBn
BnO
AcHN
BnO
LevO
OLev
OAc
HO
OH
Bn
N
Cbz

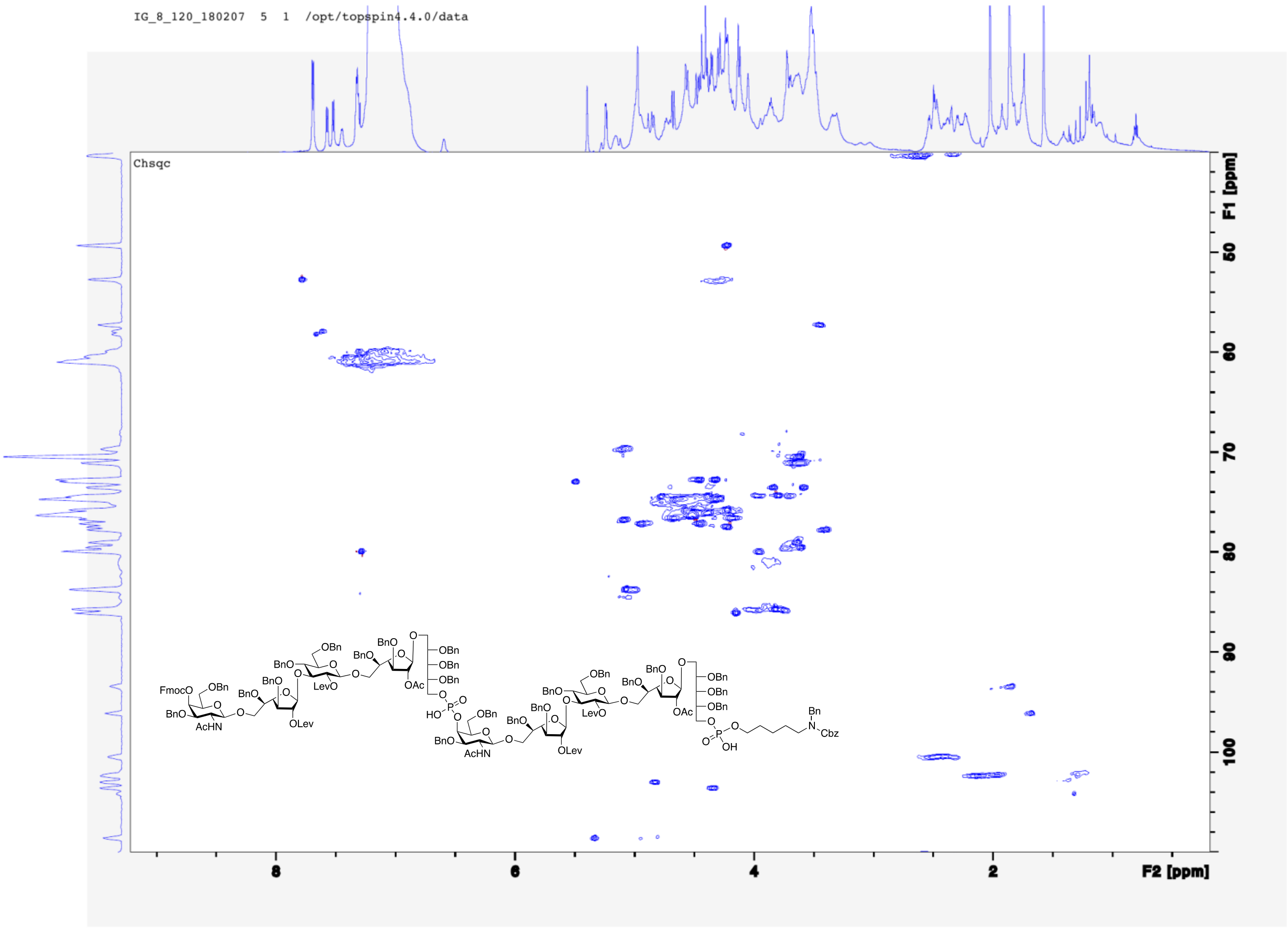

IG_8_120_180207 5 1 /opt/topspin4.4.0/data
Chsqc
F1 [ppm]
F2 [ppm]
FmocO
OBn
BnO
AcHN
OLev
LevO
OAc
OBn
HO
Bn
Cbz
OH

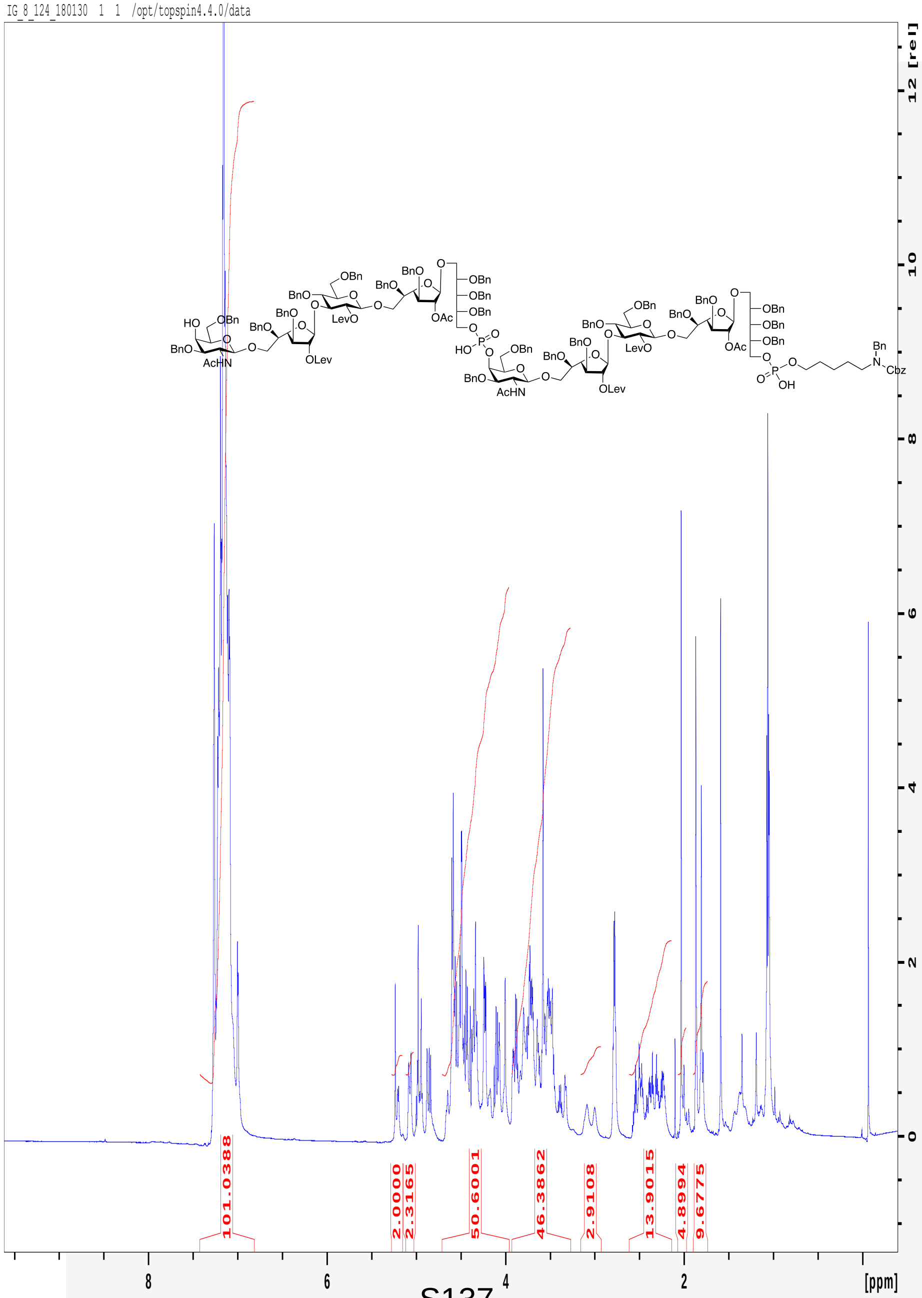
IG_8_124_180130 1 1 /opt/topspin4.4.0/data
101.0388
2.0000
2.3165
50.6001
46.3862
2.9108
13.9015
4.8994
9.6775
[ppm]
[rel]

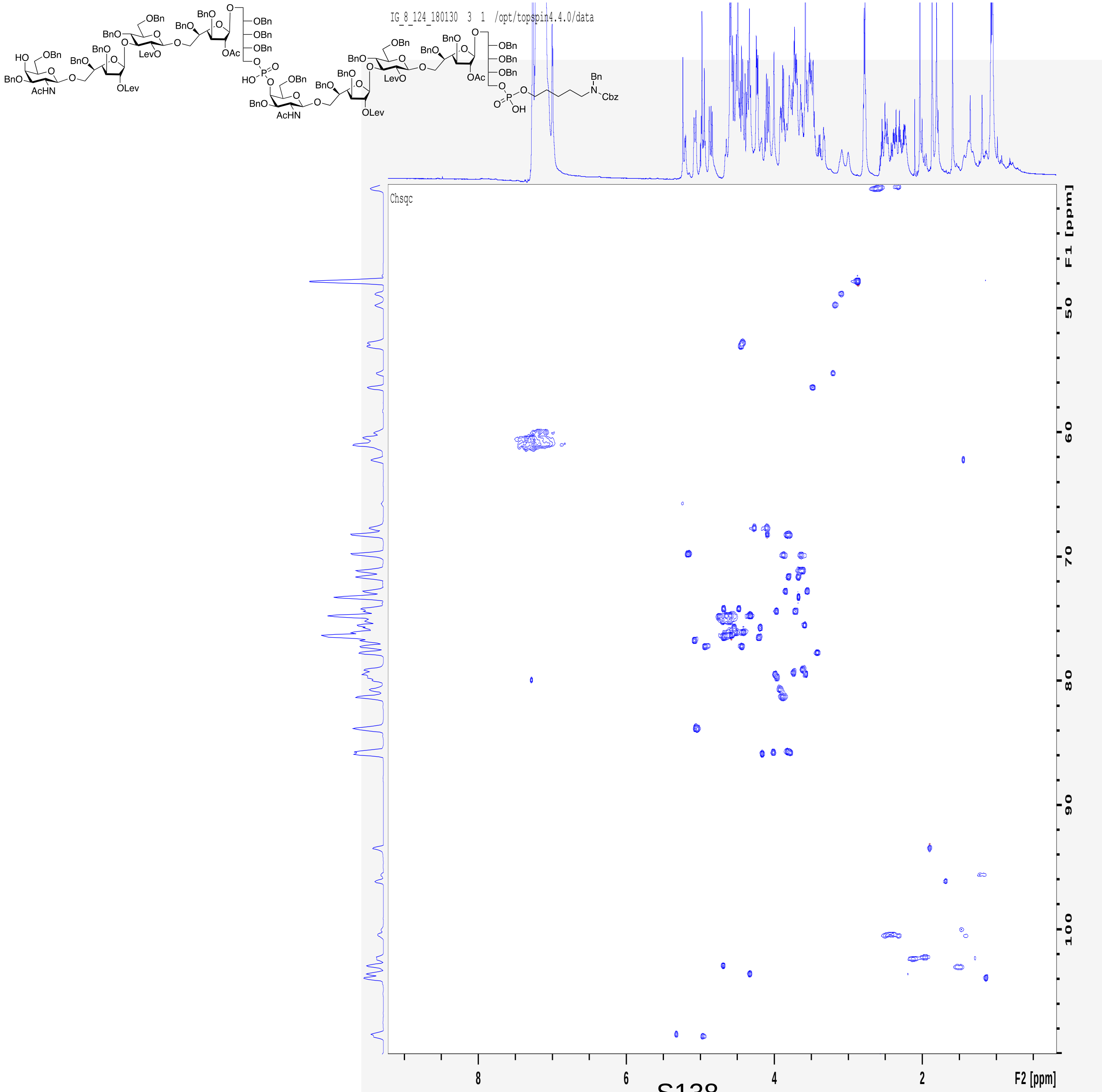
IG_8_124_180130 3 1 /opt/topspin4.4.0/data
Chsqc
F1 [ppm]
F2 [ppm]

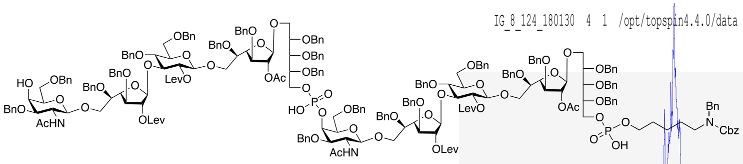

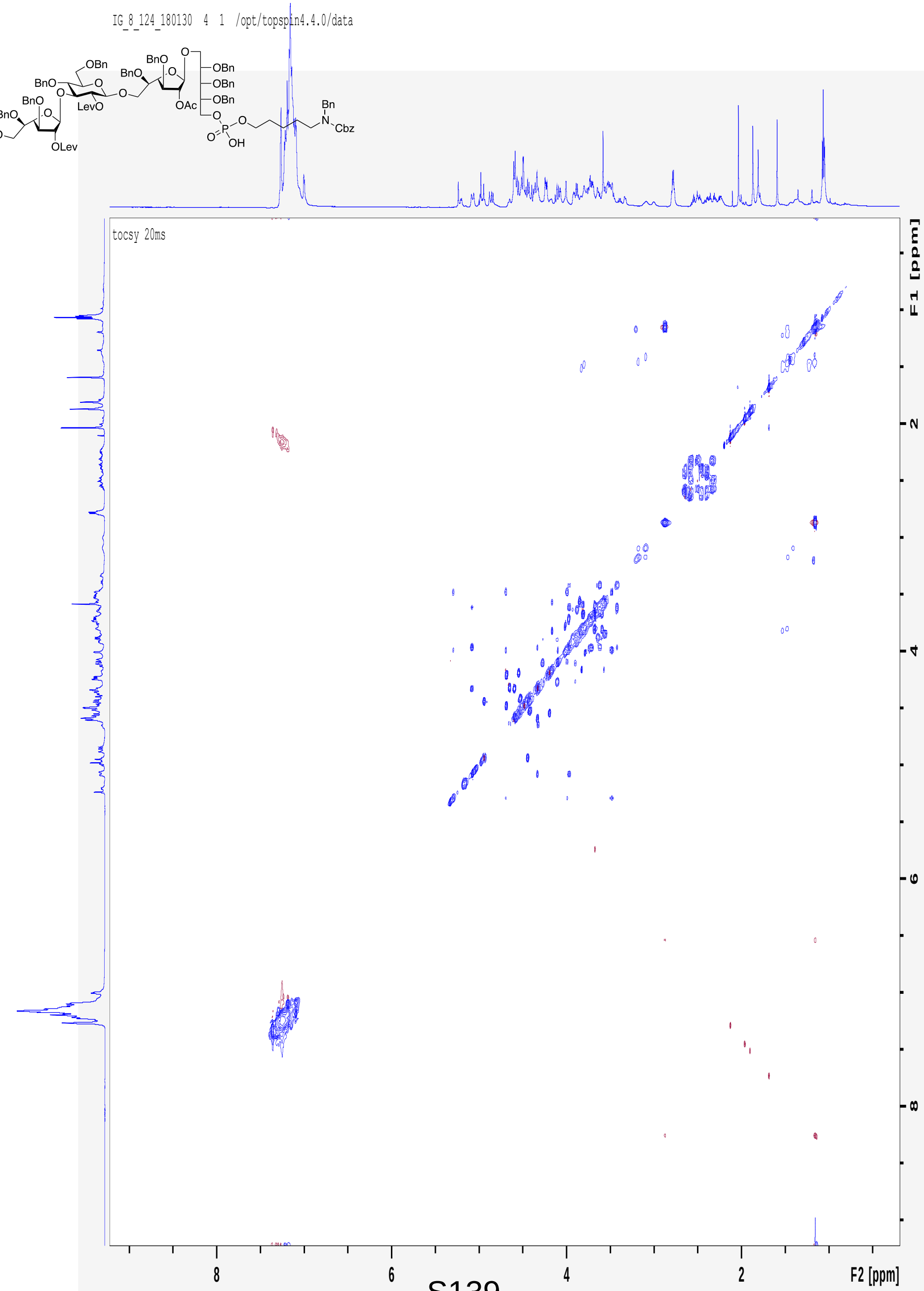
IG_8_124_180130 4 1 /opt/topspin4.4.0/data
tocsy 20ms
F1 [ppm]
F2 [ppm]
8
6
4
2

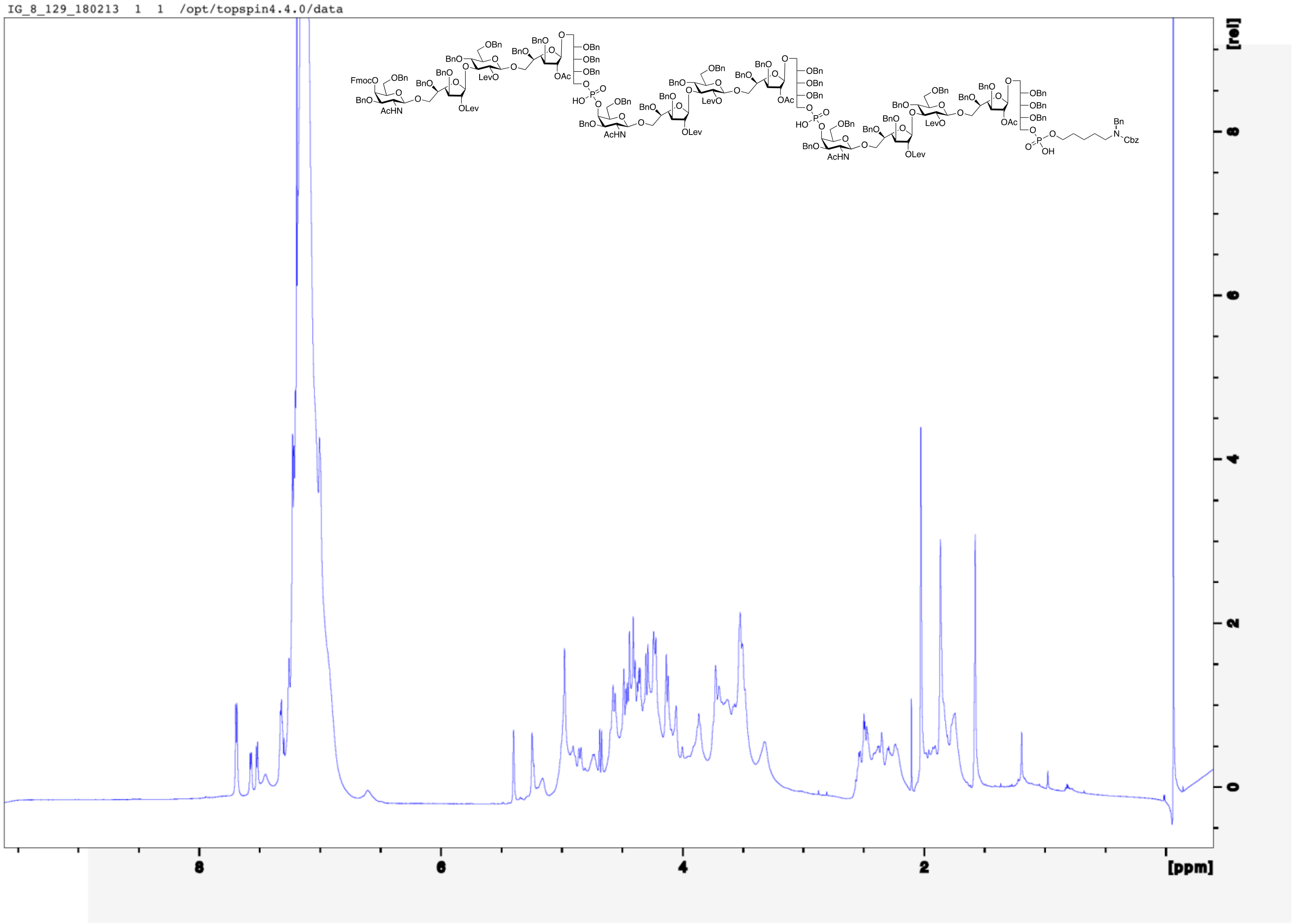
IG_8_129_180213 1 1 /opt/topspin4.4.0/data
[rel]
[ppm]

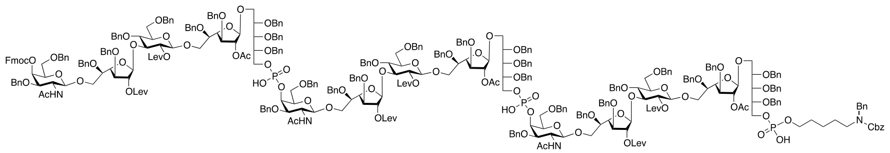

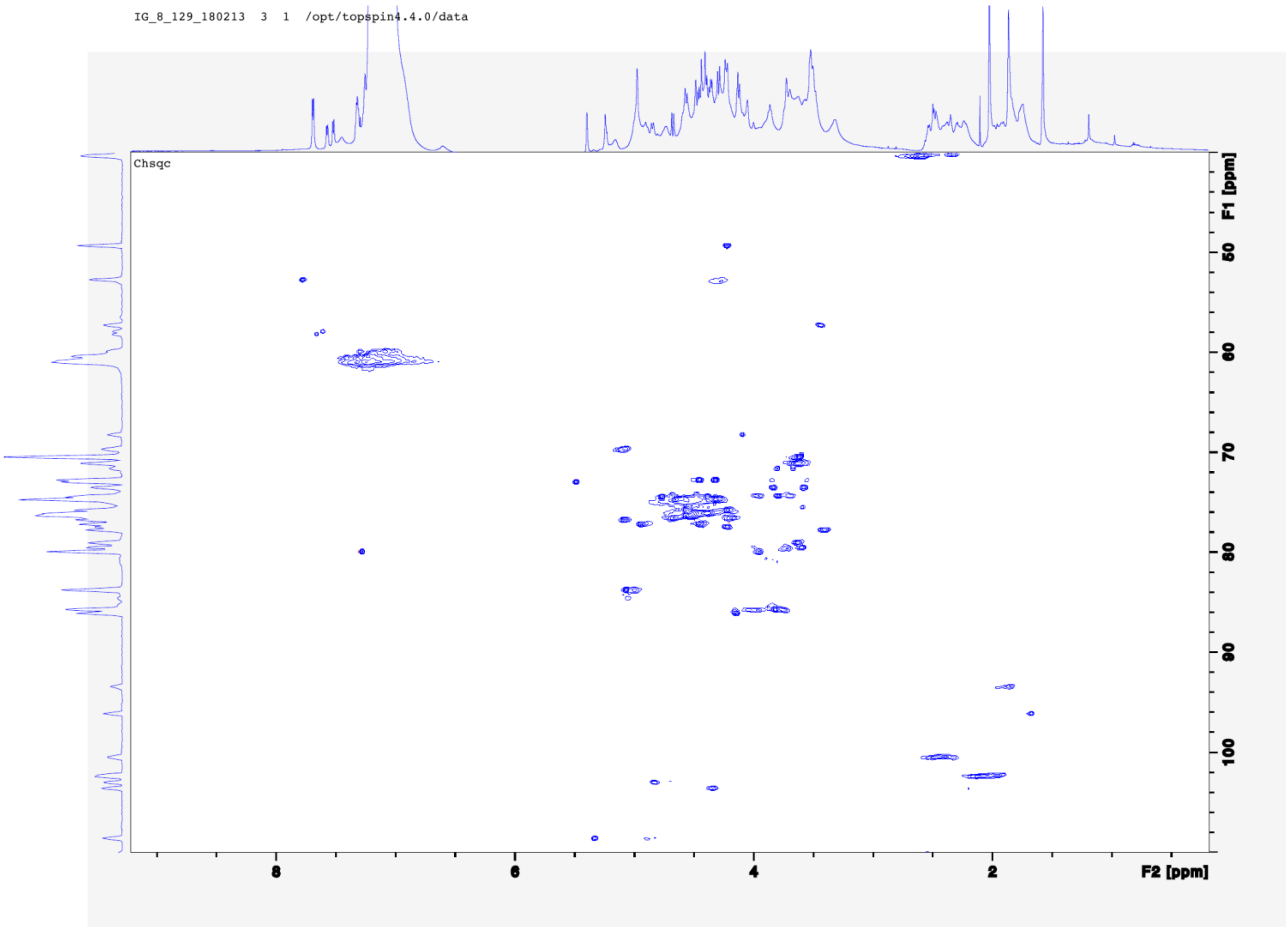
IG_8_129_180213 3 1 /opt/topspin4.4.0/data
Chsqc
F1 [ppm]
F2 [ppm]

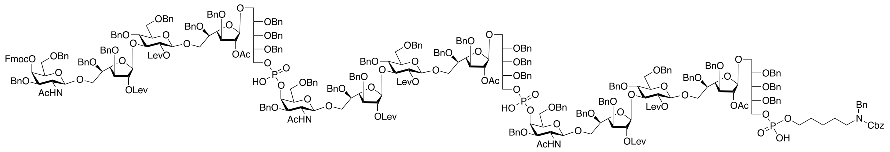

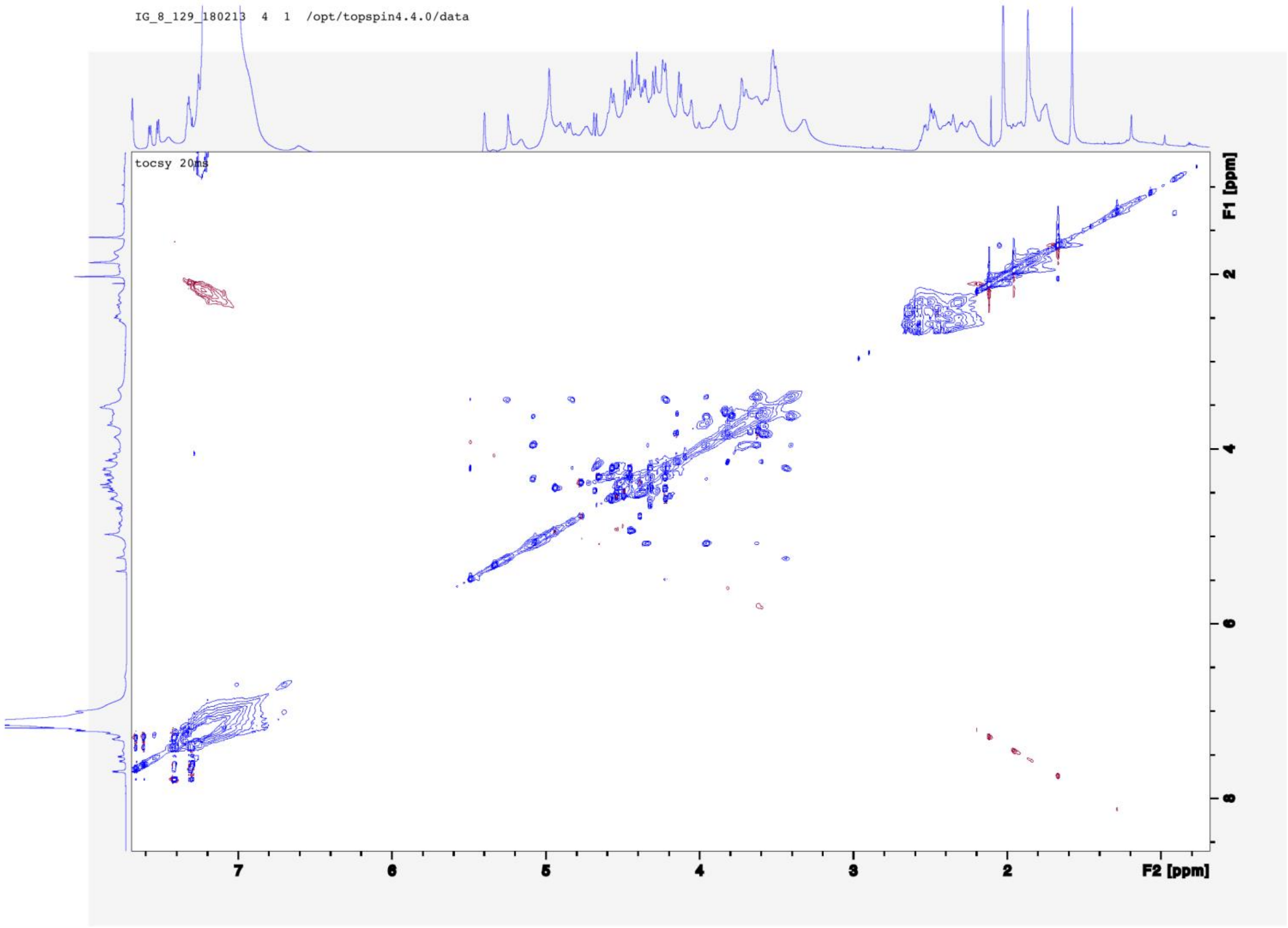

IG_8_129_180213 4 1 /opt/topspin4.4.0/data
tocsy 20ms
F1 [ppm]
F2 [ppm]

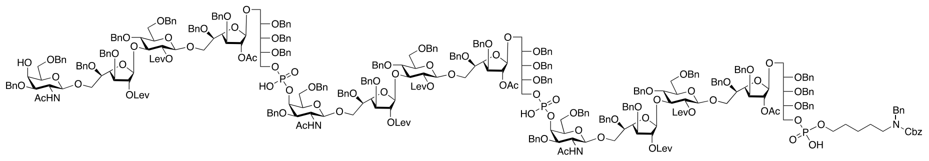

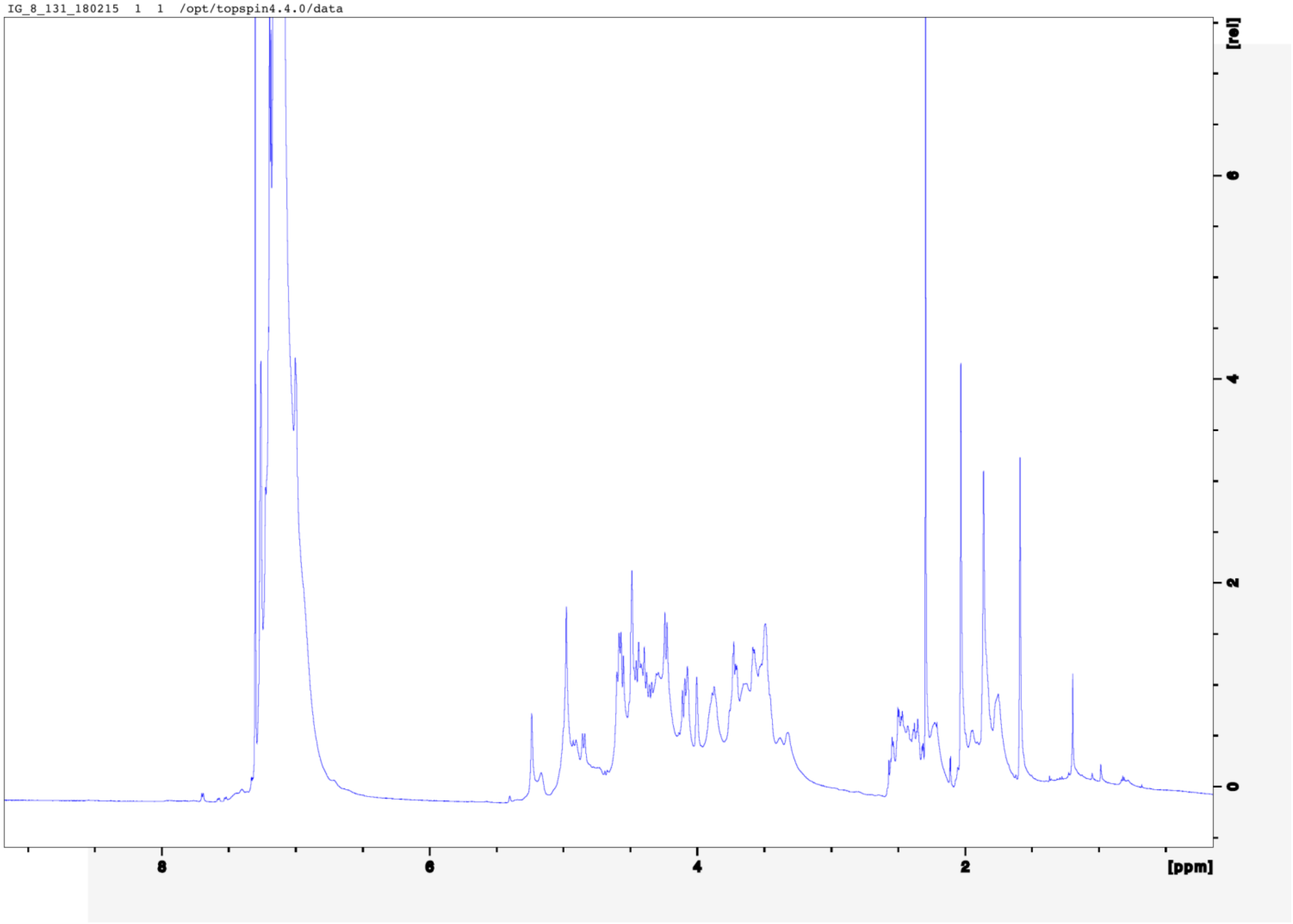
IG_8_131_180215 1 1 /opt/topspin4.4.0/data
[rel]
6
4
2
0
8
6
4
2
[ppm]

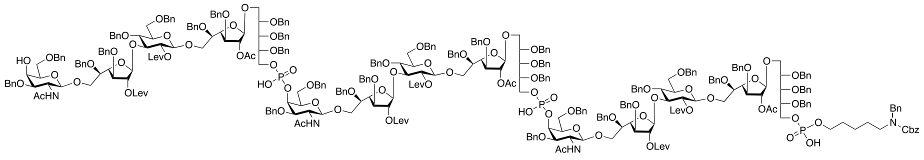

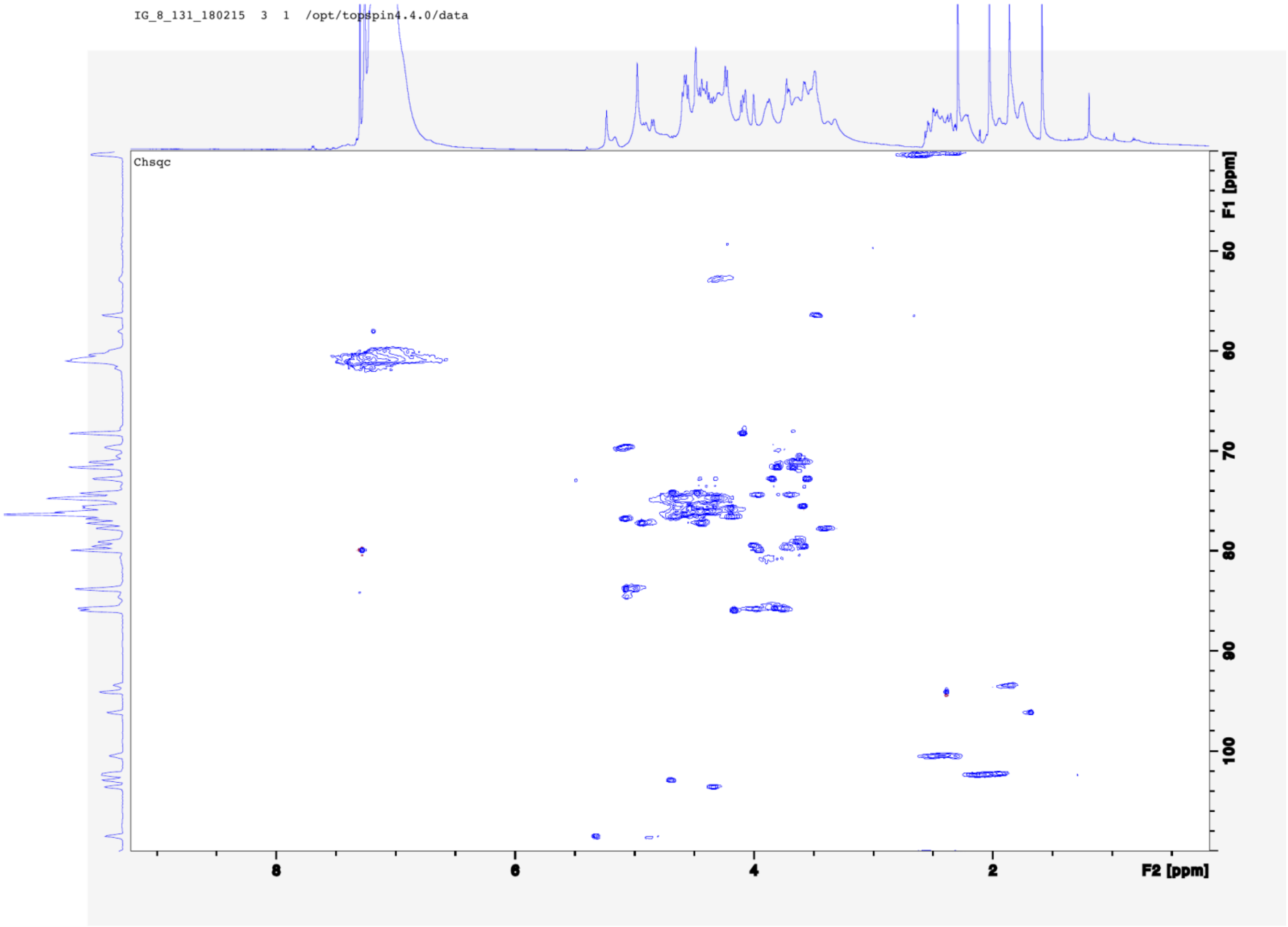
IG_8_131_180215 3 1 /opt/topspin4.4.0/data
Chsqc
F1 [ppm]
F2 [ppm]

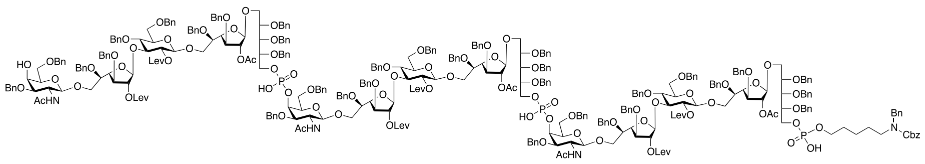

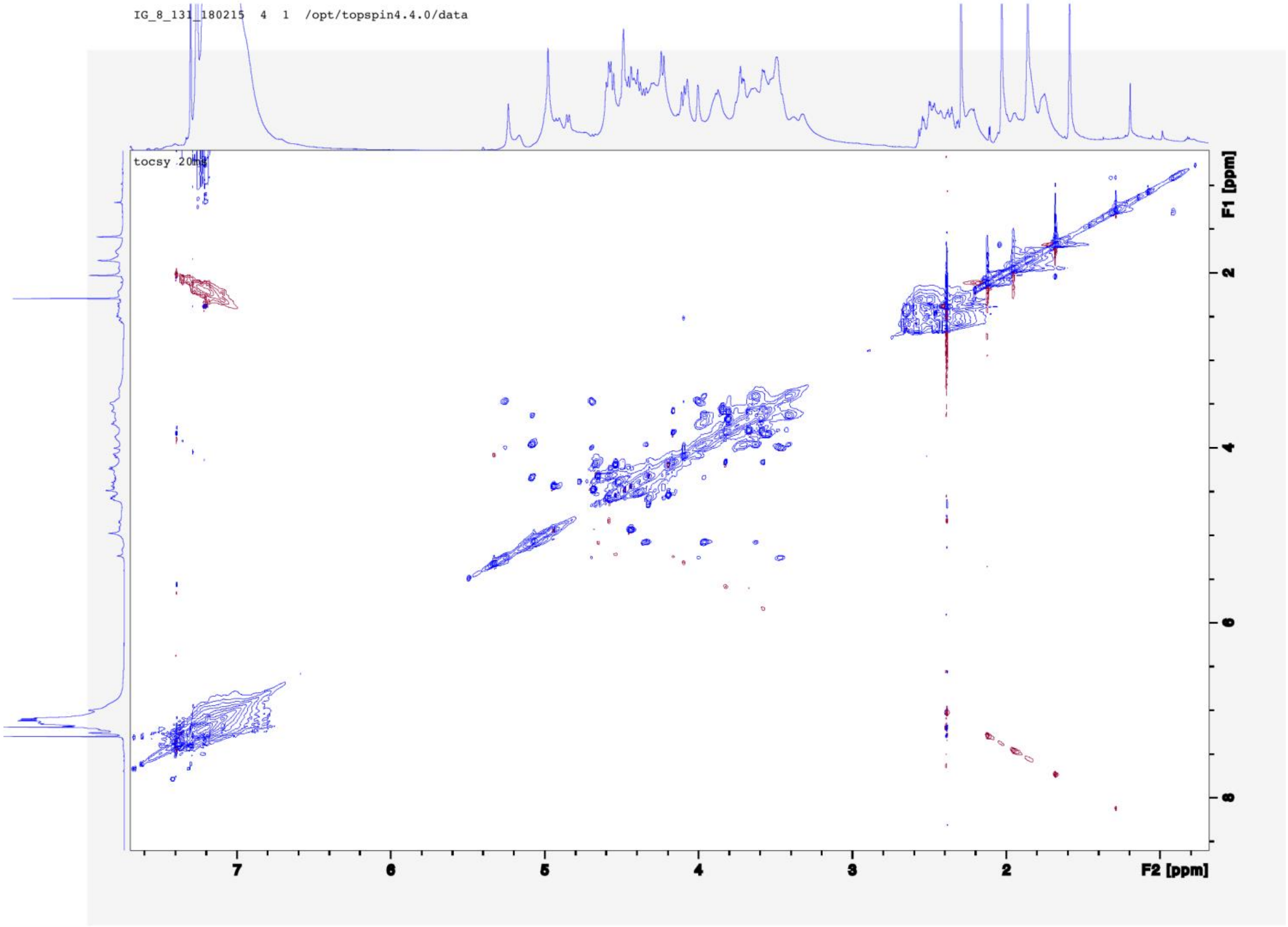
IG_8_131_180215 4 1 /opt/topspin4.4.0/data
tocsy 20mg
F1 [ppm]
F2 [ppm]

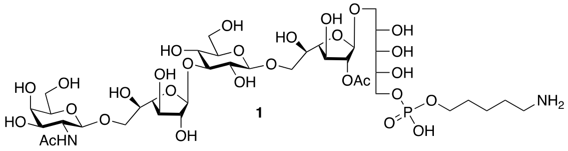
HO
OH
HO
AcHN
O
HO
OH
OH
HO
OH
HO
OAc
OH
OH
OH
NH2
O
P
OH


1

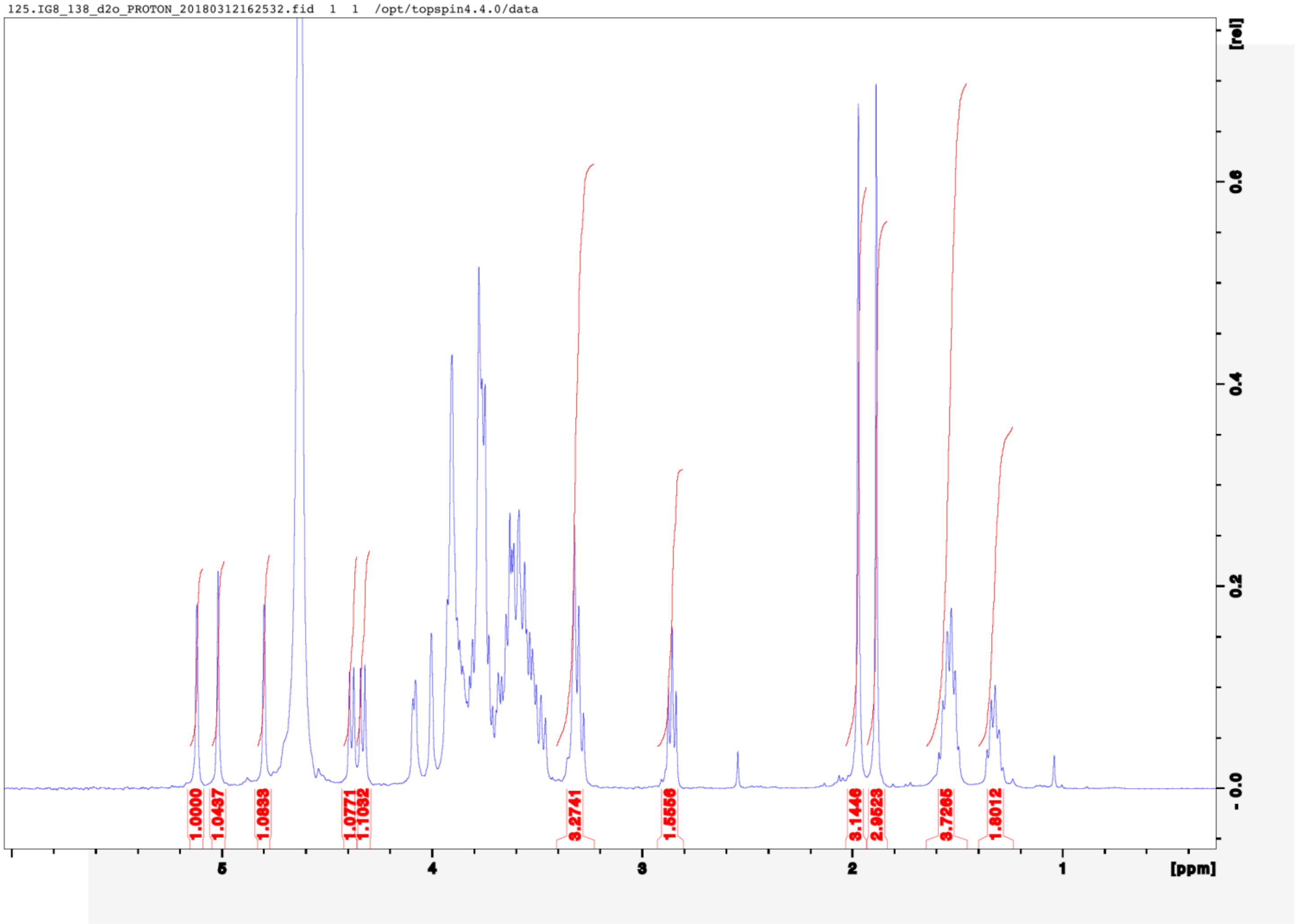
125.IG8_138_d2o_PROTON_20180312162532.fid 1 1 /opt/topspin4.4.0/data
[rel]
0.6
0.4
0.2
0.0
1.0000
1.0437
1.0833
1.0771
1.1032
3.2741
1.5556
3.1446
2.9523
3.7265
1.8012
5
4
3
2
1
[ppm]

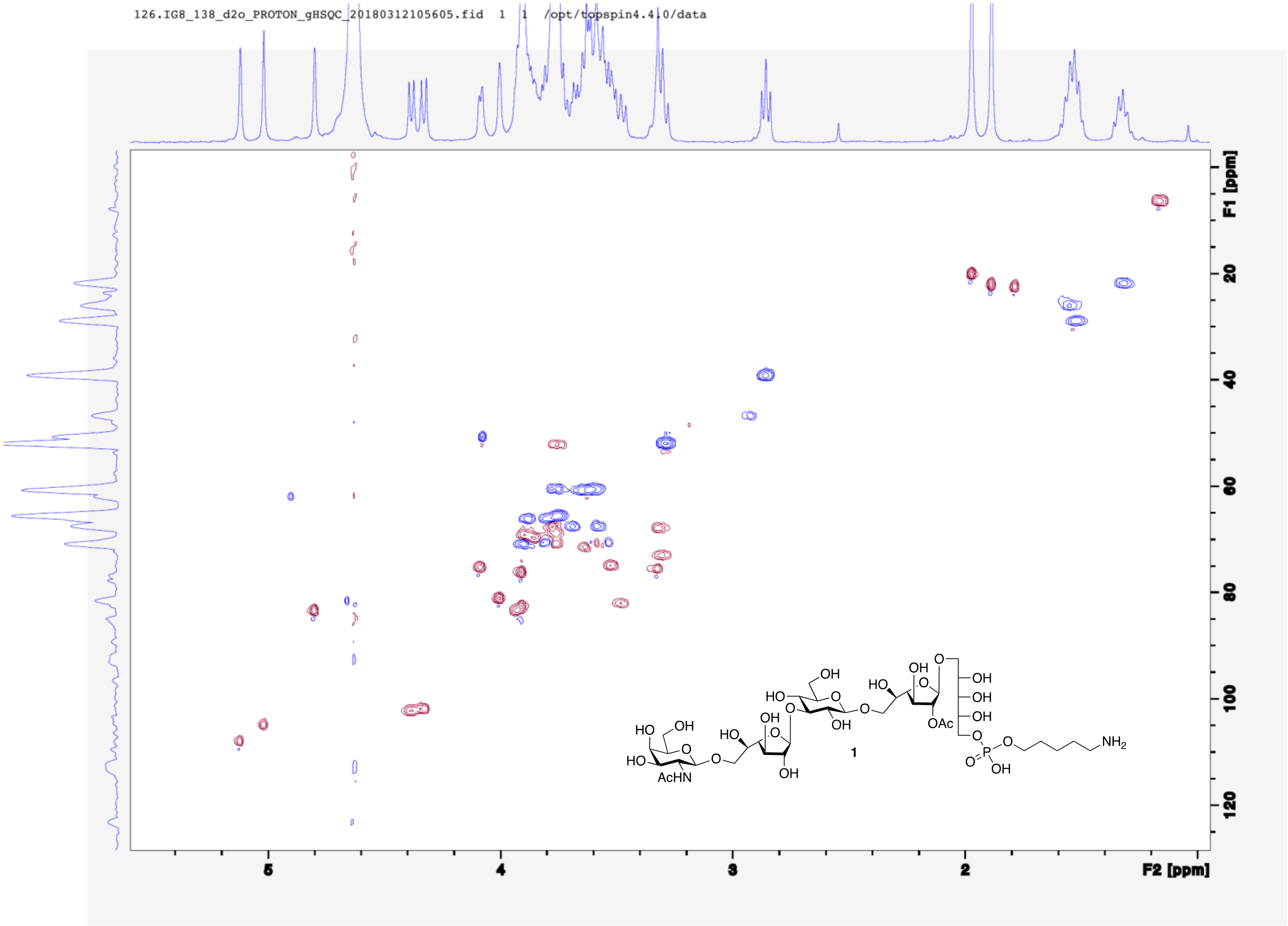

126.IG8_138_d2o_PROTON_gHSQC_20180312105605.fid 1 1 /opt/topspin4.4.0/data
F1 [ppm]
F2 [ppm]
1

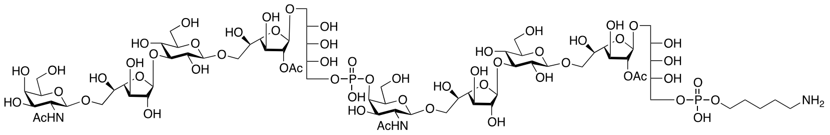

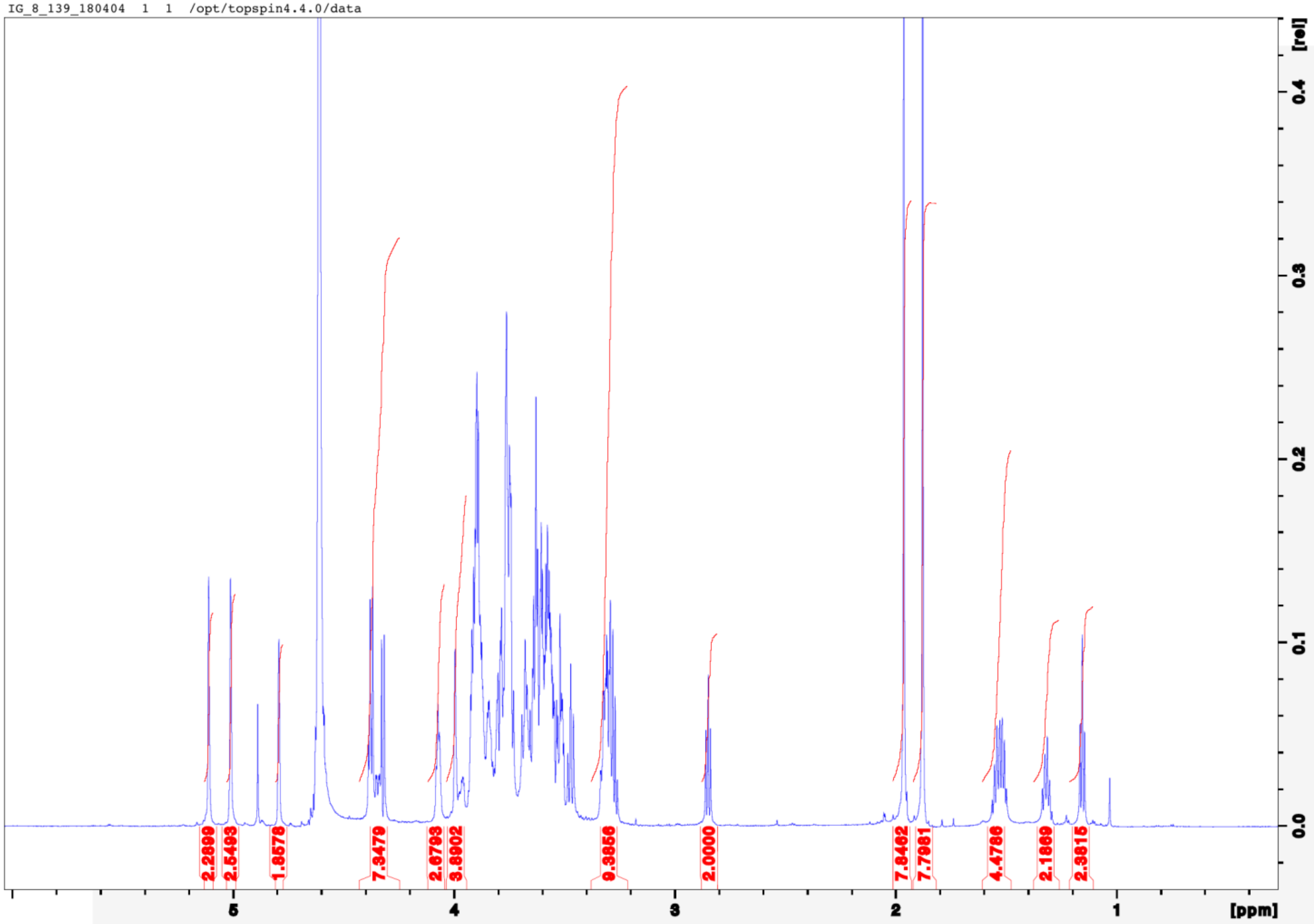
IG_8_139_180404 1 1 /opt/topspin4.4.0/data
2.2899
2.5493
1.8578
7.3479
2.6793
3.8902
9.3856
2.0000
7.8462
7.7961
4.4786
2.1869
2.3815
[rel]
[ppm]

IG_8_139_180313 2 1 /opt/topspin4.4.0/data

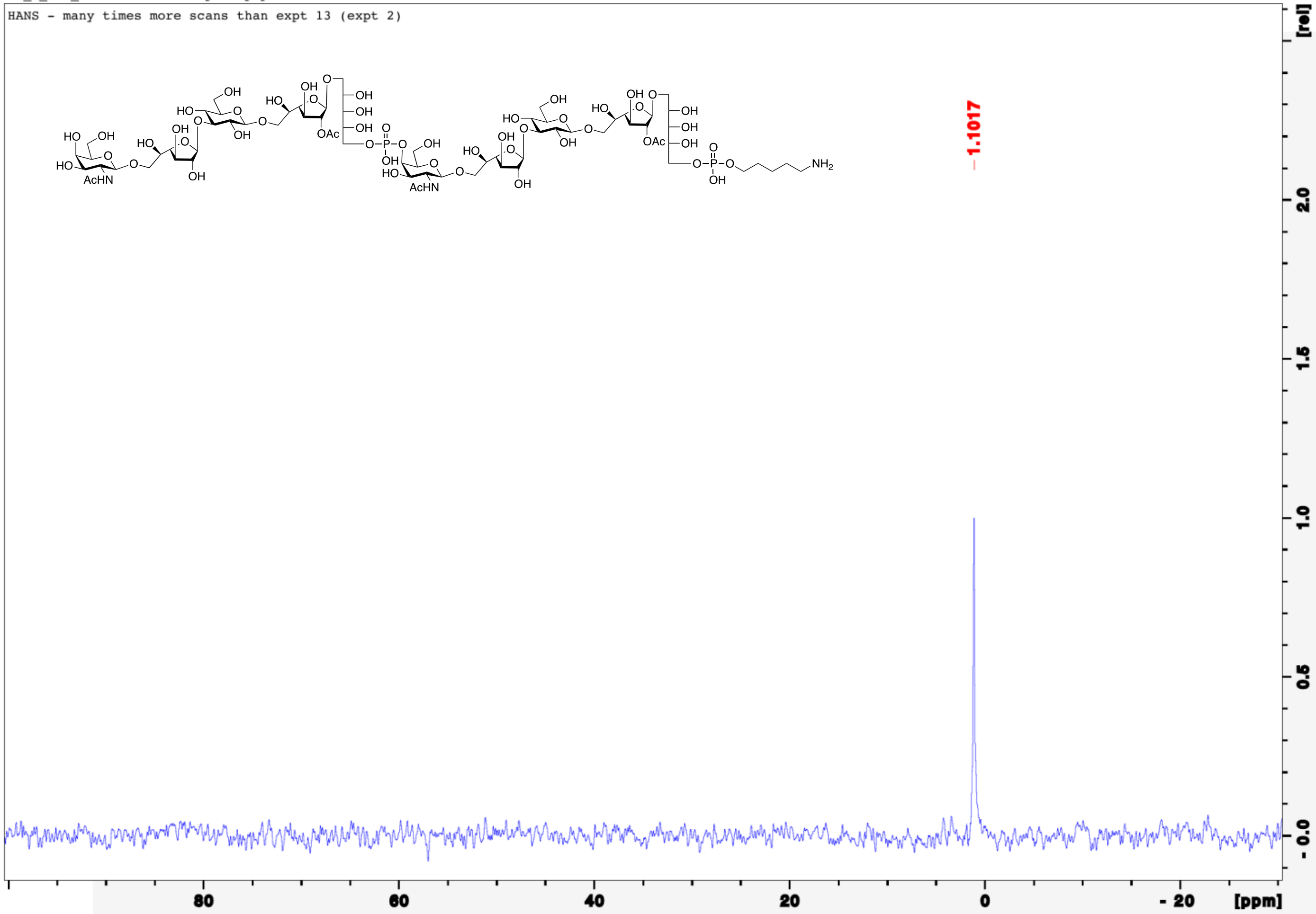

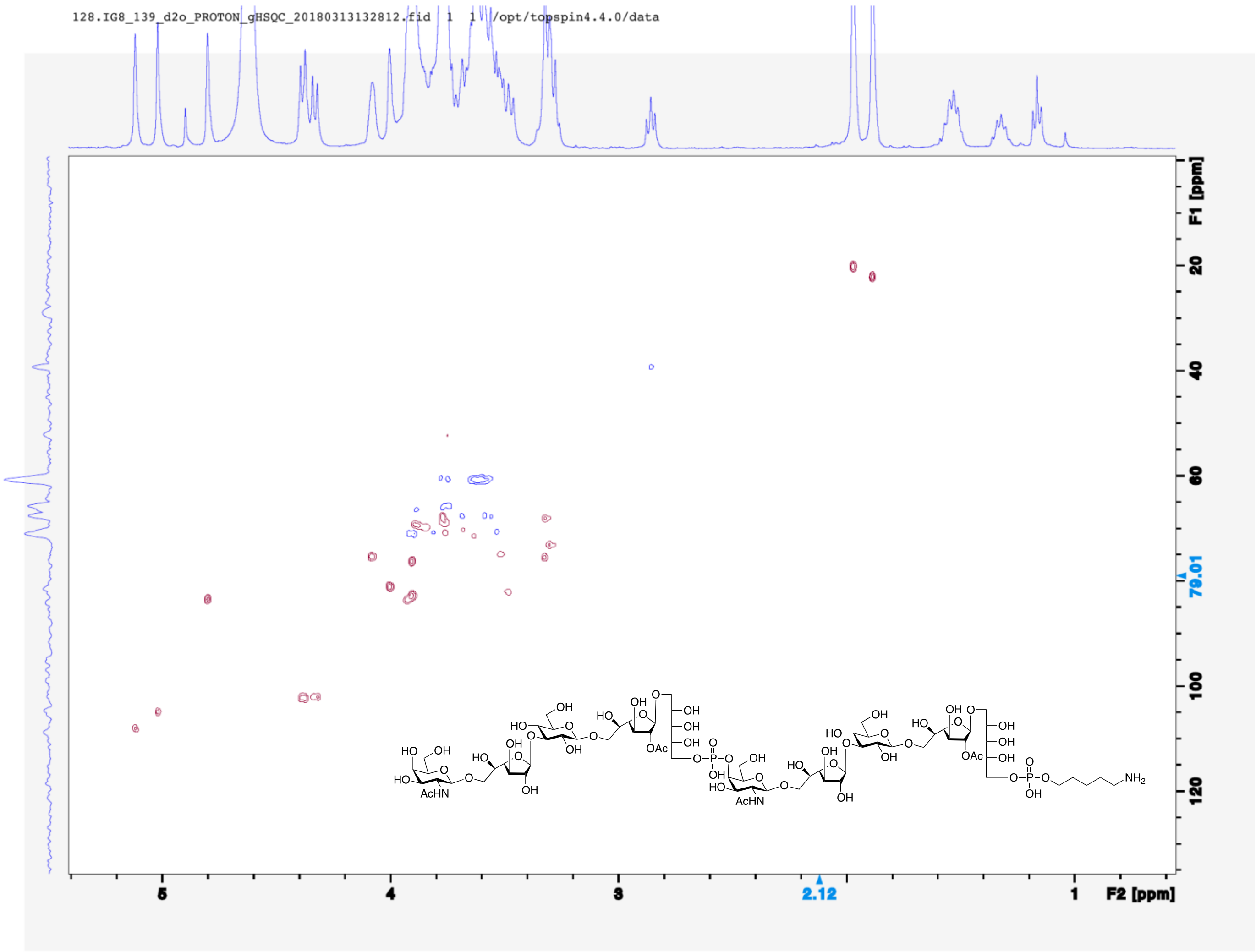
128.IG8_139_d2o_PROTON_gHSQC_20180313132812.fid 1 1 /opt/topspin4.4.0/data
F1 [ppm]
20
40
60
79.01
100
120
5
4
3
2.12
1
F2 [ppm]
AcHN
OAc
NH2

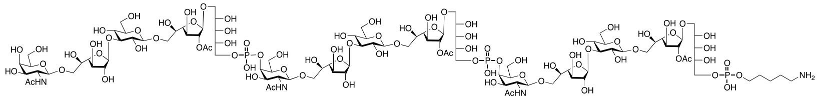

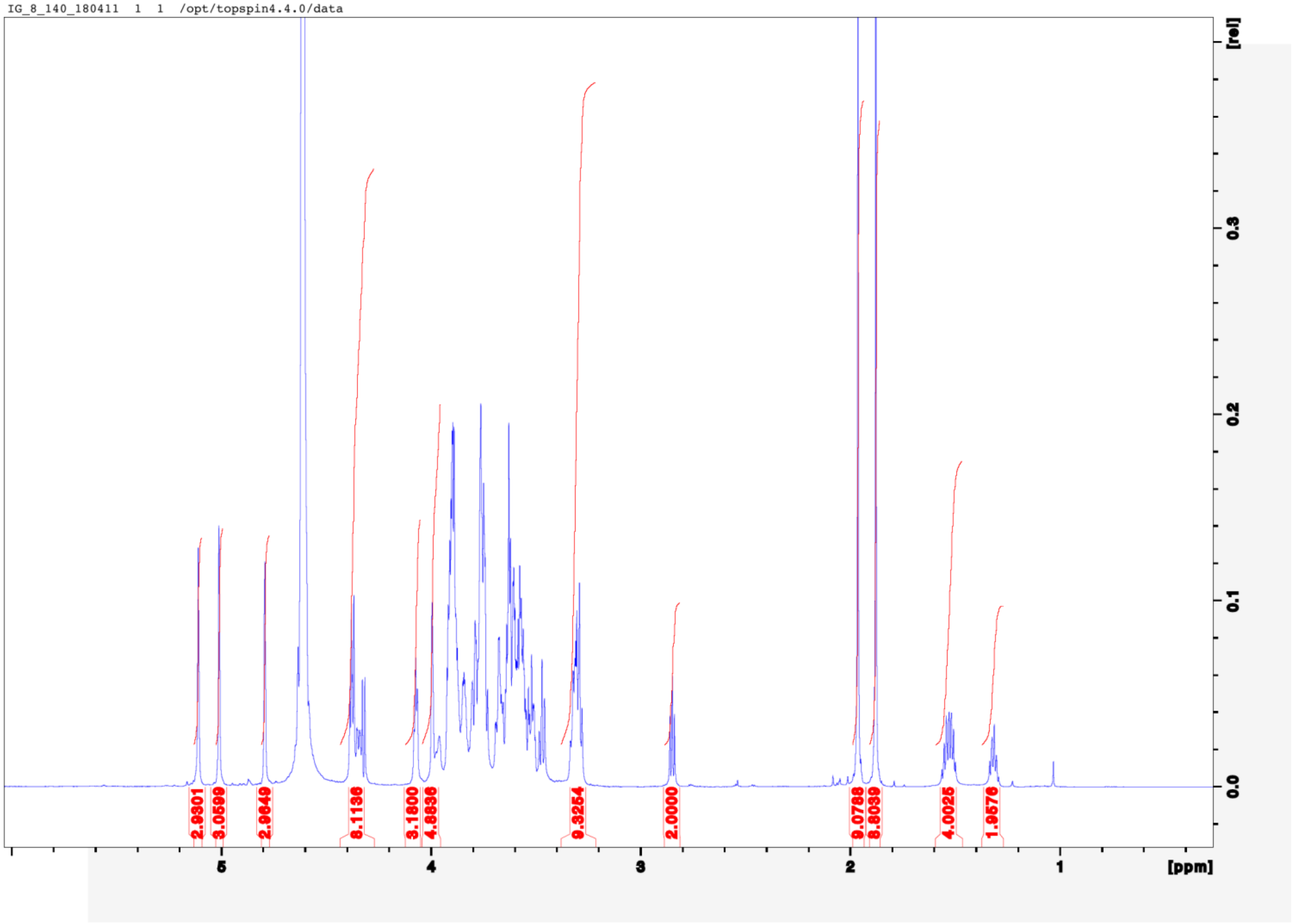

IG_8_140_180411 1 1 /opt/topspin4.4.0/data
2.9301
3.0599
2.9649
6.1136
3.1800
4.8836
9.3254
2.0000
9.0788
8.8039
4.0025
1.9576
5
4
3
2
1
[ppm]
0.0
0.1
0.2
0.3
[rel]

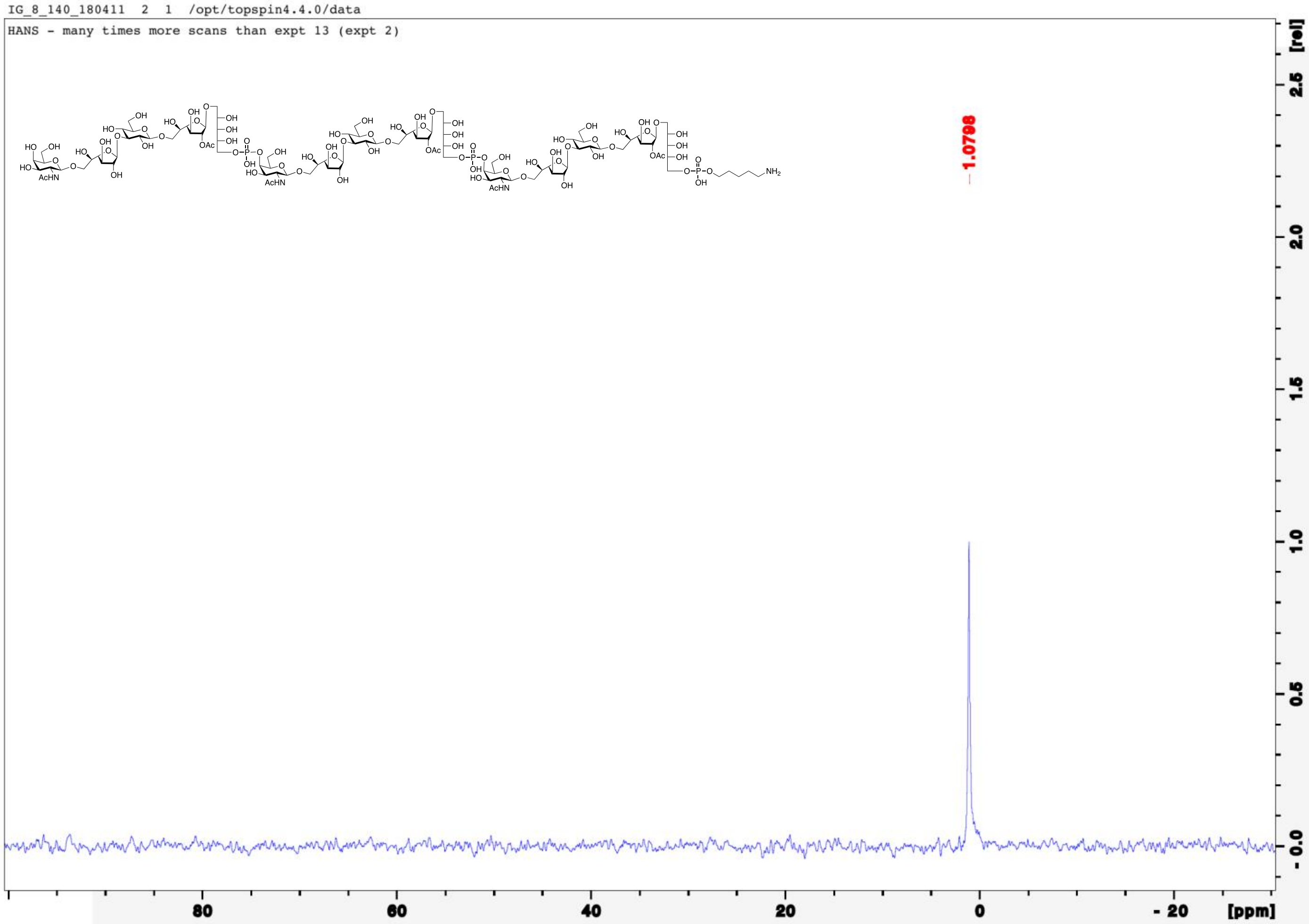
IG_8_140_180411 2 1 /opt/topspin4.4.0/data
HANS - many times more scans than expt 13 (expt 2)
-1.0798
80
60
40
20
0
- 20
[ppm]
[rel]
2.5
2.0
1.5
1.0
0.5
- 0.0

IG_8_140_180411 4 1 /opt/topspin4.4.0/data

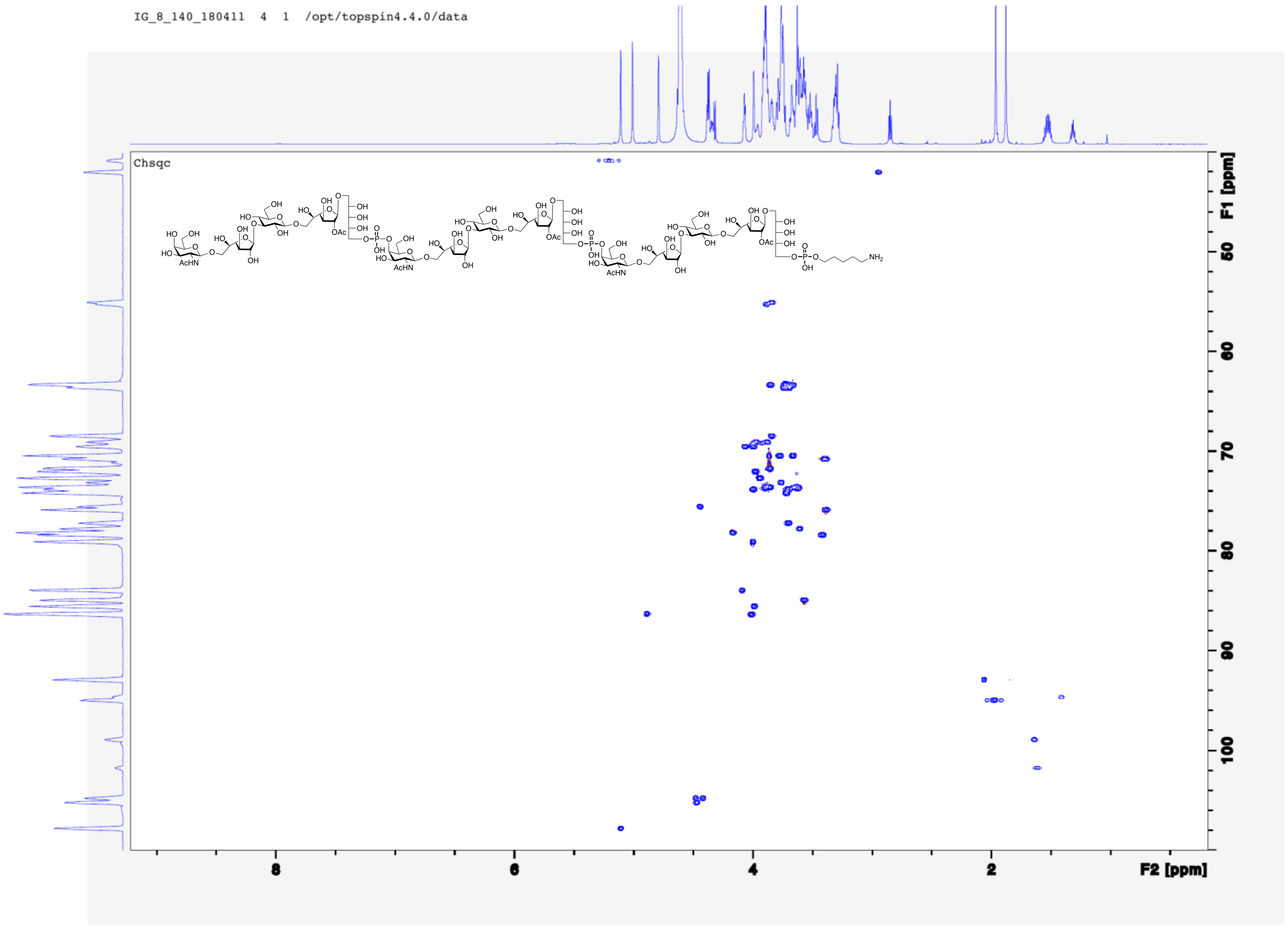

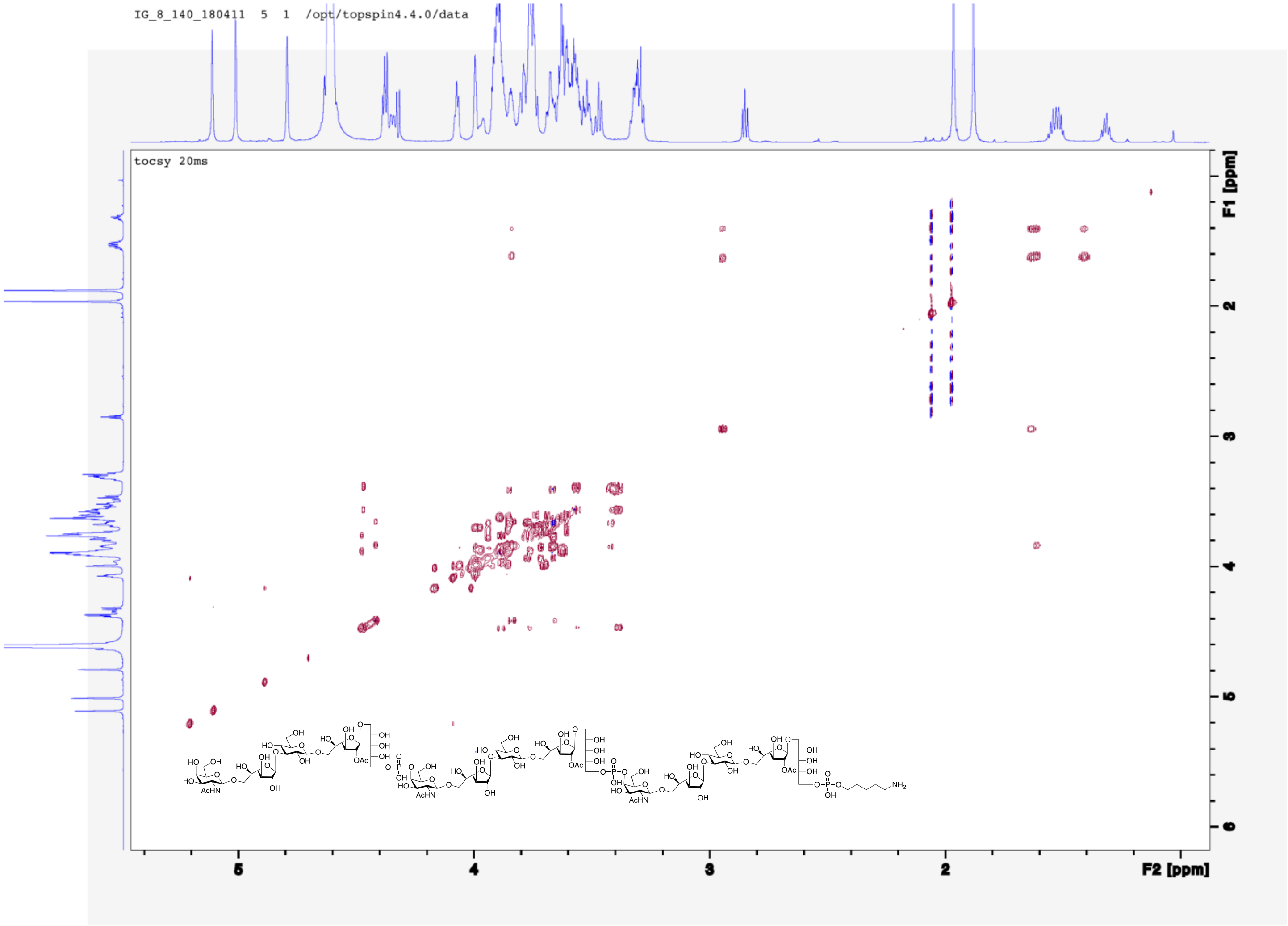
IG_8_140_180411 5 1 /opt/topspin4.4.0/data
tocsy 20ms
F1 [ppm]
F2 [ppm]
